# Multiplexed mammalian gene control via modular splicing riboswitches

**DOI:** 10.64898/2026.10.16.759771

**Authors:** Jiacheng Jiang, Tonghui Ma

**Affiliations:** School of Medicine, Nanjing University of Chinese Medicine, 138 Xianlin Rd, Nanjing, 210023, China

## Abstract

Precise, multiplexed gene regulation in mammalian cells remains constrained by a heavy reliance on heterologous protein regulators, whose associated immunogenicity and strict cargo-size limits impede clinical translation. Here, we introduce SpliceRACER (Splice-based <u>R</u>iboswitch <u>A</u>rray for <u>C</u>ombinatorial <u>E</u>xpression <u>R</u>egulation), a multiplexed RNA switch platform that achieves scalable, orthogonal, and combinatorial control of gene expression in mammalian cells through alternative splicing, eliminating the need for exogenous protein co-expression. SpliceRACER independently orchestrates three orthogonal channels to regulate fluorescent reporter outputs, yielding 3³ = 27 discrete combinatorial expression states, vastly exceeding the capacity of conventional binary gene switches. Furthermore, we engineered three generalizable strategies to effectively suppress the basal leakage inherent to synthetic riboswitches, thereby enabling tight OFF-state control of the potent pyroptotic effector GSDMD-NT for precise cell-death induction. This work offers broad utility in synthetic biology and therapeutic applications.

**Graphical abstract:** 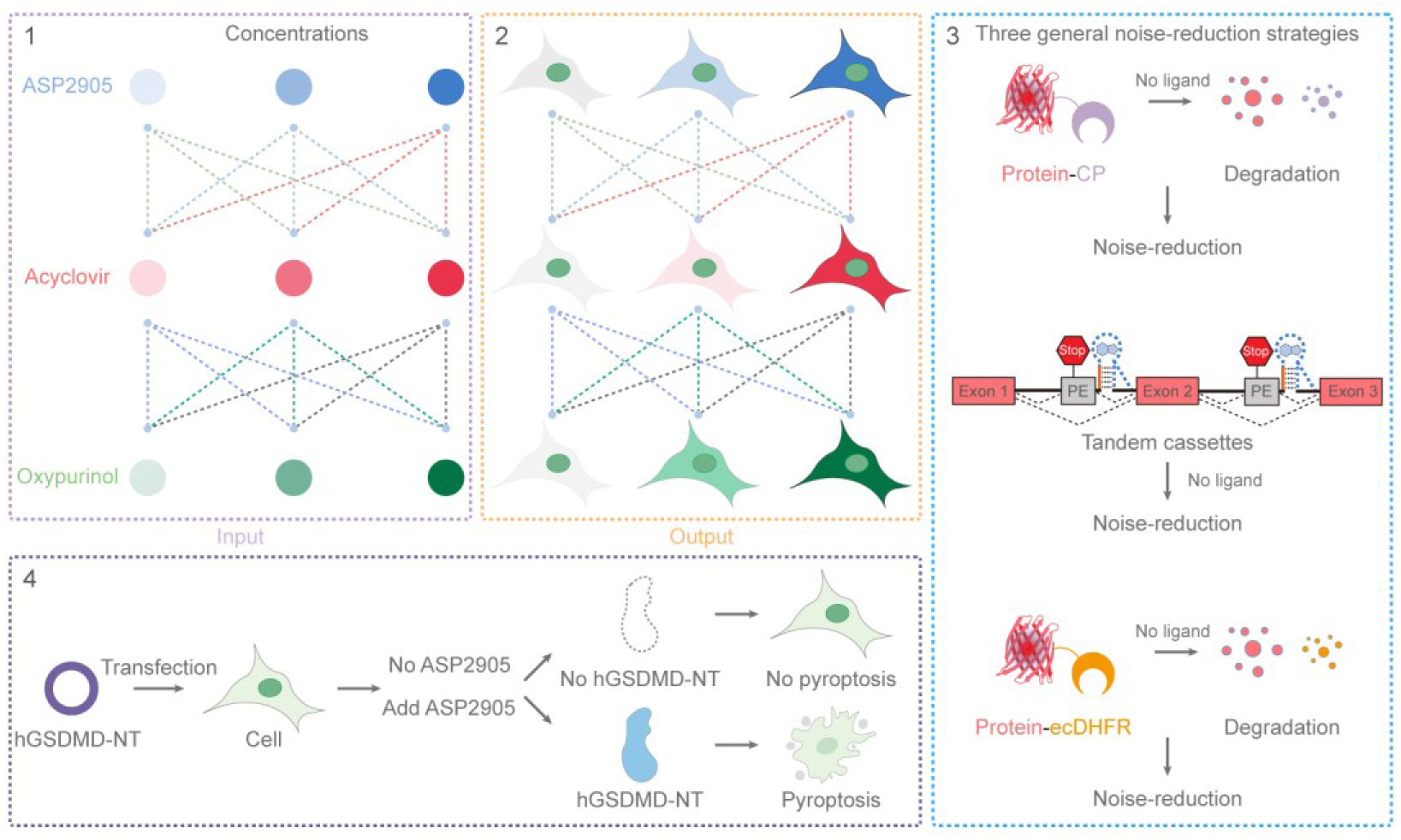

## Introduction

Mammalian synthetic circuits have evolved from simple toggle switches into multilayered networks capable of logic computation, cellular memory, and signal integration [1–5]. Yet a fundamental constraint persists: virtually all sophisticated gene programs rely on heterologous protein regulators—transcription factors, recombinases, proteases, RNA-binding proteins, or RNA-guided DNA-targeting effectors [6–10]. These heterologous components impose multifaceted burdens, manifesting as immunogenicity in clinical settings [11–14], packaging constraints for viral delivery [15], and metabolic stress on host cells [16,17]. Despite iterative advances in protein engineering, the field lacks a general strategy to decouple circuit complexity from exogenous protein expression—an indispensable prerequisite for clinical translation.

RNA-based regulatory devices have emerged as a compelling alternative to heterologous protein regulators, circumventing immunogenicity while maintaining a compact genetic footprint compatible with single-vector delivery. Among these, synthetic small-molecule-responsive riboswitches have demonstrated robust, ligand-inducible control over splicing, transcript stability, and translation in mammalian cells [18]. Diverse RNA switches built upon endogenous alternative splicing have been developed, offering a minimal architecture with high responsiveness and fidelity, plug-and-play ligand compatibility, and exceptional portability and plasticity [19–28]. However, despite their modular potential, the field remains largely confined to single-gene regulation, as scaling to multiplexed networks introduces substantial crosstalk and leakage. Cells natively process multidimensional signals through highly interconnected networks; binary switches lack the encoding capacity to recapitulate such complexity, limiting both our fundamental understanding of complex cell fates and the precise reprogramming of complex cellular phenotypes. Addressing this gap requires programmable, protein-free gene networks capable of orchestrating multidimensional cellular responses—a challenge that, if met, would redefine the programmability of mammalian synthetic biology.

Here, we present SpliceRACER, an all-RNA platform that harnesses host spliceosomal machinery to achieve scalable, orthogonal, and combinatorial gene control in mammalian cells without exogenous protein regulators. Exploiting the high specificity of RNA aptamers, we integrated three distinct aptamer–ligand pairs into a single system, preserving strict orthogonality within the complex cellular milieu, with each pair independently governing target-gene ON/OFF switching and precise expression tuning. Through combinatorial integration of these three inputs, SpliceRACER orchestrates three fluorescent reporters across low, medium, and high levels, generating 3³ = 27 discrete expression states. To address the pervasive challenge of basal leakiness in synthetic riboswitches, we engineered three general noise-reduction strategies operating at the post-transcriptional and post-translational levels. This framework endows the RNA switches with near-zero leakiness, enabling tight control over the potently toxic pyroptosis effector GSDMD-NT to drive inducible cell death.

To our knowledge, SpliceRACER establishes the first fully RNA-based, protein-free paradigm for multidimensional gene regulation in mammalian cells. By decoupling sophisticated network behavior from exogenous regulatory elements, this work offers a foundational technological architecture for safe, controllable synthetic biology and cell therapy.

## Materials and methods

### Compounds

Unless otherwise noted, all chemicals were purchased from commercial suppliers. Acyclovir (A126073), ganciclovir (G129641), penciclovir (P125160), trimethoprim (T129928), and triacetylganciclovir (T304788) were obtained from Aladdin. 6-Chloroguanosine (A800140), 8-bromo-2′-deoxyguanosine (B803865), guanosine monophosphate (G822613), *N*²-dimethylaminomethylene-2′-deoxyguanosine (*N*²-DMF-dG, N928881), 9-β-D-arabinofuranosylguanine (Ara-G, B829538), guanosine (G6043), 9-ethylguanine (E912930), and inosine (I6219) were sourced from Macklin. Valganciclovir (T1533), famciclovir (T1646), diacetylacyclovir (T65324), 2′-deoxyguanosine monohydrate (T4715), *N*²-isobutyrylguanosine monohydrate (TNU1135), 8-bromoguanosine (T7924), *N*²-methylguanosine (T7514), 6-*O*-methylguanosine (T7491), 7-methylguanosine (T7319), desciclovir (T22304), allopurinol riboside (T13539), ASP2905 (T9070), and oxypurinol (T1694) were acquired from TargetMol. Compounds were accurately weighed, dissolved in DMSO at desired concentrations with sonication assistance, filtered through a 0.22-μm syringe filter to ensure sterility, and stored in amber microcentrifuge tubes at −20 °C.

### Cell culture

HEK293T (CRL-11268), HeLa (CCL-2), MCF-7 (HTB-22), and A549 (CCL-185) cells (all obtained from ATCC) were maintained in high-glucose DMEM (C2701, Beyotime) supplemented with 10% (v/v) fetal bovine serum (FBS; A5256701, Gibco) and 1% (v/v) penicillin–streptomycin (C0222, Beyotime) at 37 °C in a humidified incubator containing 5% CO_2_. All experiments were conducted with cells at passages below 30 (passage < 30). For routine subculturing, cells were detached using 0.25% trypsin– EDTA (HY-K3009, MCE) following the manufacturer’s protocol.

### Plasmid construction

All plasmids were constructed via homologous recombination using the ClonExpress II One Step Cloning Kit (C112, Vazyme). DNA inserts were commercially synthesized by Sangon Biotech (Shanghai, China) or generated by PCR amplification. DH5α competent cells (C502, Vazyme) were transformed for plasmid propagation, and positive transformants were selected on Luria–Bertani (LB) agar supplemented with appropriate antibiotics. All constructs were validated by Sanger sequencing. Full plasmid sequences are provided in **Supplementary Note 1**, and key plasmids have been deposited with Addgene for public distribution upon publication.

### Transfection

HEK293T cells were seeded in 24-well plates and transfected 18–24 h post-seeding at ∼70% confluence. For each well, 0.5 µg of plasmid DNA was diluted in 25 µL of Opti-MEM (31985062, Gibco) and mixed with 0.8 µL of Lipo8000™ transfection reagent (C0533, Beyotime). Transfection complexes were incubated at room temperature for 15–20 min before being added dropwise to the culture medium. Transfection reagent and plasmid amounts were scaled proportionally for 6-well or 96-well plate formats. HeLa, MCF-7, and A549 cells were transfected using the same protocol.

### Microscopy

Cells were seeded in 24-well plates and transfected as described above. At 24 h post-transfection, the culture medium was replaced with phenol red-free DMEM (A1896701, Gibco). Cells were subsequently stained with Hoechst 33342 (1:1,000; C1028, Beyotime) and propidium iodide (PI; 1:1,000; C1008S, Beyotime) in the dark for 1 h at 37 °C prior to imaging. Fluorescence imaging was performed on a Leica DMi8 inverted fluorescence microscope. Images were acquired and processed using LAS X software, with identical exposure times and gain settings maintained across all comparative groups to ensure measurement fidelity.

### RT–PCR analysis

HEK293T cells were seeded in 6-well plates and transfected at ∼70% confluence as described above. At 24 h post-transfection, total RNA was extracted using RNA isolater Total RNA Extraction Reagent (R401, Vazyme) followed by isopropanol precipitation.

After genomic DNA removal, cDNA was synthesized from 1 µg of total RNA using the Hifair III 1st Strand cDNA Synthesis SuperMix (11141ES60, Yeasen) according to the manufacturer’s instructions. PCR amplification was performed using Phusion High-Fidelity DNA Polymerase (M0530S, NEB) with specific primers flanking the alternative exon (forward: 5′-GACTTTGACATGGTGGGTCAG-3′; reverse: 5′- CTTGGTATCCGGAGCCATCTA-3′). Thermal cycling conditions comprised an initial denaturation at 95 °C for 3 min; 30 cycles of 95 °C for 30 s, 55 °C for 30 s, and 72 °C for 30 s; and a final extension at 72 °C for 5 min. Amplified products were resolved on a 3% (w/v) agarose gel in 1× TAE buffer at 120 V for 1 h, visualized under UV light, and imaged using a Tanon MINI Space 1000 System.

### Dual-luciferase reporter assay

All psiCHECK-2-based reporter plasmids, including PGK-Renilla, PGK-Renilla-T2, and PGK-Renilla-Cyclone, were constructed using the psiCHECK-2 backbone (C8021, Promega). HEK293T cells were seeded in 48-well plates to achieve ∼70% confluence 24 h prior to transfection. Per well, cells were transfected with 250 ng of reporter plasmid in 12.5 µL Opti-MEM using 0.4 µL Lipo8000™ transfection reagent (C0533, Beyotime) according to the manufacturer’s instructions. Experimental groups were treated with specified concentrations of acyclovir or ASP2905, whereas control groups received equivalent volumes of vehicle (DMSO). All conditions were performed in biological triplicates. Blank wells were included for background subtraction, and wells expressing PGK-Renilla served as normalization controls for cross-plate comparisons. At 24 h post-transfection, cells were lysed and luminescence was quantified using the Dual-Lumi™ Luciferase Reporter Gene Assay Kit (RG088S, Beyotime). Firefly luciferase activity was measured first by adding 50 µL of substrate solution and incubating at room temperature for 10 min prior to reading on an EnVision multimode plate reader (PerkinElmer). Renilla luciferase activity was subsequently recorded following the addition of 50 µL of Renilla reaction/quenching buffer under identical detection parameters. After background subtraction, relative luminescence units (RLUs) were calculated as the Renilla/Firefly ratio and normalized to the PGK-Renilla reference group to account for variations in transfection efficiency and cell viability. Data are presented as mean ± SD relative expression percentages.

### Western blot

Following cell lysis with lysis buffer (WB3100, NCM Biotech) and protein quantification via the BCA Protein Assay Kit (WB6501, NCM Biotech), protein samples were normalized to equal concentrations and denatured in 5× SDS-PAGE sample loading buffer at 95 °C for 5 min. Proteins were separated by SDS-PAGE and electrically transferred onto polyvinylidene fluoride (PVDF) membranes (Millipore). Membranes were blocked with 5% (w/v) bovine serum albumin (BSA; 4240GR100, BioFroxx) in Tris-buffered saline containing 0.1% Tween-20 (TBST) for 1 h at room temperature, followed by overnight incubation at 4 °C with primary antibodies: anti-Gasdermin D (1:5,000 dilution; HA721144, HUABIO) and anti-β-actin (1:1,0000 dilution; 66009-1-Ig, Proteintech). After three washes with TBST (10 min each), membranes were incubated with appropriate HRP-conjugated secondary antibodies (1: 1,0000 dilution) for 1 h at room temperature. Following subsequent washes with TBST, protein bands were visualized using an enhanced chemiluminescence (ECL) detection kit (BMU102, Abbkine) and imaged on an Azure 500 Western Blot Imaging System (Azure Biosystems).

### CCK-8 assays

Cell viability following ligand treatment was evaluated using the Cell Counting Kit-8 (CCK-8; C0005, TargetMol) according to the manufacturer’s instructions. Cells were seeded in 96-well plates at a density of 5 × 10³ cells per well in 100 µL of complete high-glucose DMEM. Upon reaching 70–80% confluence, the culture medium was replaced with fresh medium containing indicated concentration gradients of specified ligands, followed by incubation at 37 °C for 24 h. Subsequently, 10 µL of CCK-8 reagent was added to each well, and plates were incubated at 37 °C for an additional 2 h in the dark. Absorbance at 450 nm was measured using a microplate reader. Cell viability was calculated using the formula: Cell viability (%) = [(OD_test_ − OD_blank_) / (OD_control_ − OD_blank_)] × 100%, where OD_test_, OD_control_ and OD_blank_ represent the absorbance values of ligand-treated wells, untreated vehicle control wells, and background wells containing medium without cells, respectively. All assays were conducted with at least three independent biological replicates.

### Statistics and reproducibility

Unless otherwise stated, all quantitative assays were performed with at least three independent biological replicates. Statistical analyses were conducted using GraphPad Prism (v10.1.2), and high-resolution vector figures were assembled using Adobe Illustrator 2024 (v28.7.7). Data are presented as mean ± standard deviation (s.d.). Statistical significance was assessed using two-tailed unpaired Student’s t-tests, one-way ANOVA, or Welch’s ANOVA (for unequal variances), with appropriate post-hoc tests (Tukey’s or Dunnett’s tests) applied where applicable. A *P*-value < 0.05 was considered statistically significant (\**P* < 0.05, \*\**P* < 0.01, \*\*\**P* < 0.001, \*\*\*\**P* < 0.0001; NS, not significant).

## Results

### Dual-channel RNA switches

Alternative splicing orchestrates post-transcriptional gene regulation, converting single pre-mRNAs into multiple isoforms via differential exon combinatorics, splice site selection, or intron retention [29–31]. This regulatory paradigm expands proteomic complexity, operating across >90% of the human transcriptome [32,33]. RNA switches based on small-molecule-induced poison exon splicing enable precise post-transcriptional regulation. However, existing platforms face an inescapable trade-off: stringent OFF-state silencing versus robust ON-state activation. An ideal switch demands negligible baseline leakage coupled with potent ligand-dependent induction. This trade-off is exemplified by two representative platforms: the acyclovir-responsive Cyclone system leverages a clinically approved antiviral to achieve robust induction, but exhibits appreciable basal leakiness [26] (Supplementary Fig. S2); conversely, switches governed by the potassium channel inhibitor ASP7967 or its analog ASP2905 achieve near-zero background yet display suboptimal induction efficacy, rarely exceeding 50% of wild-type expression levels as standalone modules [22]. These constraints define a critical technological gap: the absence of a single switch capable of concurrently achieving ultra-low background and high-level output.

To address this gap, we engineered an RNA splicing switch that seamlessly couples ultra-low background leakage with potent ligand-dependent induction. The system comprises three core modules: a versatile intron–poison exon–intron cassette compatible with diverse genetic loci; an RNA aptamer selective for a cognate small-molecule ligand; and a transduction domain that transduces ligand binding into an exon-skipping output (Fig. 1A). Strategically positioned within the second intron downstream of the 5′ splice site, the aptamer and transduction domain function in cis. In the absence of ligand, the metastable transduction domain promotes spliceosomal recognition, favoring poison exon inclusion. This triggers nonsense-mediated decay [34] or generates truncated, non-functional proteins (Fig. 1A). Ligand binding induces an allosteric switch in the aptamer, stabilizing the transduction domain, masking spliceosomal recognition motifs, and driving obligate poison exon skipping to produce full-length, functional proteins (Fig. 1A). System performance thus obeys a thermodynamic “Goldilocks principle”: insufficient transduction-domain stability favors constitutive inclusion and limits activation, whereas excessive stability drives constitutive skipping and elevated basal leakage [35].

**Figure 1.**
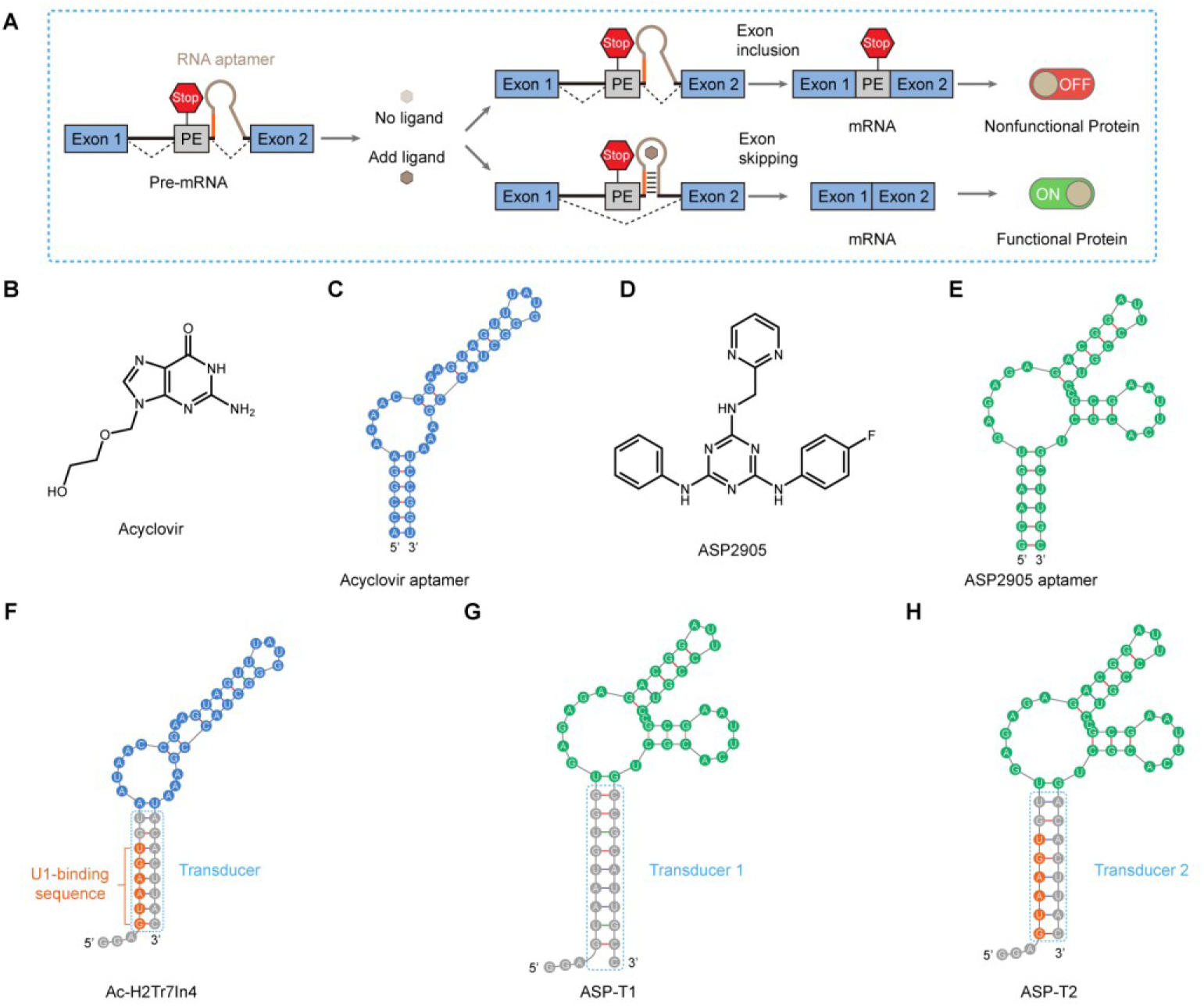
Molecular mechanism, chemical structures, and secondary structure predictions of ligand-gated poison exon (PE) splicing switches. (**A**) Schematic of PE– mediated genetic switching. Ligand binding to the embedded aptamer sterically occludes the 5′ splice site, toggling alternative splicing between PE inclusion (OFF state) and PE skipping (ON state). (**B**–**E**) Chemical structures and cognate aptamers. Acyclovir (**B**) and its aptamer (**C**); ASP2905 (**D**) and its aptamer (**E**). (**F**–**H**) Sequences and predicted secondary structures of wild-type Cyclone (Ac-H2Tr7In4; **F**), ASP-T1 (**G**), and ASP-T2 (**H**).

Despite extensive optimization of the transduction domain, ASP2905-responsive switches exhibit suboptimal induction [35]. We hypothesized that this performance bottleneck arose from the native cassette architecture rather than the transduction domain itself. To dissect this architectural constraint, we replaced the original cassette with the contiguous “intron 6_trunc_–exon 7–intron 7” segment from Cyclone [26], and engineered two transduction-domain variants, ASP-T1 and ASP-T2, to delineate the basis of allosteric control (Fig. 1G, H).

We benchmarked these designs using the bright, photostable fluorescent proteins mScarlet3H [36] and mNeonGreen [37] as reporters. Guided by established design principles [26], we precisely inserted ASP-T1/2 cassettes at defined loci within mScarlet3H (Gln¹⁰⁹–Asp¹¹⁰; CAG–GAC) and mNeonGreen (Glu⁴⁵–Glu⁴⁶; GAG–GAG) to preserve the reading frame. This yielded the PGK-3H-ASP-T1/2 (Supplementary Fig. S1D, E) and PGK-mNG-ASP-T1/2 (Supplementary Fig. S1G, H) constructs. For benchmarking, we included two control sets: (i) constructs carrying the wild-type Cyclone cassette at identical insertion sites (Supplementary Fig. S1C, F), and (ii) constitutively active reporter plasmids to define maximal achievable expression levels.

To functionally characterize these constructs, we evaluated their ligand responsiveness in HEK293T cells. Transient transfection of mScarlet3H constructs followed by 24 h treatment with acyclovir (125 µM) or ASP2905 (10 µM) revealed tight ligand-gated control: red fluorescence was virtually undetectable in the absence of ligand, whereas ligand addition drove robust expression comparable to that of unmodified PGK-3H controls (Supplementary Fig. S2A, left). Notably, contrast enhancement unmasked pronounced background fluorescence from the PGK-3H-Cyclone construct in the absence of acyclovir, confirming substantial basal leakiness (Supplementary Fig. S2A, right). In stark contrast, PGK-3H-ASP-T1/2 constructs exhibited negligible baseline fluorescence, indistinguishable from untransfected controls (Supplementary Fig. S2A, right).

Parallel assays with the mNeonGreen reporter recapitulated these observations: in the absence of ligand, PGK-mNG-Cyclone exhibited marked basal fluorescence, whereas PGK-mNG-ASP-T1/2 consistently maintained near-background signals indistinguishable from untransfected controls (Supplementary Fig. S2B, right). Upon ligand induction, both PGK-mNG-Cyclone and PGK-mNG-ASP-T2 achieved robust expression, exceeding that of unmodified PGK-mNG controls (Supplementary Fig. S2B, left). This low-leakage advantage generalized across diverse contexts: across promoters of varying strengths (CMV, SV40, Ubc) and diverse cancer cell lines (HeLa, MCF-7, A549) tested, basal expression of acyclovir-responsive switches consistently exceeded that of their ASP2905-responsive counterparts, while maximal ligand-activated outputs remained comparable between the two systems (Supplementary Figs S3–S8). Dual-luciferase reporter assays further confirmed at the biochemical level that ASP2905-responsive riboswitches exhibit significantly lower baseline activity than the acyclovir-responsive equivalents (Supplementary Figs S26 and S29). Together, these data establish the engineered ASP-T2 switches as an ultra-stringent, near-zero-leakage riboswitch platform with broad promoter and cell-type compatibility.

To further evaluate the performance of the ASP-T2 switch, we performed a head-to-head benchmarking against two recently reported splicing-based riboswitches: a theophylline-inducible multivalent aptamer switch and a tetracycline-inducible dimeric aptamer (apdimer) switch [38,39]. Although these prior studies marked notable advances in optimizing splicing-based riboswitches [38,39], direct side-by-side evaluation revealed dramatic differences. Upon transfecting the optimal constructs (TriTheo-1-8 and Tet_splice_apdimer_2) into HEK293T cells followed by treatment with cognate ligands at reported optimal concentrations, ASP2905 induced robust green fluorescence comparable to that of a constitutively active control (Supplementary Fig. S9). In contrast, theophylline- and tetracycline-induced fluorescence remained barely detectable under standard conditions and required substantial contrast enhancement to be visualized (Supplementary Fig. S9). These results establish that ASP2905-responsive riboswitches exhibit marked superiority over existing alternatives, seamlessly combining robust ligand-inducible expression with ultra-low basal leakiness.

To directly confirm that small-molecule-induced protein expression stems from regulated alternative splicing, we performed reverse transcription PCR (RT–PCR) analysis. In the ligand-free state, mNeonGreen transcripts migrated as a single predominant band (∼250 bp), consistent with poison exon inclusion, which adds ∼50 bp to the mature transcript (Supplementary Fig. S10). Ligand addition triggered efficient exon skipping, yielding a major ∼200-bp product that co-migrated with transcripts from unmodified positive controls (Supplementary Fig. S10). Sanger sequencing further revealed that in the absence of ASP2905, mNeonGreen transcripts incorporated SMN1 exon 7 sequences, whereas ASP2905 treatment yielded scarless transcripts devoid of SMN1 exon 7 (Supplementary Fig. S10C). To confirm that gene activation requires direct ligand engagement with the aptamer, we introduced point mutations into the conserved ligand-binding pocket of the ASP2905 aptamer [22], generating PGK-mNG-ASP-T2-M2/3/4 constructs (Supplementary Fig. S11B, right). These mutations completely abrogated ligand responsiveness, with neither green fluorescence nor detectable splicing shifts observed upon ASP2905 treatment (Supplementary Fig. S11B, C). Collectively, these data establish that ligand-binding-induced exon skipping is the primary regulatory mechanism governing protein output in this system.

To enable dual-channel gene regulation, we next evaluated the orthogonality between acyclovir- and ASP2905-responsive switches. Given the stark structural divergence between acyclovir and ASP2905, we reasoned that their respective SELEX-derived aptamers would exhibit negligible cross-reactivity [22,40–42], enabling concurrent, orthogonal operation. Fluorescence imaging of mScarlet3H and mNeonGreen reporters confirmed negligible inter-system crosstalk (Supplementary Fig. S2) —a finding further substantiated by RT–PCR analyses demonstrating ligand-specific splicing shifts (Supplementary Fig. S10B). This dual-channel platform thus enabled precise, independent control over two protein targets, each exhibiting dose-dependent, monotonic scaling with its cognate ligand (Supplementary Figs S12 and S13). Systematic titration of both inputs yielded a wide dynamic range of combinatorial expression states for red and green fluorescent proteins, underscoring its robust programmability (Fig. 2). Notably, this multiplexed control was achieved without compromising cell viability (Supplementary Fig. S38A). Together, these data establish a general framework for independent, orthogonal, and tunable regulation of gene expression programs in mammalian cells.

**Figure 2.**
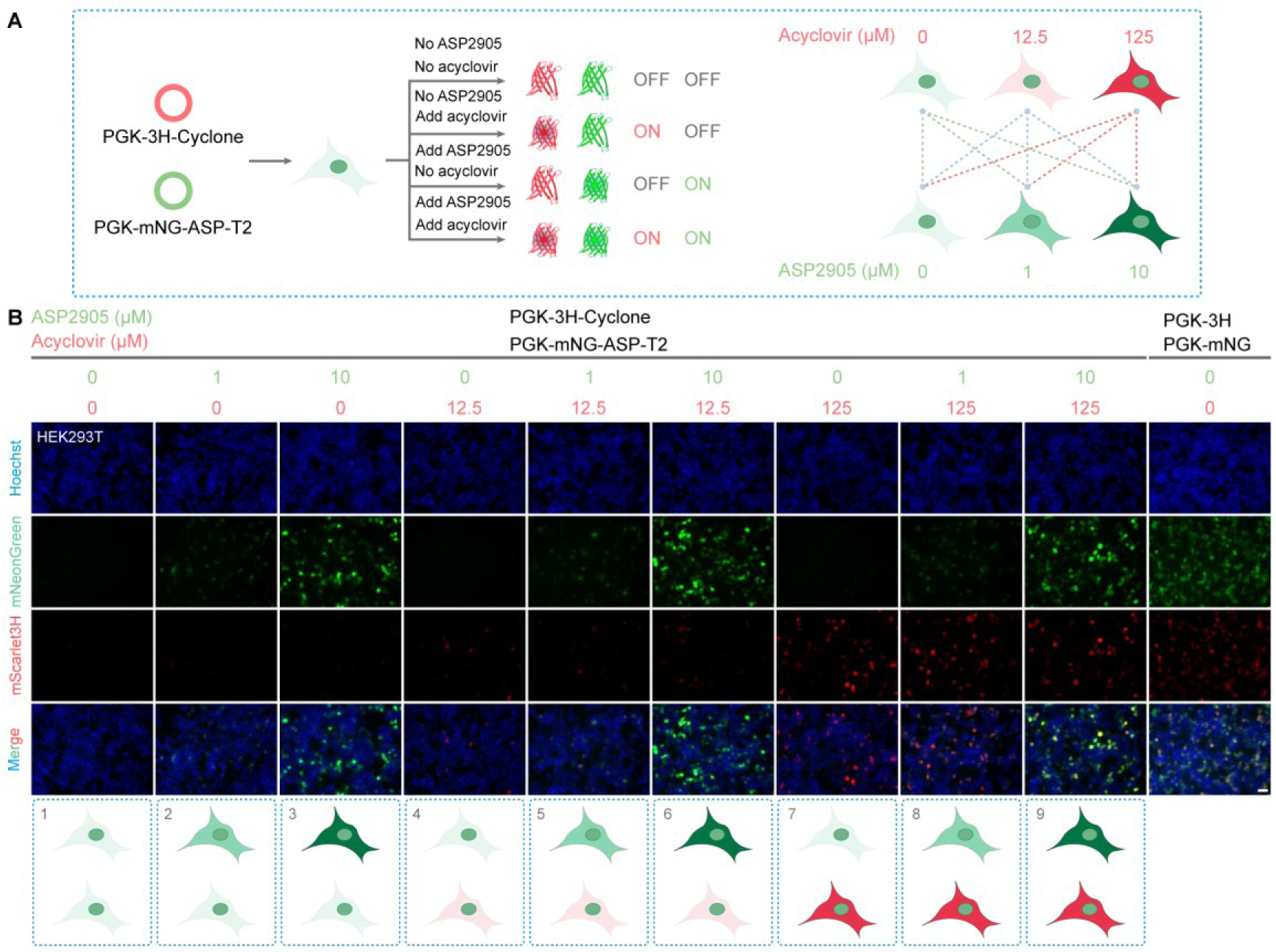
Combinatorial ligand inputs drive orthogonal dual-channel expression. (**A**) Acyclovir and ASP2905 serve as independent orthogonal inputs that selectively control mScarlet3H and mNeonGreen expression, respectively. (**B**) Dual-channel fluorescence readouts mapped across a two-dimensional ligand concentration matrix. Scale bars, 50 μm.

### Multiplexed RNA switches

Building on dual-channel designs, we sought to develop a unified platform that orchestrates multiple orthogonal RNA switches for multiplexed, small-molecule-gated control of protein outputs. This architecture offers a direct route to exponentially scalable proteome reprogramming, dramatically expanding the functional diversity of synthetic circuits. Yet, realizing this vision remains formidable. First, scalability is severely constrained by compounding circuit complexity as systems expand beyond two targets [43]. Second, current strategies lack the precision required to fine-tune multiple protein levels within a single cellular context [44]. Third, widespread reliance on exogenous protein effectors imposes systemic complexity and clinical immunogenicity risks [1,3]. Finally, most existing platforms fail to support dynamic, reversible regulation across multiple genes in parallel [45]. These collective constraints represent a critical bottleneck impeding the engineering of sophisticated, multiplexed gene circuits.

To overcome these barriers, we expanded the regulatory repertoire by introducing a third chemically inducible switch responsive to oxypurinol and its structural analogs [24]. Mirroring the regulatory logic of the ASP2905- and acyclovir-gated systems, oxypurinol drives dose-dependent expression by toggling between poison exon inclusion and skipping (Supplementary Fig. S14). Crucially, all three RNA switches displayed strict mutual orthogonality, with imperceptible crosstalk across all pairwise combinations (Supplementary Figs S15–S18). Together, these modular switches establish a versatile toolkit for parallel integration, enabling the seamless orchestration of multiple independent RNA circuits within a unified regulatory architecture.

Building on this unified architecture, we experimentally validated the platform by integrating three distinct, orthogonal RNA switches into a single system. In this triple-channel design, ASP2905, acyclovir, and oxypurinol independently controlled the expression of mTagBFP2 (blue) [46], mScarlet3H (red), and mNeonGreen (green), respectively (Supplementary Fig. S19A–C). This architecture enabled robust, orthogonal regulation across three distinct protein outputs (Supplementary Fig. S19D). In principle, systematic ligand titration enables independent, quantitative tuning of each reporter, mapped across a multidimensional combinatorial output space (Fig. 3). Mathematically, a system with *N* discrete concentration steps per ligand yields up to *N*³ distinct expression states. Experimentally, we defined three concentration levels per input and successfully resolved all 3³ = 27 predicted expression combinations (Supplementary Fig. S20). We designated this platform SpliceRACER (Splice-based <u>R</u>iboswitch <u>A</u>rray for <u>C</u>ombinatorial <u>E</u>xpression <u>R</u>egulation).

**Figure 3.**
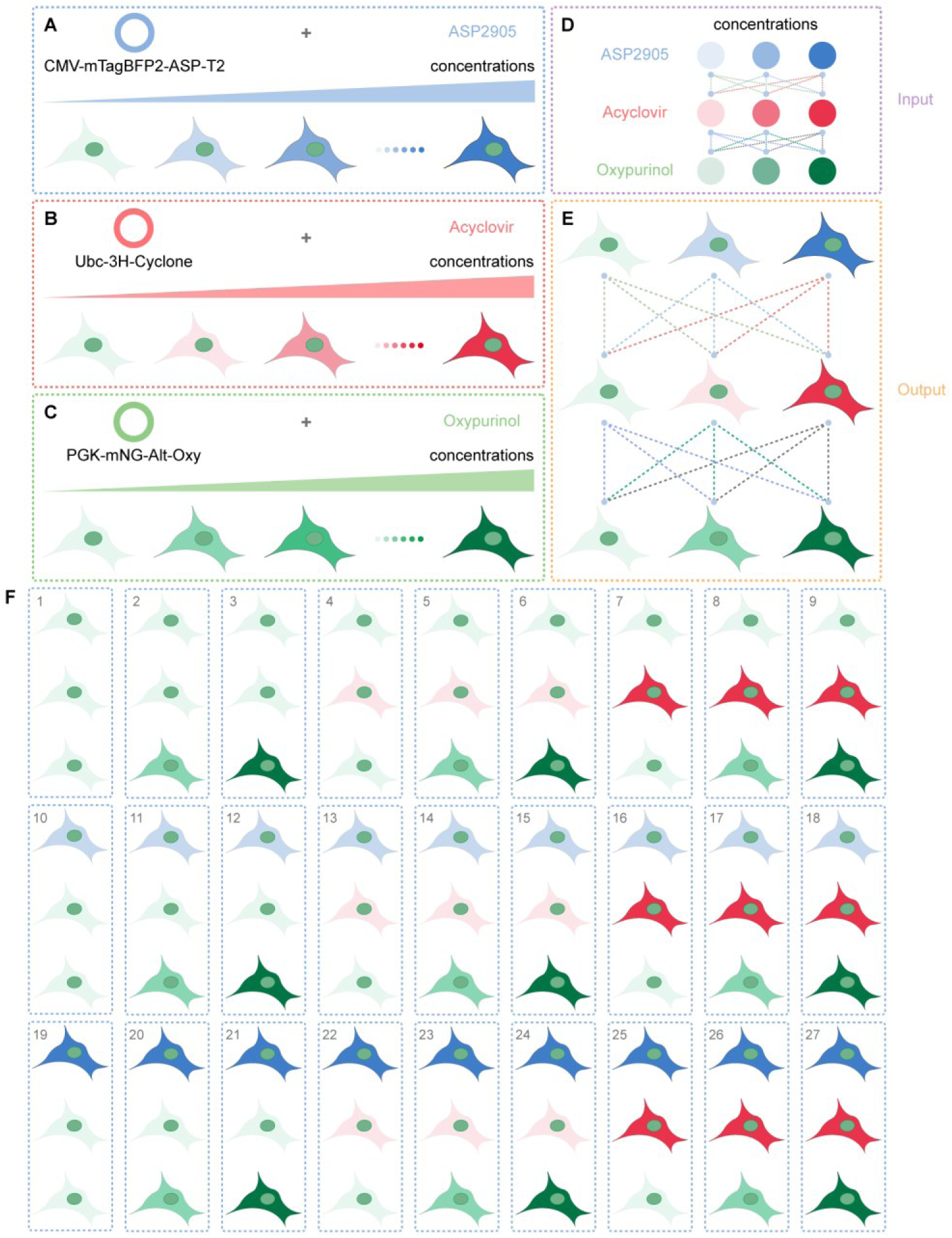
A three-input ligand repertoire expands the output space to 27 discrete states. (**A**–**C**) ASP2905, acyclovir, and oxypurinol serve as orthogonal inputs that dose-dependently control mTagBFP2 (blue), mScarlet3H (red), and mNeonGreen (green) expression, respectively. (**D**) Experimental scheme for the three-input combinatorial layout. Ternary ligand inputs expand the phenotypic state space to 27 resolvable outputs. (**E**, **F**) Three-dose treatment (none, low, high) per ligand yields 3^3^ = 27 combinatorial fluorescence states.

Thus, by virtue of its modular architecture and intrinsic orthogonality, SpliceRACER provides a versatile foundation for engineering increasingly sophisticated gene networks. Multiple ligands can be harnessed to combinatorially gate the expression of a single target protein (Supplementary Fig. S21); conversely, a single ligand can coordinate the expression of multiple targets within a single cell (Supplementary Fig. S22). In principle, with each ligand independently regulating *N* distinct proteins, SpliceRACER enables coordinated control over up to 3 × *N* distinct protein species. Fine-tuning ligand concentration gradients further renders these parallel outputs quantitatively tunable, facilitating the construction of complex, programmable gene networks (Supplementary Fig. S23). This scalability and precision fundamentally expand the design space of synthetic biology, establishing a new paradigm for mammalian gene circuit engineering.

### General noise-reduction strategies

Although the ASP2905-gated switch exhibits negligible basal leakage, the lack of clinical safety data for ASP2905 limits its immediate translational utility [47]. In contrast, acyclovir, a clinically approved, first-line anti-herpesvirus drug, possesses an established safety profile [48], positioning it as the preferred candidate for clinical translation. However, the wild-type Cyclone switch suffers from substantial basal leakage—a major bottleneck for applications requiring stringent OFF-state silencing. This limitation is especially acute when controlling constitutively active editors like SpCas9, where even trace background expression drives persistent DNA editing and genotoxic risks [49]. Similarly, leaky expression of potent cytotoxic effectors induces lethal toxicity in host cells [50]. To overcome these constraints, we systematically re-engineered the Cyclone architecture to achieve stringent, near-zero basal leakage.

To dissect the molecular origin of basal leakiness in wild-type Cyclone, we profiled its response to a panel of acyclovir analogs—valacyclovir, ganciclovir, valganciclovir, and penciclovir. All tested analogs potently induced mNeonGreen expression, unmasking substantial binding promiscuity within the aptamer (Supplementary Fig. S24A). This broad cross-reactivity suggests that basal leakiness stems from the illegitimate recognition of endogenous nucleoside pools by the acyclovir aptamer, which triggers constitutive poison exon skipping. Further supporting this model, exogenous guanosine and related nucleosides robustly induced reporter expression (Supplementary Fig. S24). Conversely, several analogs failed to trigger expression despite exhibiting no cytotoxicity (Supplementary Figs S24 and S38C), likely due to steric occlusion within the binding pocket. Critically, *N*²-methylation of the guanine C2-amino group completely abolished reporter activity, confirming that even minimal steric hindrance at this site disrupts aptamer binding (Supplementary Fig. S24B). Notably, guanosine, 2′-deoxyguanosine, and GMP each drove expression even more efficiently than acyclovir, implying that endogenous purine pools constitutively occupy the aptamer pocket and establish a high basal expression baseline.

To establish causality between endogenous nucleosides and basal leakiness in wild-type Cyclone, we engineered three aptamer variants targeting the acyclovir-binding pocket: Cyclone-M2 and Cyclone-M4, which abolish binding to acyclovir and its analogs, alongside Cyclone-M5 as a ligand-responsive control [42] (Supplementary Fig. S25, left). Functional characterization confirmed that M2 and M4 failed to respond to either acyclovir or guanosine (Supplementary Fig. S25, center). Crucially, M2 and M4 exhibited markedly reduced basal leakiness in the ligand-free state (Supplementary Fig. S25, right), directly linking the loss of nucleoside binding to suppressed background expression. Collectively, these data establish that spurious recognition of endogenous nucleotides is the primary driver of basal leakiness in the wild-type Cyclone system.

To suppress basal leakiness in wild-type Cyclone, we developed two engineering strategies: the C-terminal fusion of degradation tags to recruit the ubiquitin– proteasome system (UPS) [51], and the tandem duplication of the regulatory cassette (Fig. 4A, B). For the degradation-based approach, we fused a composite CP tag comprising the CL1 peptide [52] and PEST degron [53] to the reporter C-terminus, targeting nascent polypeptides for rapid proteasomal degradation. In the ligand-free state, poison exon inclusion predominates, and trace full-length proteins arising from escape-splicing events are rapidly cleared by the CP tag, minimizing basal expression (Supplementary Fig. S27). Upon ligand addition, poison exon skipping increases markedly, yielding a surge of translatable transcripts that saturates UPS degradation and drives robust reporter accumulation (Supplementary Fig. S27). For the tandem-duplication strategy, poison exon inclusion predominates at both splicing junctions in the ligand-free state (Fig. 4B). The multiplicative attenuation across sequential, independent splicing events compounds to favor poison exon inclusion (Fig. 4B), thereby substantially reducing basal expression (Supplementary Fig. S28). Dual-luciferase reporter assays quantitatively confirmed that the CP-fusion (Cyclone-CP) and tandem-duplication (2×Cyclone) constructs suppressed basal expression by 4.1-fold and 2.0-fold, respectively, albeit with a modest reduction in maximal output (Supplementary Fig. S29).

**Figure 4.**
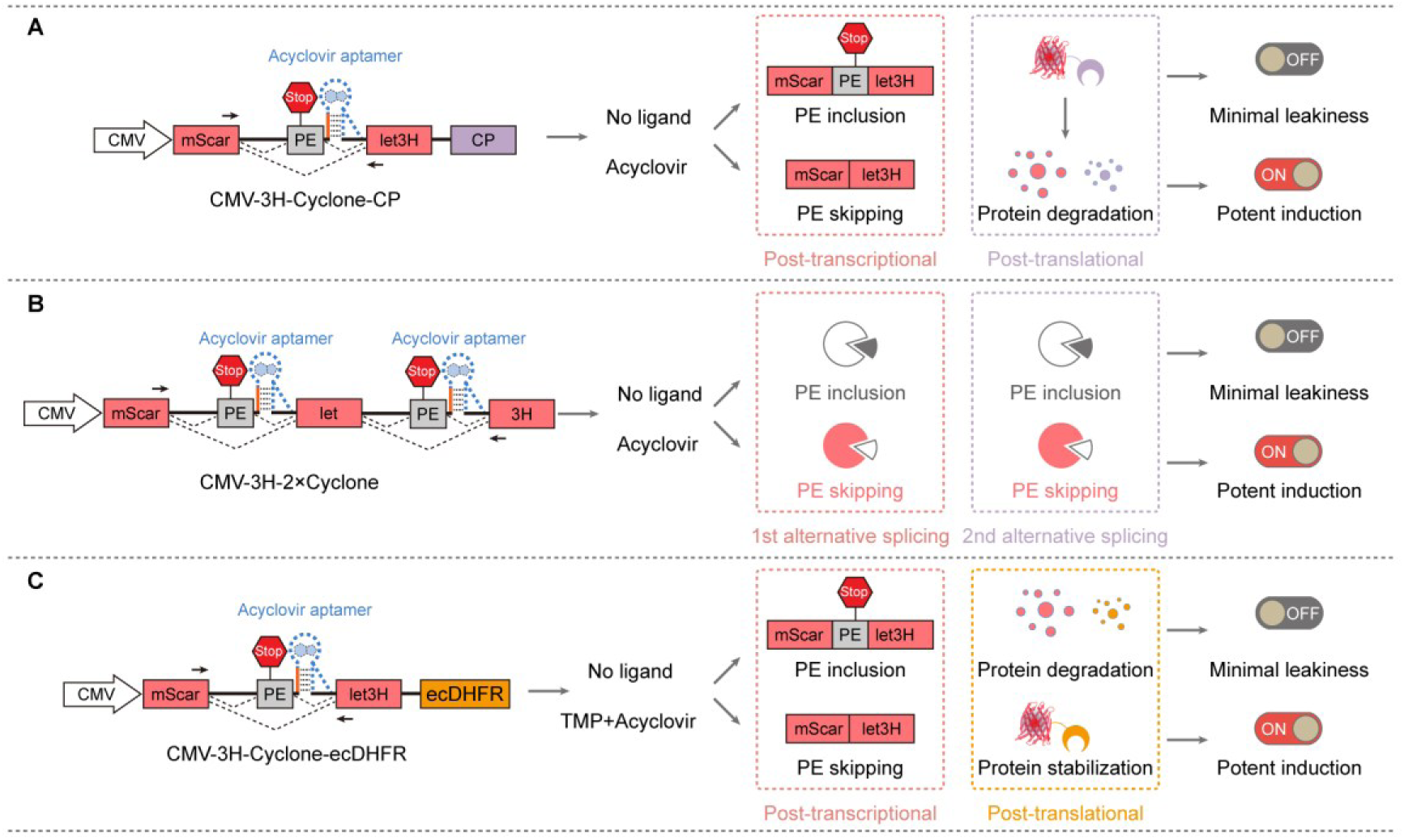
Three noise-reduction strategies for suppressing basal expression. (**A**) CP-degron fusion. Leaky fusion proteins produced in the ligand-free state are constitutively targeted for proteasomal degradation, mitigating basal protein accumulation independently of splicing regulation. (**B**) Tandem splicing switches. Basal leakage requires two independent PE skipping events; multiplicative probabilistic filtering favors PE inclusion, diverting leaky transcripts to NMD. (**C**) ecDHFR destabilizing domain fusion. Leaky proteins are constitutively degraded in the absence of ligand, while TMP binding stabilizes the fusion and restores full dynamic range, decoupling stringent basal repression from maximal induction.

In synthetic gene circuit engineering, minimizing basal leakiness often incurs a penalty in maximal induction, creating a critical performance bottleneck. A strategy capable of decoupling stringent OFF-state silencing from ON-state output capacity has therefore remained highly sought after. Previous studies have established that destabilizing domains (DDs) mediate the rapid, constitutive degradation of fused proteins—an effect rescued upon binding a cognate synthetic ligand that shields the domain from proteolysis [54,55]. The *Escherichia coli* dihydrofolate reductase (ecDHFR) mutant is a well-characterized DD that targets fused proteins for degradation in the absence of trimethoprim (TMP) (Supplementary Fig. S30). We thus fused ecDHFR to the C-terminus of the reporter embedded within the Cyclone expression cassette (Cyclone-ecDHFR) (Fig. 4C). In the ligand-free state, this dual-action design suppresses target protein leakage at both the post-transcriptional and post-translational levels. Upon the simultaneous addition of acyclovir and TMP, poison exon skipping is triggered concurrently with TMP-mediated stabilization of the reporter. This combinatorial strategy effectively minimizes basal leakiness without compromising maximal induced output, thereby resolving the canonical performance trade-off (Supplementary Fig. S31).

### Control pyroptosis with ASP2905-gated switch

To demonstrate the translational utility of this platform, we applied it to achieve precise, ligand-gated control over cytotoxic effectors. The N-terminal domain of Gasdermin D (GSDMD-NT) is a potent executor of pyroptosis that oligomerizes to form plasma membrane pores, driving cell lysis, the lytic release of damage-associated molecular patterns (DAMPs), and robust inflammatory cascades [56,57]. In resting cells, full-length GSDMD remains autoinhibited, as its C-terminal domain sterically occludes the cytotoxic N-terminal domain via an interdomain linker [58]. While inflammasome activation triggers caspase-mediated cleavage of this linker to execute pyroptosis [59], reliance on this endogenous cascade restricts its spatiotemporal control for therapeutic intervention [60]. Although heterologous systems (e.g., Cre/loxP, Tet-On 3G) can bypass upstream sensing to achieve GSDMD-NT-mediated tumor clearance [61,62], they inherently trigger host immunogenicity, exceed AAV packaging limits, and impose substantial metabolic burdens. Consequently, a traceless, protein-free RNA platform enabling tight GSDMD-NT regulation represents a major unmet need in precision oncology.

To address this limitation, we engineered an all-RNA regulatory system for GSDMD-NT expression that eliminates immunogenic foreign proteins and minimizes cellular metabolic load (Fig. 5B). Such a system must satisfy two stringent requirements: (i) near-zero baseline leakage in the OFF state to maintain GSDMD-NT levels strictly below the lytic threshold; and (ii) robust, rapid ON-state induction to push GSDMD-NT accumulation above this cytotoxic threshold. As a proof of concept, we initially harnessed the ASP2905-responsive riboswitch given its intrinsic, near-zero leakage. However, post-translational noise reduction via degradation tags suffers from kinetic hysteresis, failing to counteract the rapid, irreversible membrane perforation executed by even trace GSDMD-NT. We therefore implemented tandem riboswitch arrays to suppress escape-splicing events at the pre-mRNA level, preventing the generation of full-length GSDMD-NT transcripts (Fig. 5A).

**Figure 5.**
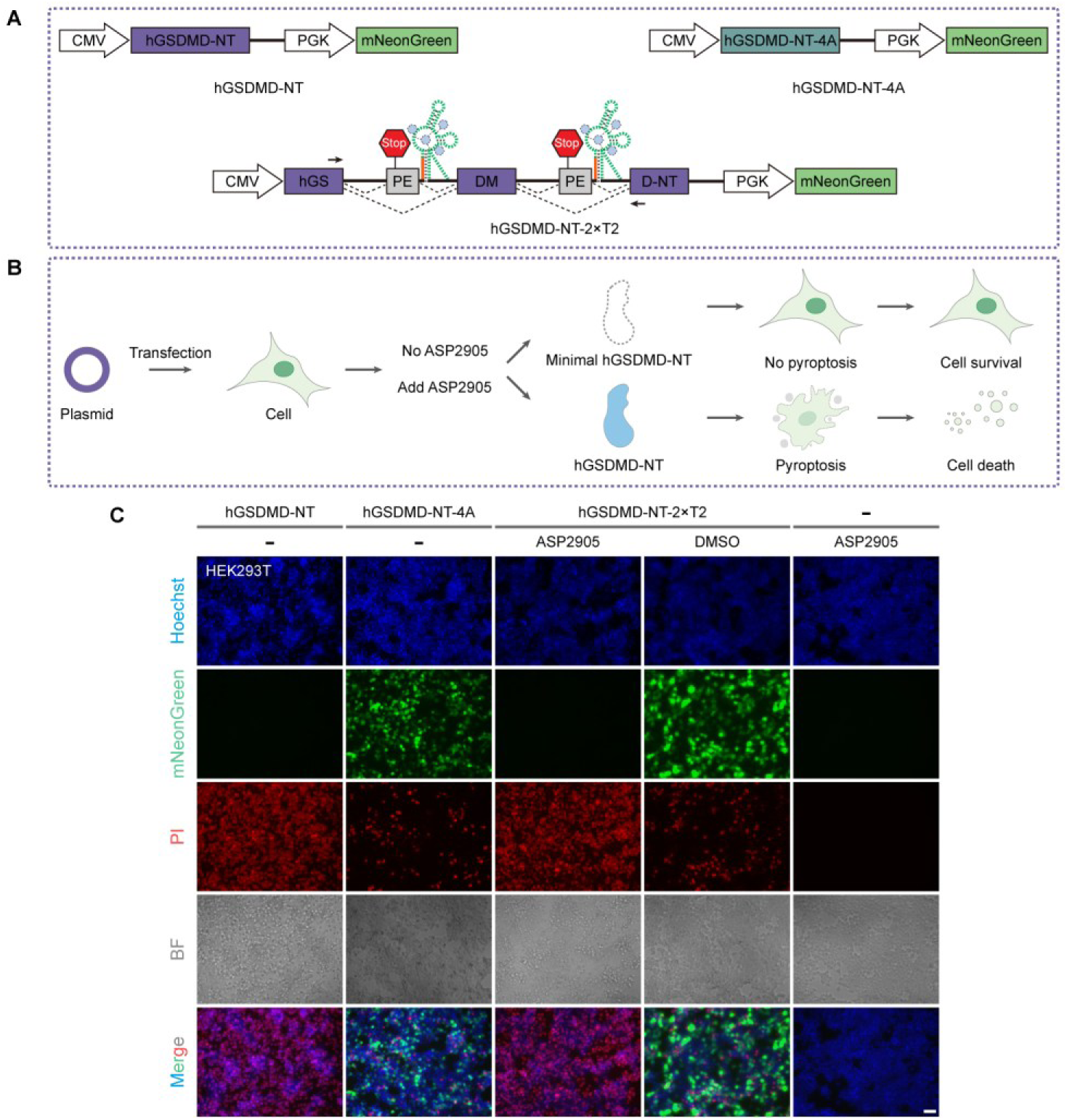
Precision chemical control of hGSDMD-NT-mediated pyroptosis via an ASP2905-responsive riboswitch. (**A**) Genetic architecture of the ASP2905-gated hGSDMD-NT expression construct. (**B**) ASP2905-induced hGSDMD-NT expression selectively triggers pyroptosis with tight OFF-state control. (**C**) Representative fluorescence micrographs showing PI-positive membrane permeabilization and cell death. Scale bars, 50 μm.

To enable direct phenotypic evaluation, we combined fluorescence microscopy with propidium iodide (PI) staining, which revealed tight, ligand-dependent control of hGSDMD-NT-2×T2 expression by the ASP2905-responsive tandem riboswitch (Fig. 5C and Supplementary Fig. S32). In the ligand-free state, basal hGSDMD-NT-2×T2 leakage remained indistinguishable from that of cells expressing the hGSDMD-NT-4A mutant—a well-characterized variant lacking membrane-perforating activity [57]. Upon ASP2905 administration, hGSDMD-NT-2×T2 induction drove a robust, dose-dependent increase in PI-positive cells, reaching levels comparable to those driven by constitutive expression (Fig. 5C and Supplementary Fig. S32). Crucially, ASP2905 treatment alone elicited no background cytotoxicity or pyroptotic phenotypes (Fig. 5C). Western blot analysis further confirmed the specific, ASP2905-dependent production of GSDMD-NT in cells transfected with hGSDMD-NT-2×T2 (Supplementary Fig. S33).

Crucially, despite the ultra-low basal leakage of the single-switch architecture—with dual-luciferase reporter assays revealing a baseline leakage of ∼1% under the CMV promoter (Supplementary Fig. S26) —transfection with hGSDMD-NT-T2 still triggered substantial ligand-independent pyroptosis (Supplementary Fig. S34). These results indicated that even trace amounts of GSDMD-NT suffice to drive membrane perforation, underscoring the extreme lethal potency of this effector and the imperative for absolute OFF-state silencing. Furthermore, when benchmarked against the gold-standard Tet-On 3G system, the tandem ASP2905-gated splicing switch displayed markedly lower basal leakage in the ligand-free state, whereas Tet-On 3G exhibited abundant background PI-positive cells. Upon ligand addition, both systems achieved comparable fractions of PI-positive cells (Supplementary Fig. S35). Modulating construct context by subcloning hGSDMD-NT-2×T2 from pcDNA3.1 into the pLVX lentiviral backbone further dampened baseline leakage. The high background mortality observed in uninduced Tet-On 3G controls likely stems from continuous, cumulative basal leakage over time (Supplementary Fig. S36). Strikingly, this persistent background lethality in doxycycline-inducible systems has been largely overlooked in prior studies employing Tet-On-driven GSDMD-NT [63,64]. Finally, this ASP2905-gated pyroptotic switch demonstrated robust generalizability across diverse human cancer cell lines, including HeLa, MCF-7, and A549 (Supplementary Fig. S37).

Together, these findings establish an all-RNA, traceless platform for ligand-gated pyroptosis, operating with stringent OFF-state silencing and high ON-state potency. By bypassing the inherent immunogenic and packaging constraints of heterologous systems, this modular framework provides vast translational potential—ranging from spatiotemporally precise cell ablation to programmable cancer immunotherapies.

## Discussion

SpliceRACER establishes a fully modular, protein-free RNA architecture for multiplexed gene regulation in mammalian systems. By eliminating reliance on exogenous protein trans-factors, this platform transcends the binary limitations of classic switches to enable multidimensional logic control. The seamless integration of three orthogonal aptamer–ligand pairs—acyclovir, ASP2905, and oxypurinol— validates the foundational principle that structurally divergent small molecules can orchestrate independent RNA circuits within a unified regulatory framework. This combinatorial expression manifold establishes an exponentially scalable route to high-dimensional proteome programming—transitioning isolated RNA switches into composable, higher-order gene networks and equipping synthetic biology with a versatile class of modular tools.

A central challenge in deploying such riboswitches with potent therapeutic effectors is the stringent requirement for OFF-state repression [27,49,65]. We identified spurious recognition of endogenous nucleoside pools by the acyclovir aptamer as the primary driver of basal leakiness. To overcome this, we engineered three general noise-reduction strategies. While tandem riboswitch arrays and CP-degron fusions effectively attenuated leakage, they typically imposed a penalty on maximal output. Integrating an ecDHFR destabilization domain establishes orthogonal, post-transcriptional and post-translational dual checkpoints, thereby decoupling the intrinsic trade-off between basal leakage and maximal output in synthetic gene circuits. This combinatorial design enables stringent, near-zero basal expression while preserving robust induction capacity—defining a precisely tunable operational window for applications ranging from precision genome editing to cell-based therapeutics.

Achieving traceless, protein-free regulation is paramount when deploying irreversible cytotoxic effectors like GSDMD-NT. Our data, consistent with previous studies, demonstrate that even trace amounts of GSDMD-NT suffice to drive membrane perforation [57,66], rendering post-translational degradation strategies insufficient due to kinetic hysteresis. By implementing tandem splicing arrays, we achieved OFF-state leakiness indistinguishable from that of an inactive mutant. Crucially, head-to-head benchmarking against the conventional Tet-On 3G system exposed cumulative background mortality in doxycycline-inducible platforms—a critical liability largely overlooked in prior reports [62–64]. Consequently, tandem splicing switches provide a safer, more reliable chassis for inducible cell ablation and next-generation cancer immunotherapies, while circumventing the host immunogenicity and AAV packaging bottlenecks inherent to trans-factor-dependent systems.

Beyond cell death control, the orthogonal SpliceRACER architecture is readily extensible to programmable cell-fate specification. Somatic cell reprogramming via defined transcription factors (Oct4, Sox2, Klf4, c-Myc) represents a classic paradigm for engineering cell fate [67,68]. Deploying SpliceRACER for the orthogonal, combinatorial regulation of core pluripotency factors (Oct4, Sox2, Klf4) thus establishes a powerful framework for chemically timed iPSC generation. Here, the stringent, near-zero baseline leakage of the tandem RNA architecture is particularly advantageous, as uncoordinated ectopic expression of reprogramming factors can trigger aberrant lineage differentiation or oncogenic transformation [69,70].

Despite its versatility, SpliceRACER currently relies on three distinct small-molecule ligands, and their high-dose combinatorial administration can induce off-target cytotoxicity. Future engineering should expand this ligand repertoire by identifying aptamers responsive to bio-orthogonal or FDA-approved small molecules. Furthermore, while SpliceRACER exhibits robust performance across diverse mammalian cell lines, *in vivo* evaluation in relevant animal models remains essential to fully realize its translational potential. Structural resolution of aptamer–ligand cross-reactivity will yield deeper insights into the chemical origin of basal leakiness, thereby guiding the rational design of next-generation, zero-leakage switches.

In summary, SpliceRACER transcends incremental optimization of conventional riboswitches, establishing a generalizable, multidimensional RNA framework. By interfacing orthogonal chemical inputs directly with host spliceosomal machinery, it sets a new benchmark for mammalian gene control—equipping the research community with a modular, all-RNA toolkit for engineering higher-order, programmable gene networks.

---

**Supplementary Figure 1.**
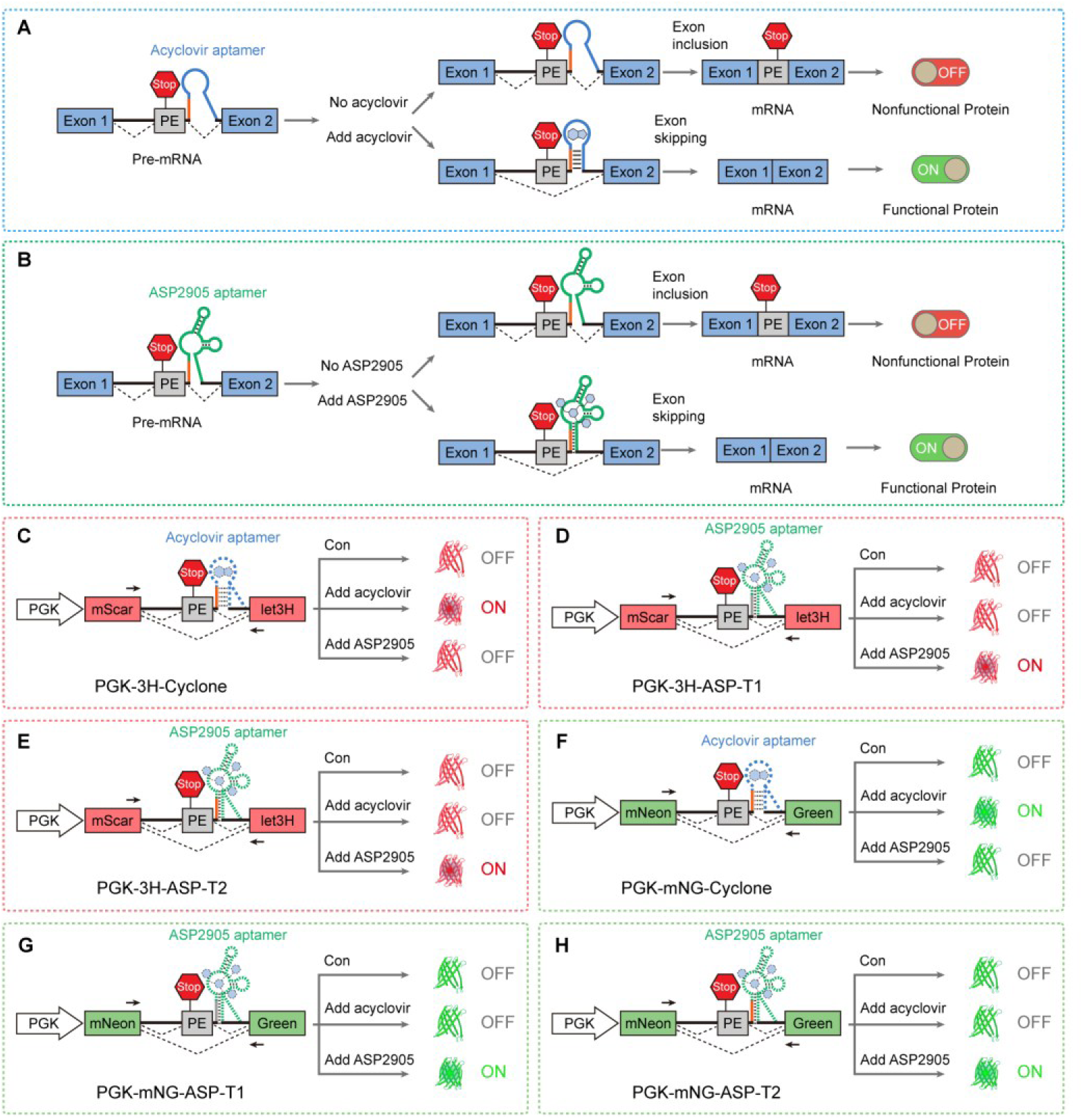
Design and mechanism of ligand-gated alternative splicing switches. (**A**, **B**) Schematic mechanism of ligand-gated poison exon (PE) splicing regulated by acyclovir (**A**) and ASP2905 (**B**). In the unliganded OFF state, the accessible 5′ splice site promotes U1 snRNP recognition, biasing splicing toward PE inclusion and yielding prematurely terminated, non-functional products. Ligand binding stabilizes the aptamer architecture, sterically occluding U1 access and enforcing PE skipping to restore full-length protein expression. (**C**–**H**) Small-molecule–dependent activation of reporter outputs. Acyclovir-dependent PE skipping drives mScarlet3H expression in (**C**) and mNeonGreen expression in (**F**). ASP2905-dependent PE skipping drives mScarlet3H expression in (**D**, **E**) and mNeonGreen expression in (**G**, **H**).

**Supplementary Figure 2.**
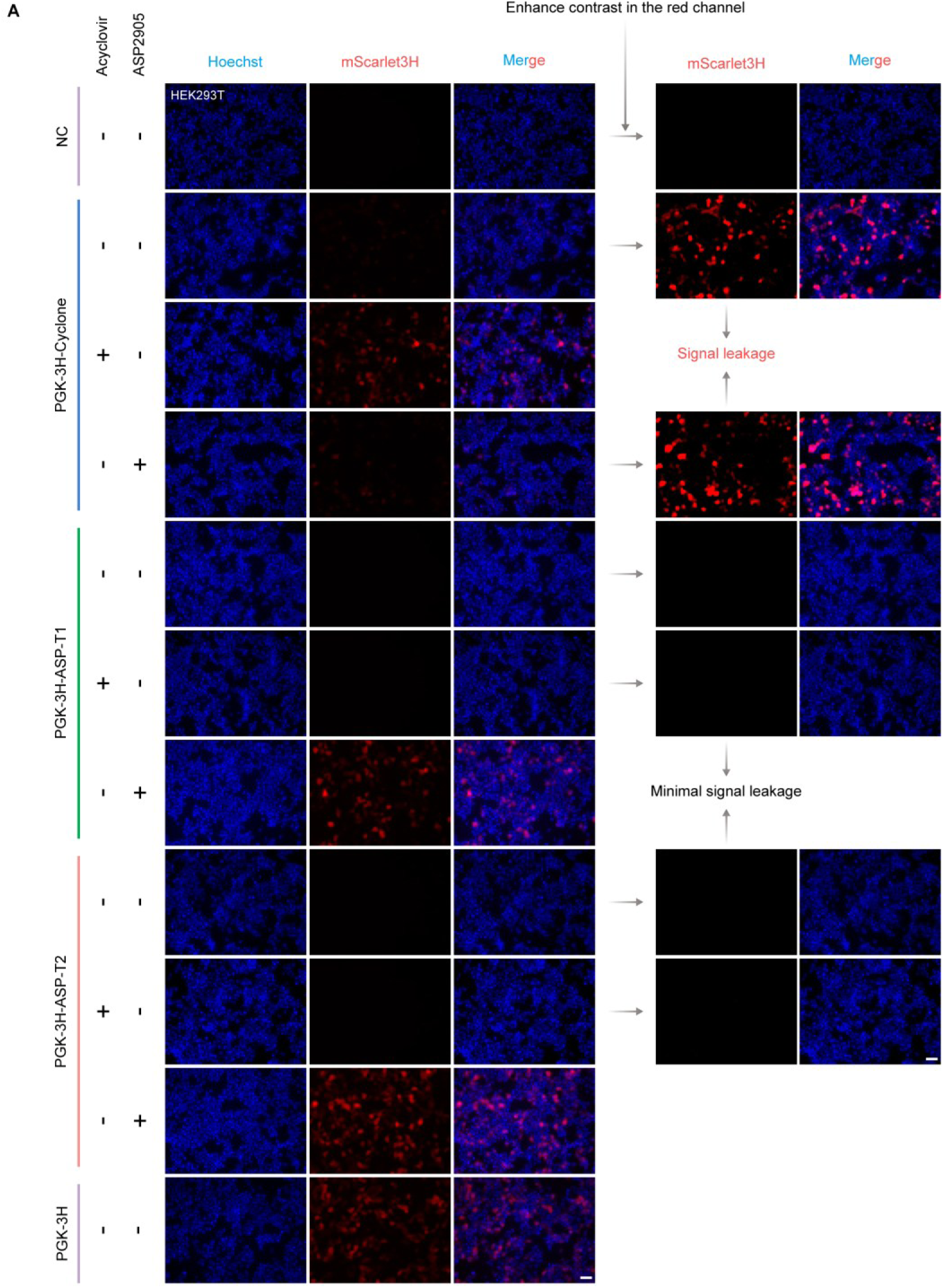
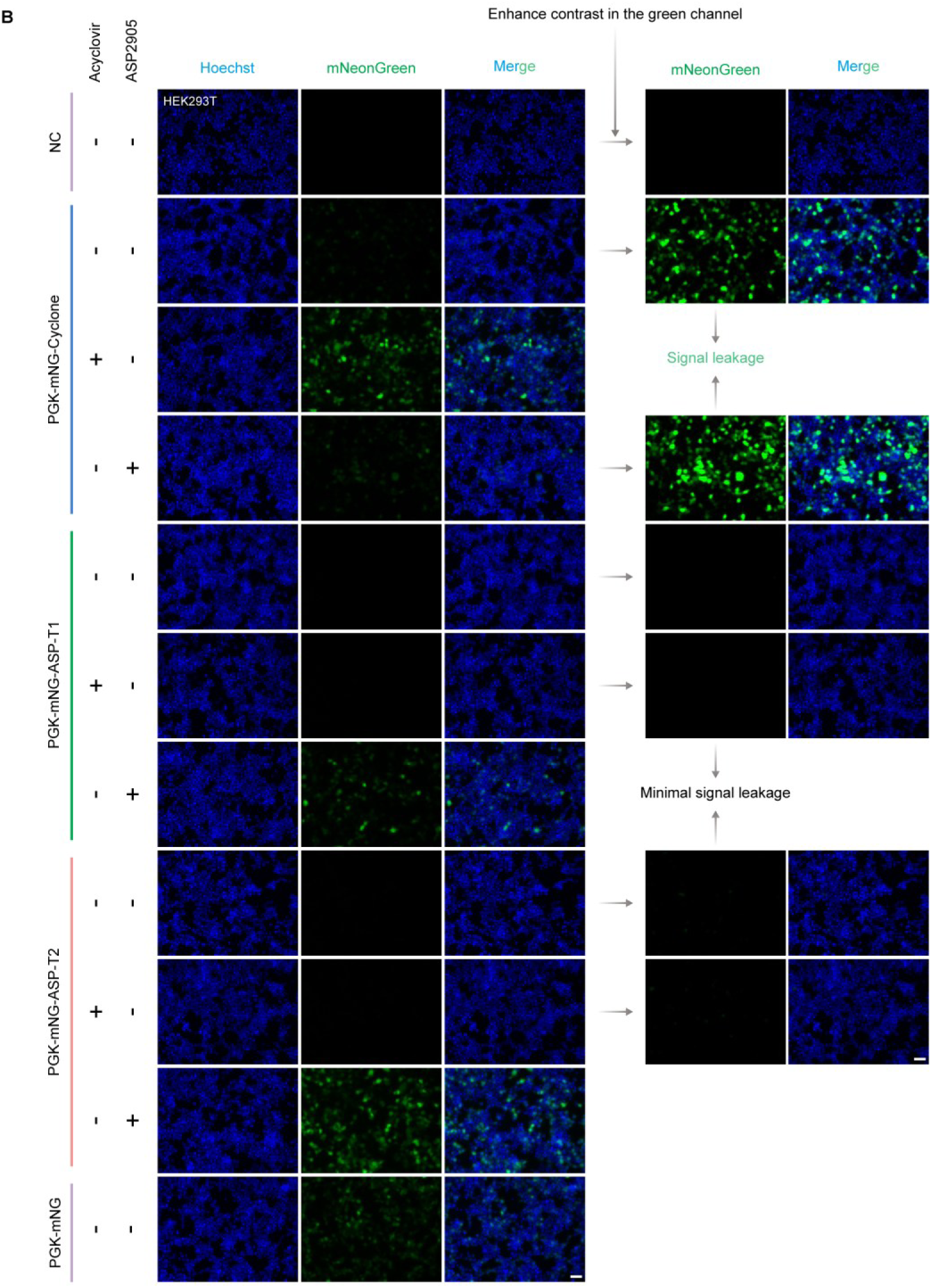
Basal expression leakiness and channel orthogonality of ligand-gated splicing switches. (**A**, **B**) Distinct basal leakiness of acyclovir- and ASP2905-gated circuits. Acyclovir-gated circuits exhibit pronounced ligand-free leakage, whereas ASP2905-gated circuits show negligible baseline activity, with no detectable inter-channel cross-talk. HEK293T cells were transfected with the indicated constructs and treated with 125 μM acyclovir or 10 μM ASP2905 for 24 h. Scale bars, 50 μm. Representative images from at least three independent biological replicates.

**Supplementary Figure 3.**
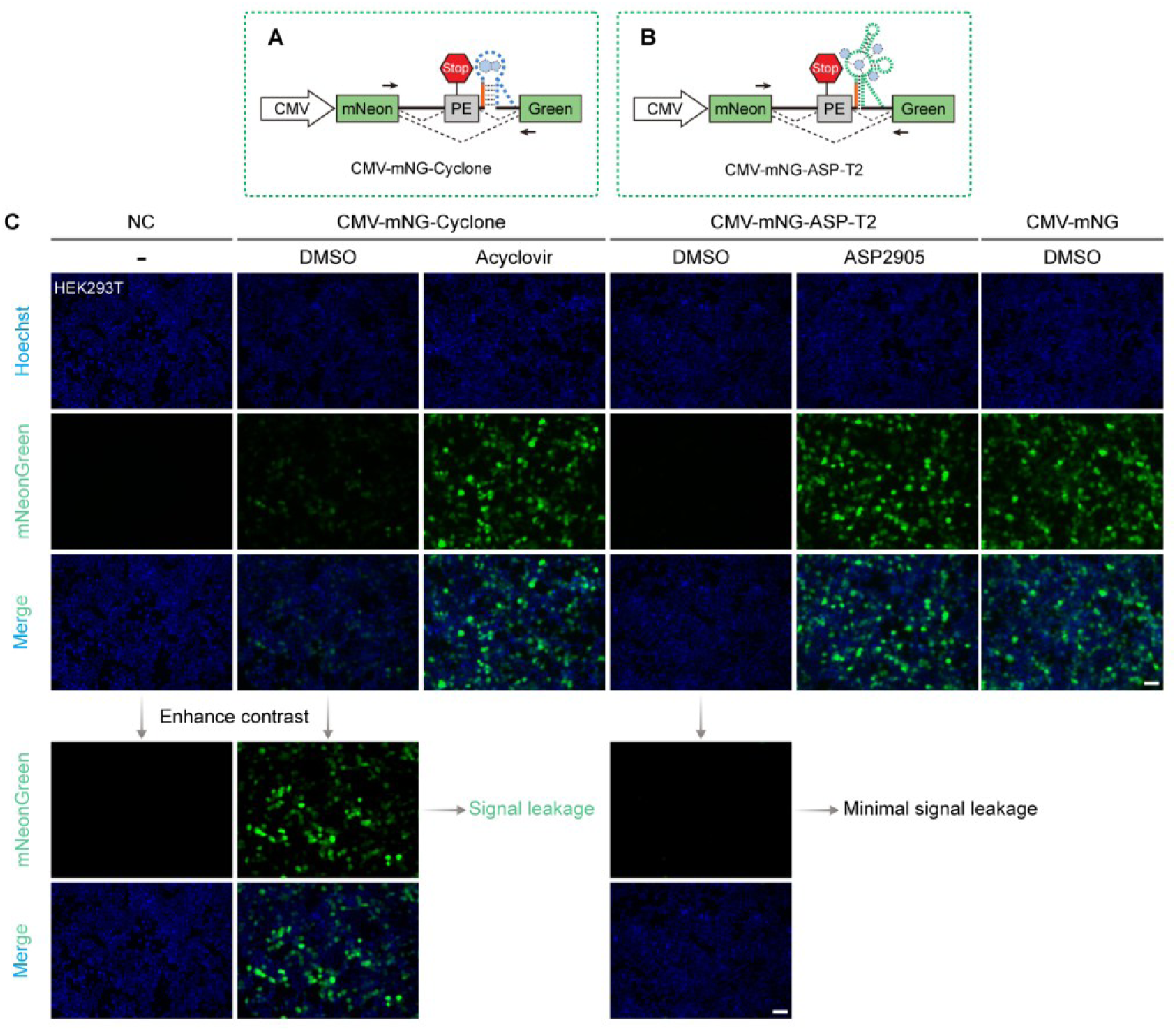
Performance comparison of CMV-driven ligand-gated splicing switches. (**A**, **B**) CMV-driven cassettes in which acyclovir and ASP2905 induce mNeonGreen expression via ligand-induced PE skipping. (**C**) Differential OFF-state stringency. Acyclovir-gated circuits exhibit pronounced ligand-free leakage, whereas ASP2905-gated circuits remain tightly repressed, yielding superior ON/OFF expression contrast. HEK293T cells were transfected with the indicated constructs and treated with 125 μM acyclovir or 10 μM ASP2905 for 24 h. Scale bars, 50 μm. Representative images from at least three independent biological replicates.

**Supplementary Figure 4.**
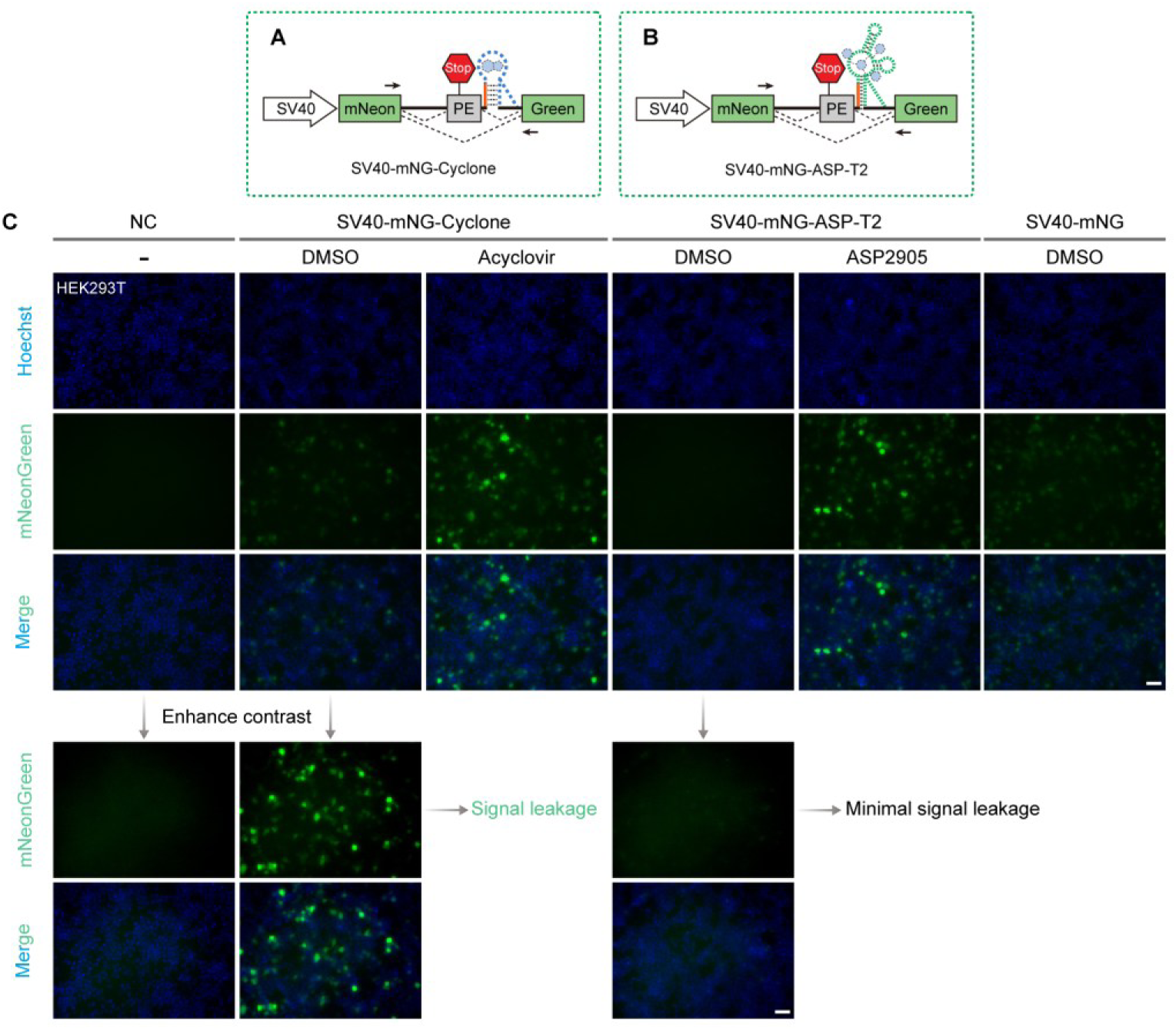
Performance comparison of SV40-driven ligand-gated splicing switches. (**A**, **B**) SV40-driven cassettes in which acyclovir and ASP2905 induce mNeonGreen expression via ligand-induced PE skipping. (**C**) Differential OFF-state stringency. Acyclovir-gated circuits exhibit pronounced ligand-free leakage, whereas ASP2905-gated circuits remain tightly repressed, yielding superior ON/OFF expression contrast. HEK293T cells were transfected with the indicated constructs and treated with 125 μM acyclovir or 10 μM ASP2905 for 24 h. Scale bars, 50 μm. Representative images from at least three independent biological replicates.

**Supplementary Figure 5.**
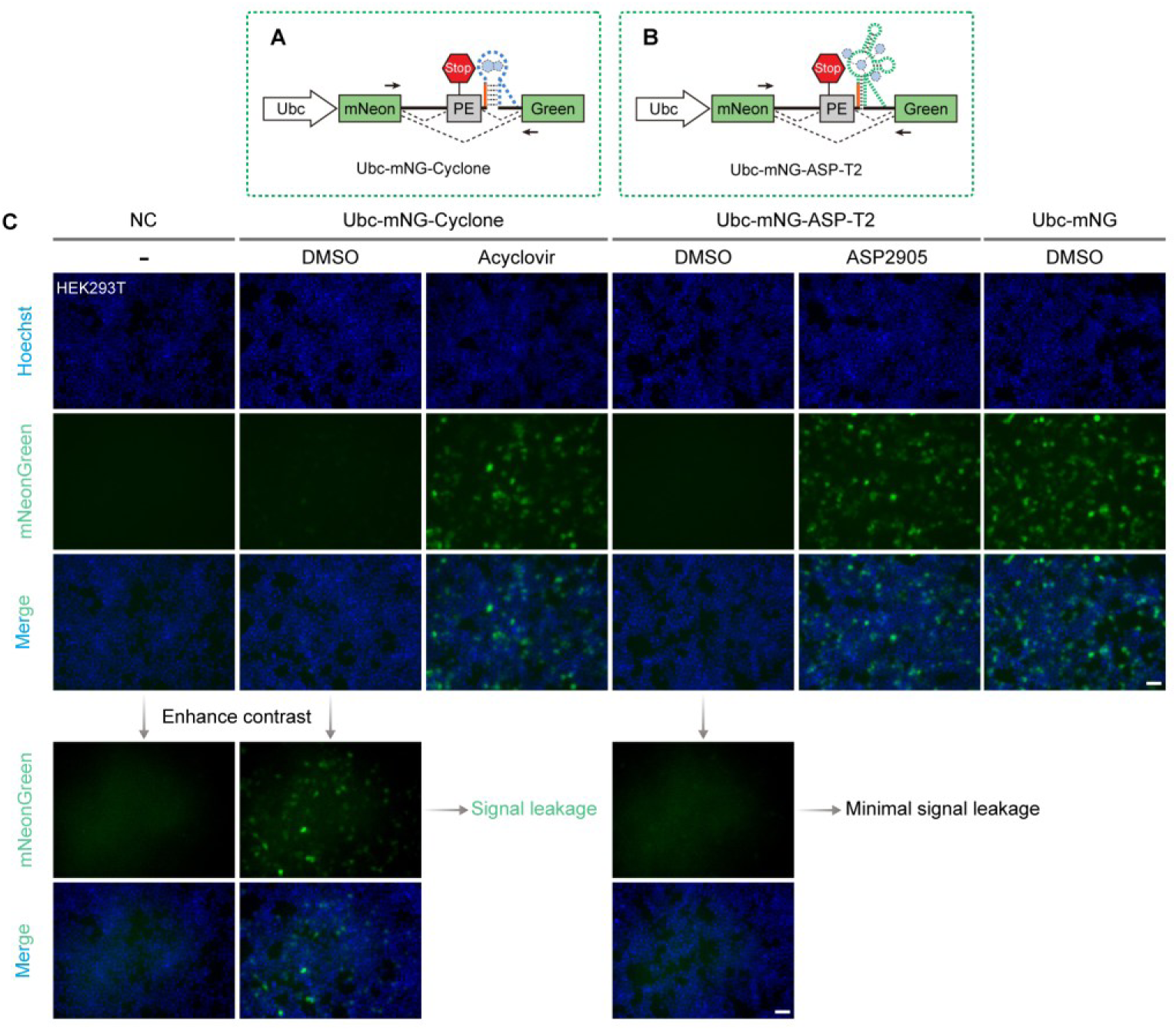
Performance comparison of Ubc-driven ligand-gated splicing switches. (**A**, **B**) Ubc-driven cassettes in which acyclovir and ASP2905 induce mNeonGreen expression via ligand-induced PE skipping. (**C**) Differential OFF-state stringency. Acyclovir-gated circuits exhibit pronounced ligand-free leakage, whereas ASP2905-gated circuits remain tightly repressed, yielding superior ON/OFF expression contrast. HEK293T cells were transfected with the indicated constructs and treated with 125 μM acyclovir or 10 μM ASP2905 for 24 h. Scale bars, 50 μm. Representative images from at least three independent biological replicates.

**Supplementary Figure 6.**
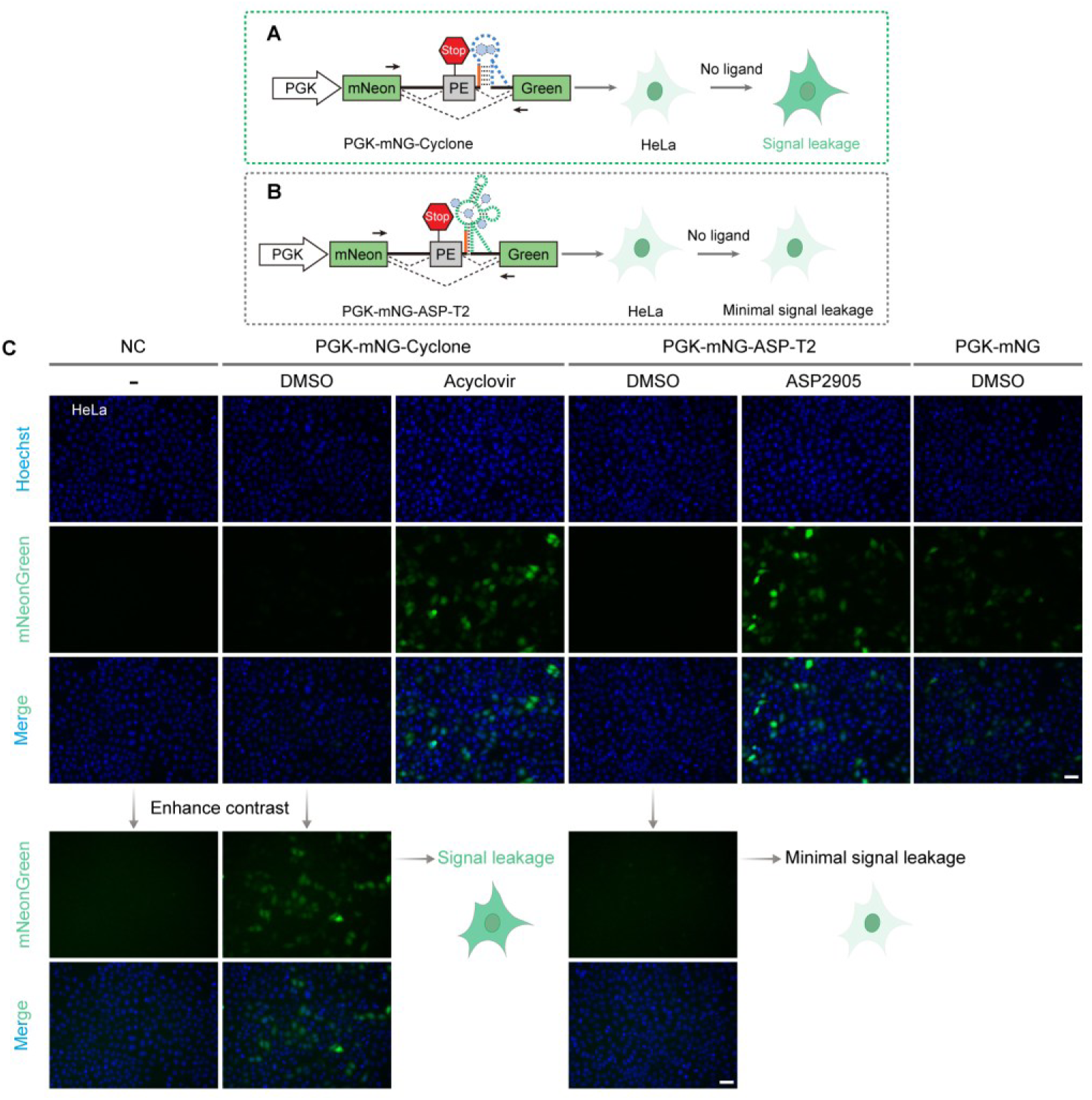
Performance comparison of ligand-gated splicing switches in HeLa cells. (**A**, **B**) PGK-driven expression cassettes in which acyclovir and ASP2905 induce mNeonGreen expression via ligand-induced PE skipping. (**C**) Differential OFF-state stringency. Acyclovir-gated circuits exhibit pronounced ligand-free leakage, whereas ASP2905-gated circuits remain tightly repressed, yielding superior ON/OFF expression contrast. HeLa cells were transfected with the indicated constructs and treated with 125 μM acyclovir or 10 μM ASP2905 for 24 h. Scale bars, 50 μm. Representative images from at least three independent biological replicates.

**Supplementary Figure 7.**
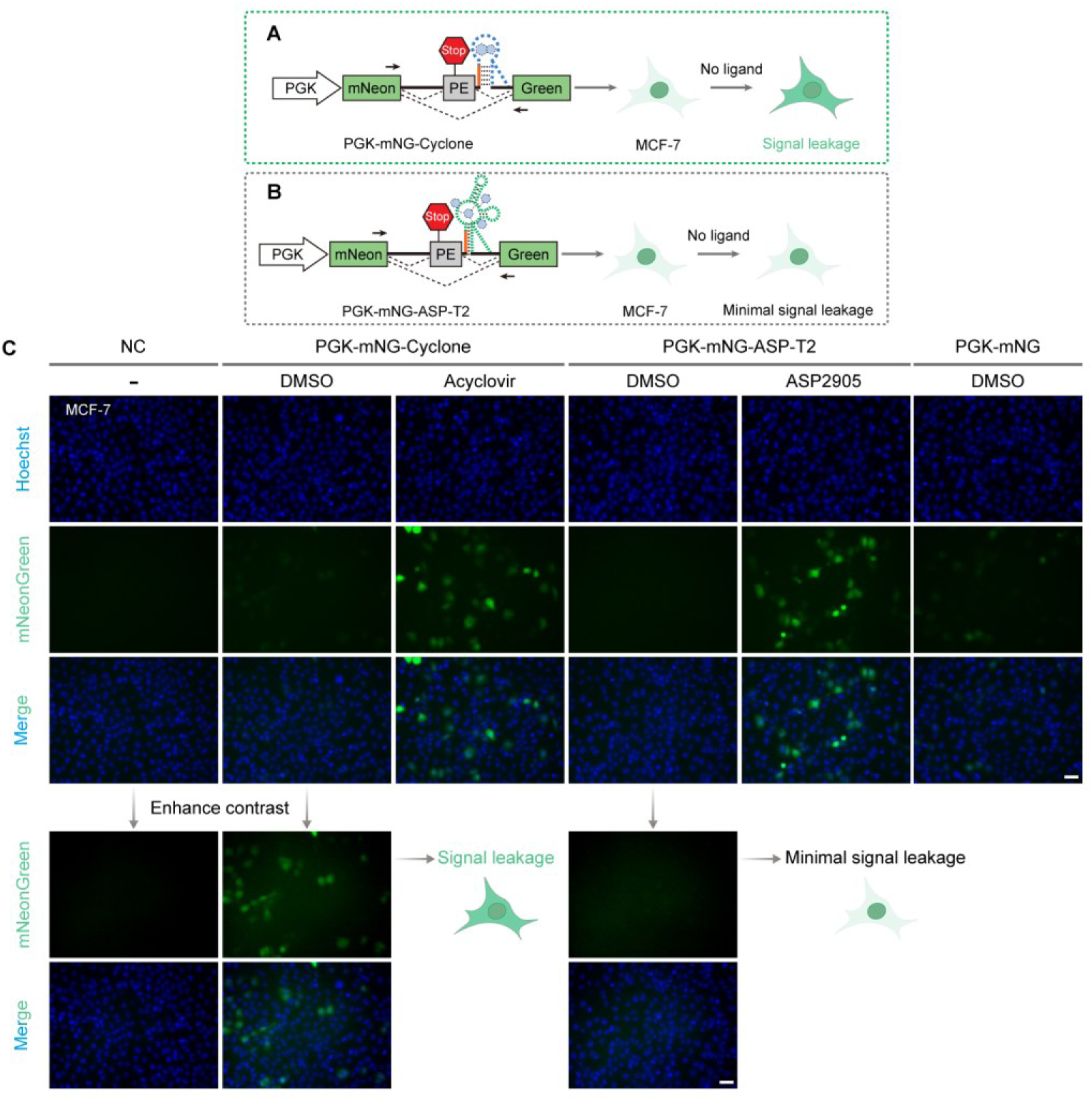
Performance comparison of ligand-gated splicing switches in MCF-7 cells. (**A**, **B**) PGK-driven expression cassettes in which acyclovir and ASP2905 induce mNeonGreen expression via ligand-induced PE skipping. (**C**) Differential OFF-state stringency. Acyclovir-gated circuits exhibit pronounced ligand-free leakage, whereas ASP2905-gated circuits remain tightly repressed, yielding superior ON/OFF expression contrast. MCF-7 cells were transfected with the indicated constructs and treated with 125 μM acyclovir or 10 μM ASP2905 for 24 h. Scale bars, 50 μm. Representative images from at least three independent biological replicates.

**Supplementary Figure 8.**
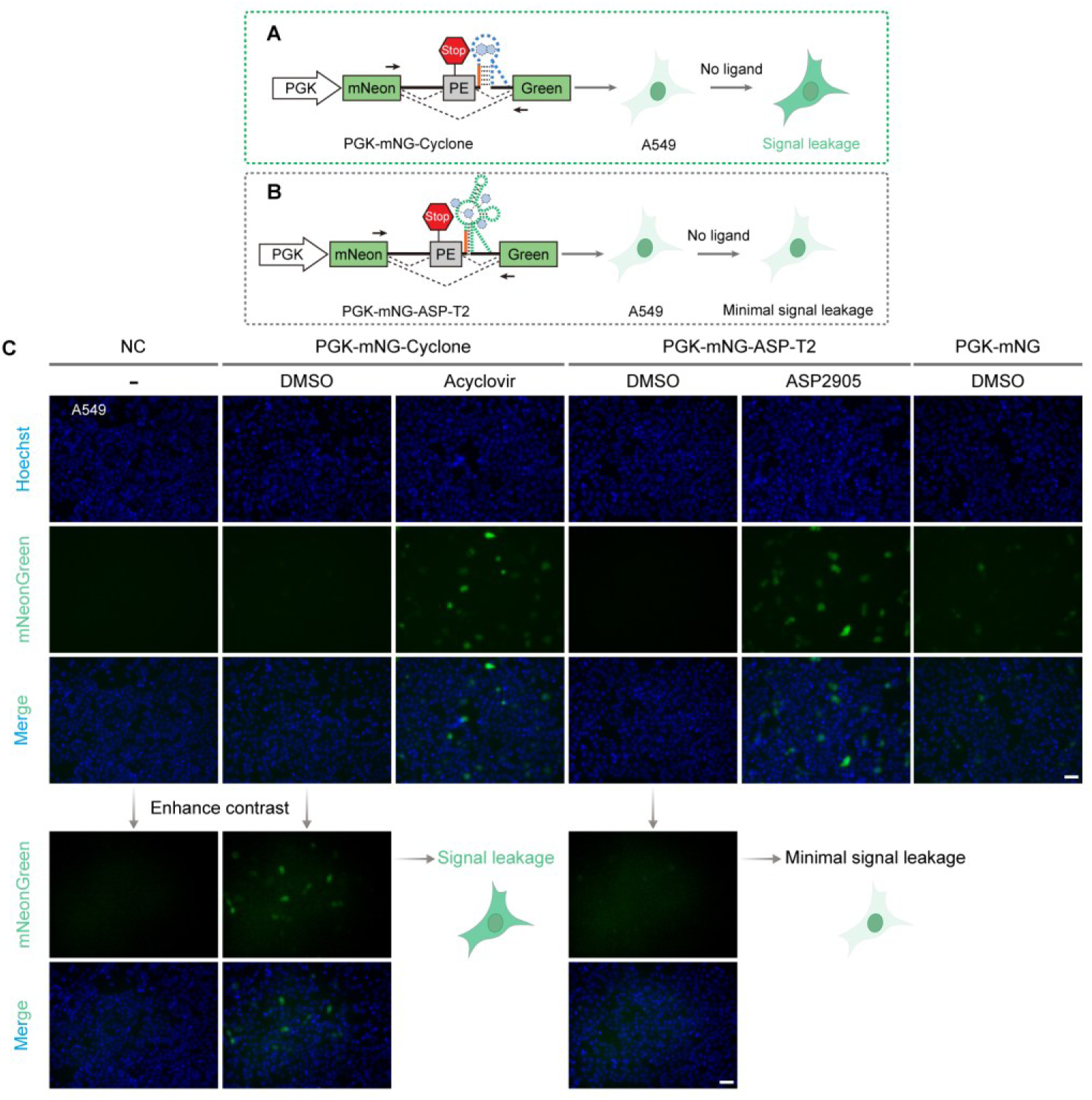
Performance comparison of ligand-gated splicing switches in A549 cells. (**A**, **B**) PGK-driven expression cassettes in which acyclovir and ASP2905 induce mNeonGreen expression via ligand-induced PE skipping. (**C**) Differential OFF-state stringency. Acyclovir-gated circuits exhibit pronounced ligand-free leakage, whereas ASP2905-gated circuits remain tightly repressed, yielding superior ON/OFF expression contrast. A549 cells were transfected with the indicated constructs and treated with 125 μM acyclovir or 10 μM ASP2905 for 24 h. Scale bars, 50 μm. Representative images from at least three independent biological replicates.

**Supplementary Figure 9.**
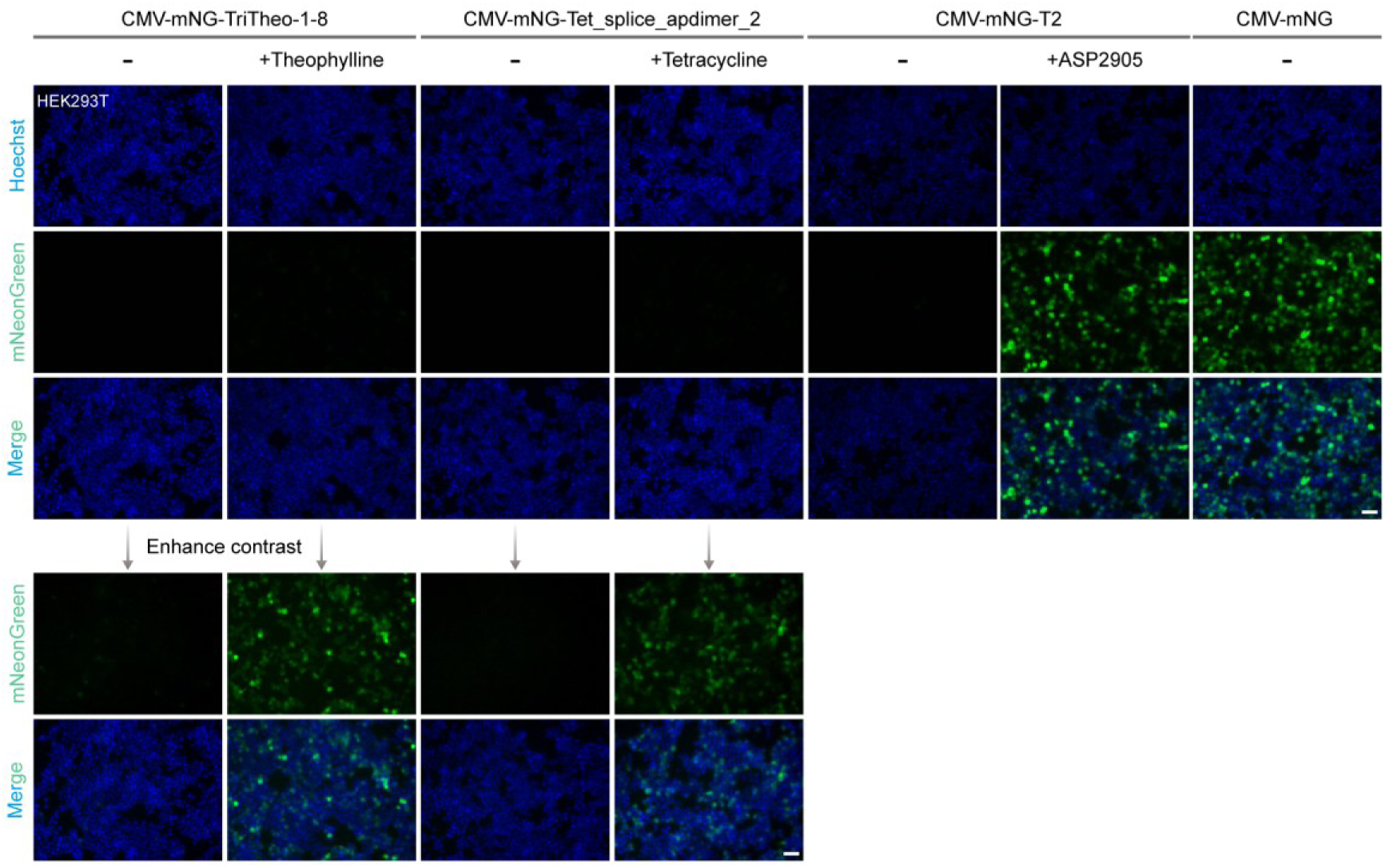
Head-to-head comparison with existing small-molecule– gated splicing switches. In the ON state, classical theophylline-and tetracycline-gated switches showed barely detectable expression, whereas ASP2905-gated switches achieved expression levels comparable to the constitutive control. HEK293T cells were transfected with the indicated constructs and treated with cognate ligands (1 mM theophylline, 50 μM tetracycline, or 10 μM ASP2905) for 24 h. Scale bars, 50 μm. Representative images from at least three independent biological replicates.

**Supplementary Figure 10.**
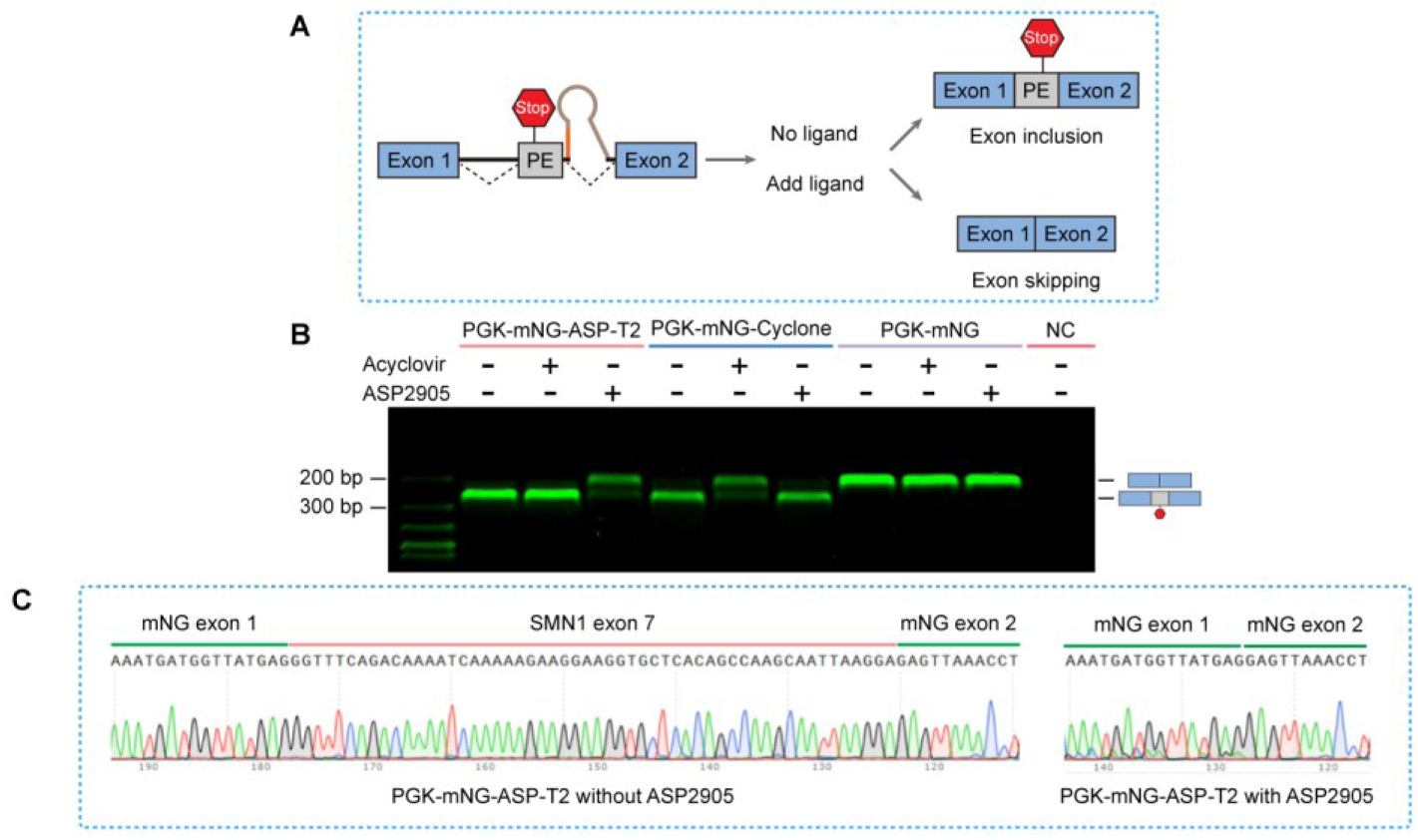
Molecular basis of ligand-gated splicing specificity. (**A**) Ligand-induced conformational switching governs splice-site accessibility. Binding of acyclovir or ASP2905 to their cognate aptamers stabilizes the RNA architecture, thereby gating 5′ splice site accessibility to enforce either PE inclusion (lower band, OFF state) or PE skipping (upper band, ON state). (**B**) RT–PCR analysis of alternative splicing isoforms. HEK293T cells transfected with the indicated constructs were treated with 125 μM acyclovir or 10 μM ASP2905 for 24 h. Total RNA was extracted, reverse-transcribed, and PCR-amplified, with splicing isoforms resolved by 3% agarose gel electrophoresis. (**C**) Base-resolution Sanger sequencing chromatograms confirming precise splice-junction fidelity across ON and OFF states.

**Supplementary Figure 11.**
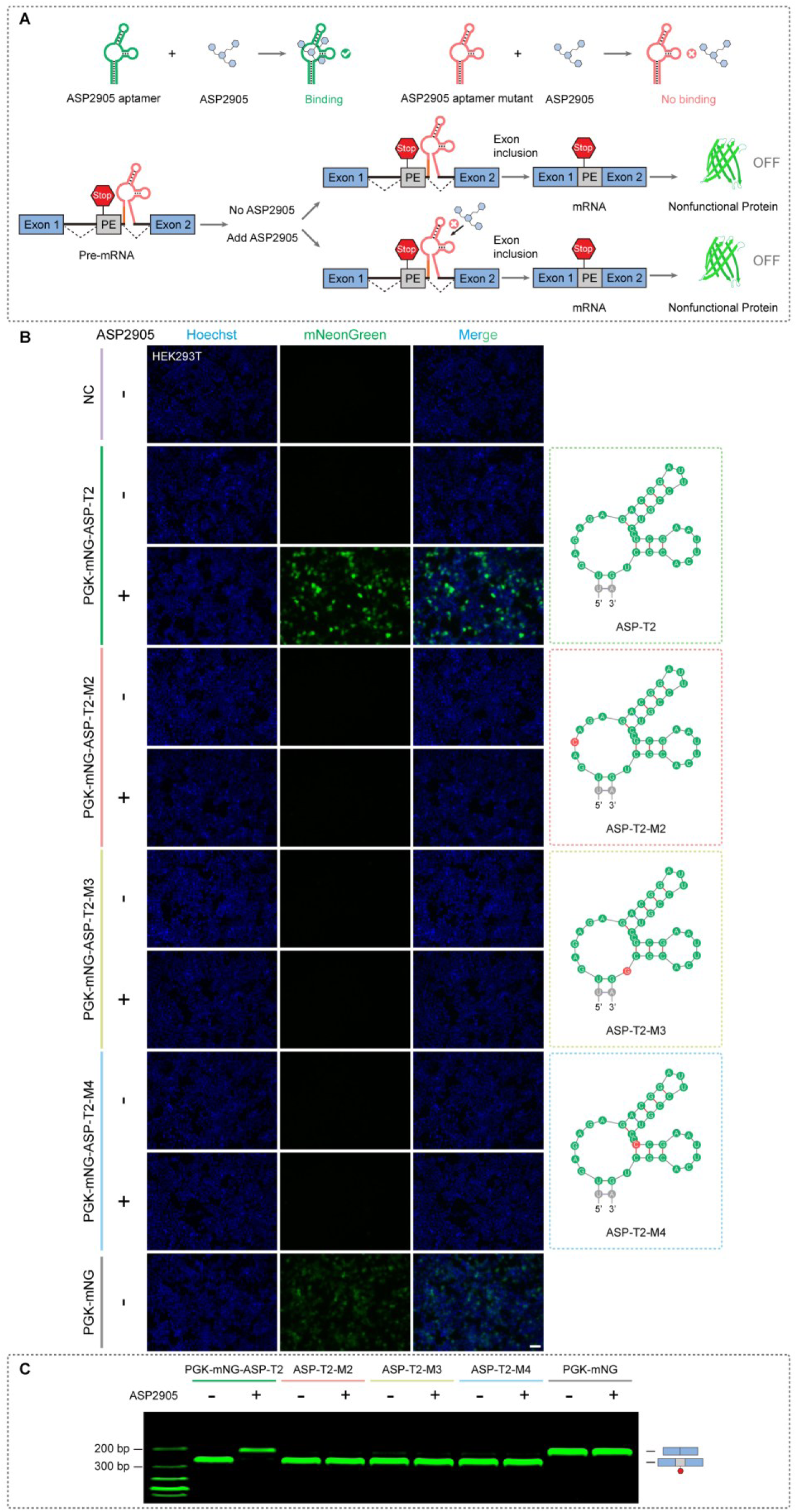
Structural integrity of the ASP2905 aptamer is essential for ligand-gated splicing control. (**A**) Single-nucleotide variants (SNVs) disrupt the ASP2905-binding pocket, abrogating ligand-binding affinity and responsiveness. (**B**) SNVs abolish ASP2905-induced mNeonGreen expression. HEK293T cells transfected with the indicated SNV constructs were treated with or without 10 μM ASP2905 for 24 h; red circles denote mutation sites. Scale bars, 50 μm. (**C**) RT–PCR analysis demonstrates constitutive PE inclusion and complete loss of ligand-induced PE skipping across all mutants. Total RNA from transfected cells treated with or without 10 μM ASP2905 was extracted, reverse-transcribed, and PCR-amplified, with products resolved on a 3% agarose gel, revealing PE-included (lower band, OFF state) and PE-skipped (upper band, ON state) bands. Representative images from at least three independent biological replicates.

**Supplementary Figure 12.**
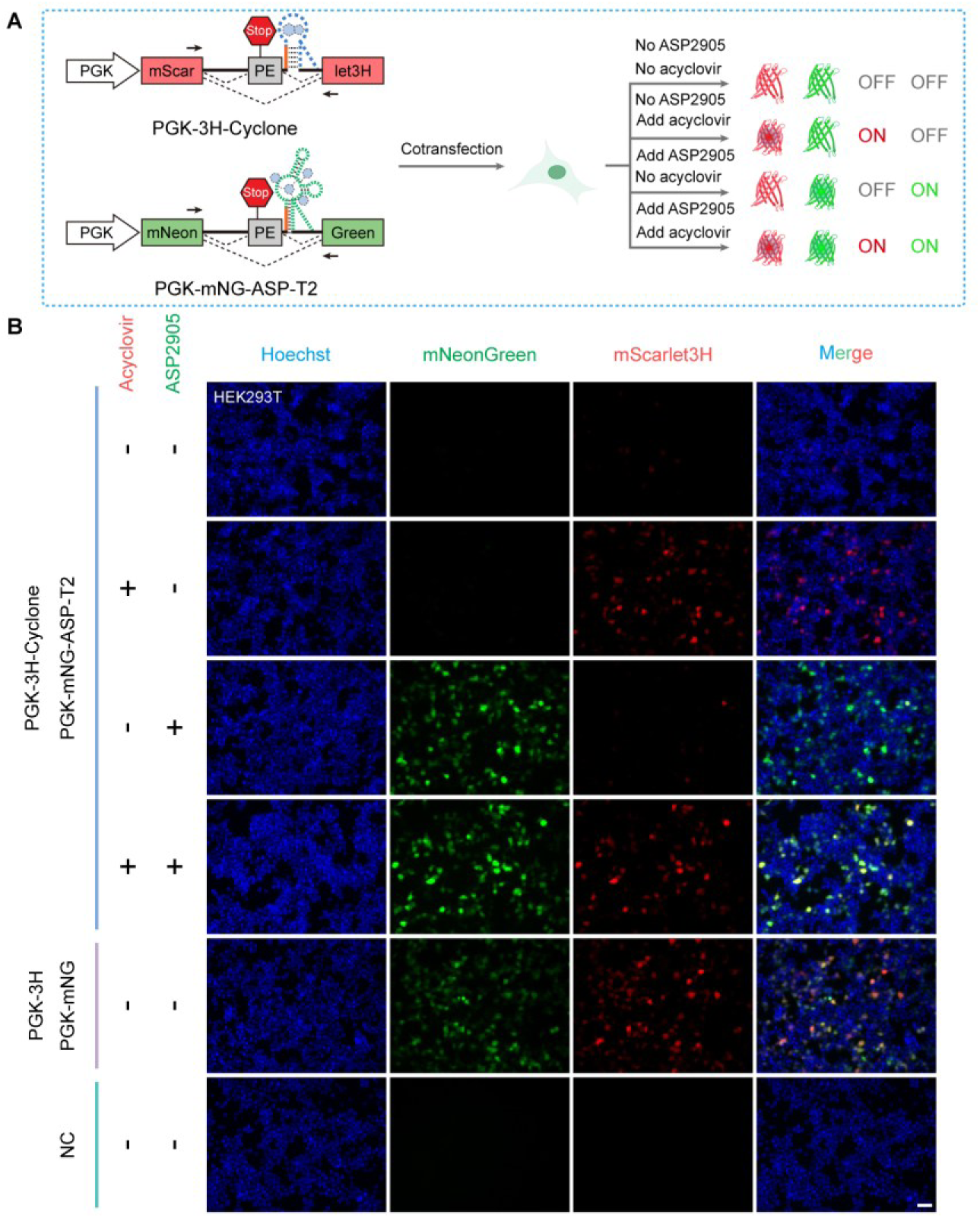
Orthogonal dual-channel regulation of distinct reporter genes by small-molecule–gated splicing switches. (**A**) Schematic of orthogonal dual-channel control. Acyclovir and ASP2905 bind their respective cognate aptamers to independently enforce PE skipping in mScarlet3H and mNeonGreen expression cassettes. (**B**) Dual-color live-cell imaging demonstrating channel orthogonality. HEK293T cells co-expressing both reporter constructs were treated with vehicle control (DMSO), 125 μM acyclovir, 10 μM ASP2905, or the two-ligand combination for 24 h. Acyclovir selectively activates mScarlet3H expression without inducing mNeonGreen, whereas ASP2905 specifically triggers mNeonGreen without cross-activating mScarlet3H, yielding spectrally resolvable, mutually exclusive outputs with negligible inter-channel cross-talk. Scale bars, 50 μm. Representative images from at least three independent biological replicates.

**Supplementary Figure 13.**
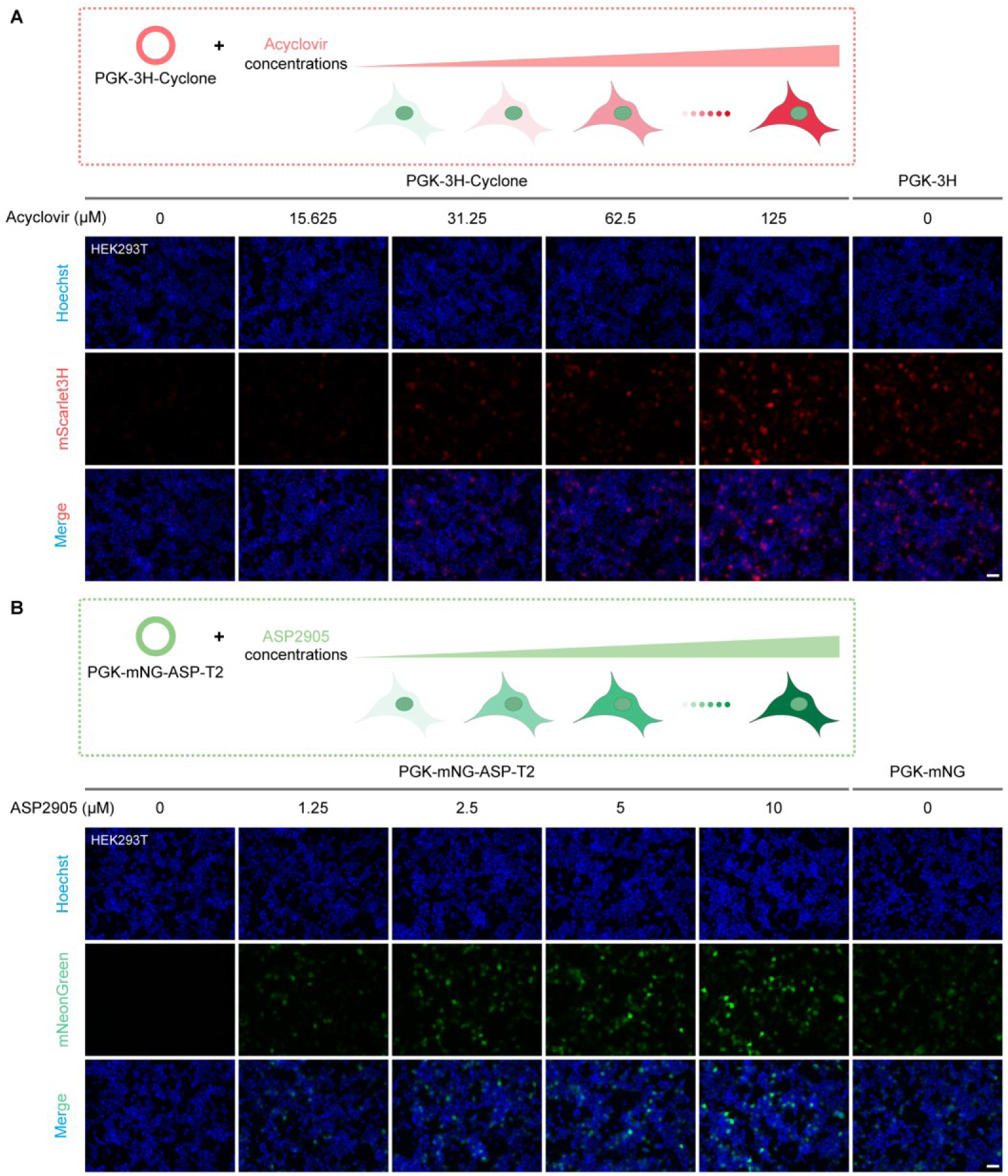
Dose-dependent tunability of ligand-gated splicing switches. (**A**) Acyclovir induces mScarlet3H expression in a concentration-dependent manner across a wide dynamic range. (**B**) ASP2905 enables precise, dose-responsive titration of mNeonGreen expression. HEK293T cells transfected with the indicated constructs were treated with gradient concentrations of acyclovir or ASP2905 for 24 h. Scale bars, 50 μm. Representative images from at least three independent biological replicates.

**Supplementary Figure 14.**
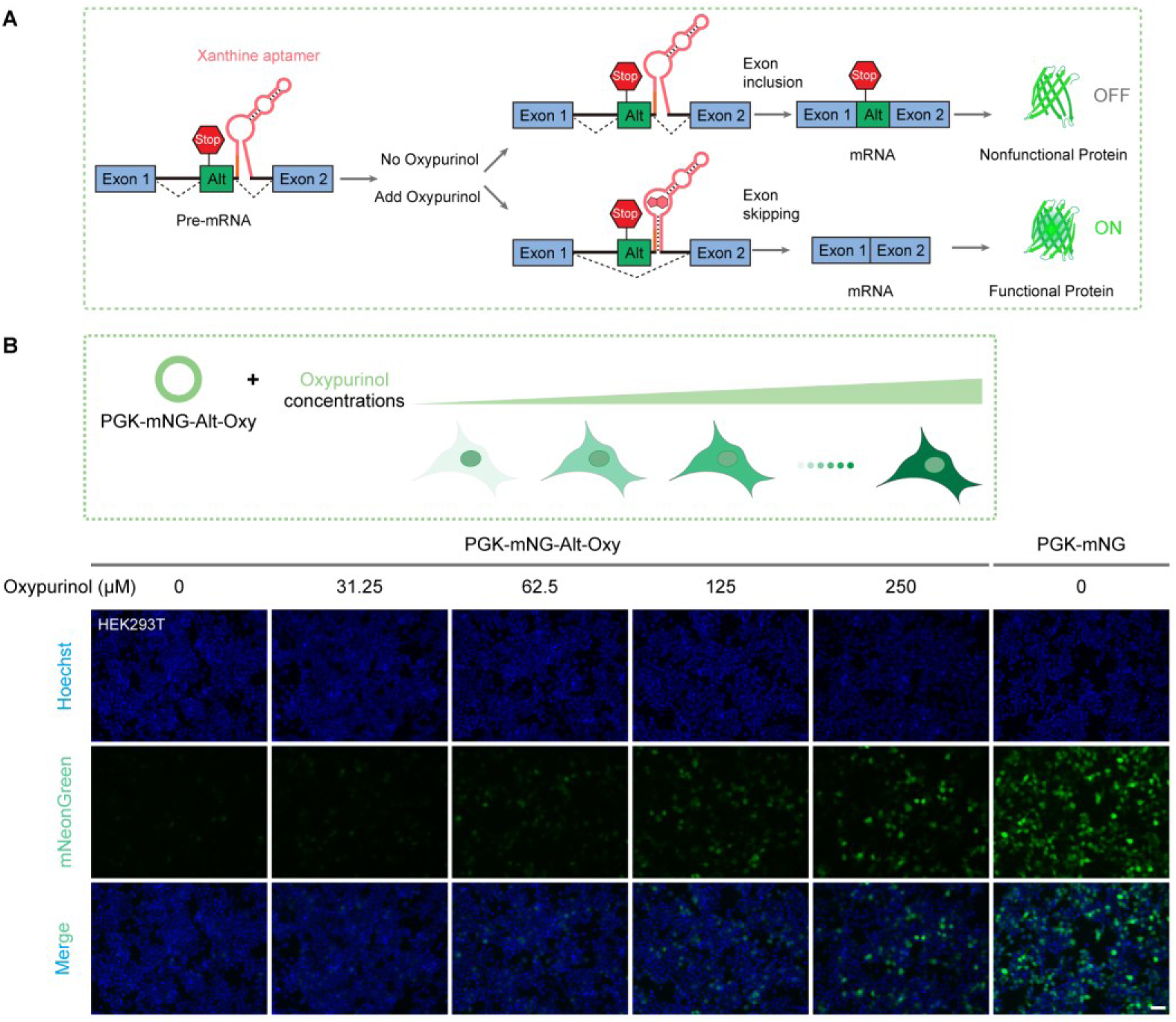
Ligand-tunable splicing switch with oxypurinol. (**A**) Schematic of the oxypurinol-responsive expression cassette driving ligand-induced PE skipping. (**B**) Dose-responsive induction of mNeonGreen expression by oxypurinol. HEK293T cells transfected with the indicated constructs were treated with gradient concentrations of oxypurinol for 24 h. Scale bars, 50 μm. Representative images from at least three independent biological replicates.

**Supplementary Figure 15.**
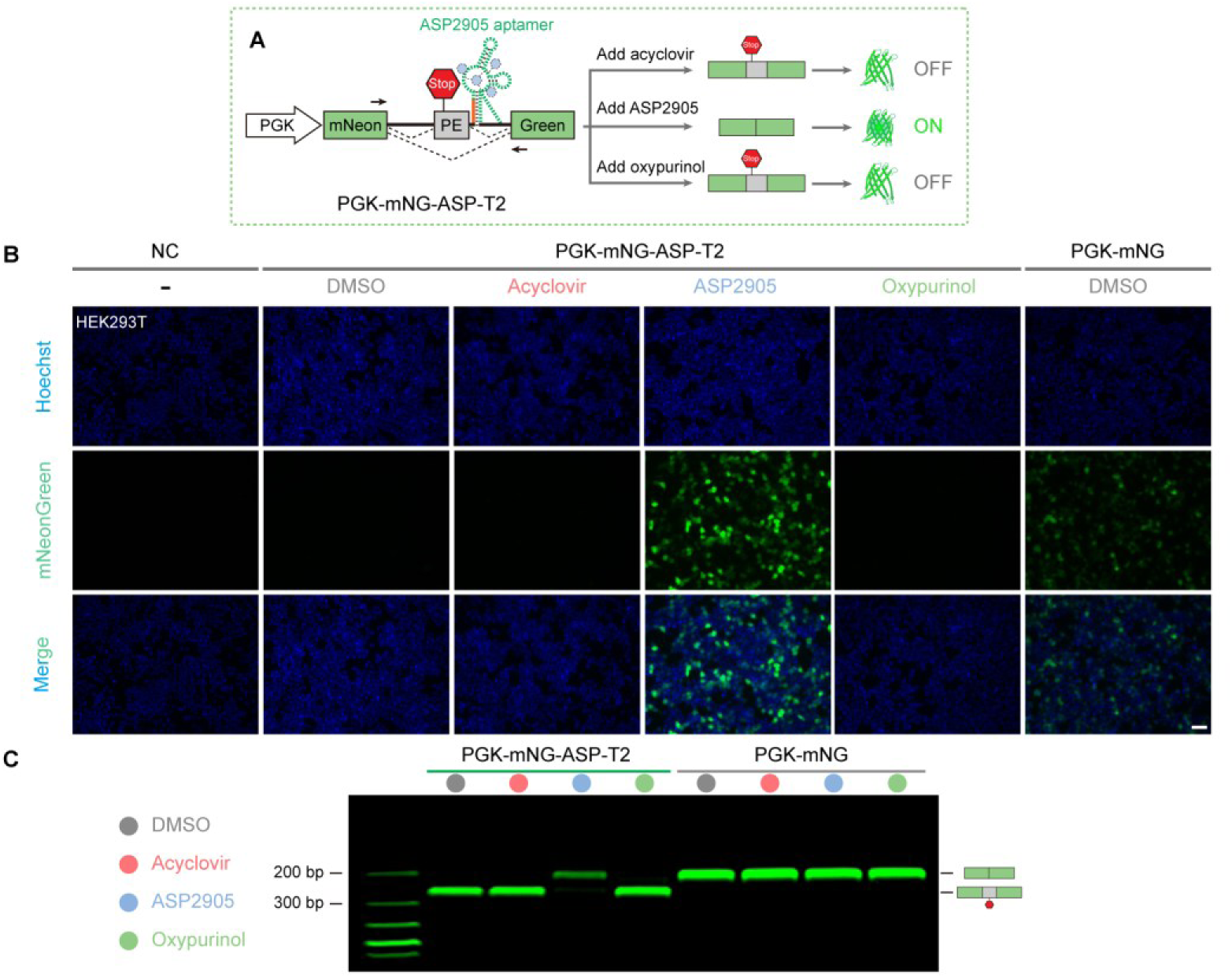
Ligand selectivity of the ASP2905-gated splicing switch. (**A**) Schematic of aptamer ligand selectivity. ASP2905 binding drives PE skipping (ON state), whereas acyclovir and oxypurinol fail to bind the ASP2905 aptamer, thereby retaining constitutive PE inclusion (OFF state). (**B**) Ligand specificity in live cells. HEK293T cells transfected with the ASP2905-gated reporter construct were treated with 125 μM acyclovir, 10 μM ASP2905, or 250 μM oxypurinol for 24 h. Only ASP2905 robustly induces mNeonGreen expression, whereas off-target ligands produce no detectable signal. Scale bars, 50 μm. (**C**) RT–PCR analysis confirming exclusive ligand selectivity. Total RNA was extracted, reverse-transcribed, and PCR-amplified, with splicing isoforms resolved by 3% agarose gel electrophoresis. ASP2905 specifically triggers PE skipping (upper band, ON state), whereas acyclovir and oxypurinol retain the PE-inclusion profile (lower band, OFF state). Representative images from at least three independent biological replicates.

**Supplementary Figure 16.**
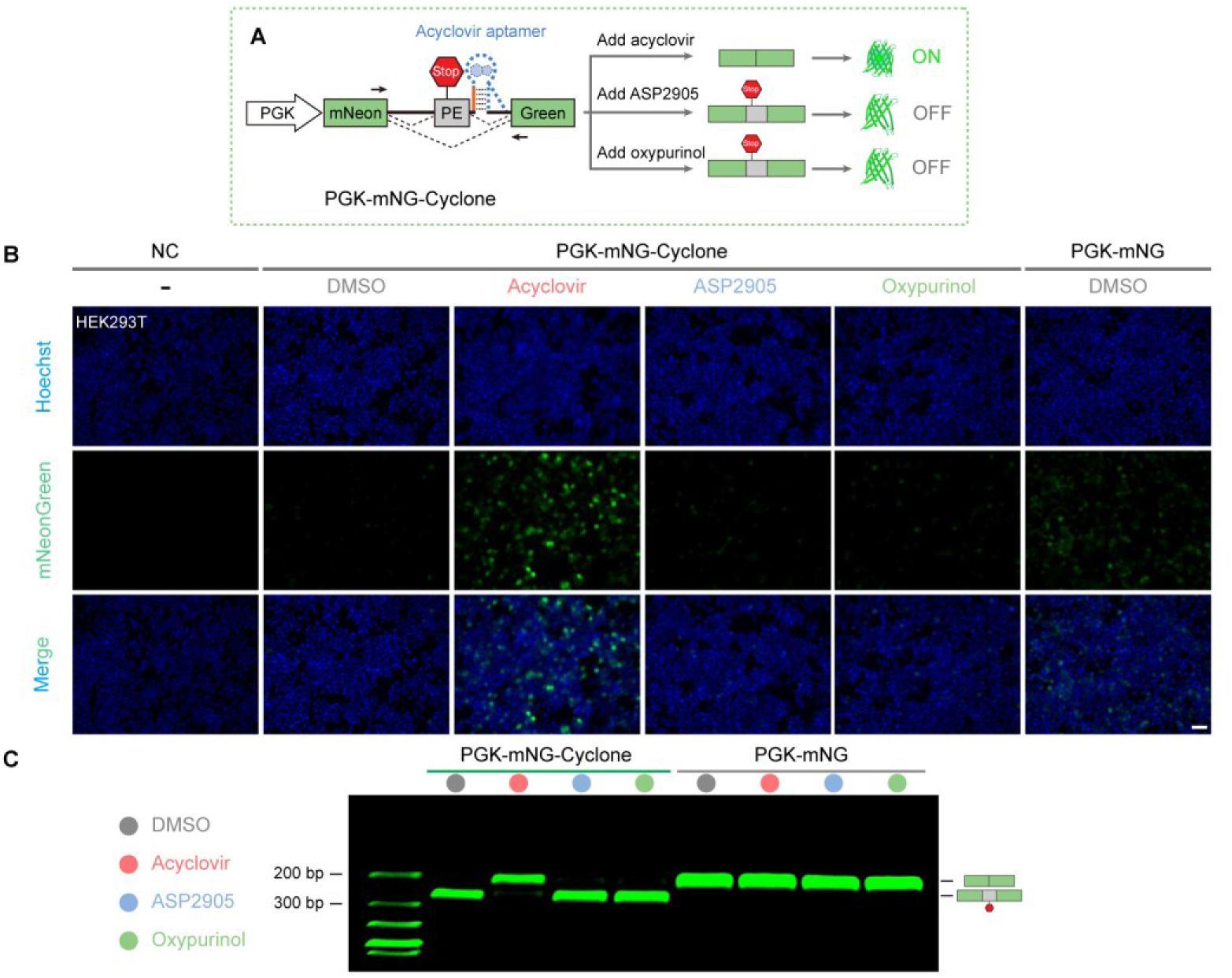
Ligand selectivity of the acyclovir-gated splicing switch. (**A**) Schematic of aptamer ligand selectivity. Acyclovir binding drives PE skipping (ON state), whereas ASP2905 and oxypurinol fail to bind the acyclovir aptamer, thereby retaining constitutive PE inclusion (OFF state). (**B**) Ligand specificity in live cells. HEK293T cells transfected with the acyclovir-gated reporter construct were treated with 125 μM acyclovir, 10 μM ASP2905, or 250 μM oxypurinol for 24 h. Only acyclovir robustly induces mNeonGreen expression, whereas off-target ligands produce no detectable signal. Scale bars, 50 μm. (**C**) RT–PCR analysis confirming exclusive ligand selectivity. Acyclovir specifically triggers PE skipping (upper band, ON state), whereas ASP2905 and oxypurinol retain the PE-inclusion profile (lower band, OFF state). Representative images from at least three independent biological replicates.

**Supplementary Figure 17.**
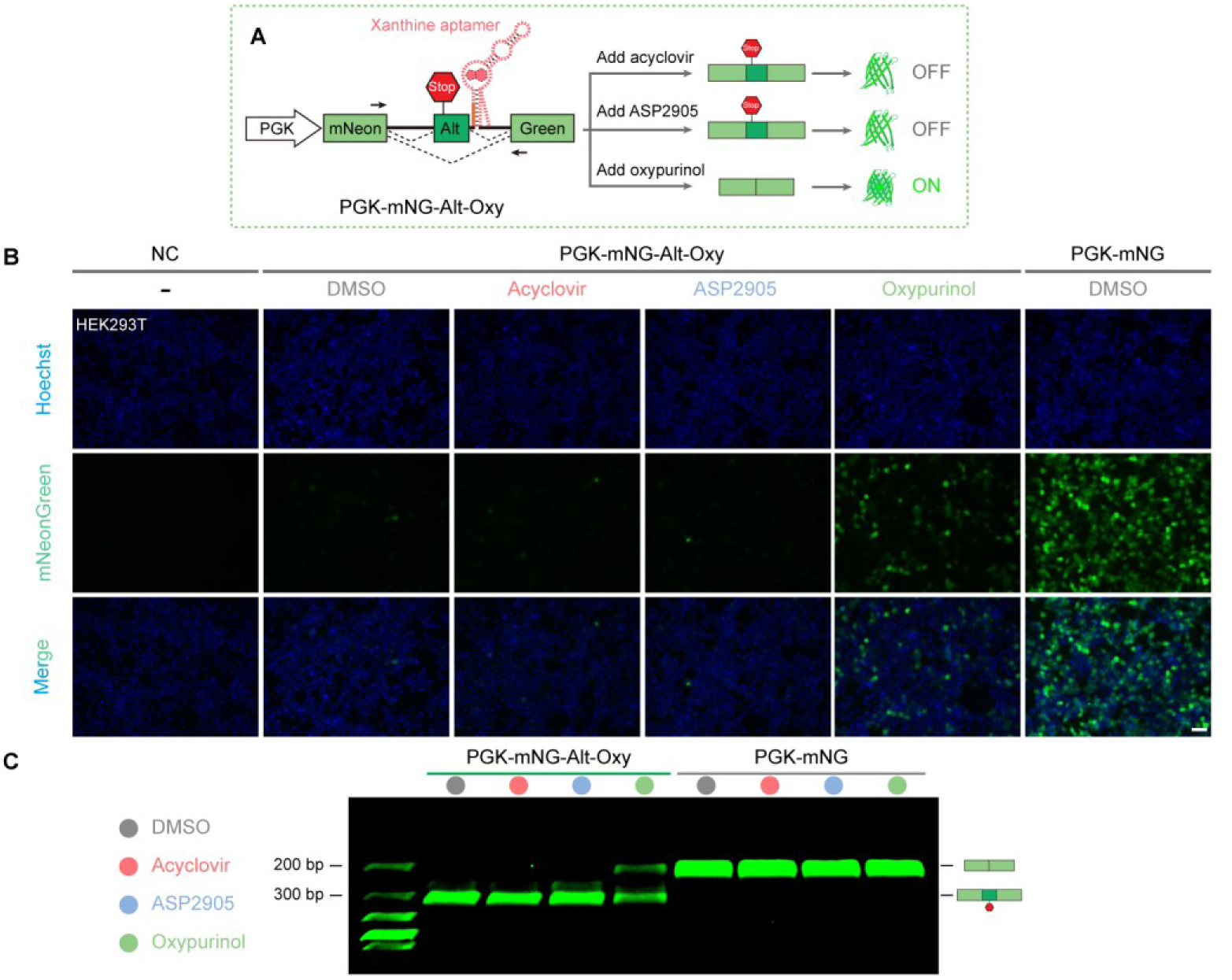
Ligand selectivity of the oxypurinol-gated splicing switch. (**A**) Schematic of aptamer ligand selectivity. Oxypurinol binding drives PE skipping (ON state), whereas ASP2905 and acyclovir fail to bind the oxypurinol aptamer, thereby retaining constitutive PE inclusion (OFF state). (**B**) Ligand specificity in live cells. HEK293T cells transfected with the oxypurinol-gated reporter construct were treated with 125 μM acyclovir, 10 μM ASP2905, or 250 μM oxypurinol for 24 h. Only oxypurinol robustly induces mNeonGreen expression, whereas off-target ligands produce no detectable signal. Scale bars, 50 μm. (**C**) RT–PCR analysis confirming exclusive ligand selectivity. Oxypurinol specifically triggers PE skipping (upper band, ON state), whereas ASP2905 and acyclovir retain the PE-inclusion profile (lower band, OFF state). Representative images from at least three independent biological replicates.

**Supplementary Figure 18.**
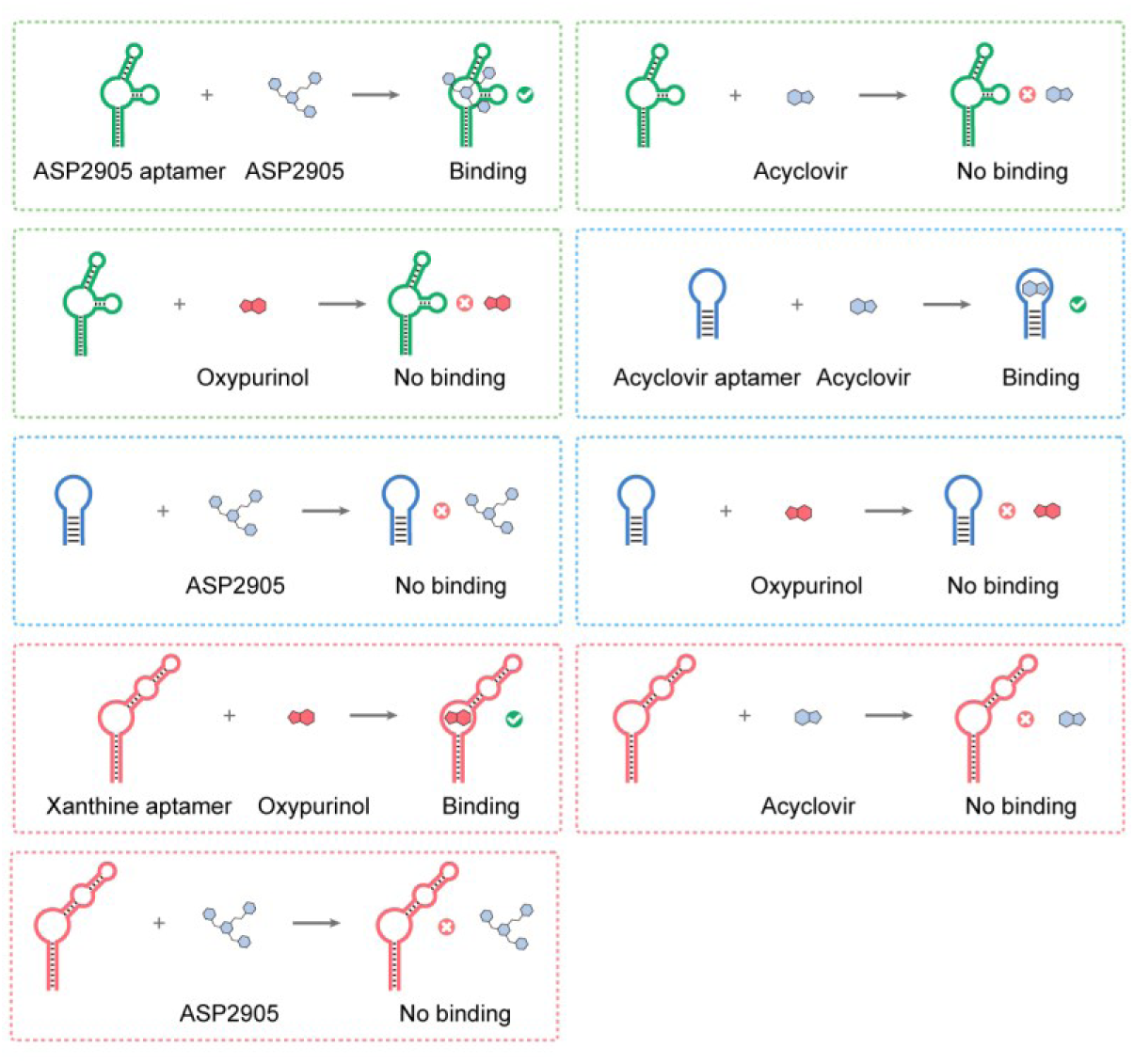
Orthogonal ligand–aptamer pairings. Schematic depicting the exclusive pairing between each ligand and its cognate aptamer, establishing the molecular basis for cross-talk-free orthogonal splicing control.

**Supplementary Figure 19.**
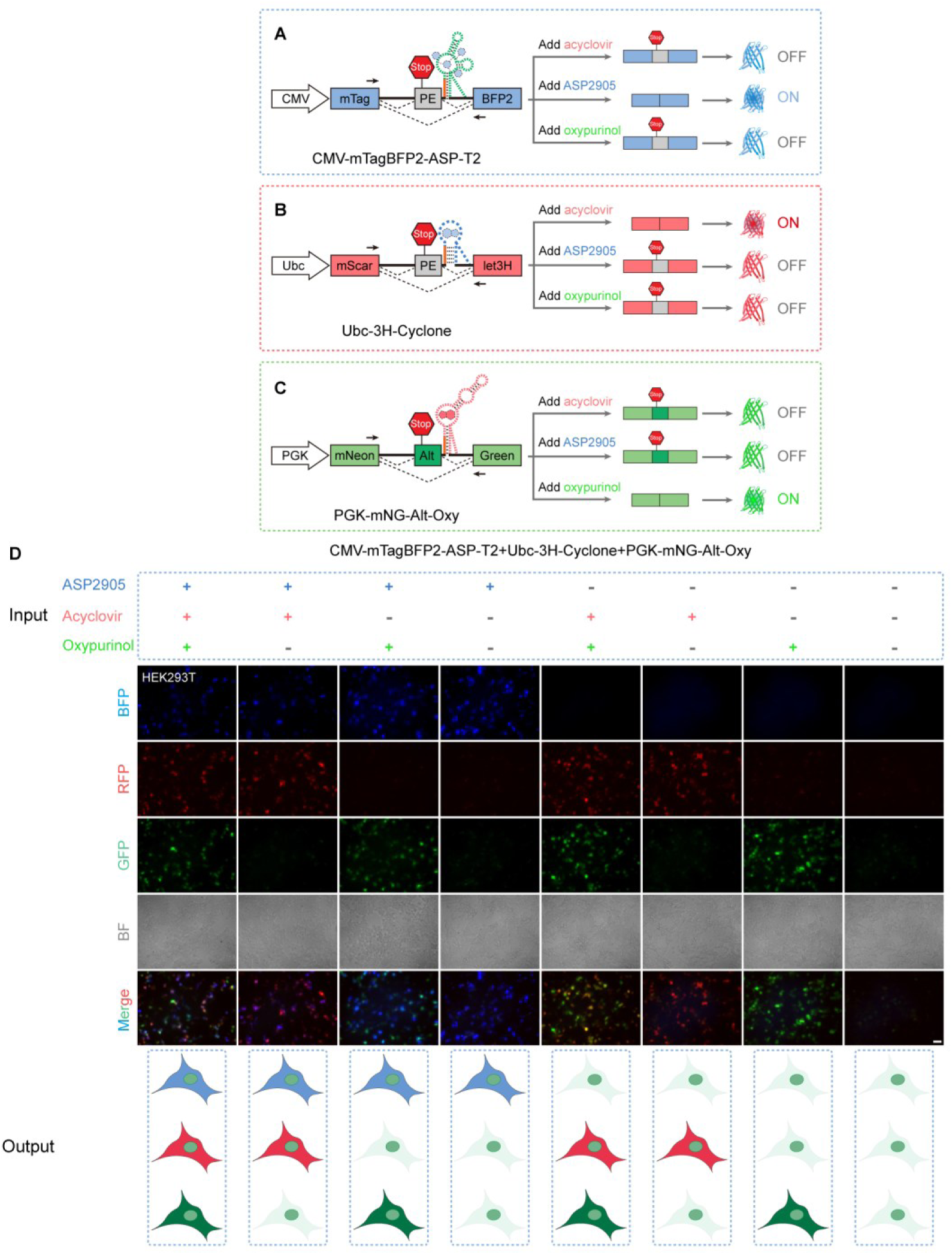
Orthogonal three-channel multiplexed gene control by small-molecule–gated splicing switches. (**A**–**C**) Schematics of three-channel orthogonal control. ASP2905, acyclovir, and oxypurinol selectively engage their respective cognate aptamers to independently drive PE skipping, thereby selectively activating mTagBFP2 (blue), mScarlet3H (red), and mNeonGreen (green) expression cassettes, respectively. (**D**) Three-color live-cell imaging demonstrating channel orthogonality. HEK293T cells co-expressing all three reporter constructs were treated with individual ligands (10 μM ASP2905, 125 μM acyclovir, or 200 μM oxypurinol) or their combinations for 24 h, yielding spectrally resolvable, mutually exclusive outputs with negligible inter-channel cross-talk. Scale bars, 50 μm. Representative images from at least three independent biological replicates.

**Supplementary Figure 20.**
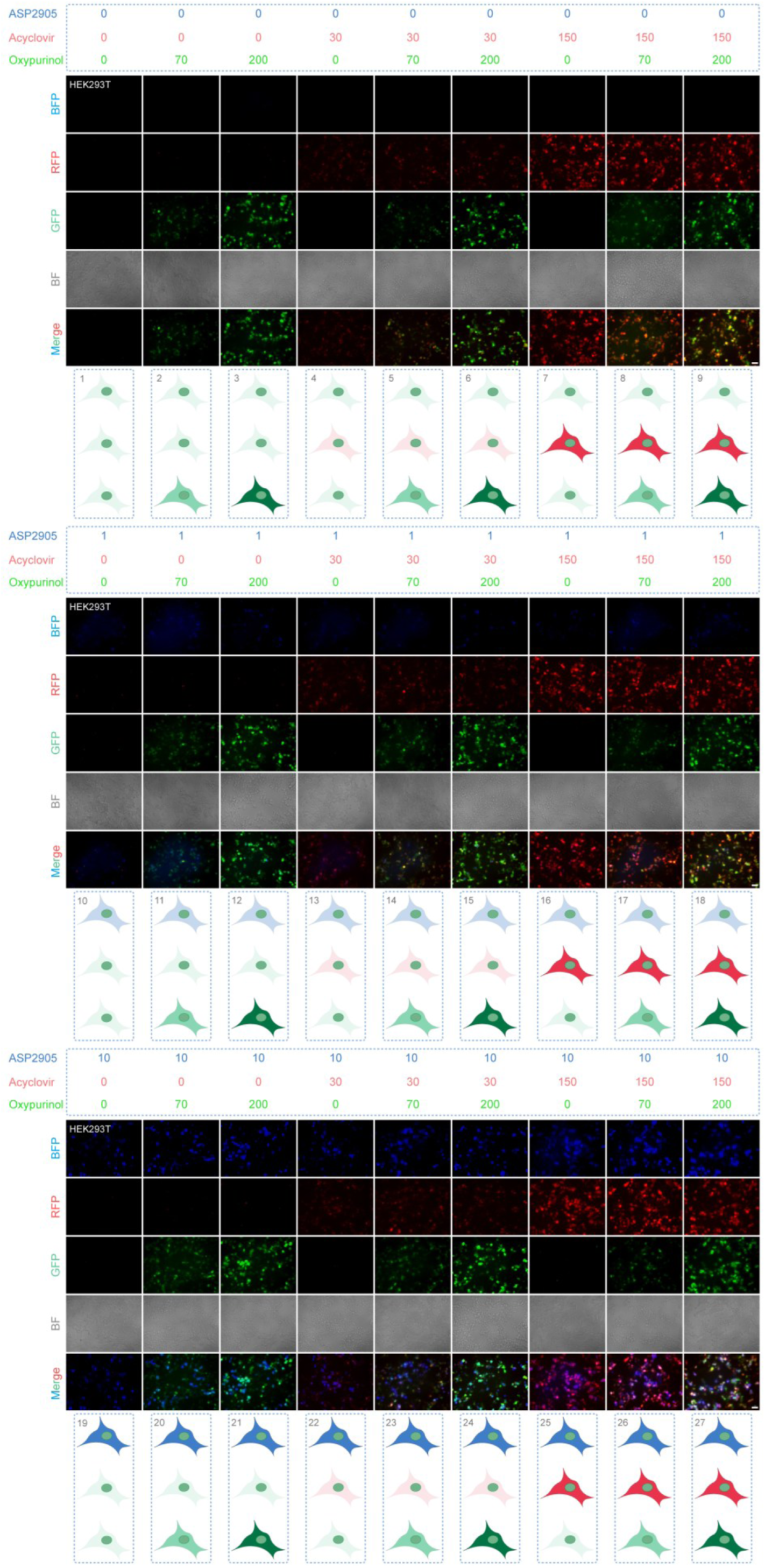
Combinatorial output matrix of three orthogonal small-molecule–gated splicing switches. Panoramic three-color fluorescence matrix illustrating all 27 (3^3^) combinatorial output states accessible through orthogonal ligand pairing, demonstrating multi-input logic gating with negligible inter-channel cross-talk. Scale bars, 50 μm. Representative images from at least three independent biological replicates.

**Supplementary Figure 21.**
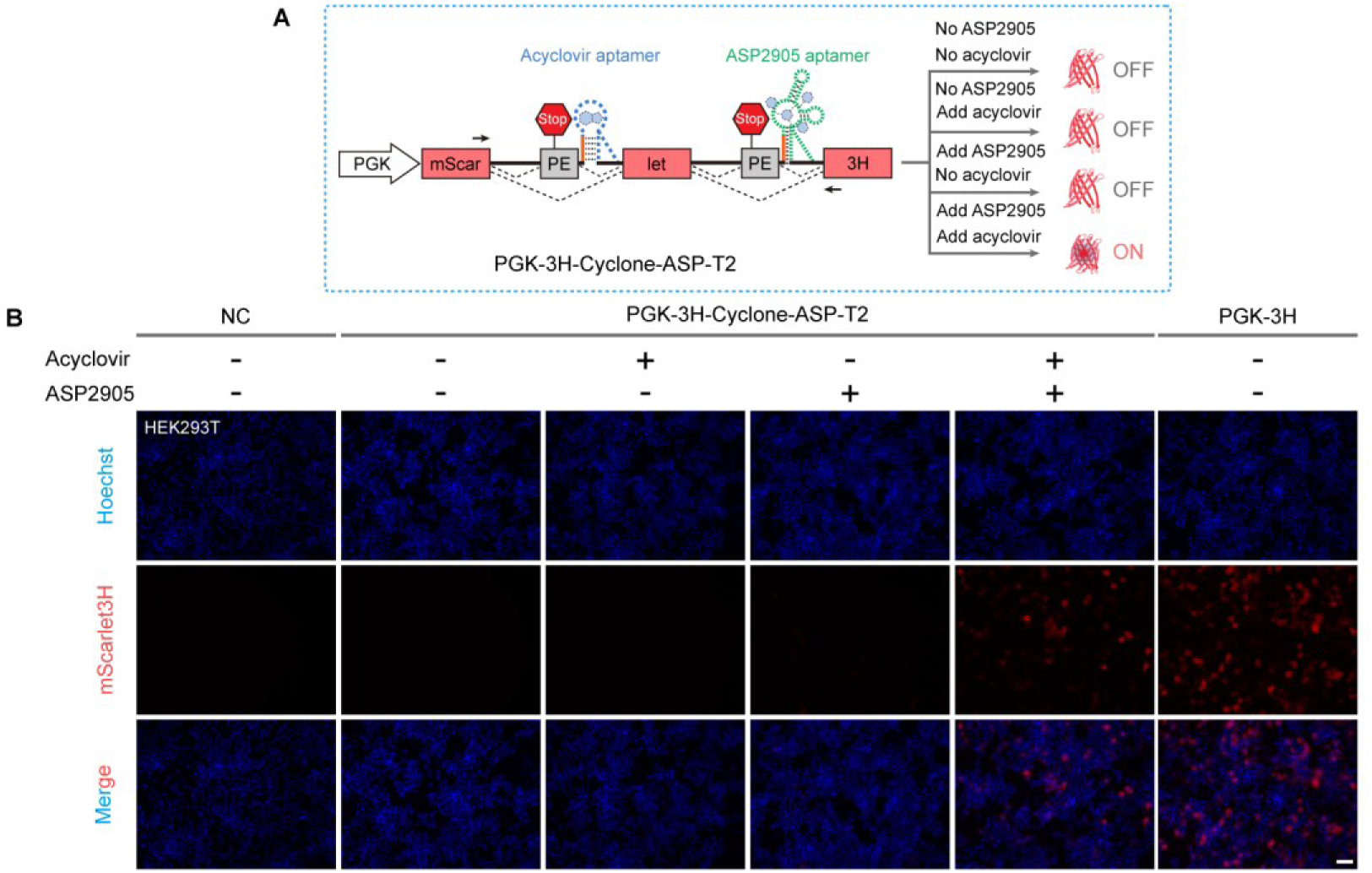
Dual-ligand AND-gate regulation by small-molecule–gated splicing switches. (**A**) Schematic of the dual-ligand AND-gate circuit. Cooperative binding of acyclovir and ASP2905 to their respective cognate aptamers drives PE skipping to activate target gene expression. (**B**) Live-cell fluorescence imaging validating AND-gate logic behavior. HEK293T cells transfected with the AND-gate reporter construct were treated with vehicle control (DMSO), 125 μM acyclovir, 10 μM ASP2905, or the two-ligand combination for 24 h. Robust mScarlet3H expression is exclusively observed upon simultaneous dual-ligand treatment. Scale bars, 50 μm. Representative images from at least three independent biological replicates.

**Supplementary Figure 22.**
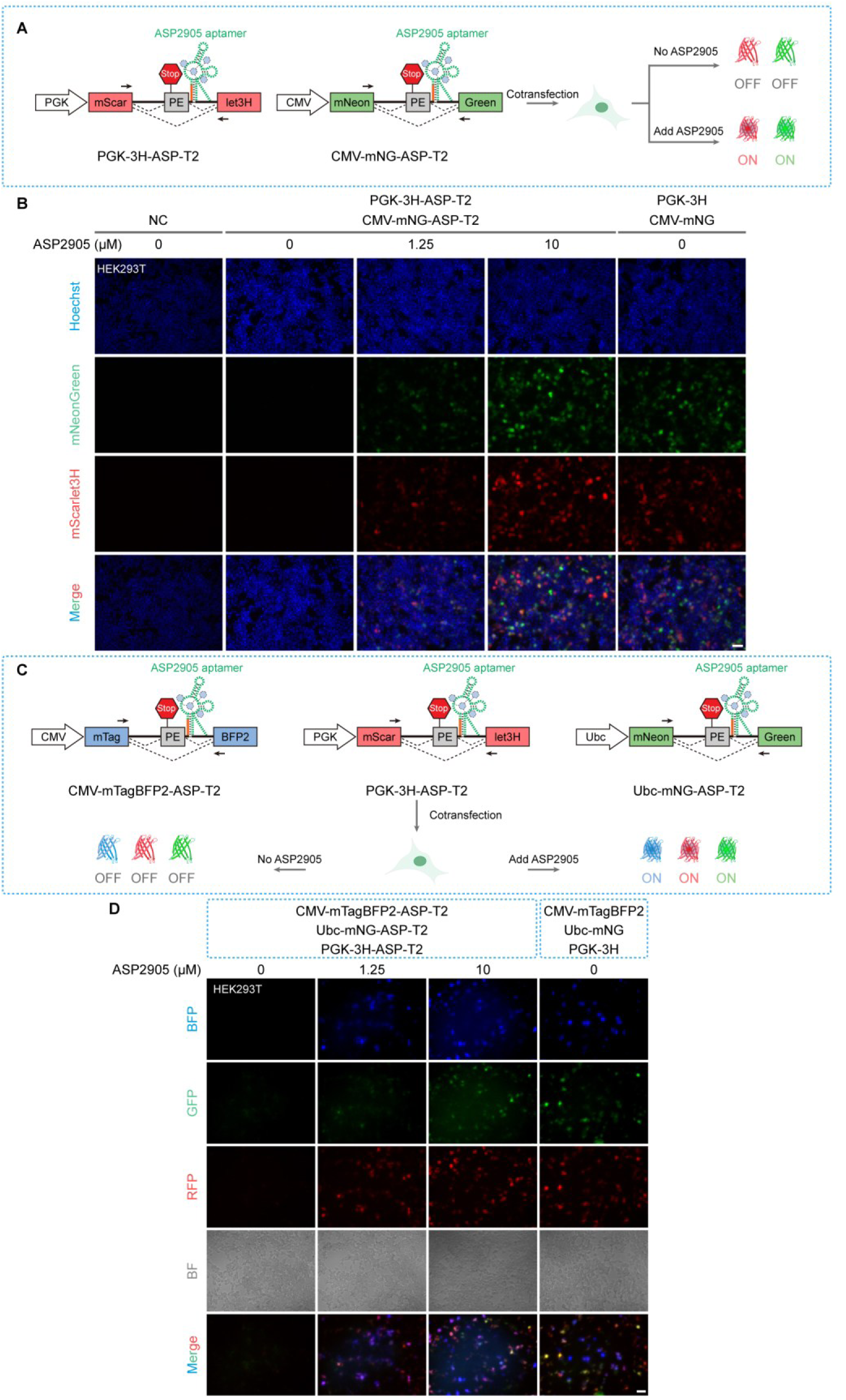
Single-ligand multi-gene activation. (**A**) Schematic of dual-gene activation. A single ASP2905 input triggers simultaneous PE skipping across two independent loci. (**B**) Dual-gene live-cell imaging. HEK293T cells co-transfected with two distinct ASP2905-responsive reporters were treated with 10 μM ASP2905 for 24 h, driving concurrent, robust mScarlet3H and mNeonGreen expression. (**C**) Schematic of tri-gene activation. A single ASP2905 input synchronously drives coordinated PE skipping across three independent loci. (**D**) Tri-gene live-cell imaging. HEK293T cells co-transfected with three distinct ASP2905-responsive reporters were treated with 10 μM ASP2905 for 24 h, yielding synchronized three-channel fluorescence output (mTagBFP2, mScarlet3H, and mNeonGreen). Scale bars, 50 μm. Representative images from at least three independent biological replicates.

**Supplementary Figure 23.**
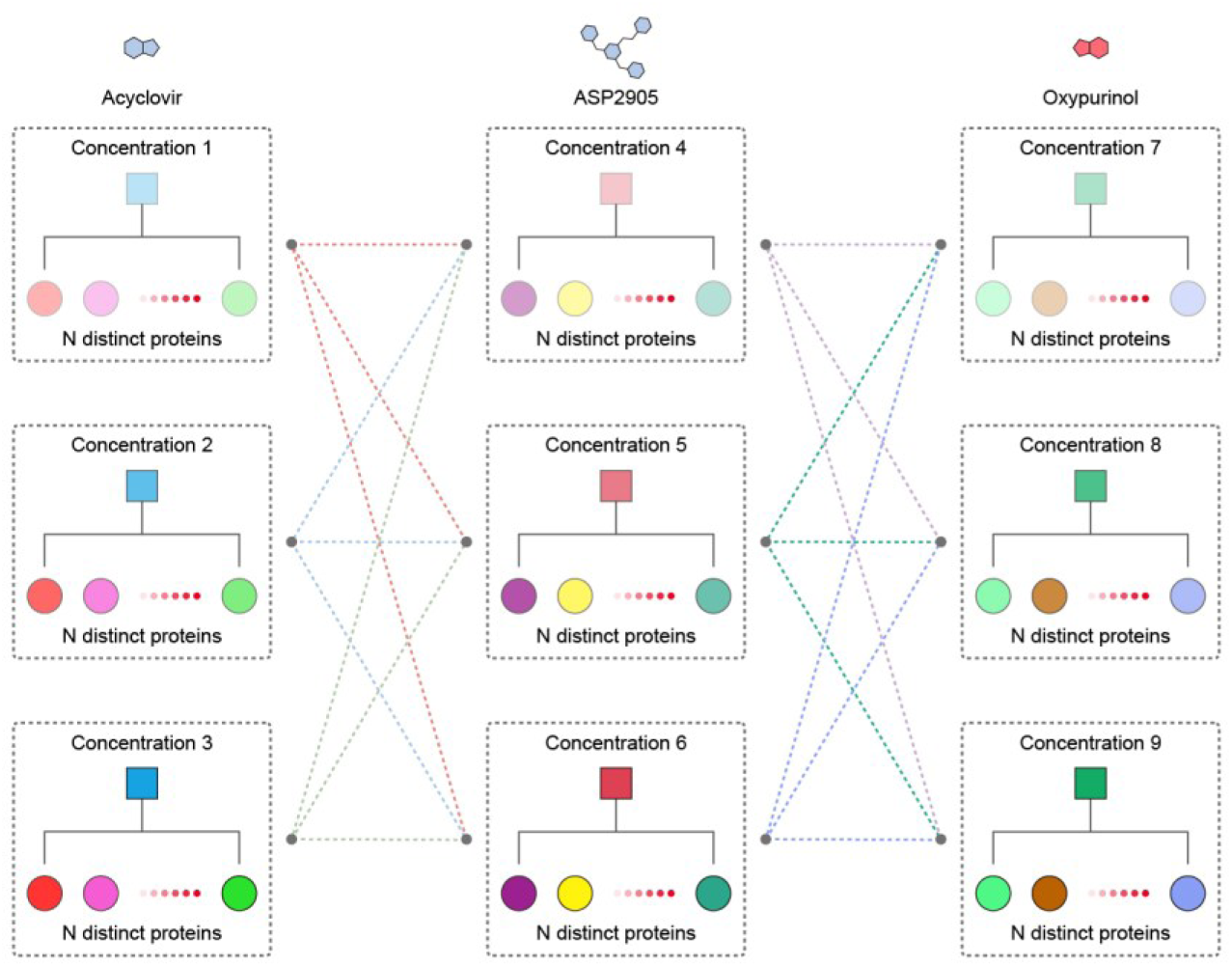
Conceptual framework for parallel and orthogonal multi-gene control by small-molecule–gated splicing switches. Schematic illustrating the theoretical design principle for parallel multi-protein regulation. Because each orthogonal ligand–aptamer module independently governs *N* distinct target proteins, three orthogonal splicing switches collectively enable simultaneous regulation of 3*N* proteins. Ligand titration within each channel thus provides an additional layer of continuous, finely tuned expression control while preserving strict orthogonality.

**Supplementary Figure 24.**
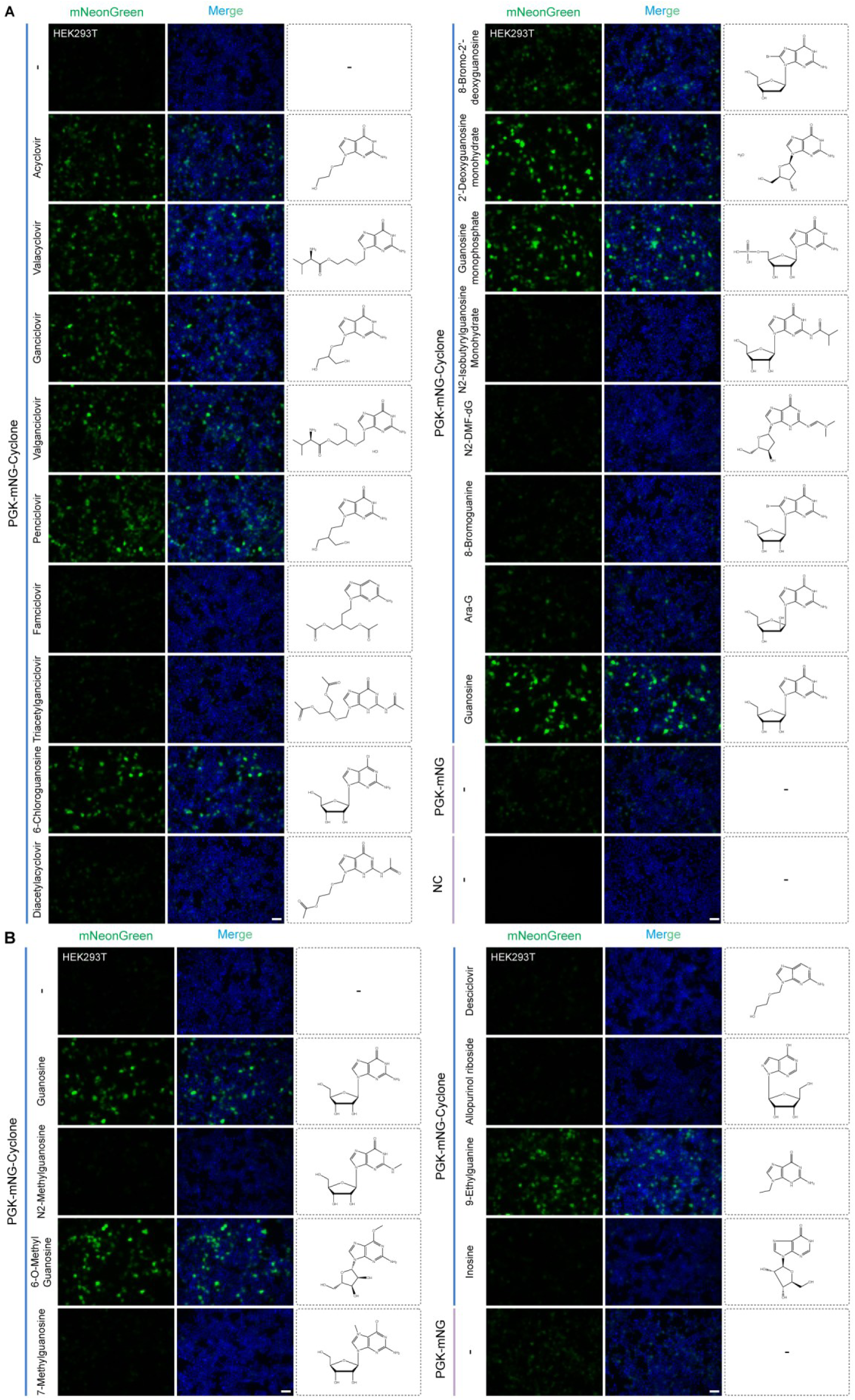
Structure–activity relationship and molecular determinants of acyclovir analogues. (**A**, **B**) Live-cell fluorescence imaging of the acyclovir-gated splicing reporter following treatment with 125 μM acyclovir analogues for 24 h. Splicing activation strictly requires an intact guanine pharmacophore: an unmodified *N*^2^-amino group is essential to retain critical hydrogen-bonding interactions within the aptamer binding pocket, whereas the *N*^9^ position accommodates acyclic side chains or small alkyl substitutions. Conversely, bulky *N*^2^-substitutions (e.g., isobutyryl or dimethylaminomethylene), altered sugar puckering geometries (e.g., arabinofuranosyl), or heterocycle modifications (e.g., inosine or allopurinol) completely abrogate PE skipping. Scale bars, 50 μm. Representative images from at least three independent biological replicates.

**Supplementary Figure 25.**
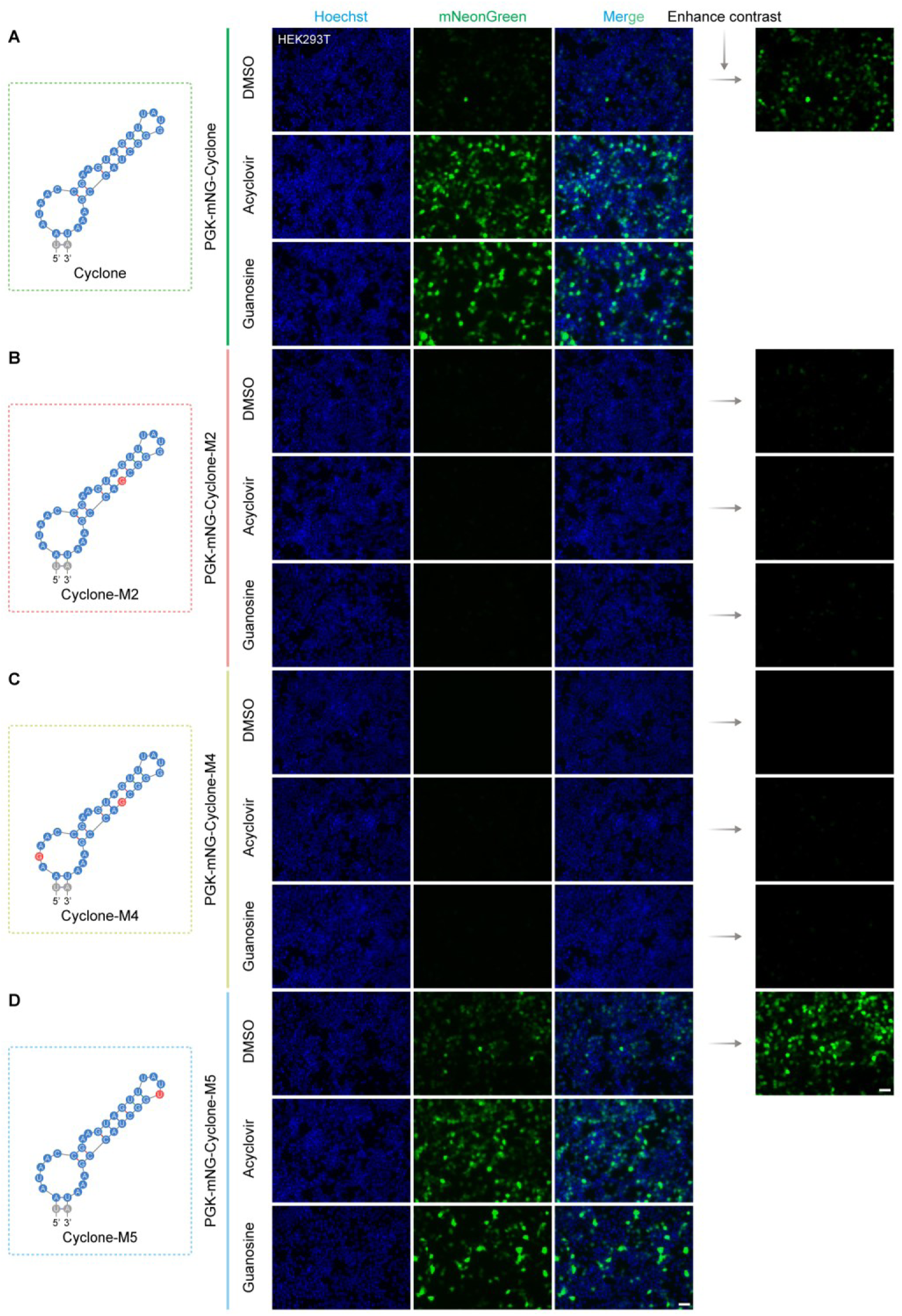
Reverse-genetic evidence that endogenous nucleosides drive Cyclone basal leakage. (**A**–**D**) Live-cell fluorescence imaging of wild-type and variant Cyclone reporters treated with vehicle control (DMSO), 125 μM acyclovir, or 125 μM guanosine for 24 h. The wild-type Cyclone (**A**) and the ligand-responsive M5 variant (**D**) respond to both ligands and exhibit pronounced basal fluorescence. By contrast, the binding-deficient M2 (**B**) and M4 (**C**) variants, harboring targeted mutations at the ligand-binding interface, are completely unresponsive to either ligand, with fluorescence remaining negligible under all conditions. Red circles denote mutation sites. Scale bars, 50 μm. Representative images from at least three independent biological replicates.

**Supplementary Figure 26.**
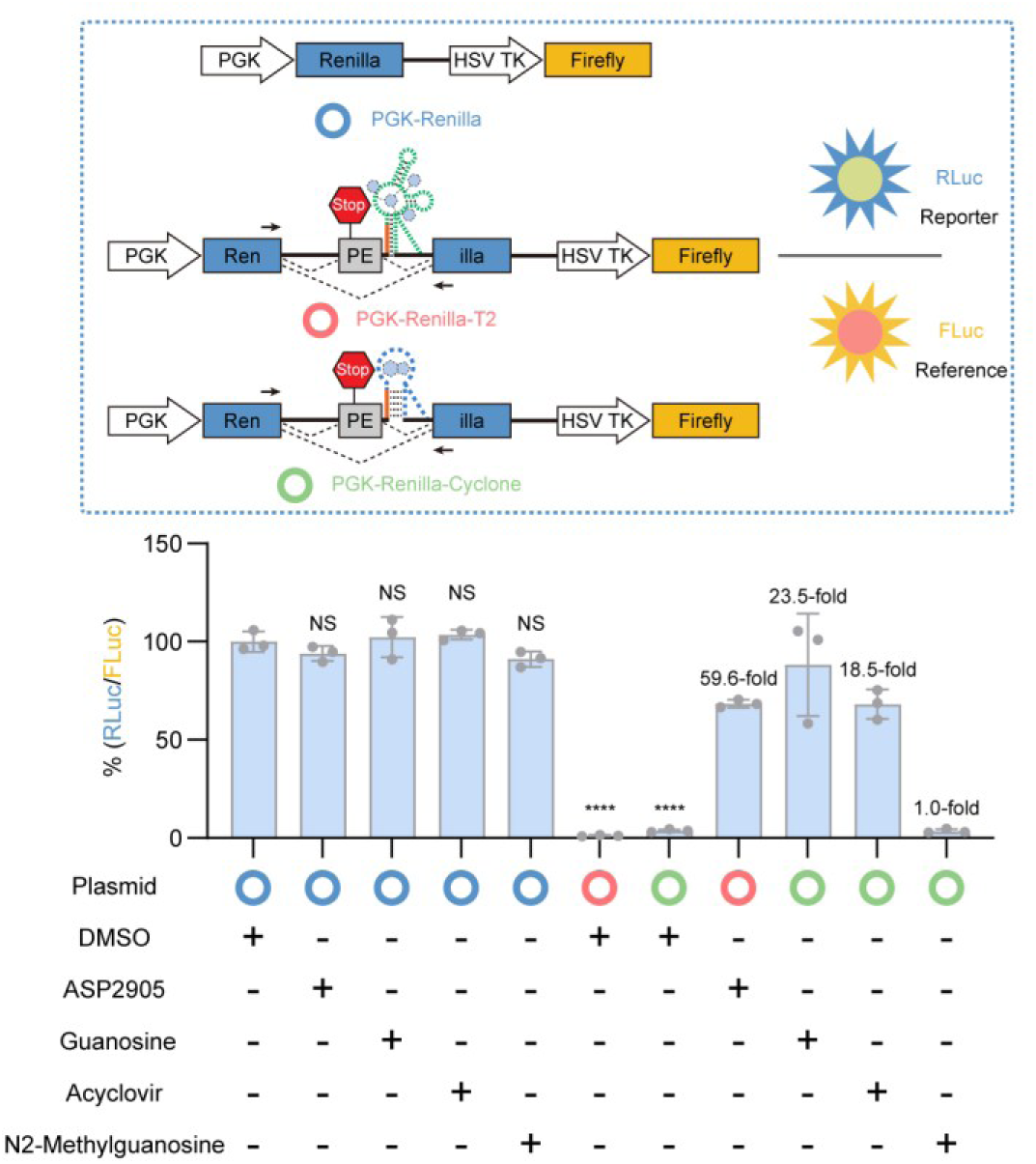
Dynamic range analysis of ligand-gated splicing switches by dual-luciferase assay. Acyclovir (125 μM) and ASP2905 (10 μM) elicit comparable maximal luciferase output at 24 h; the ASP2905-gated circuit displays markedly suppressed basal activity, translating to a ∼3.2-fold improvement in both signal-to-noise ratio and dynamic range relative to the acyclovir-gated system. *N*^2^-methylguanosine (125 μM) fails to trigger luciferase expression above vehicle control baseline. Data represent mean ± s.d. from ≥3 independent experiments. \*\*\*\**P* < 0.0001; NS, not significant.

**Supplementary Figure 27.**
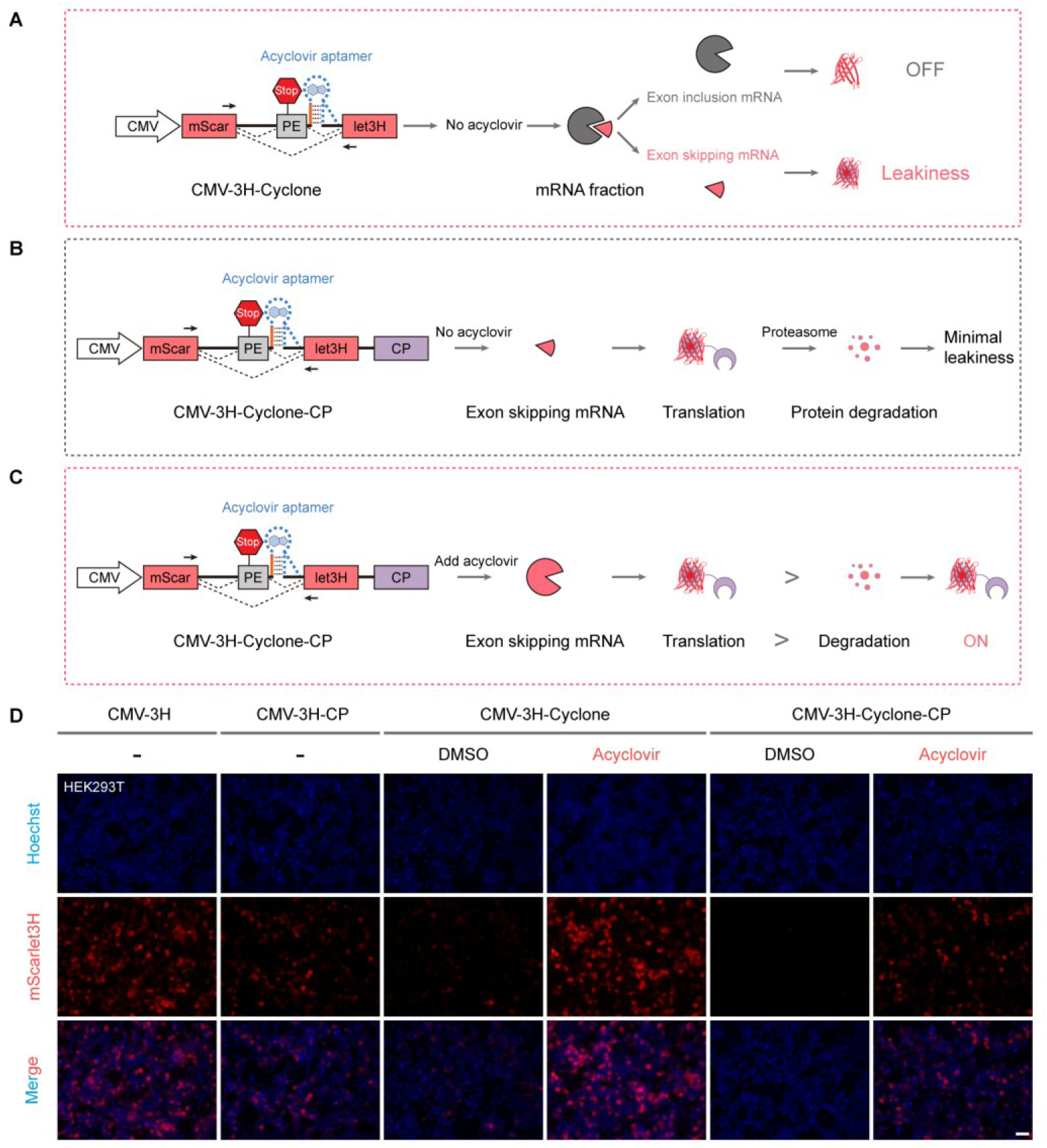
Mitigation of Cyclone basal leakage via the Ubiquitin– Proteasome System (UPS). (**A**–**C**) Schematic of UPS-mediated basal leakage suppression. In the absence of ligand, the trace reporter generated by constitutive low-level PE skipping is efficiently cleared by the C-terminal CP degron through proteasomal degradation. Upon ligand binding, robust PE skipping exceeds the threshold of UPS degradative capacity, permitting net reporter accumulation and measurable fluorescence. (**D**) Live-cell fluorescence imaging validating effective background suppression by the CP degron. Scale bars, 50 μm. Representative images from at least three independent biological replicates.

**Supplementary Figure 28.**
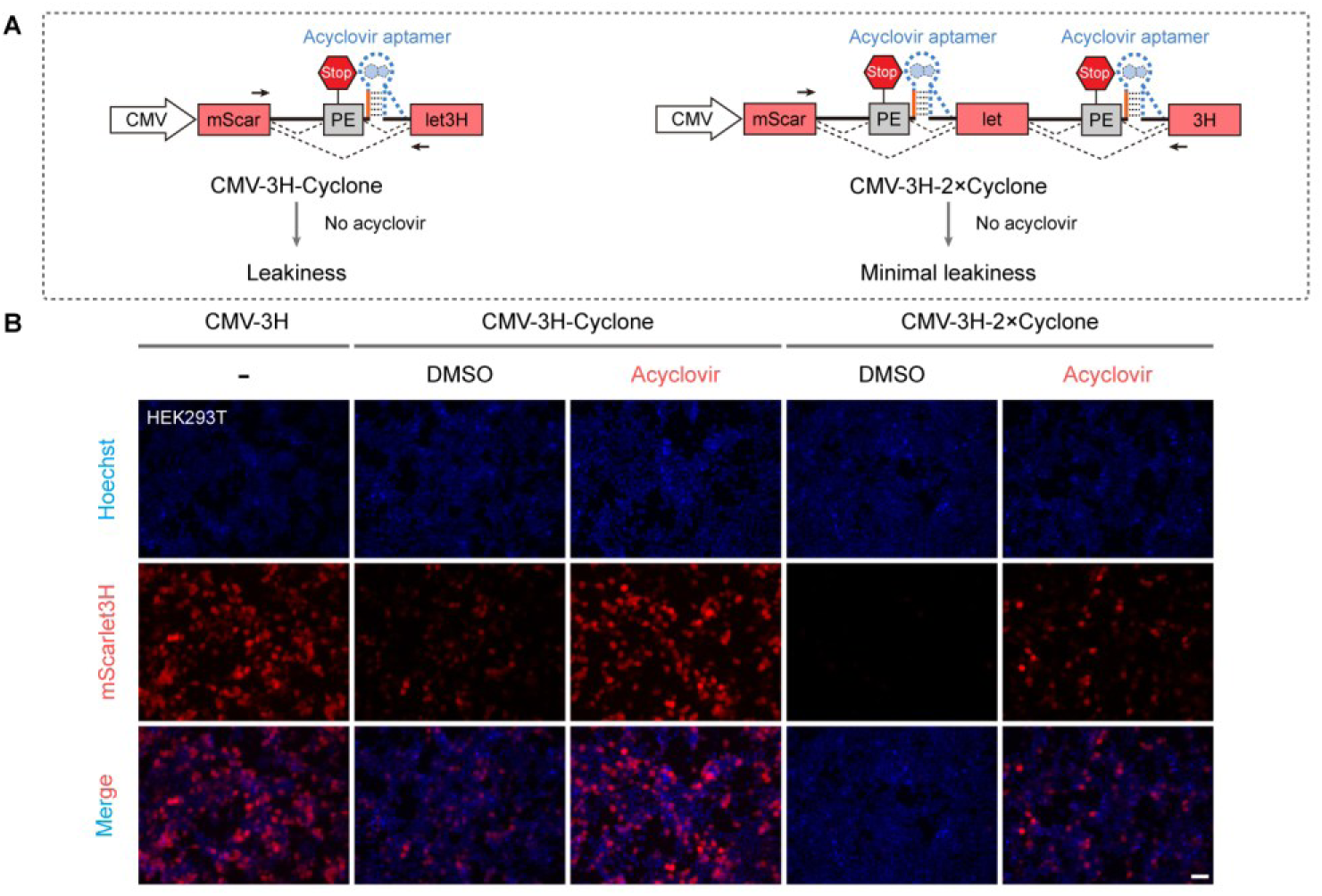
Tandem riboswitches suppress Cyclone basal leakage at the post-transcriptional level. (**A**) Schematic of basal leakage suppression via tandem riboswitch integration. In the ligand-free state, spontaneous PE skipping occurs with low stochastic probability at each autonomous splicing junction. Because productive leakage necessitates concurrent skipping across all tandem junctions, this configuration reduces background expression to near-zero levels. (**B**) Live-cell fluorescence imaging of reporter constructs bearing tandem riboswitches. Scale bars, 50 μm. Representative images from at least three independent biological replicates.

**Supplementary Figure 29.**
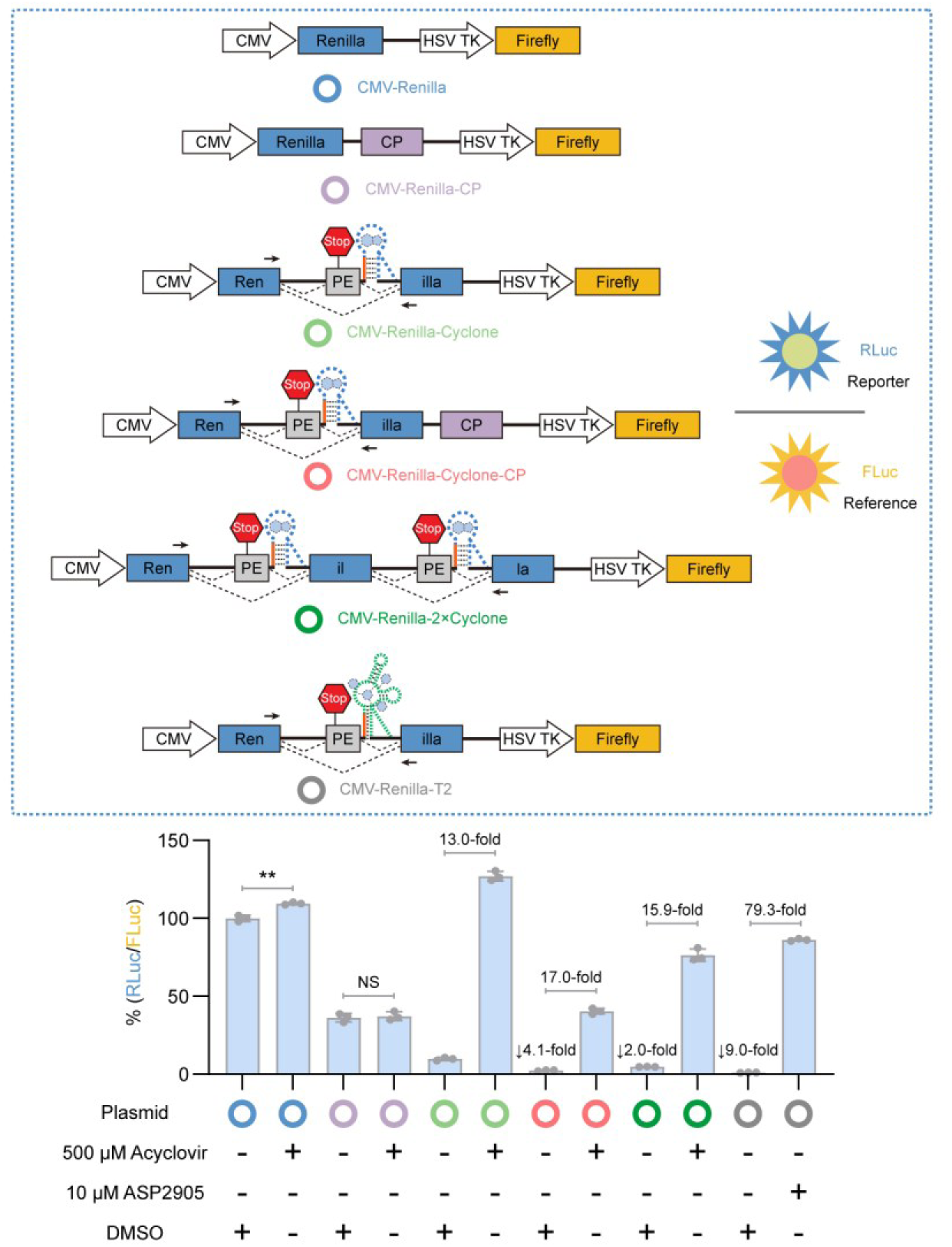
Quantitative comparison of general noise-reduction strategies by dual-luciferase assay. Relative to the wild-type Cyclone system, both engineering strategies markedly suppress basal leakage. In particular, the CP-tag fusion (Cyclone-CP) and the tandem-riboswitch construct (2×Cyclone) attenuate basal expression by 4.1-fold and 2.0-fold, respectively. Data represent mean ± s.d. from ≥3 independent biological replicates. \*\**P* < 0.01; NS, not significant.

**Supplementary Figure 30.**
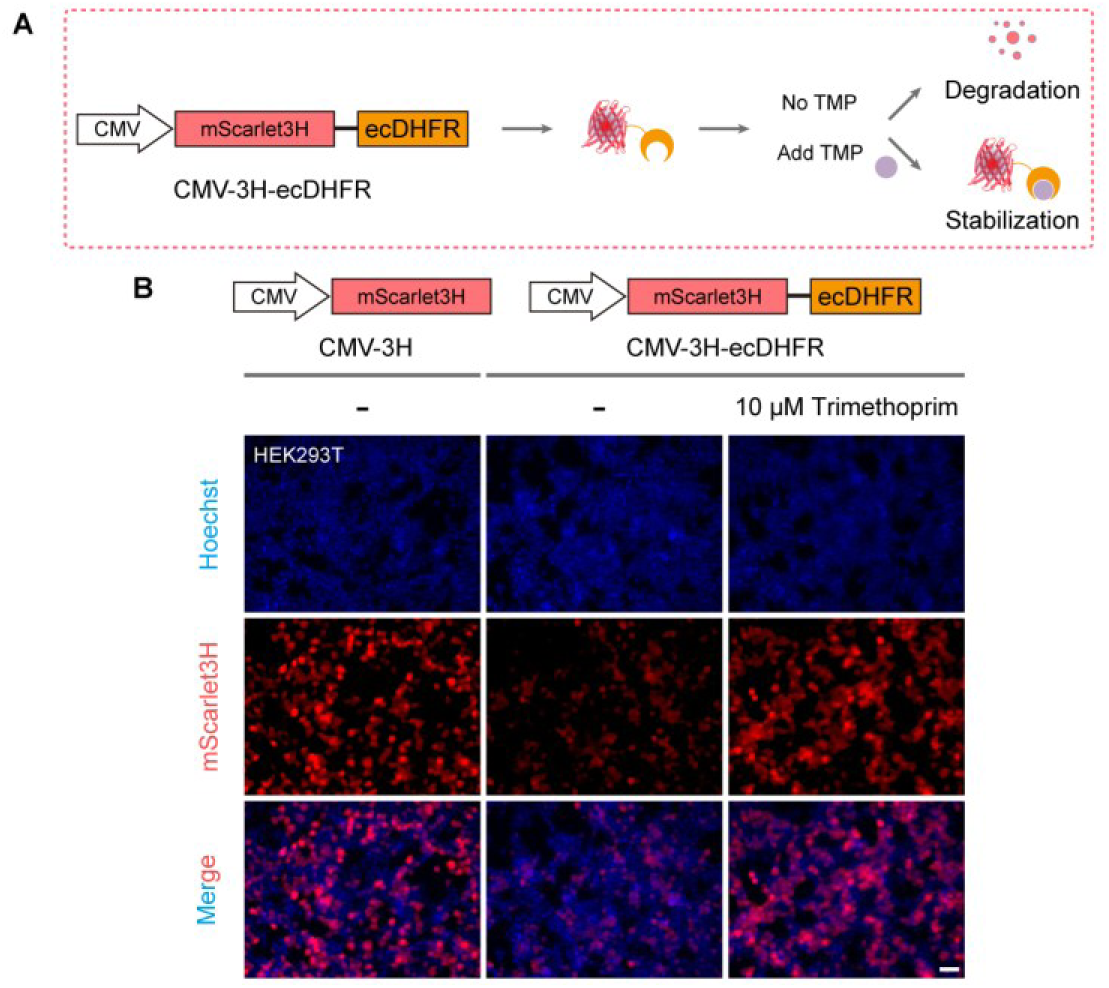
Trimethoprim (TMP) prevents destabilization of ecDHFR-fused proteins. (**A**) TMP binds to and stabilizes ecDHFR-fused proteins. (**B**) Live-cell fluorescence imaging. Upon transfection of the corresponding plasmids and TMP treatment, reporter proteins fused to ecDHFR are stabilized at the post-translational level. Scale bars, 50 μm. Representative images from at least three independent biological replicates.

**Supplementary Figure 31.**
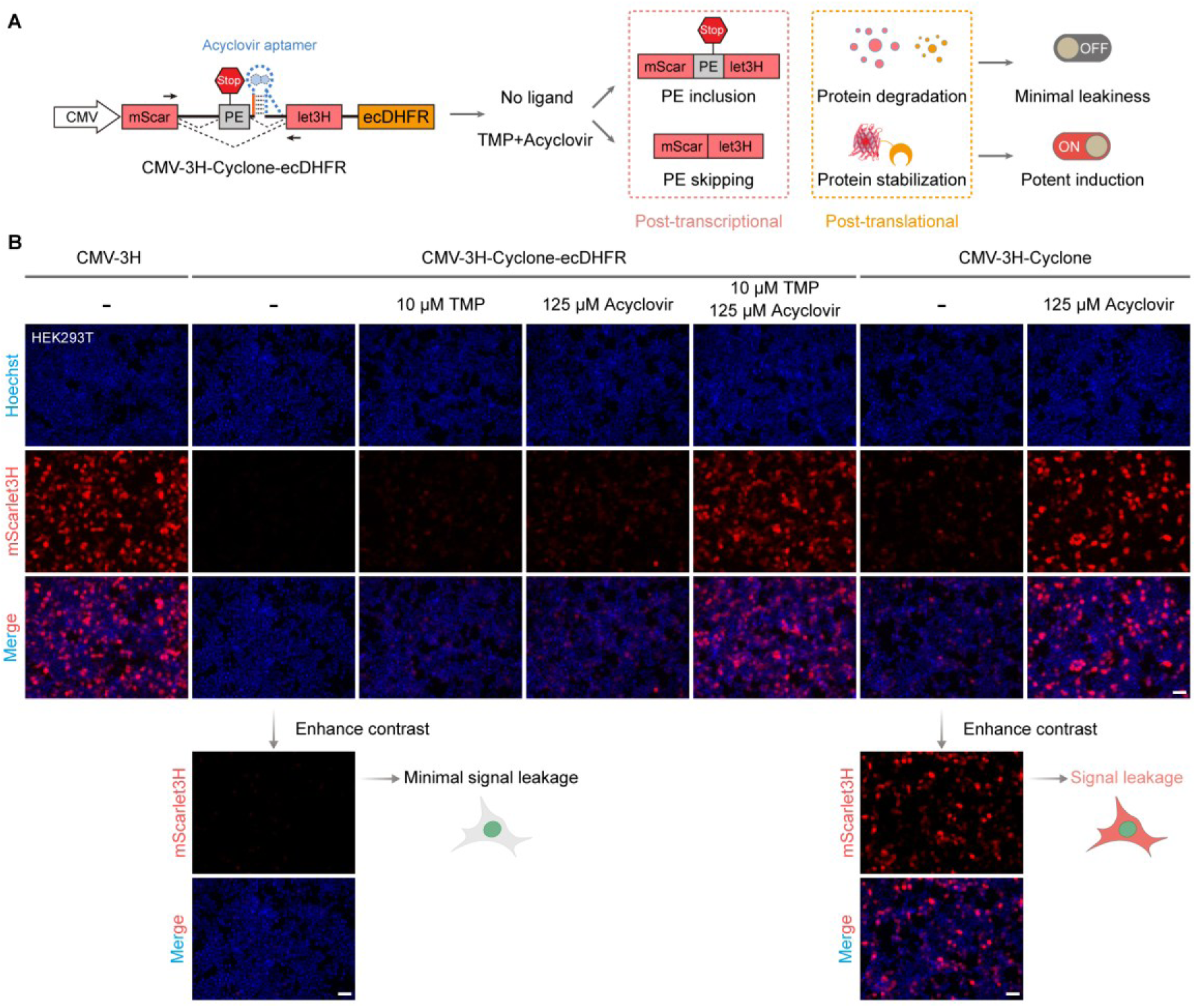
Noise reduction via C-terminal ecDHFR fusion. (**A**) Dual noise-reduction mechanism operating at both post-transcriptional and post-translational levels without compromising maximal induced expression. (**B**) Live-cell fluorescence imaging confirming background suppression. In the ligand-free state, the ecDHFR-fused reporter exhibits near-zero basal leakage; co-treatment with both cognate ligands (TMP and acyclovir) triggers robust expression with negligible attenuation of maximal output. Scale bars, 50 μm. Representative images from at least three independent biological replicates.

**Supplementary Figure 32.**
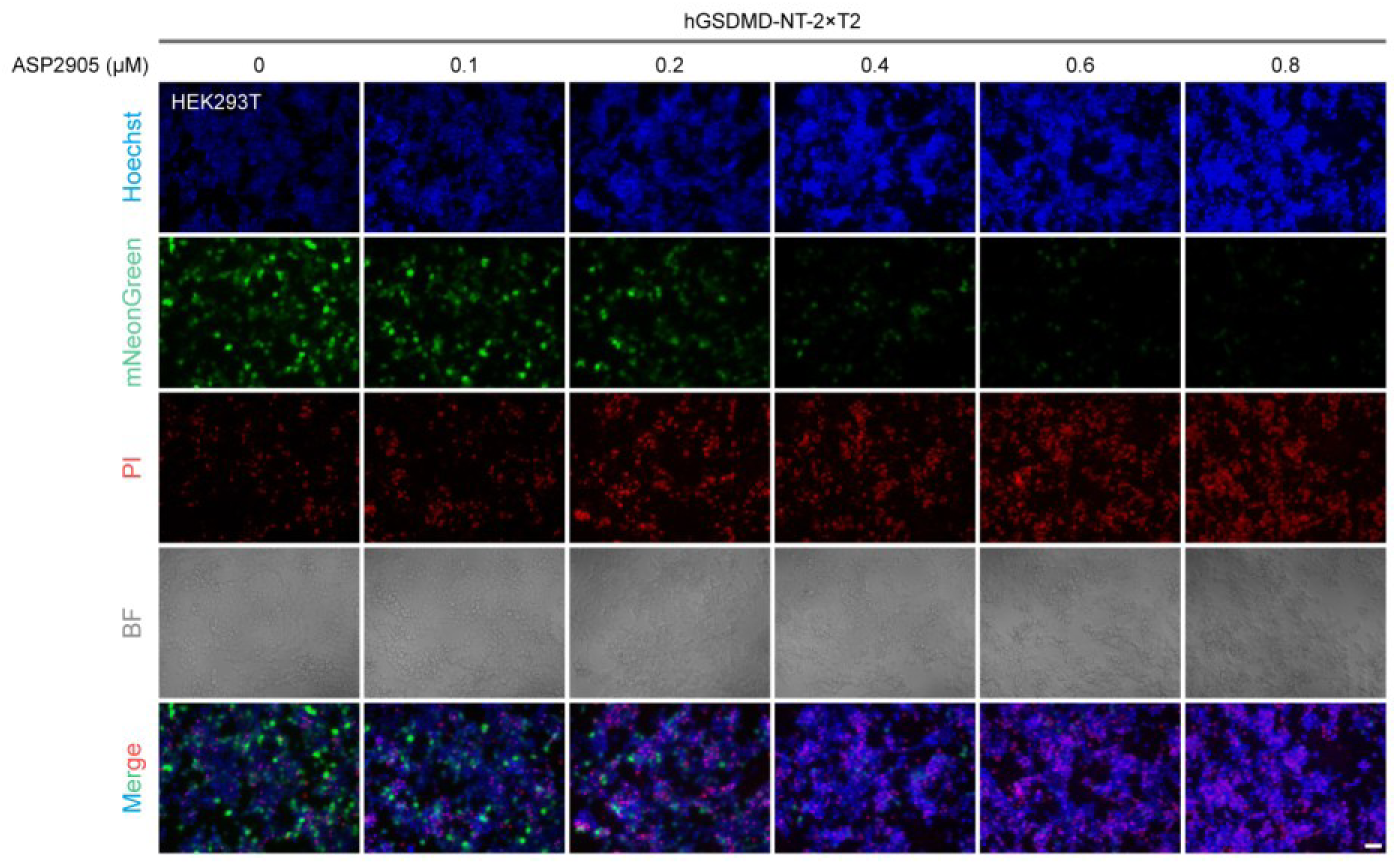
Dose-dependent pyroptosis induction by ASP2905 in cells expressing the ASP2905-gated hGSDMD-NT construct. Live-cell fluorescence imaging with propidium iodide (PI) staining. Cells expressing the ASP2905-gated hGSDMD-NT construct were treated with the indicated concentrations of ASP2905 for 24 h. Increasing ligand concentrations drive a dose-dependent increase in PI-positive cells, confirming ligand-gated pyroptotic cell death. Scale bars, 50 μm. Representative images from at least three independent biological replicates.

**Supplementary Figure 33.**
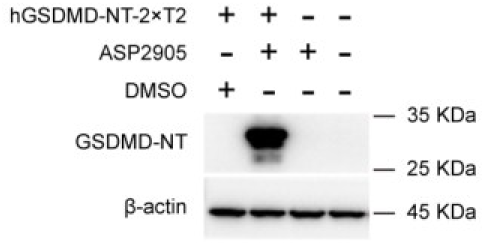
ASP2905-dependent induction of GSDMD-NT protein. Immunoblot analysis of lysates from HEK293T cells expressing the ASP2905-gated hGSDMD-NT construct or mock control cells, following treatment with ASP2905 (10 μM) or vehicle control (DMSO) for 24 h. ASP2905 treatment elicits a specific GSDMD-NT band exclusively in cells harboring the gated construct, whereas no signal is detectable in the absence of ligand or in control cells lacking the construct. GAPDH serves as a loading control. Representative images from at least three independent biological replicates.

**Supplementary Figure 34.**
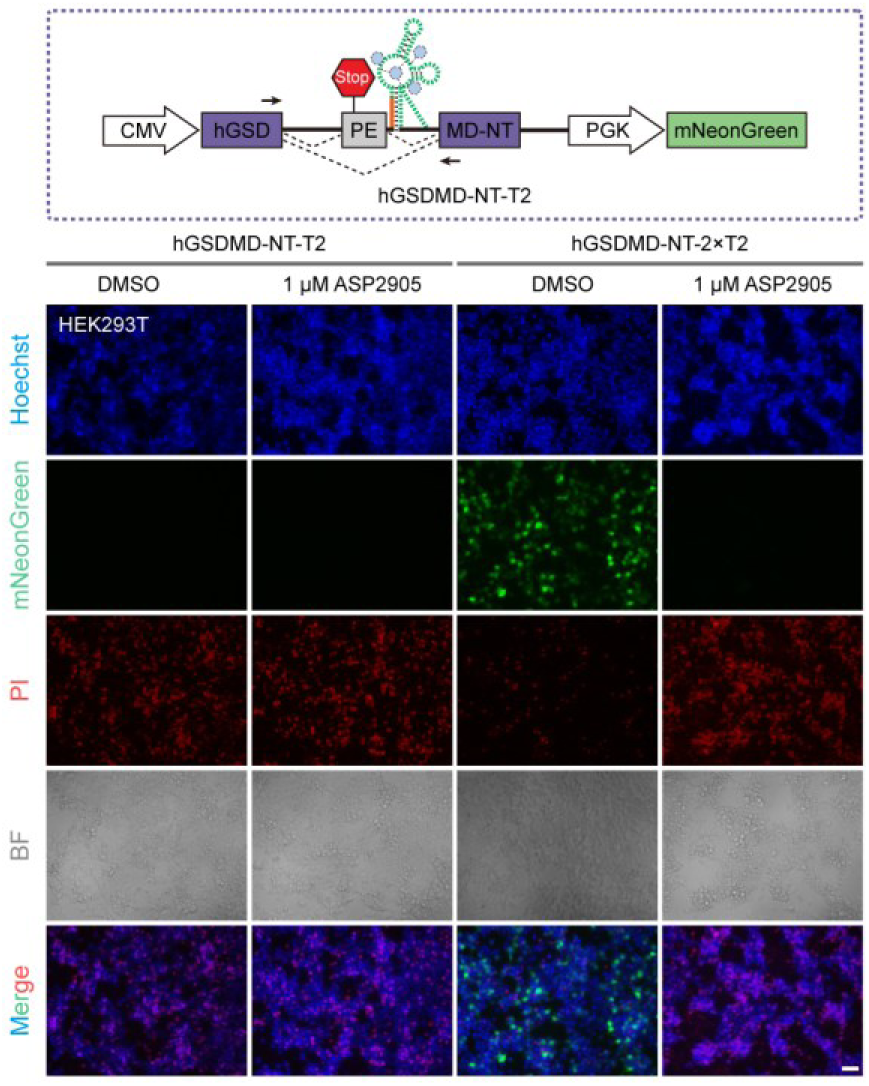
Single versus tandem ASP2905-gated splicing switches in GSDMD-NT leakage. Live-cell fluorescence imaging evaluating basal leakage suppression. A single ASP2905-gated splicing switch permits constitutive GSDMD-NT leakage, causing cell death and virtual loss of the internal green fluorescent protein (GFP) control regardless of ligand status. In contrast, tandem ASP2905-gated switches effectively suppress basal leakage, maintaining robust GFP fluorescence and cell survival in the ligand-free state. Scale bars, 50 μm. Representative images from at least three independent biological replicates.

**Supplementary Figure 35.**
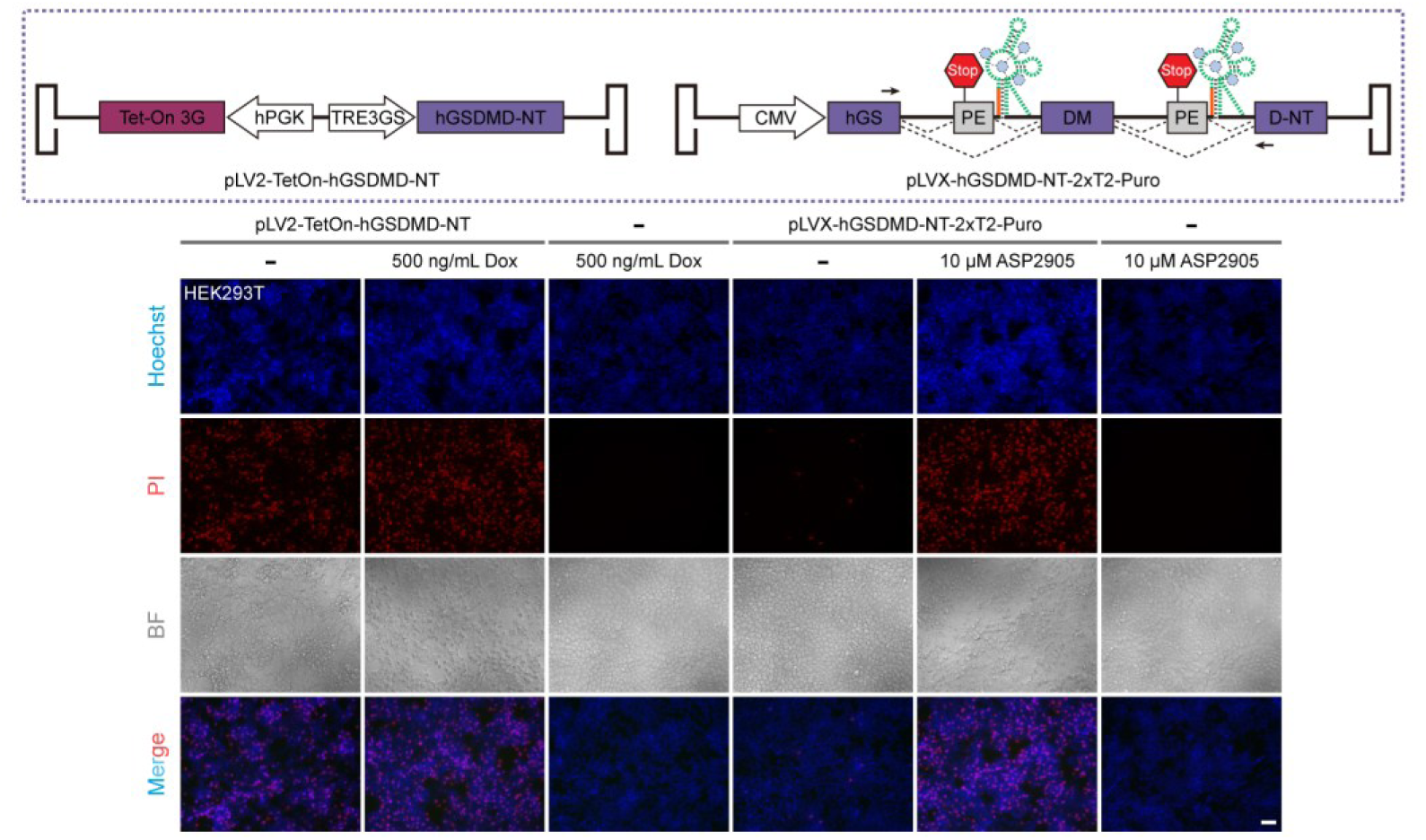
Tet-On 3G versus 2×T2 tandem switches in GSDMD-NT-mediated pyroptosis control. Live-cell fluorescence imaging with PI staining evaluating basal leakage and activation profiles. In the ligand-free state, Tet-On 3G exhibits substantial basal leakage, whereas 2×T2 maintains minimal background. Upon ligand addition, both systems trigger comparable PI uptake, confirming effective pyroptotic induction. Scale bars, 50 μm. Representative images from at least three independent biological replicates.

**Supplementary Figure 36.**
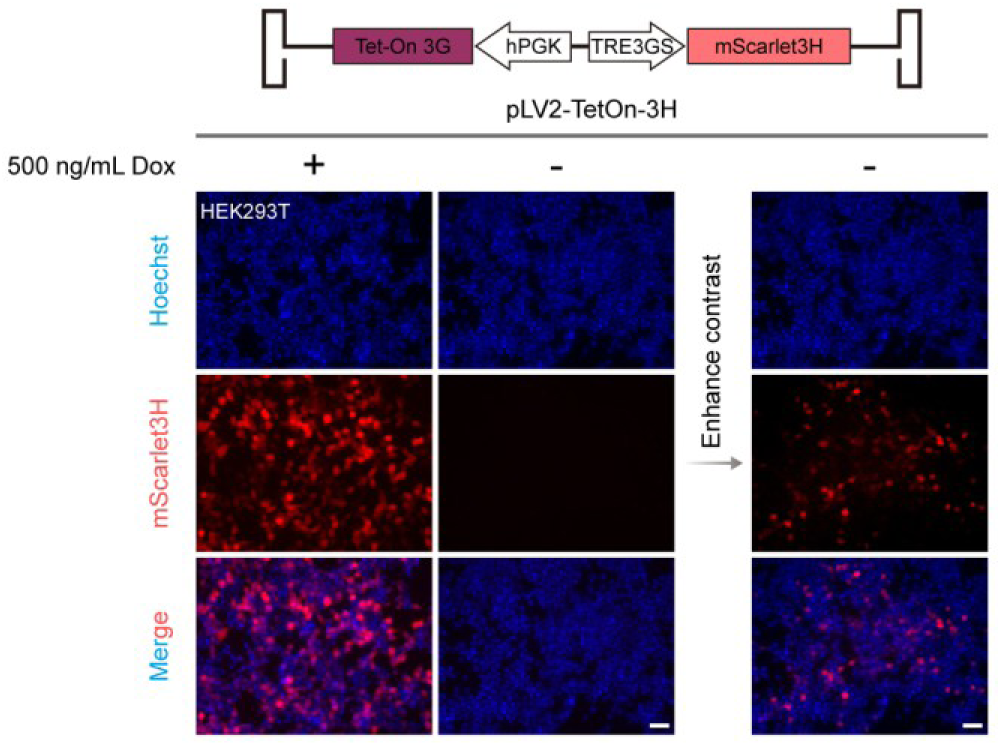
Basal leakage of the Tet-On 3G system. Live-cell fluorescence imaging. Despite its widespread use as an inducible expression system, Tet-On 3G remains prone to appreciable basal leakage in the ligand-free state. Scale bars, 50 μm. Representative images from at least three independent biological replicates.

**Supplementary Figure 37.**
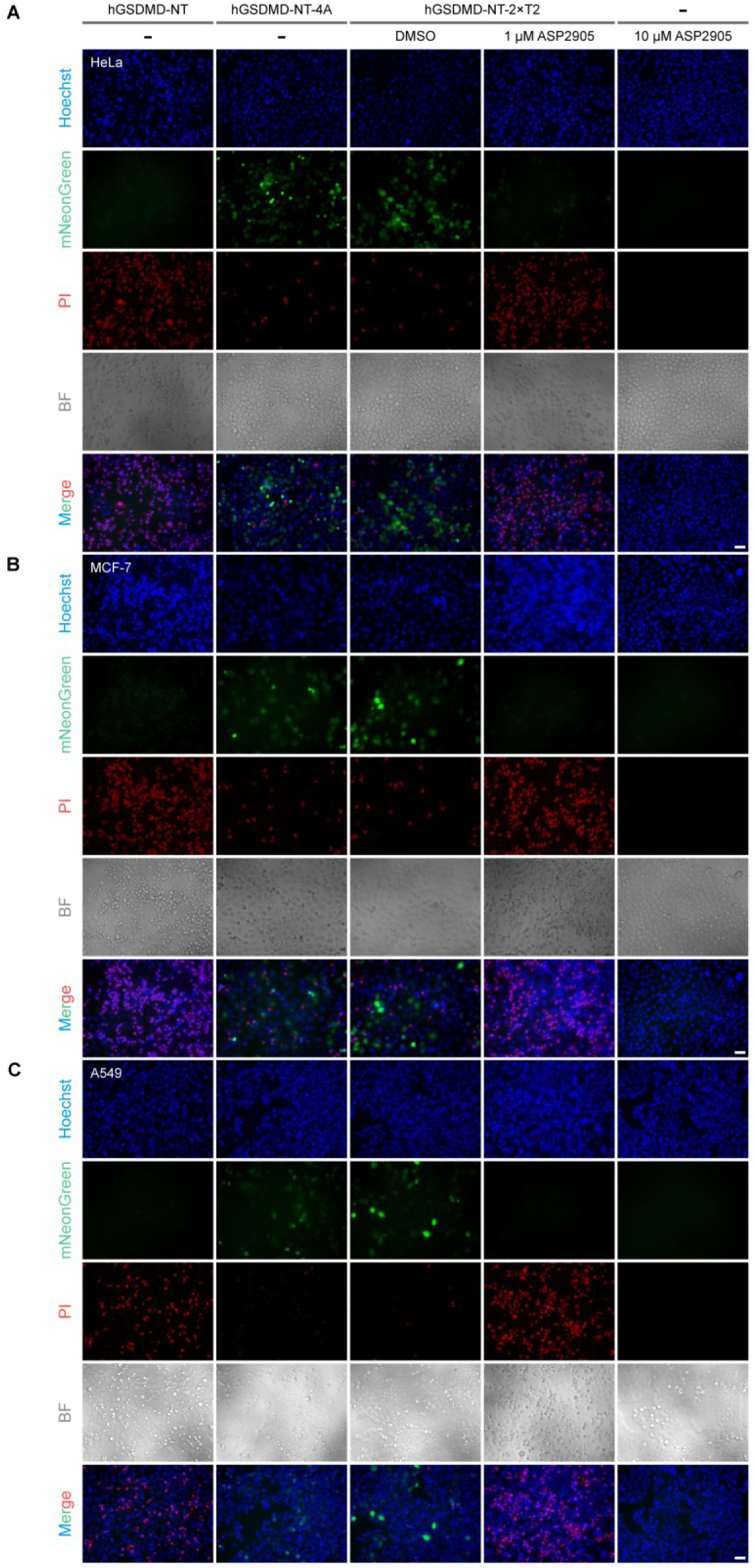
ASP2905-gated pyroptosis extends to diverse cancer cell lines. (**A**) HeLa, (**B**) MCF-7, and (**C**) A549 cells transfected with hGSDMD-NT-2×T2 and treated with ASP2905 (1 μM) or DMSO for 24 h, followed by PI staining. ASP2905 induces robust PI uptake across all lines, confirming pyroptotic cell death, whereas vehicle controls show minimal background. Scale bars, 50 μm. Representative images from at least three independent biological replicates.

**Supplementary Figure 38.**
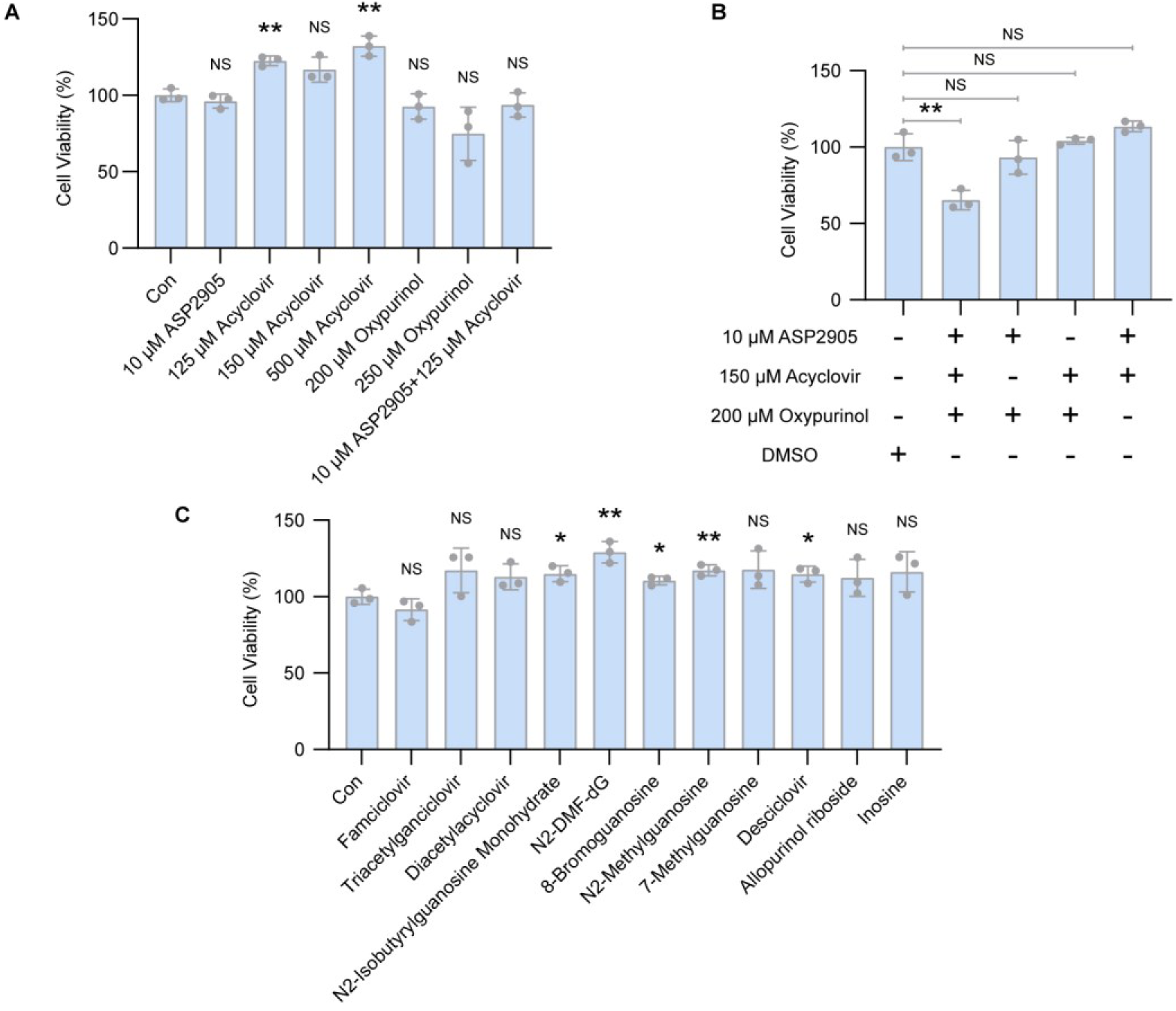
Ligand-dependent cytotoxicity in HEK293T cells. (**A**, **B**) Cell viability of HEK293T cells following treatment with ASP2905, acyclovir, or oxypurinol, alone or in combination. (**C**) Cell viability following treatment with inactive acyclovir analogues (125 μM) that fail to activate Cyclone. Data represent mean ± s.d. from at least three independent biological replicates. \**P* < 0.05, \*\**P* < 0.01; NS, not significant.

## Supplementary information

**Supplementary Figure 39.**
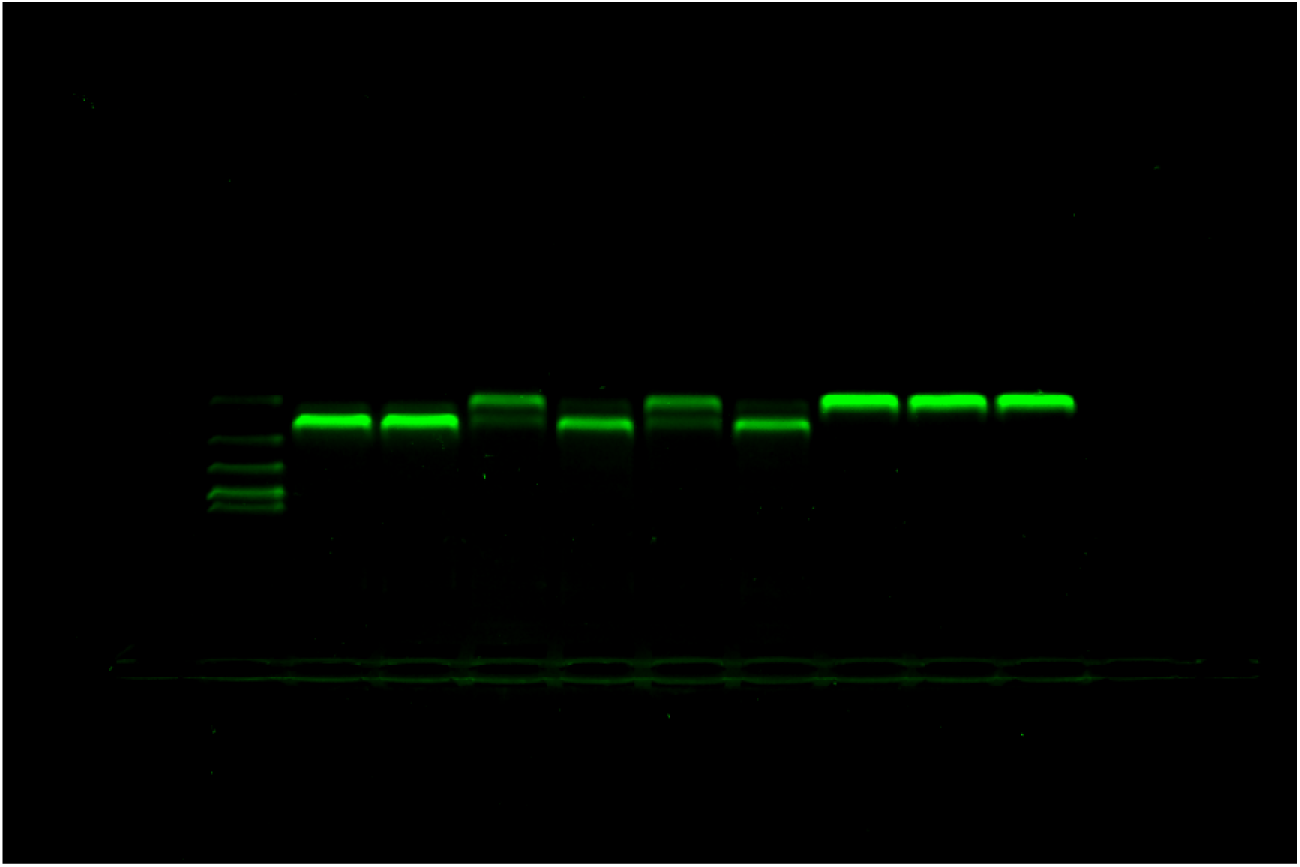
Uncropped agarose gel from Supplementary Fig. S10B.

**Supplementary Figure 40.**
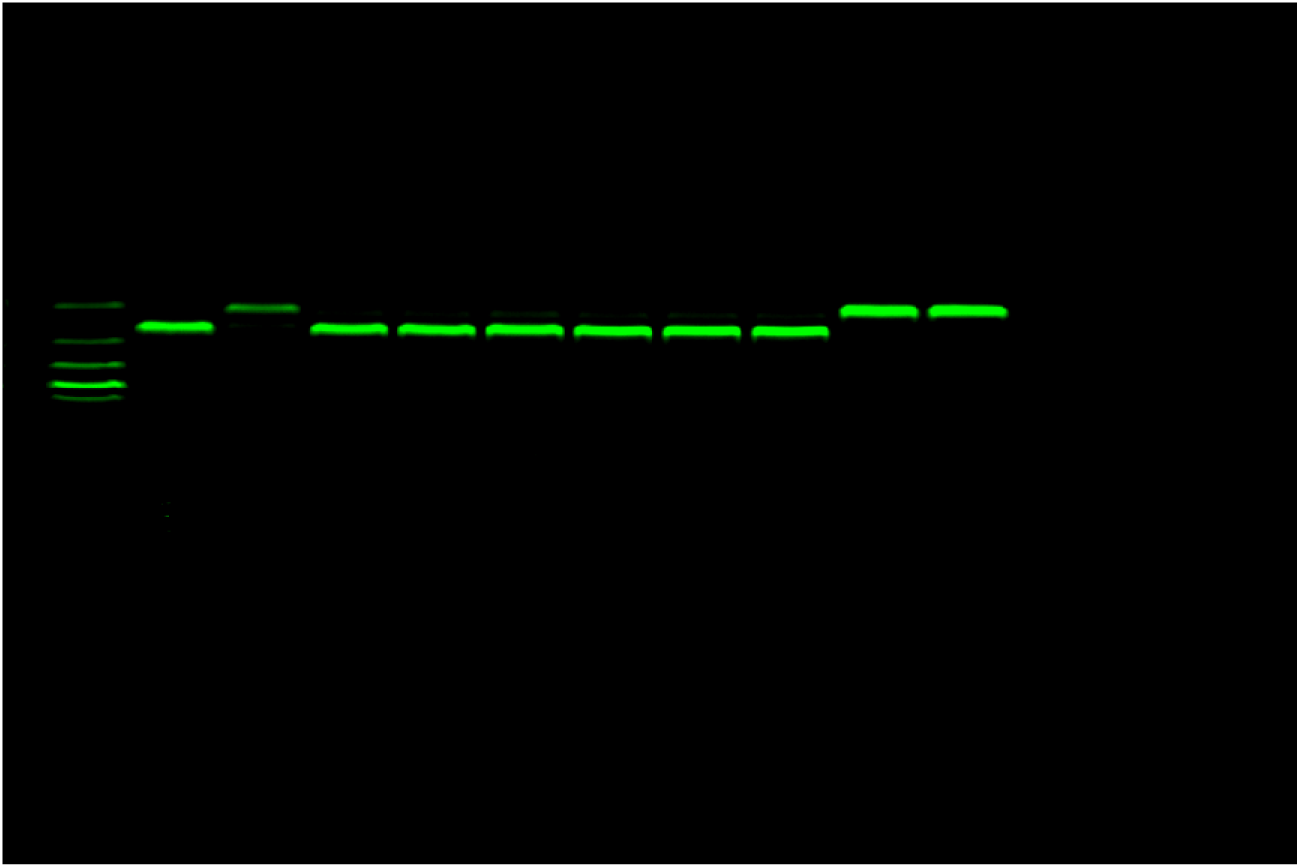
Uncropped agarose gel from Supplementary Fig. S11C.

**Supplementary Figure 41.**
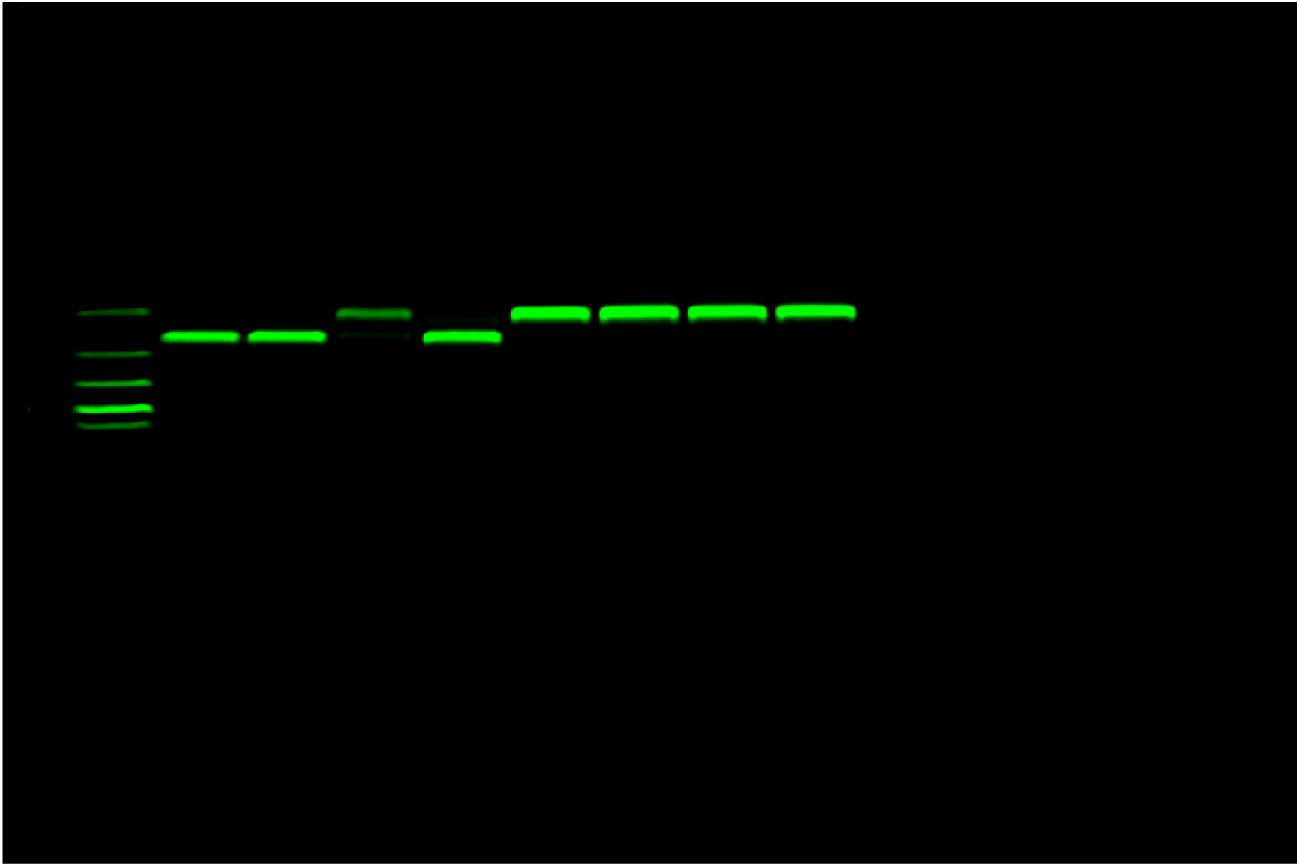
Uncropped agarose gel from Supplementary Fig. S15C.

**Supplementary Figure 42.**
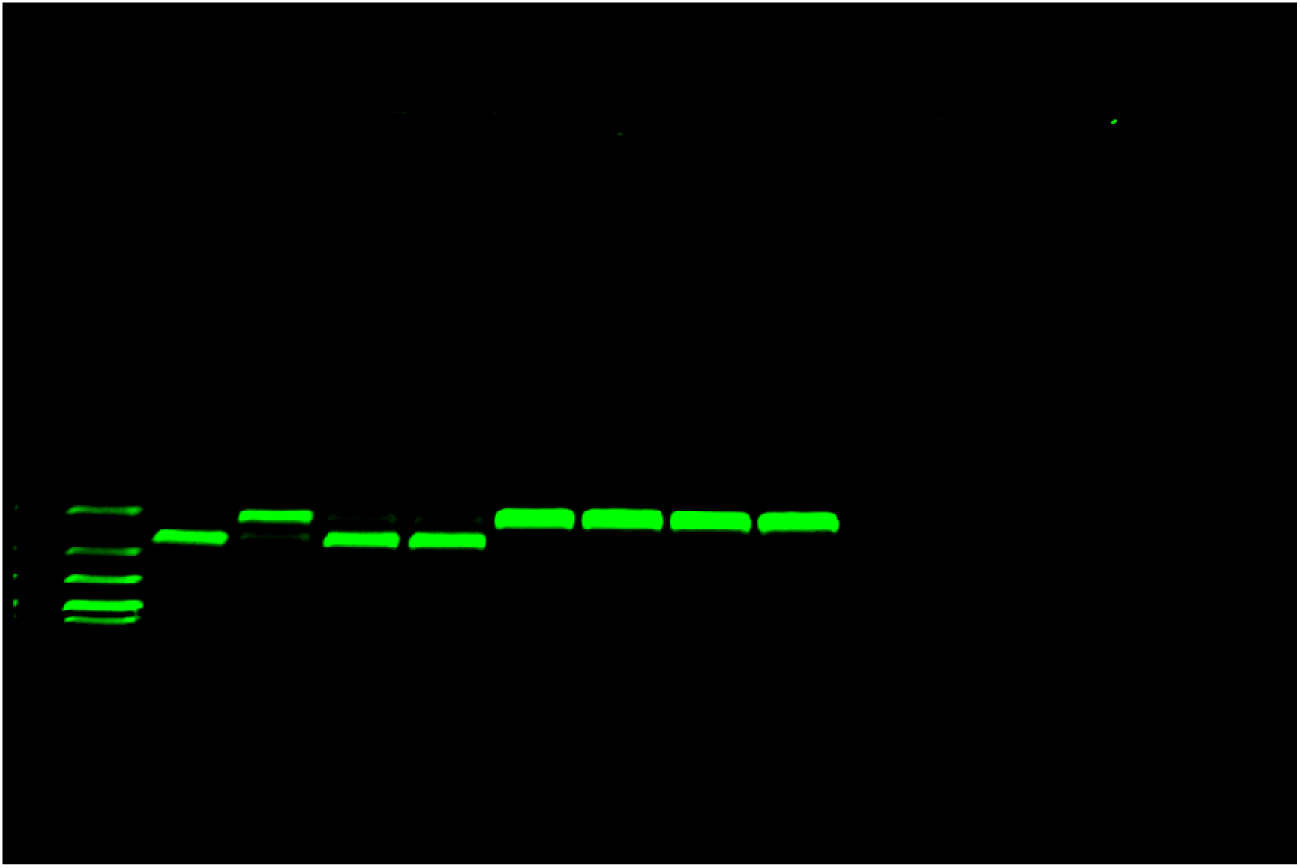
Uncropped agarose gel from Supplementary Fig. S16C.

**Supplementary Figure 43.**
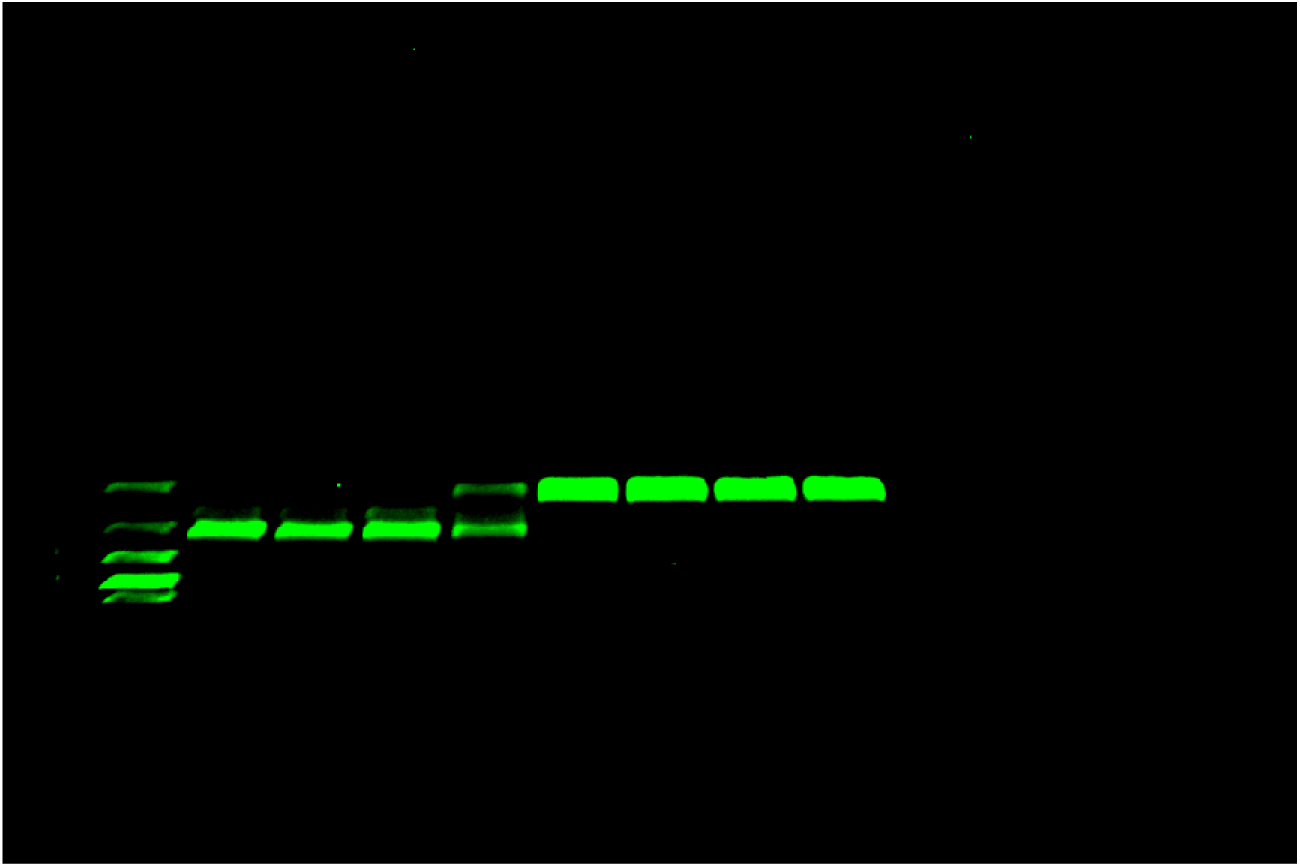
Uncropped agarose gel from Supplementary Fig. S17C.

**Supplementary Figure 44.**
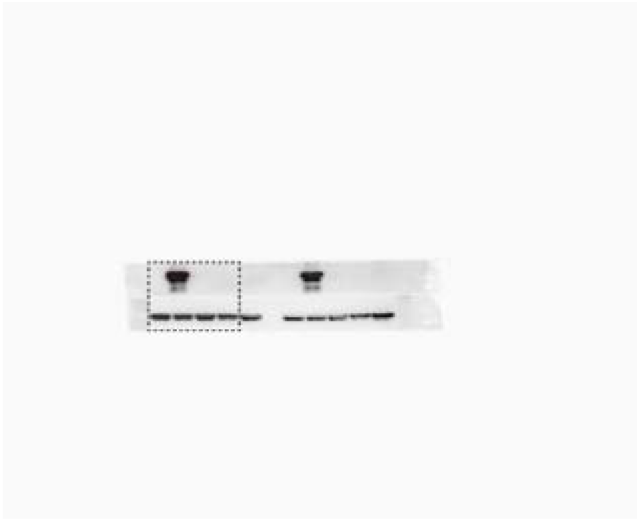
Uncropped Western blot from Supplementary Fig. S33. Dashed lines indicate the region shown in the main figure.

## Supplementary Note 1. Plasmid Sequences

Plasmids will be deposited at Addgene (https://www.addgene.org) for public distribution.

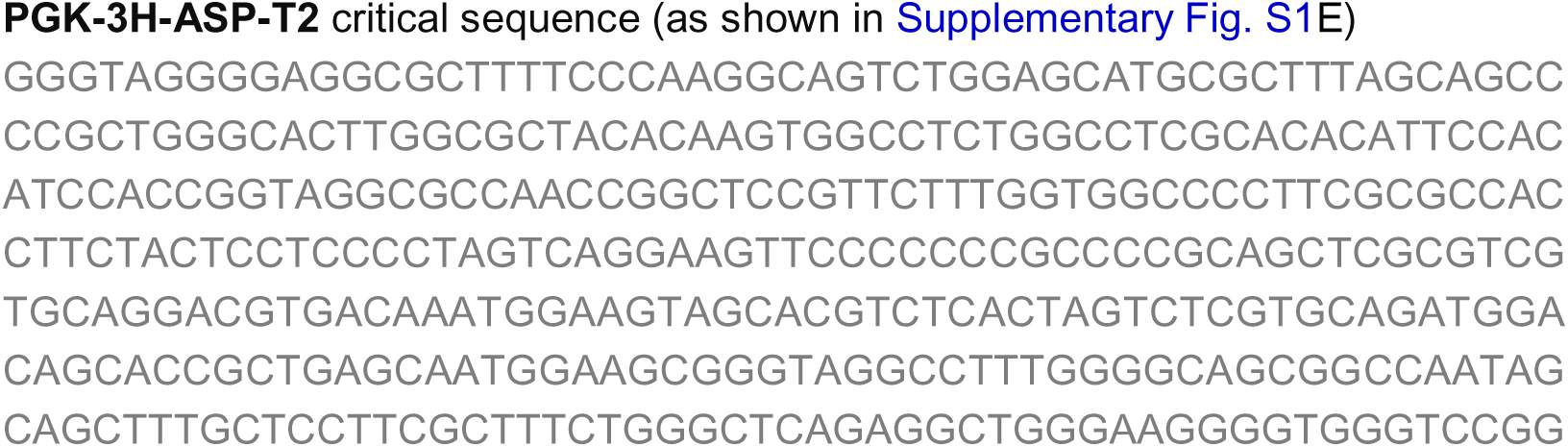

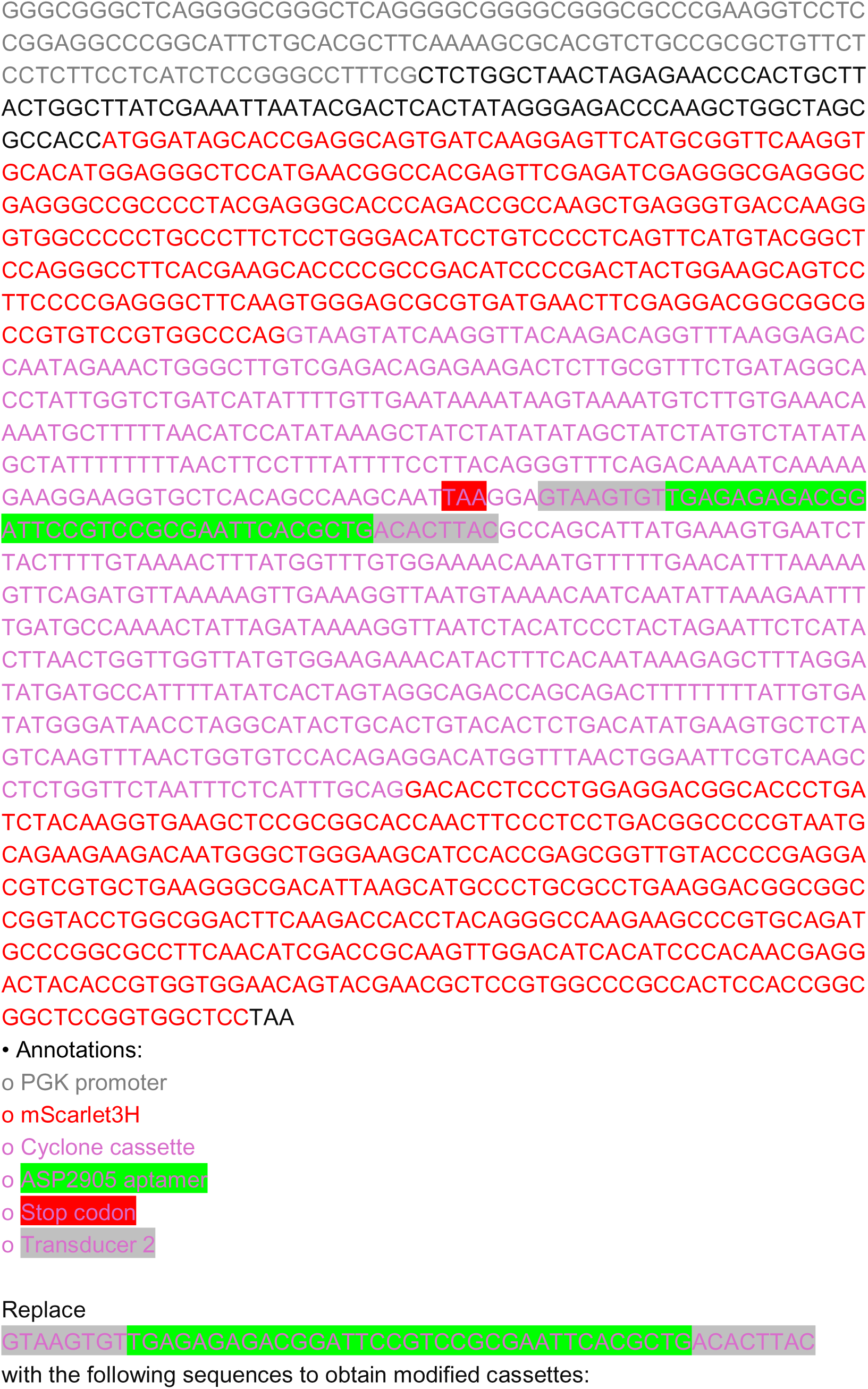

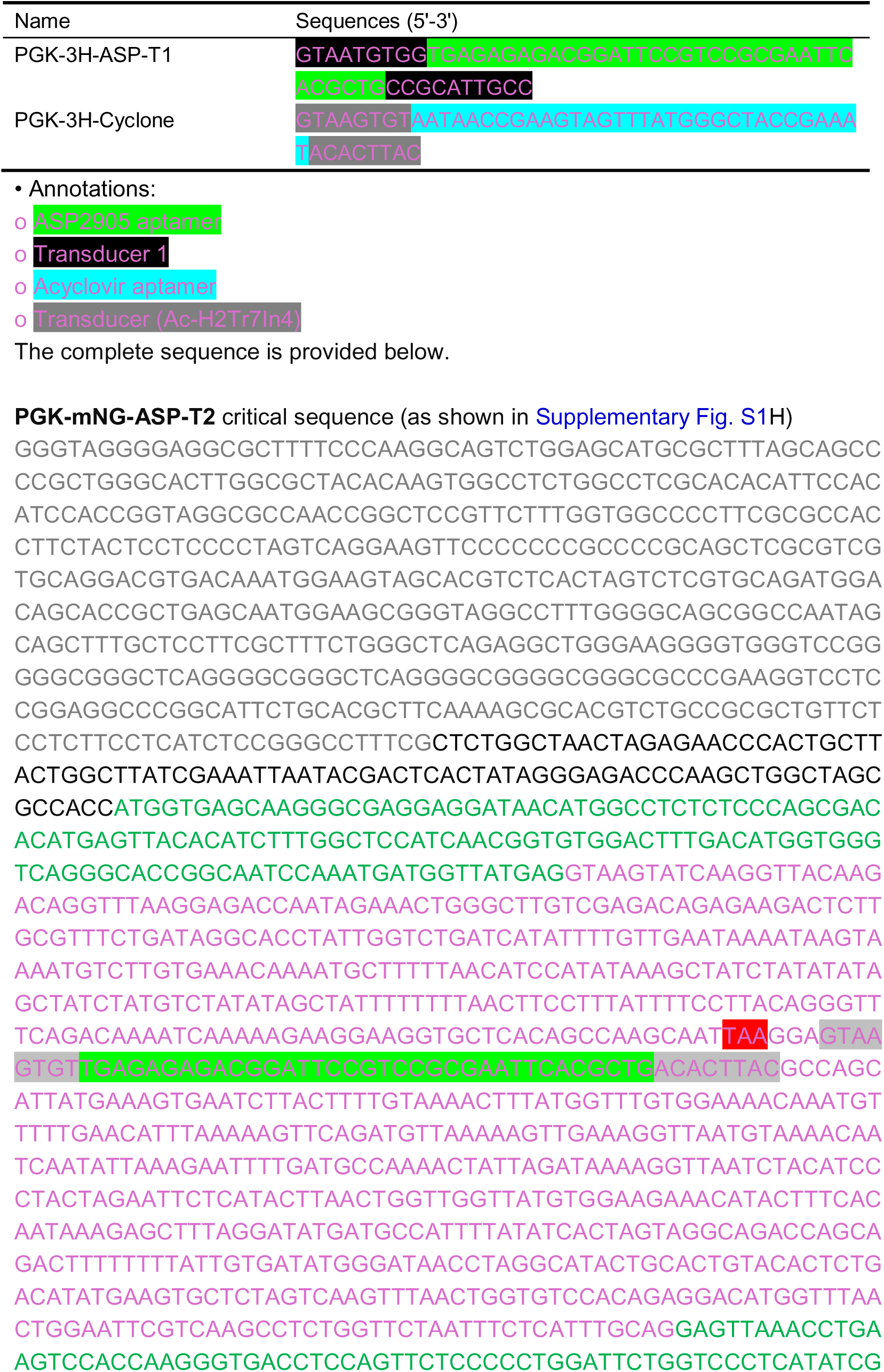

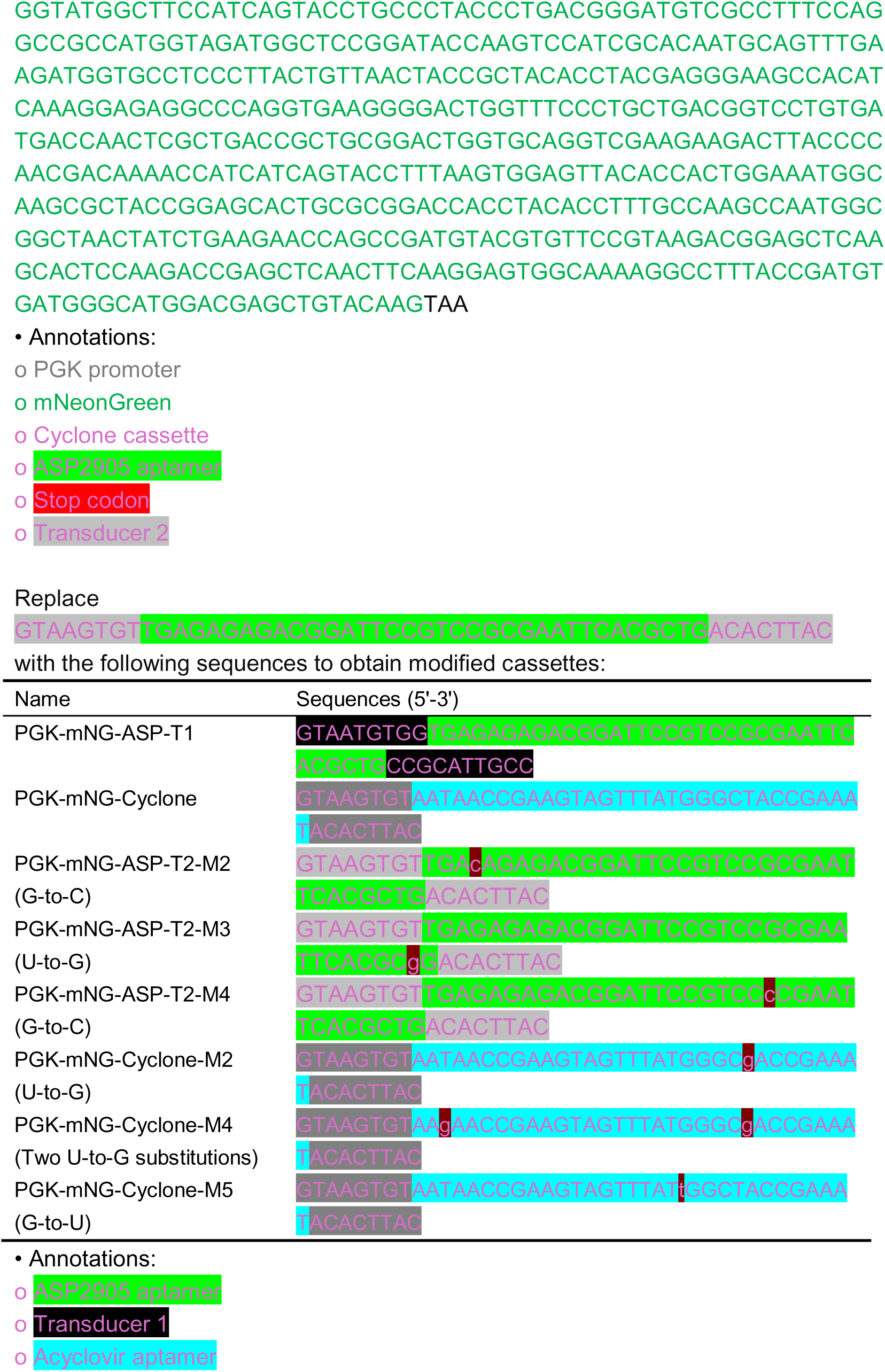

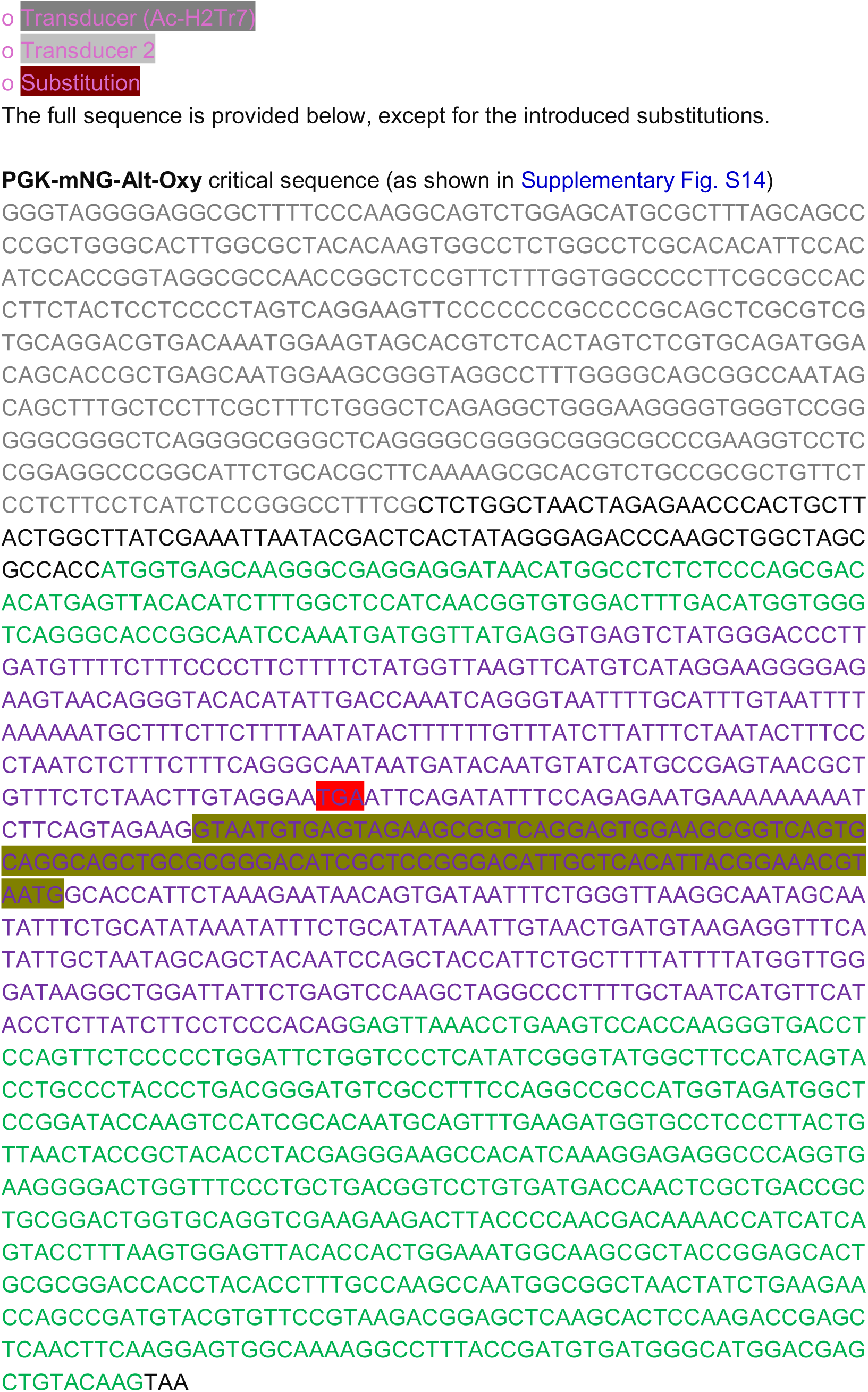

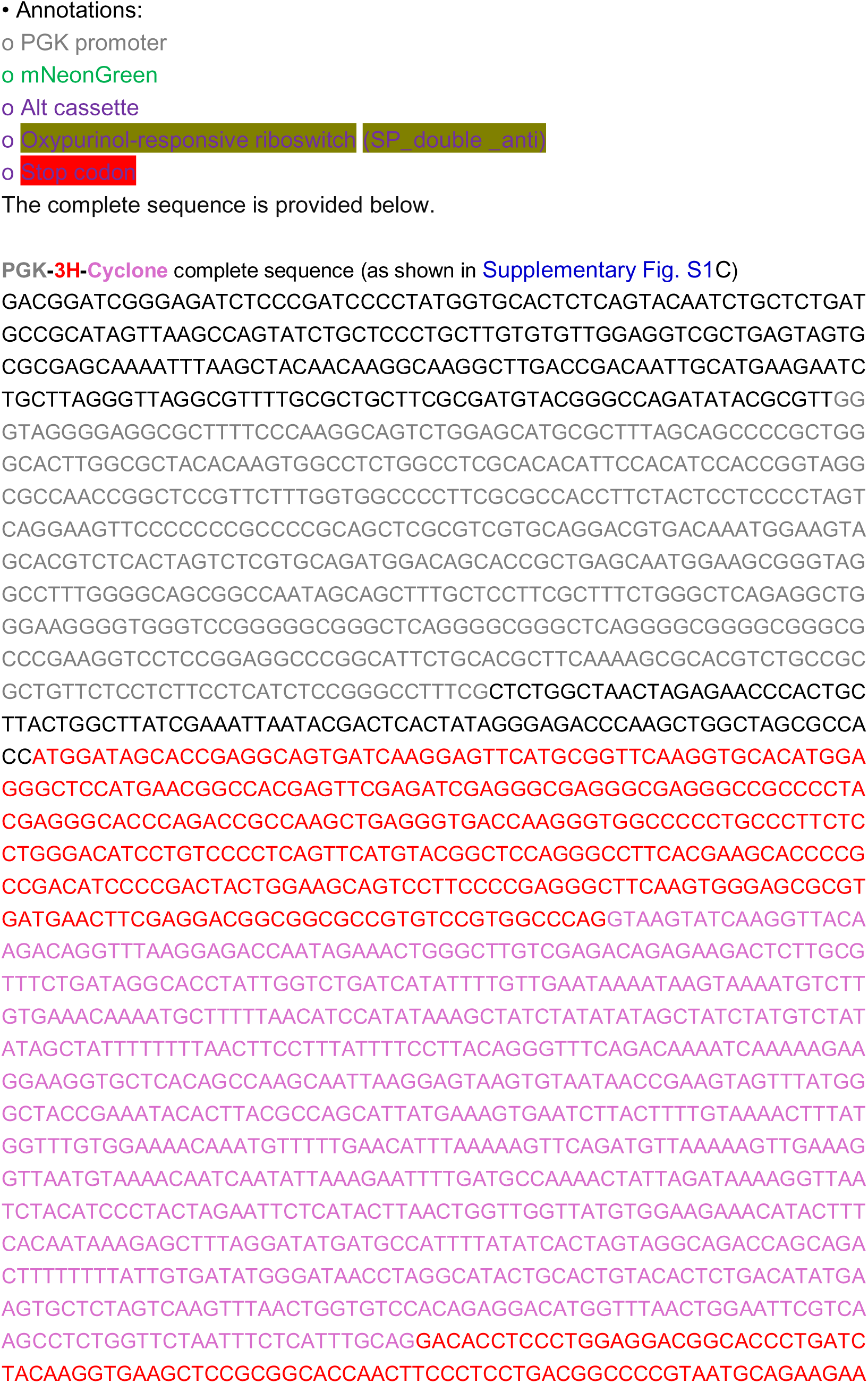

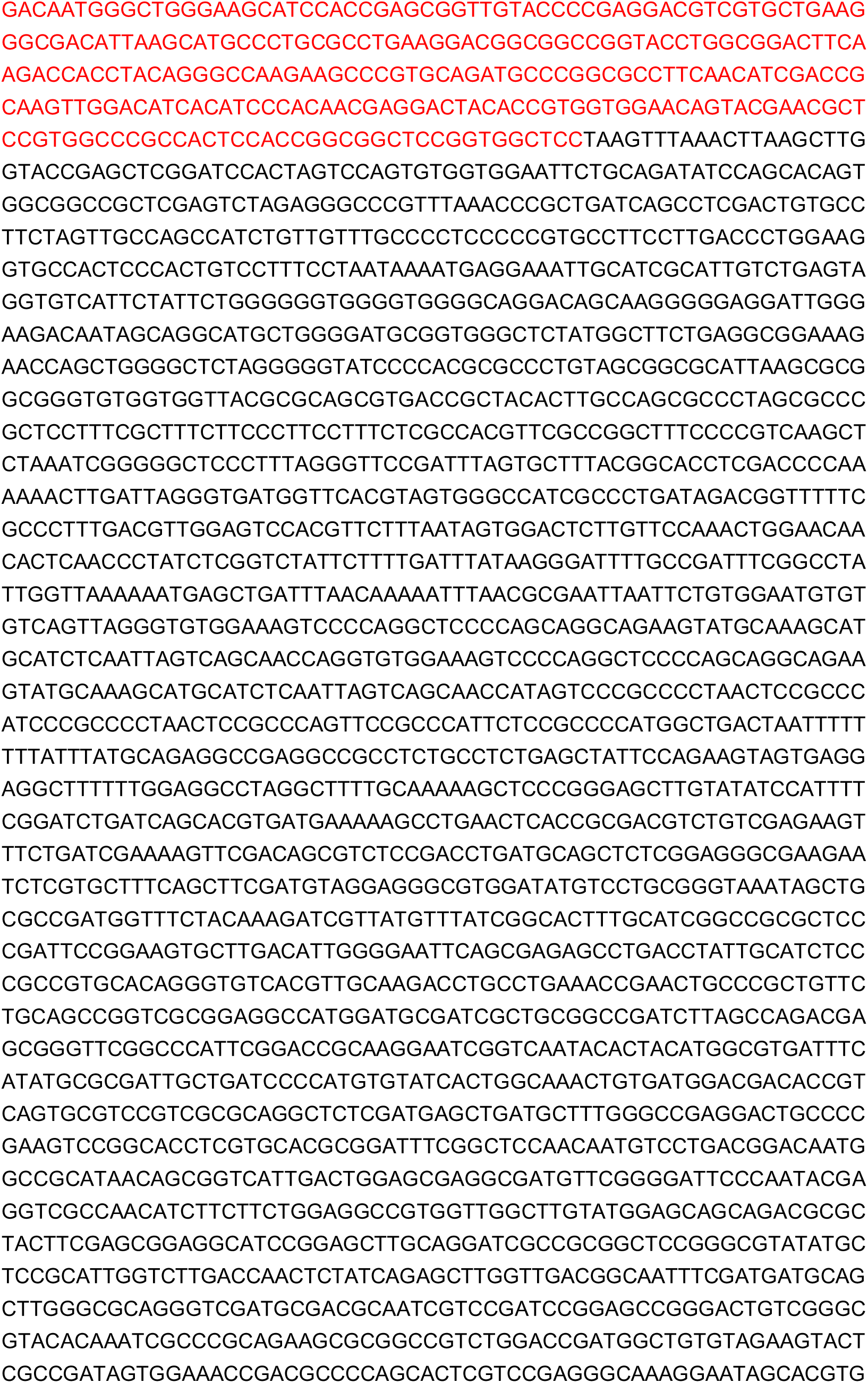

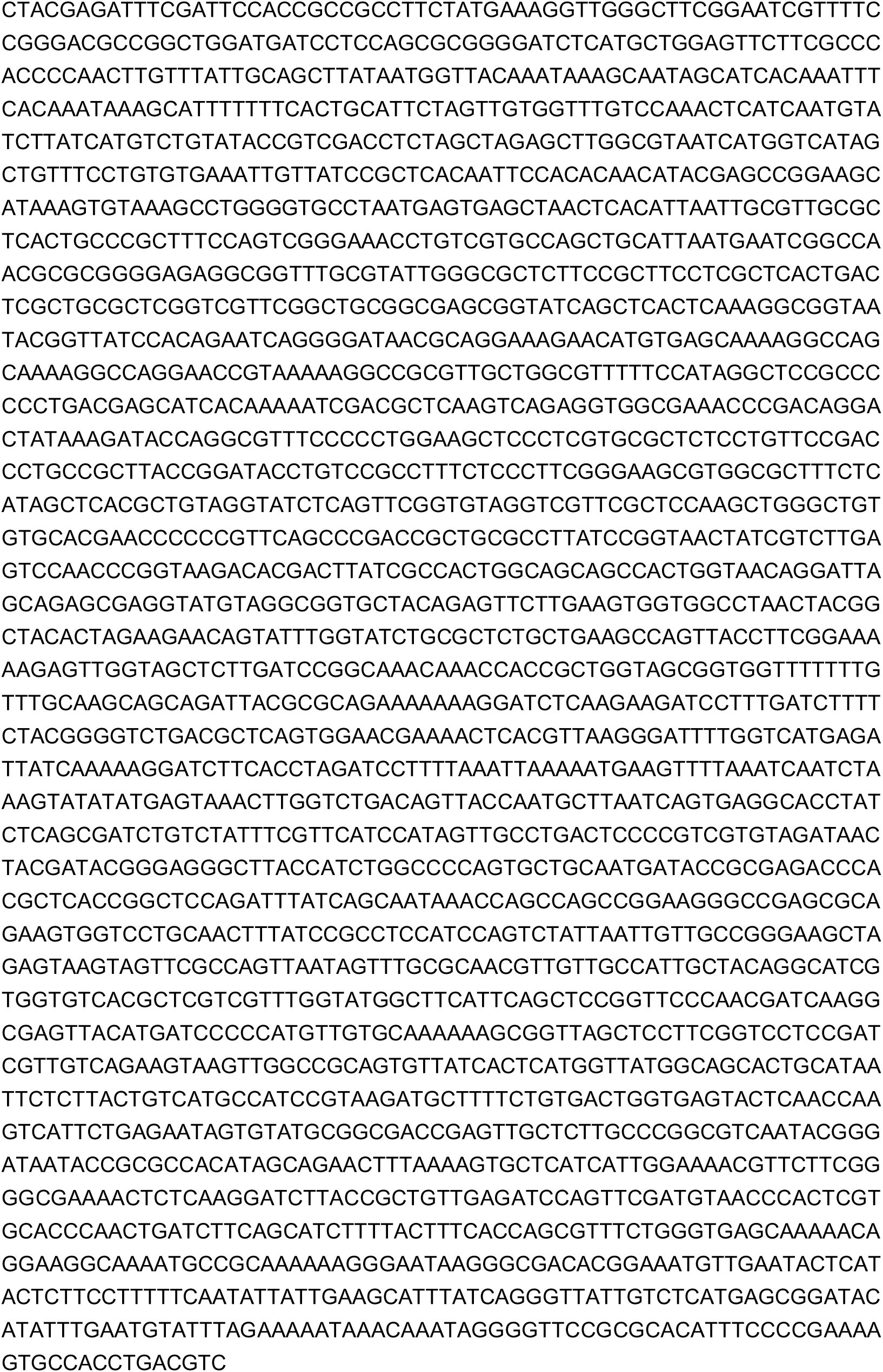

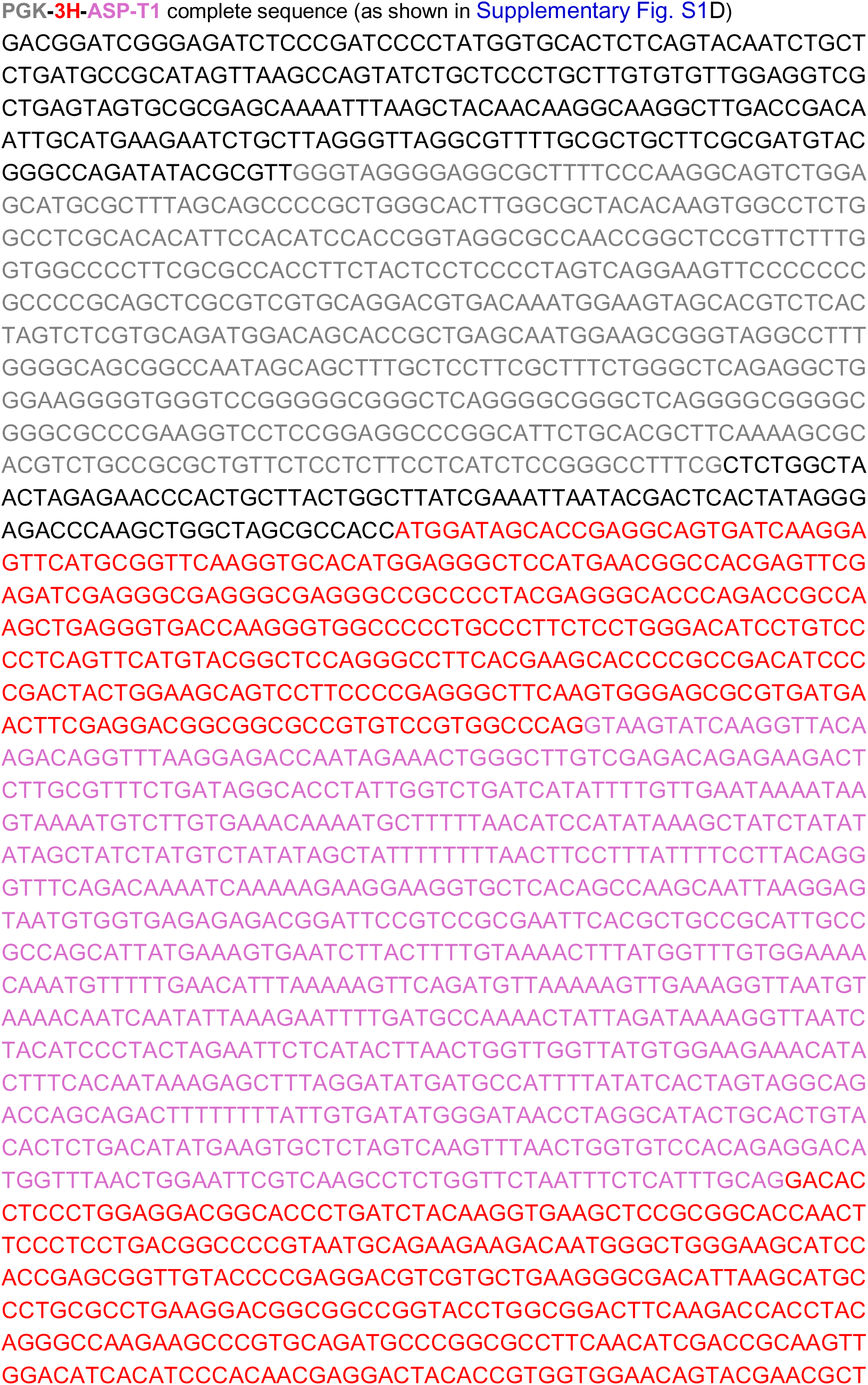

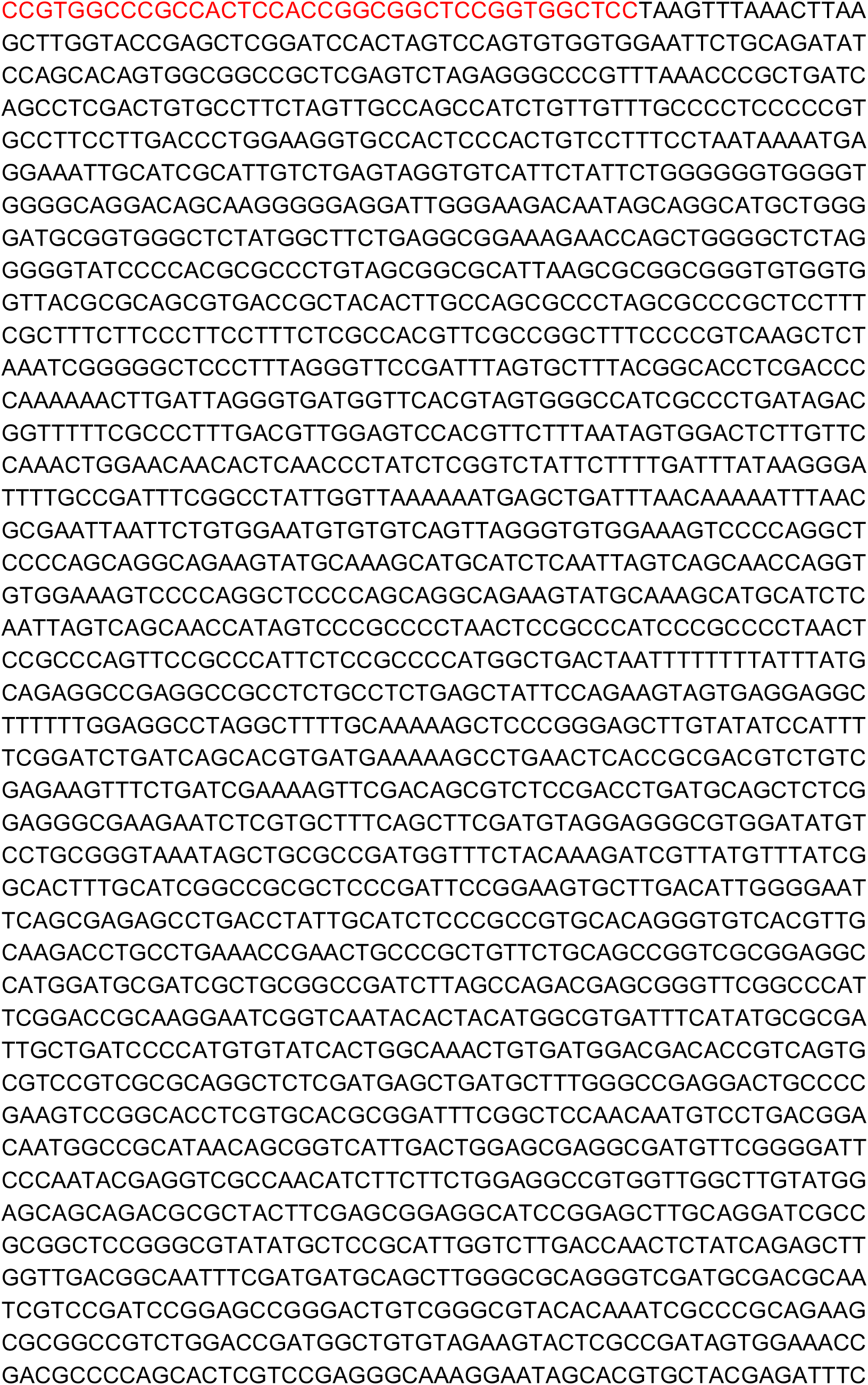

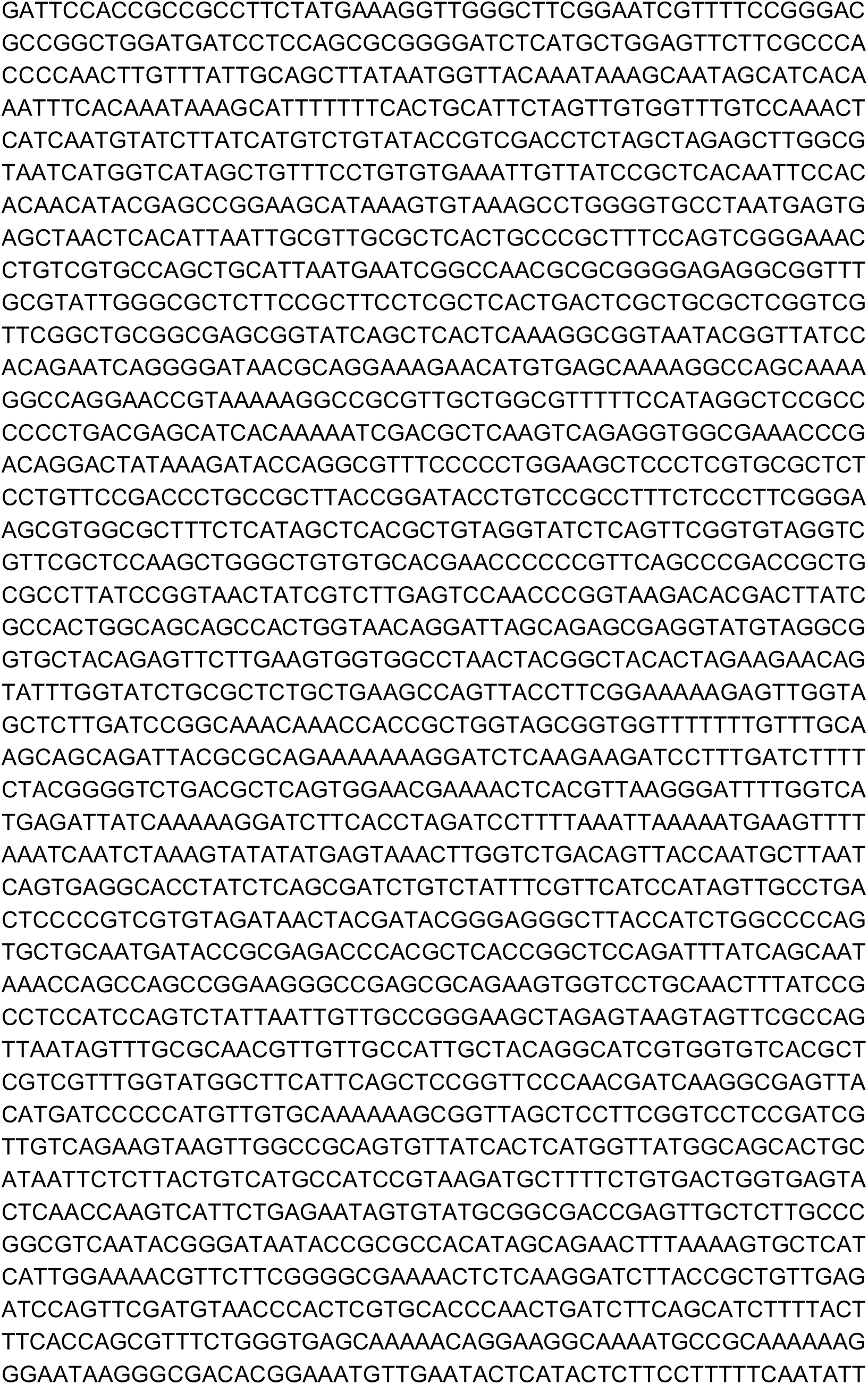

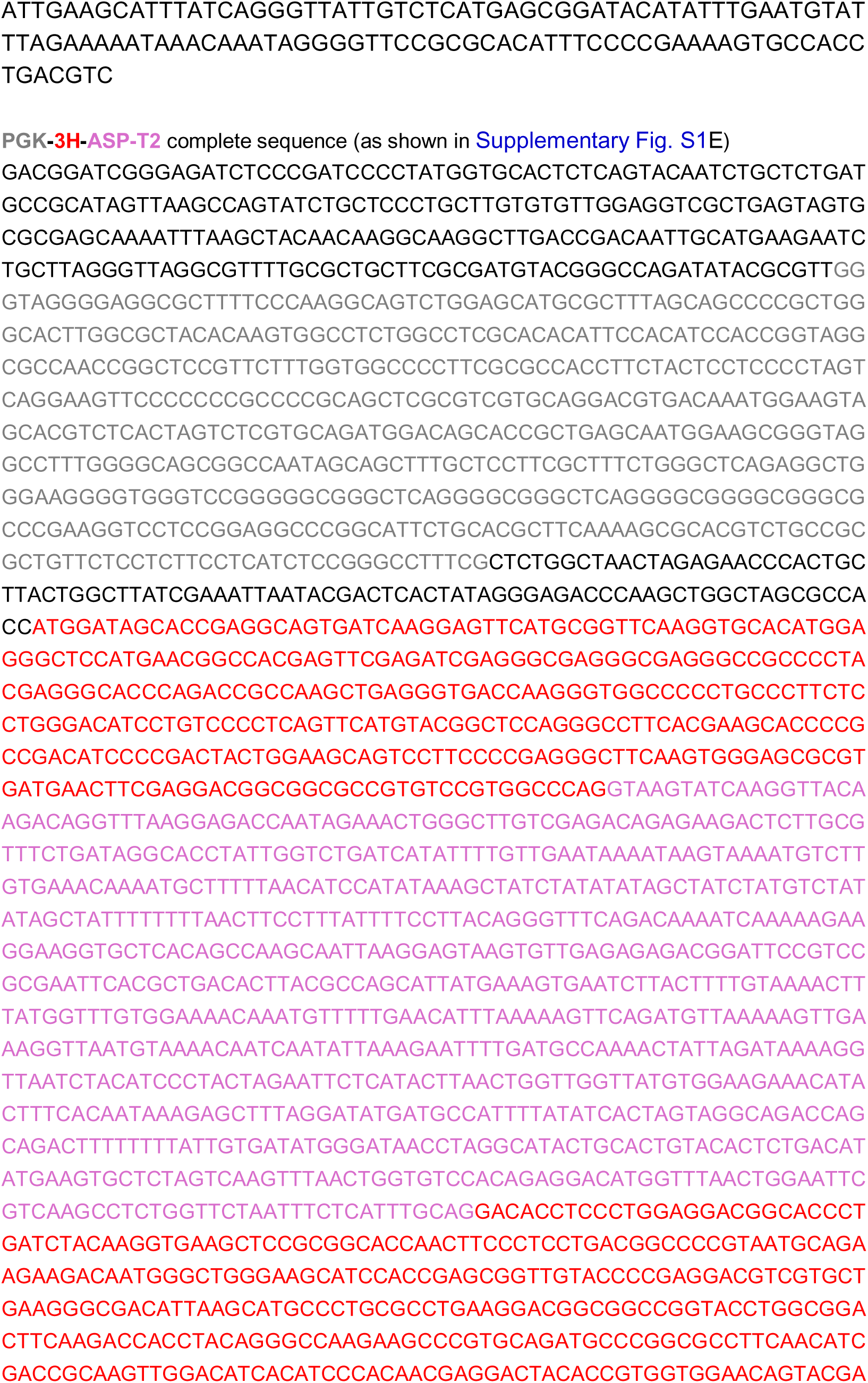

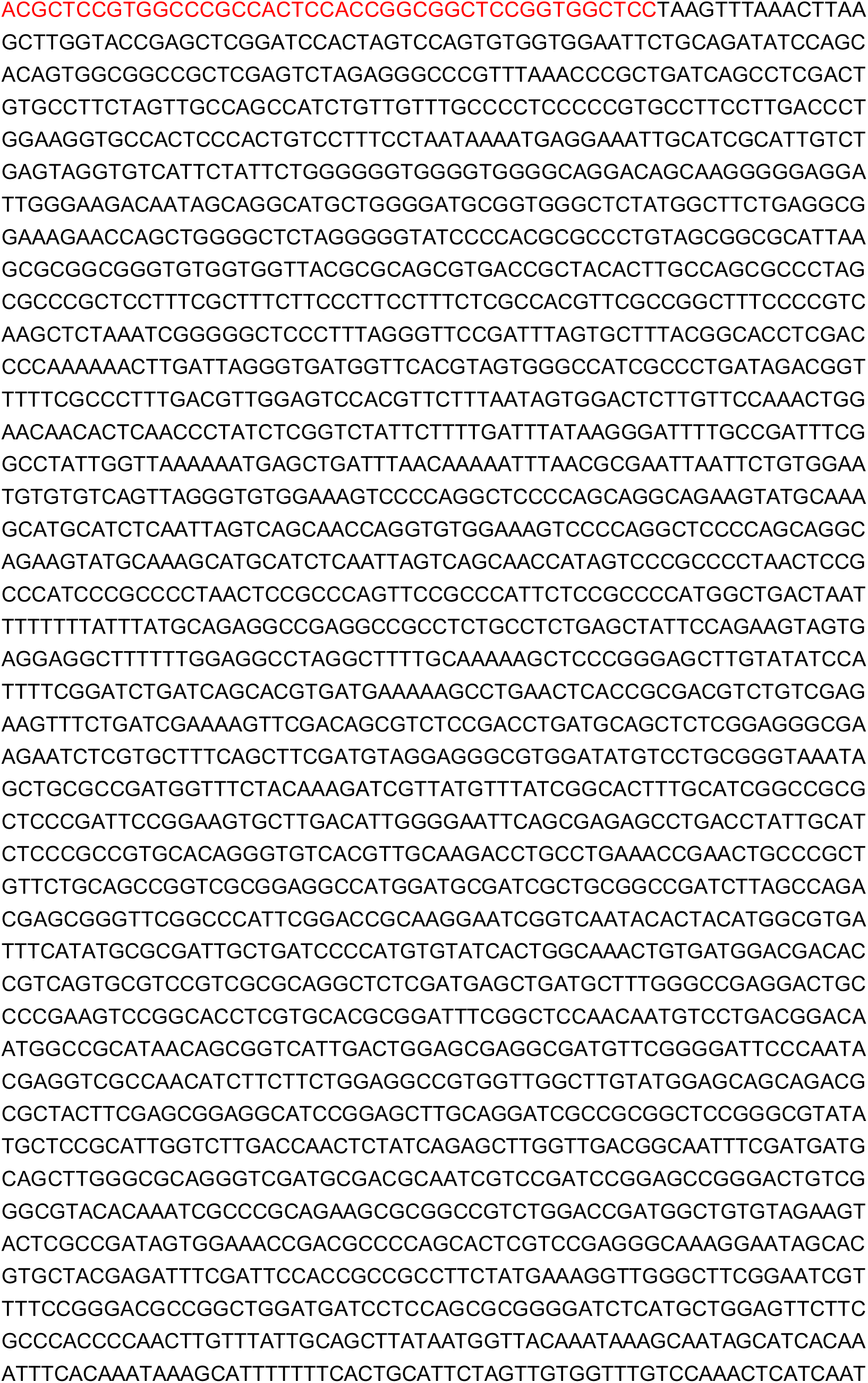

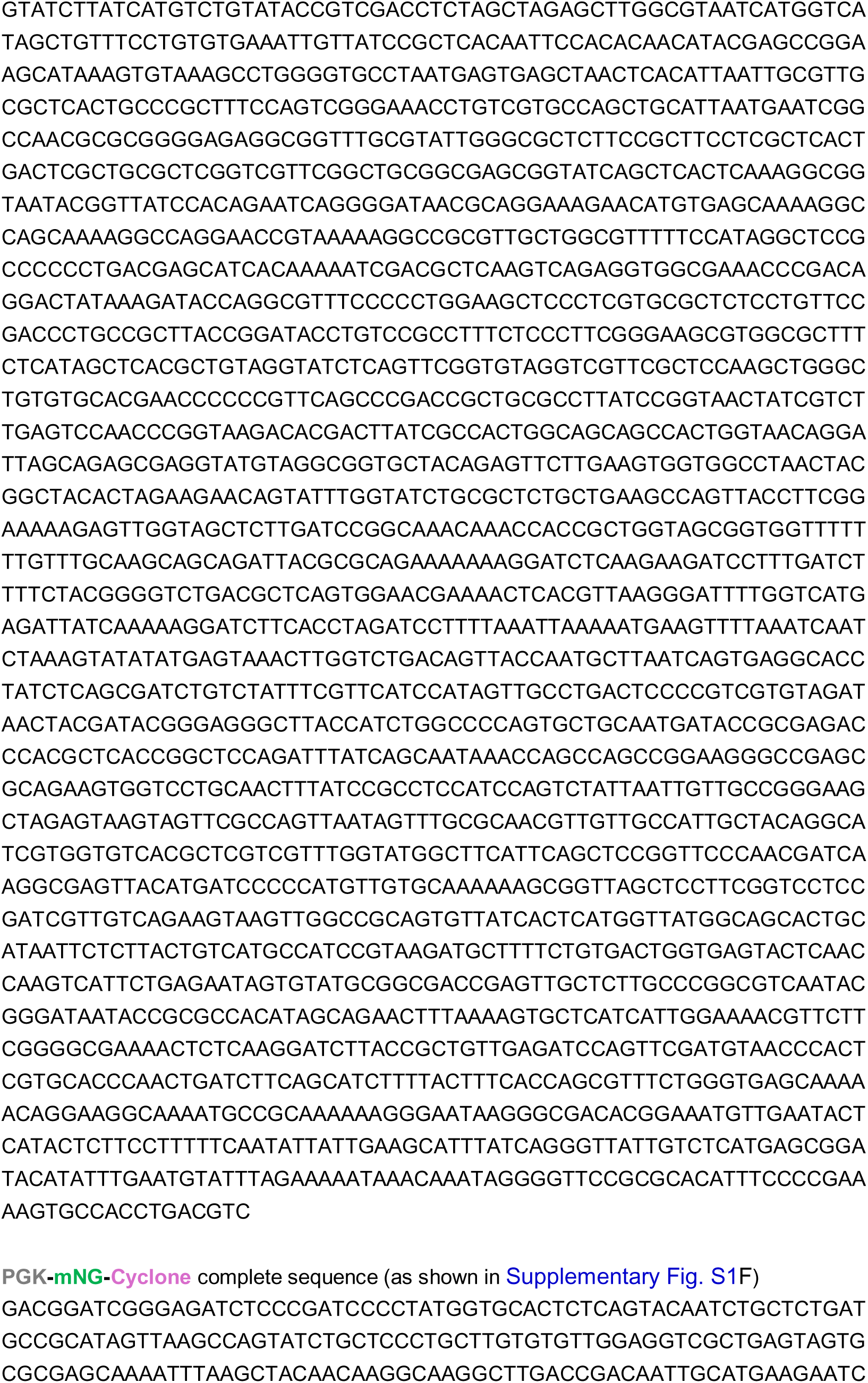

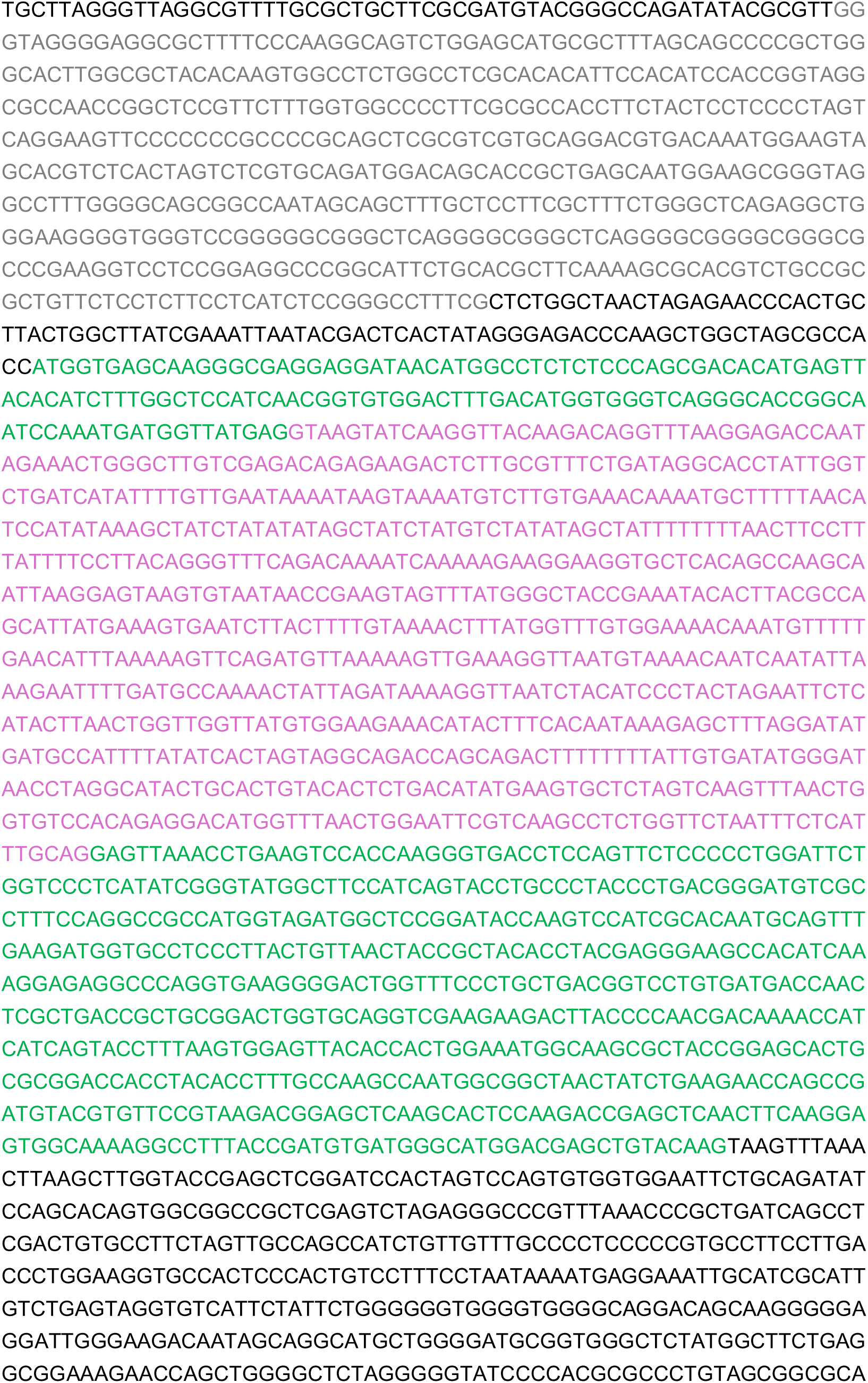

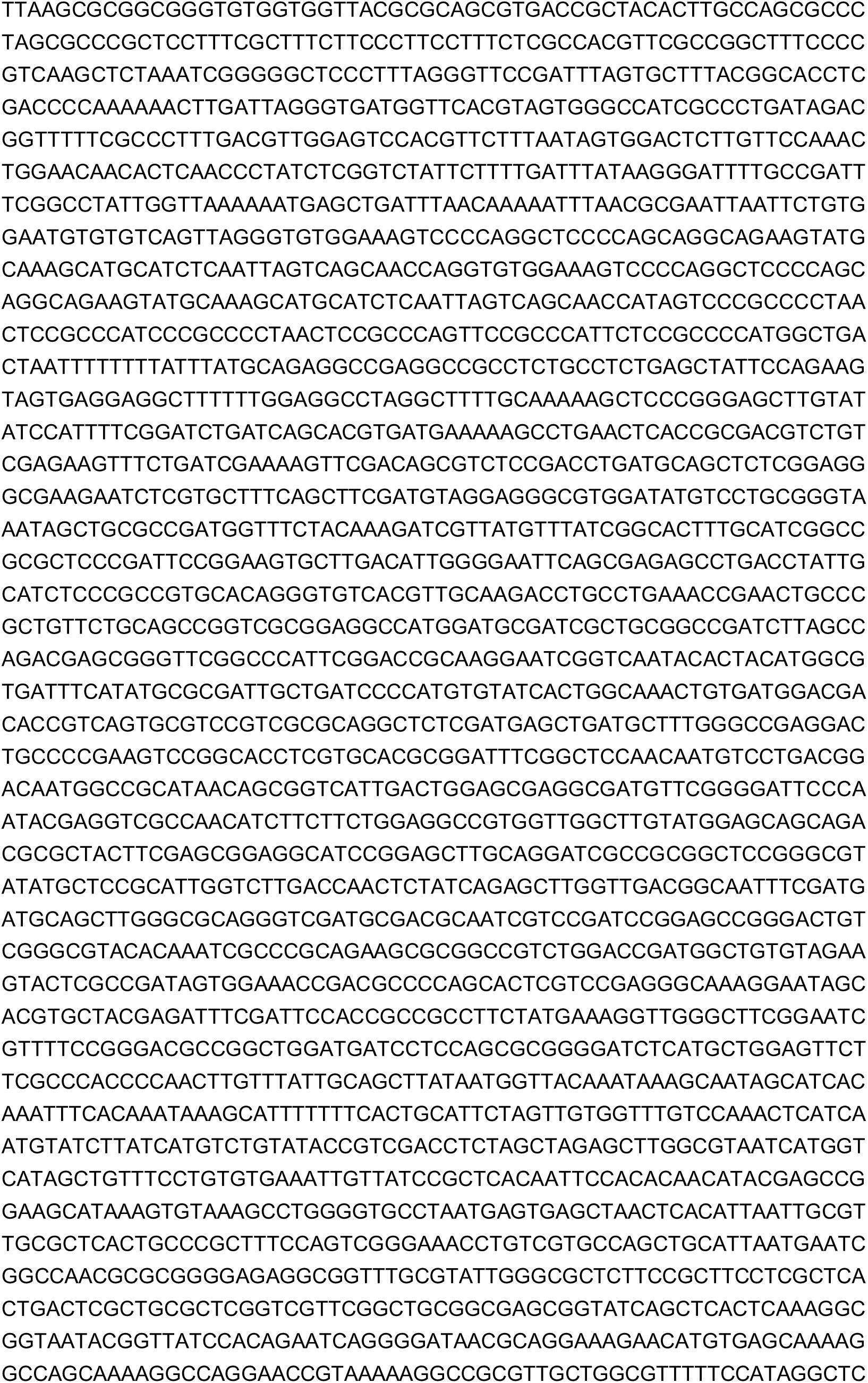

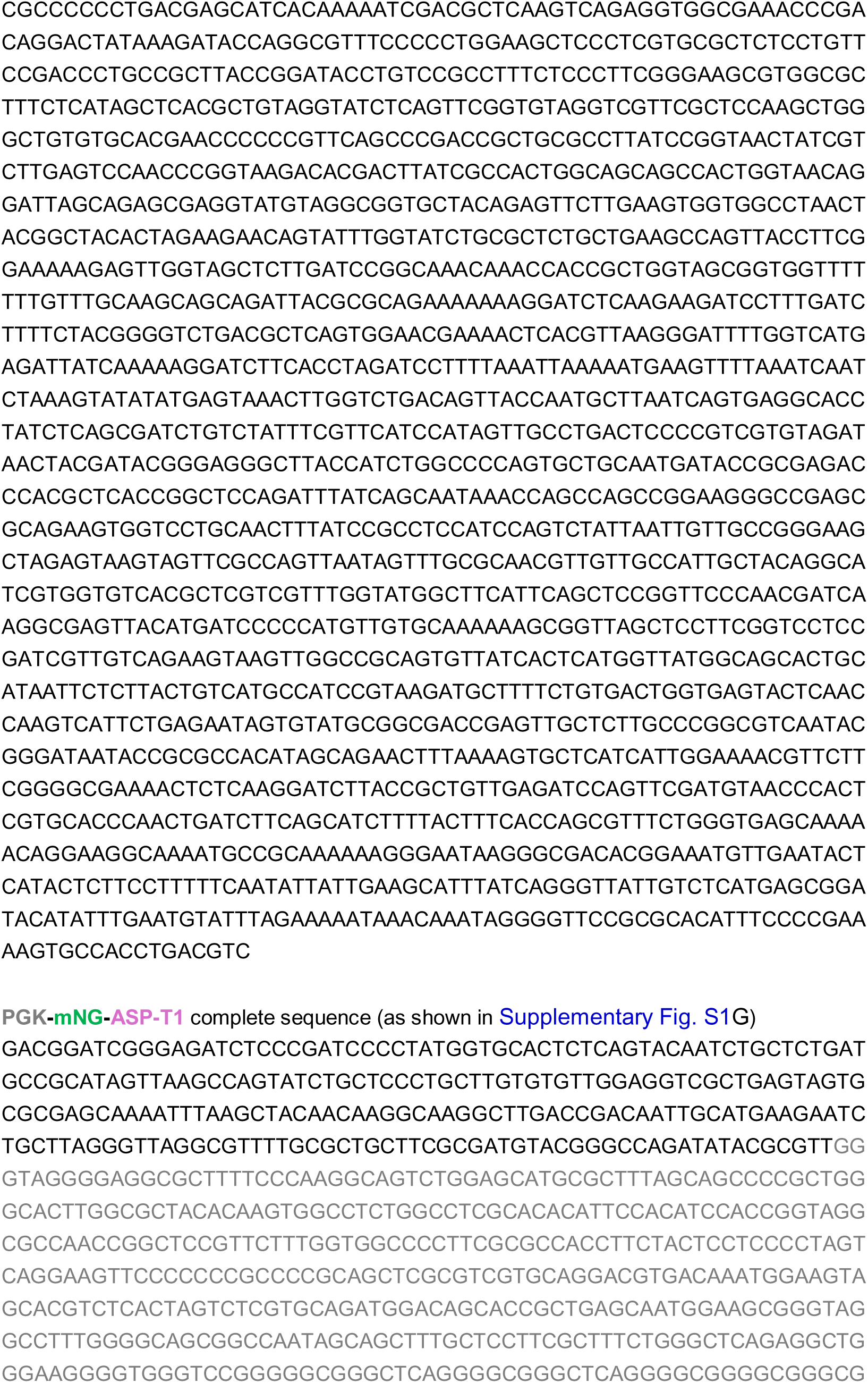

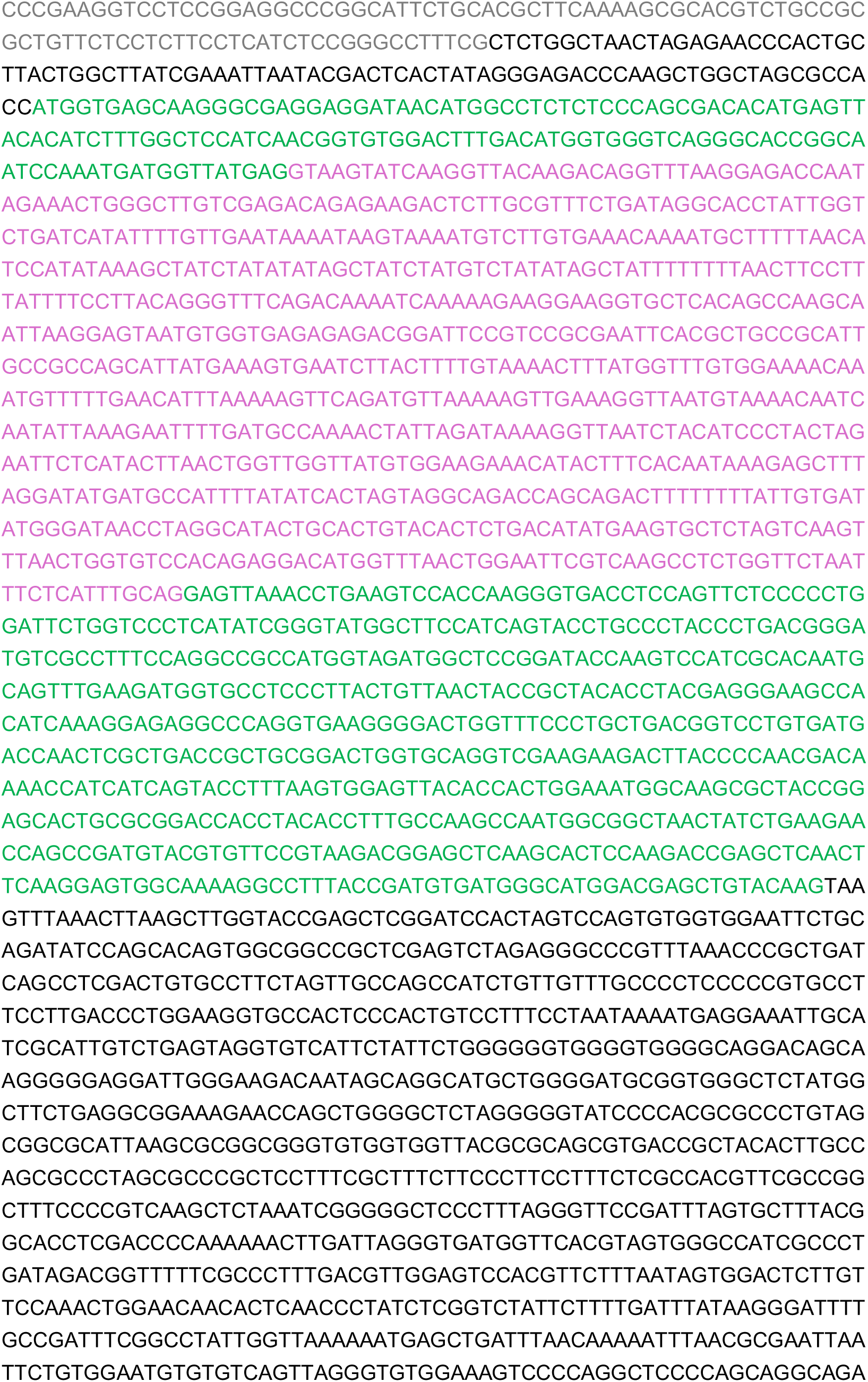

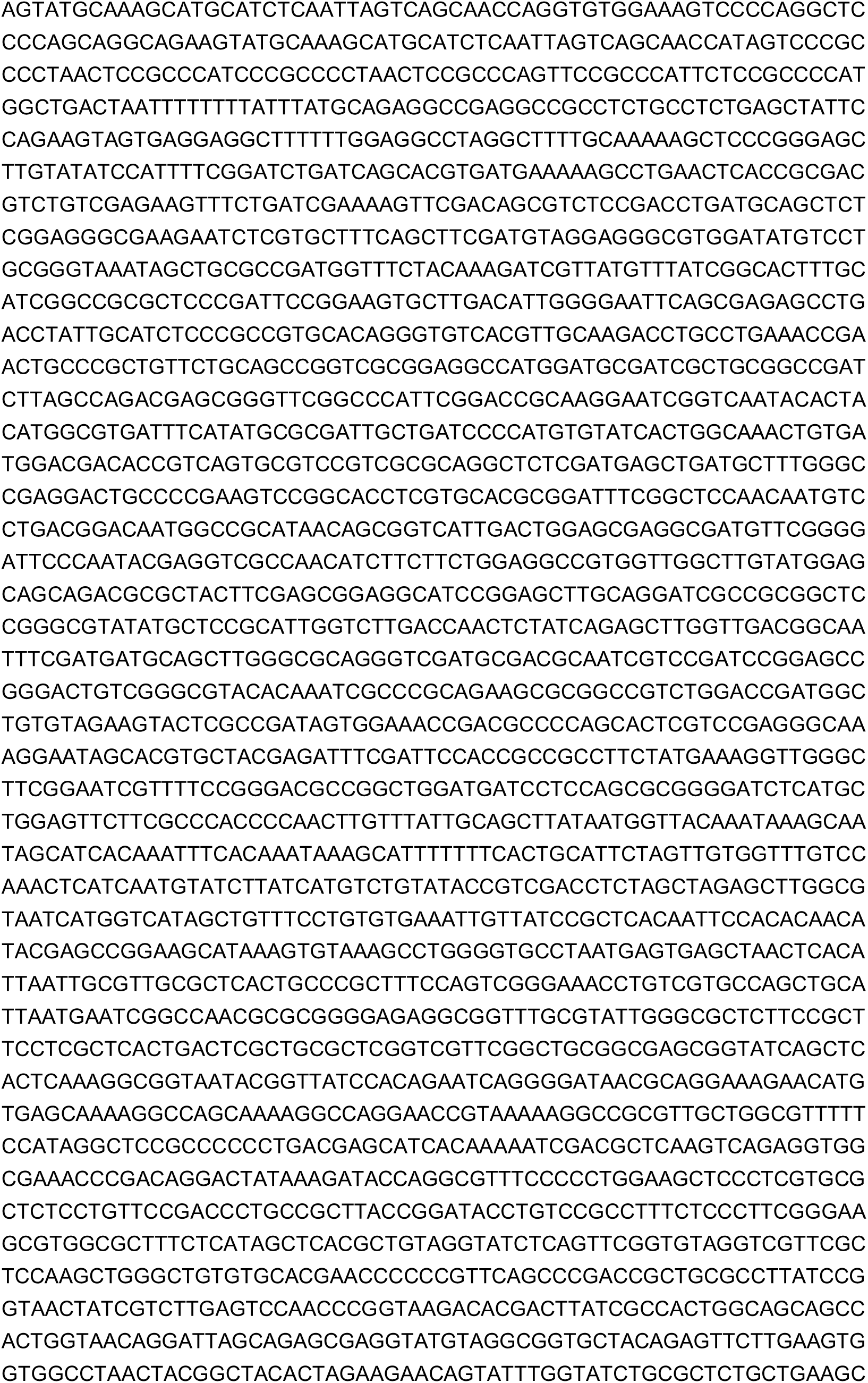

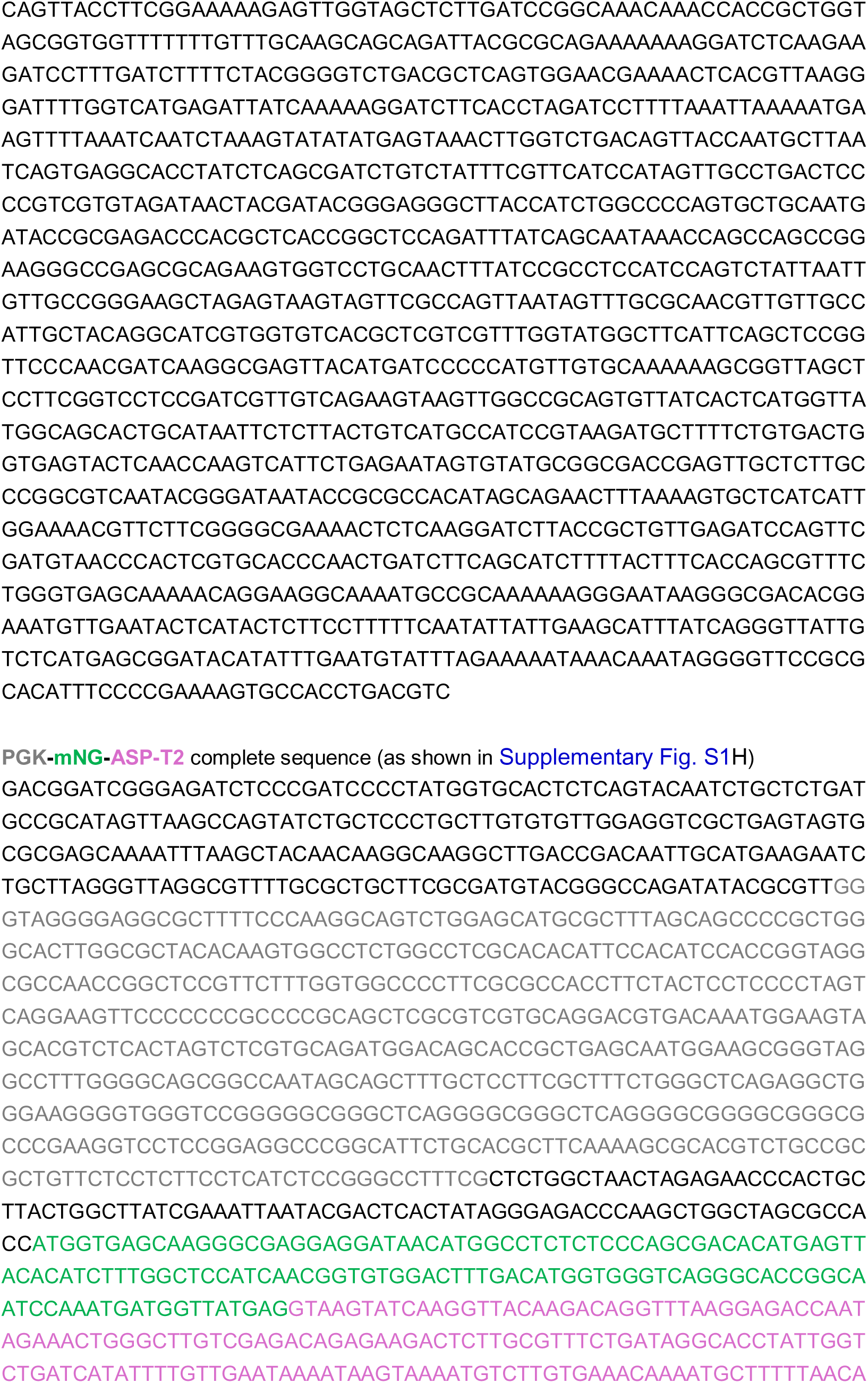

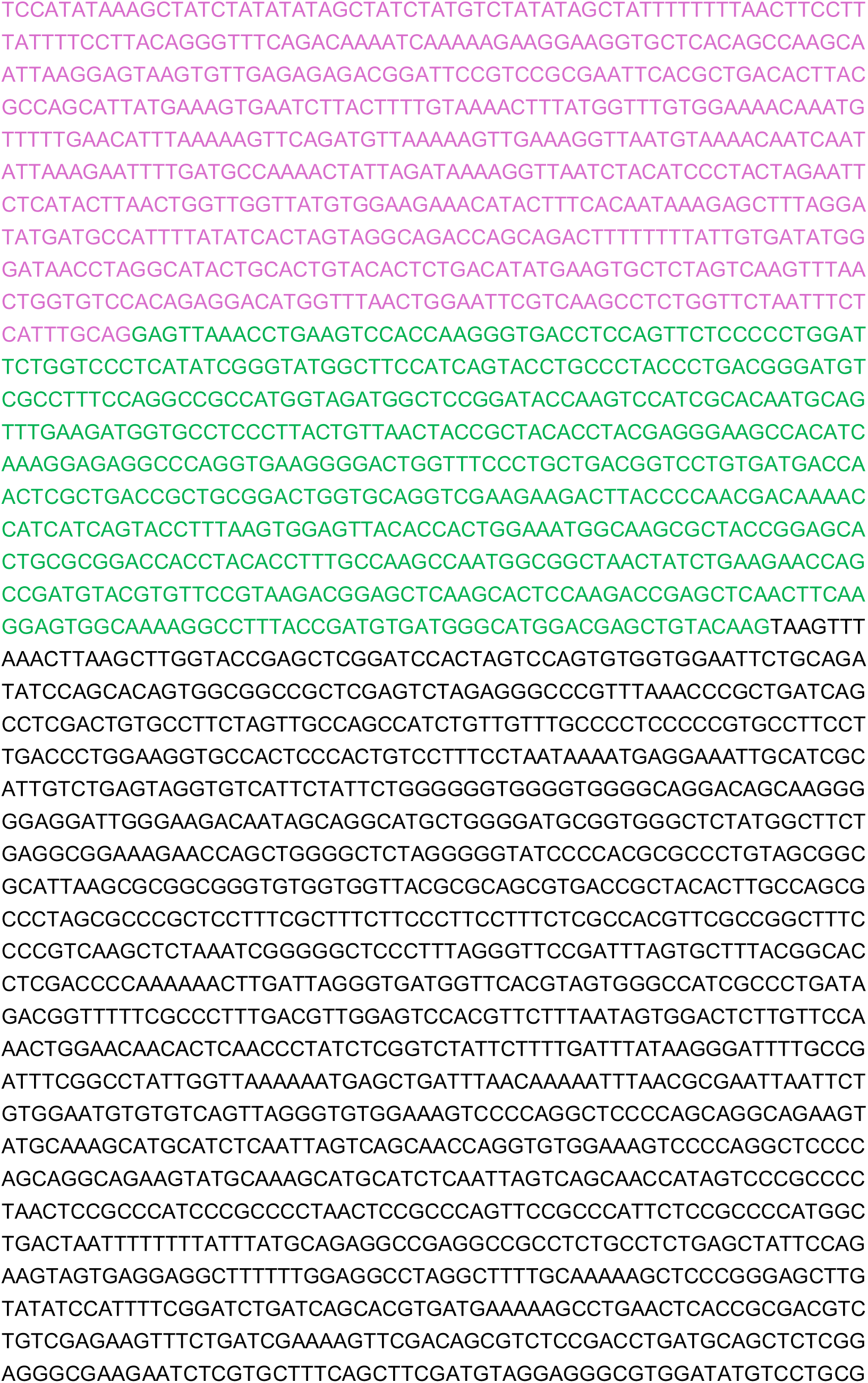

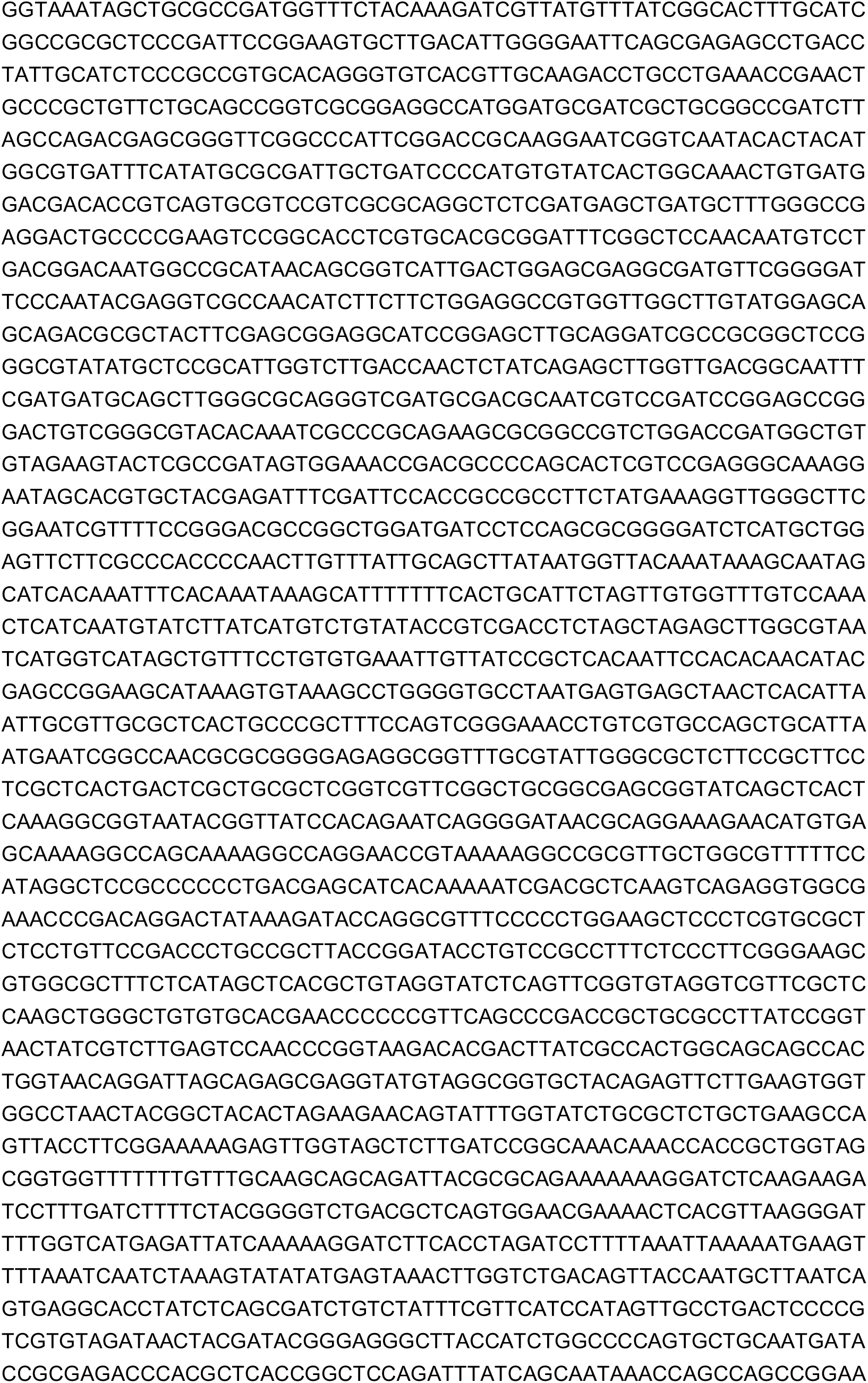

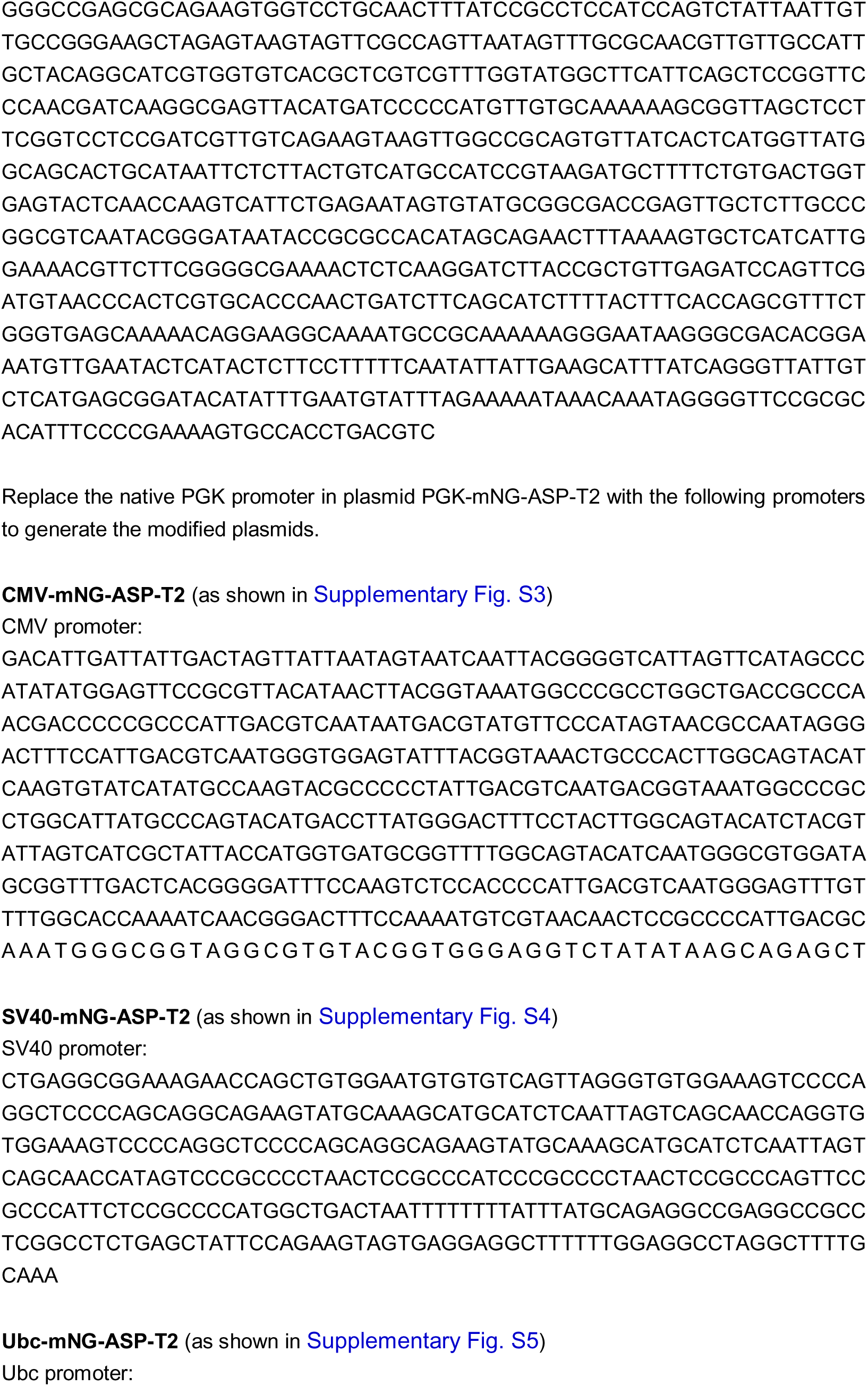

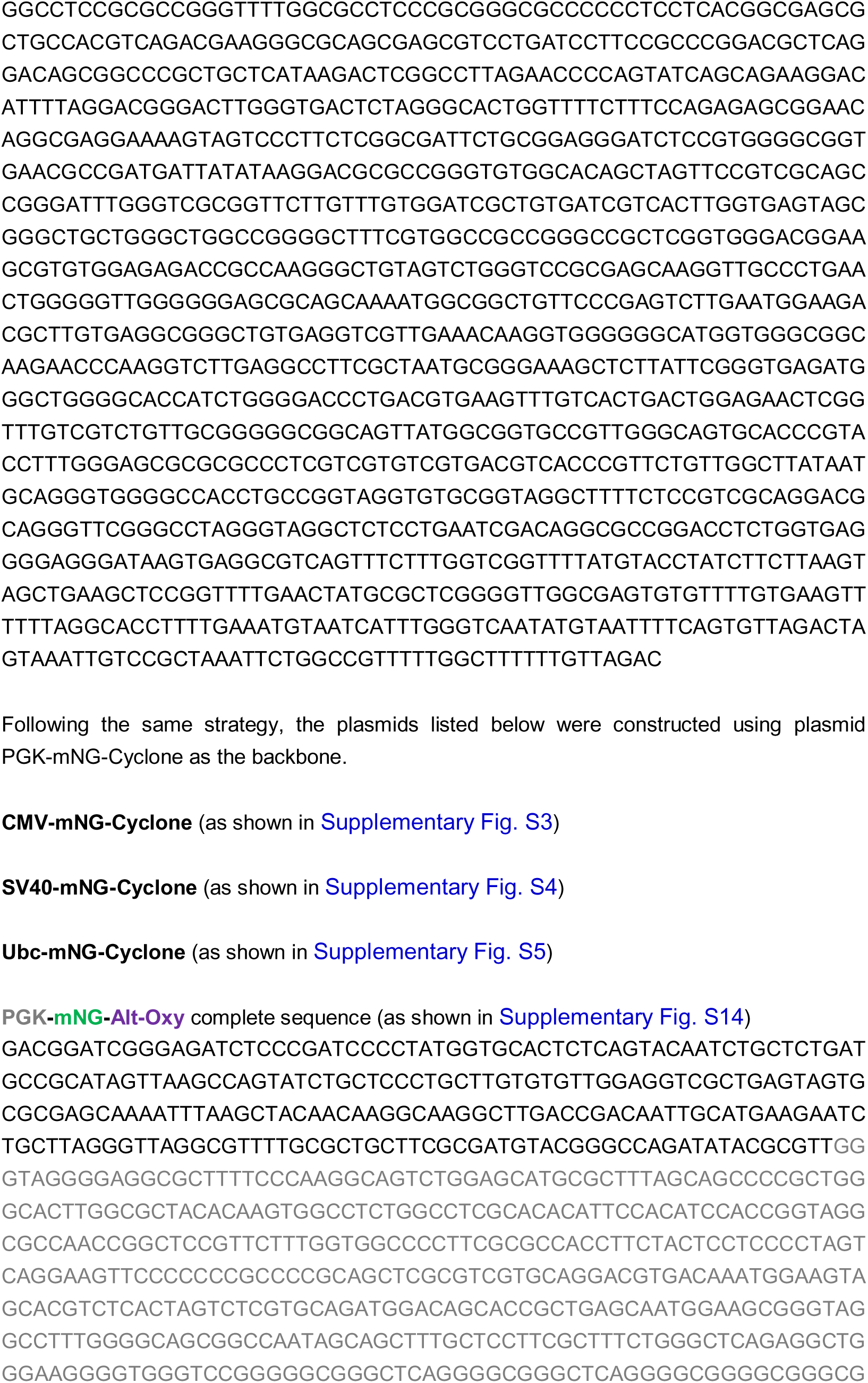

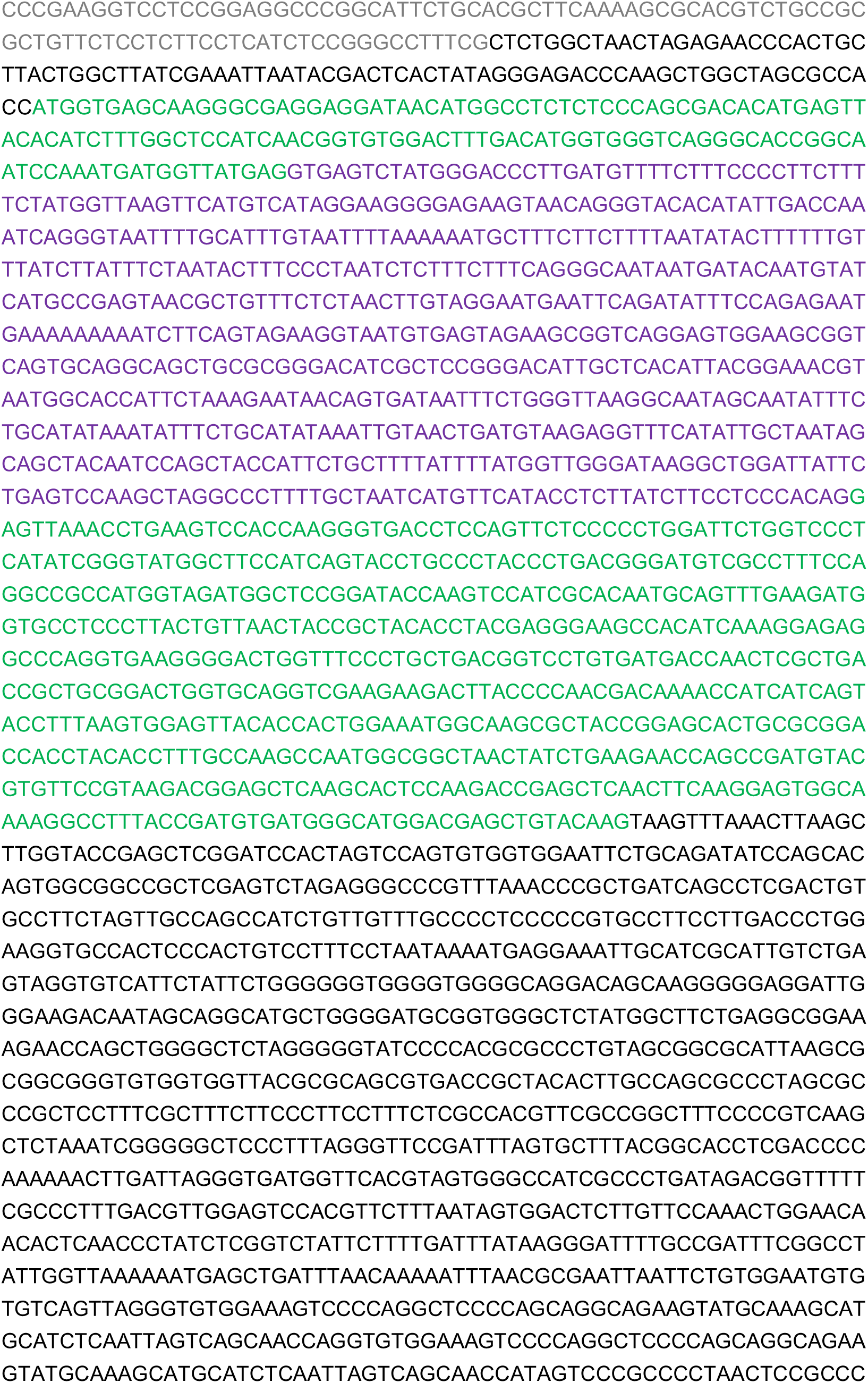

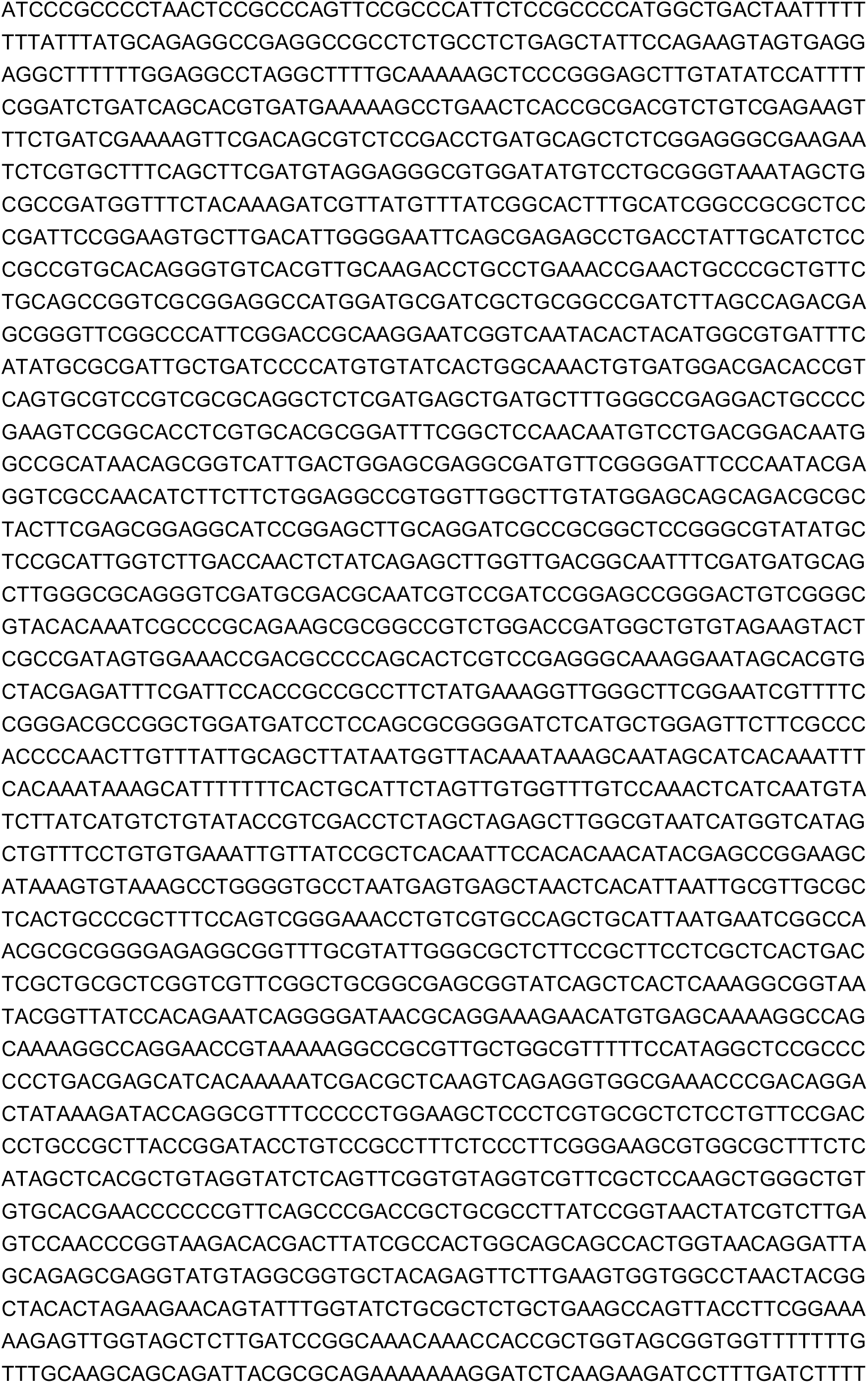

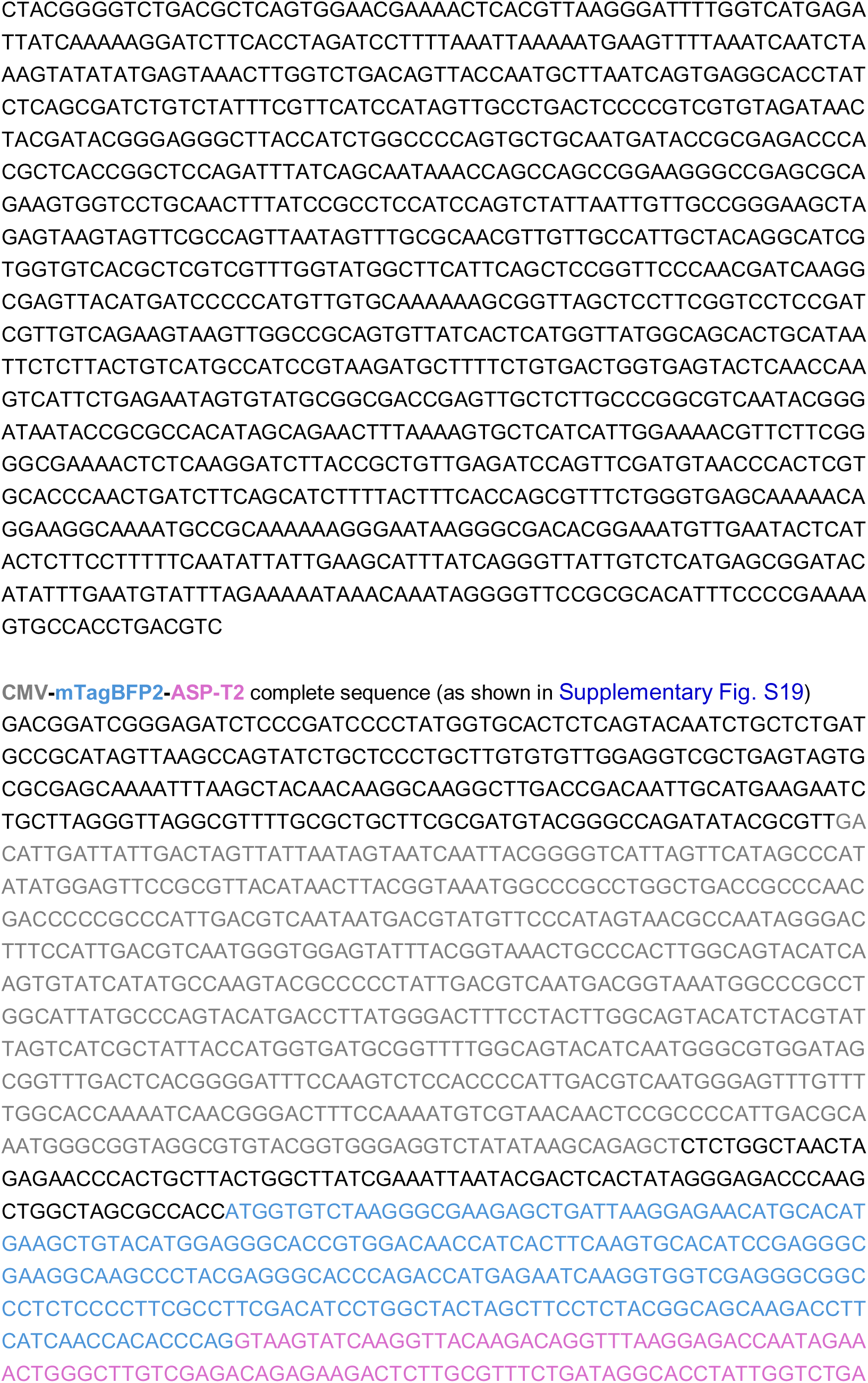

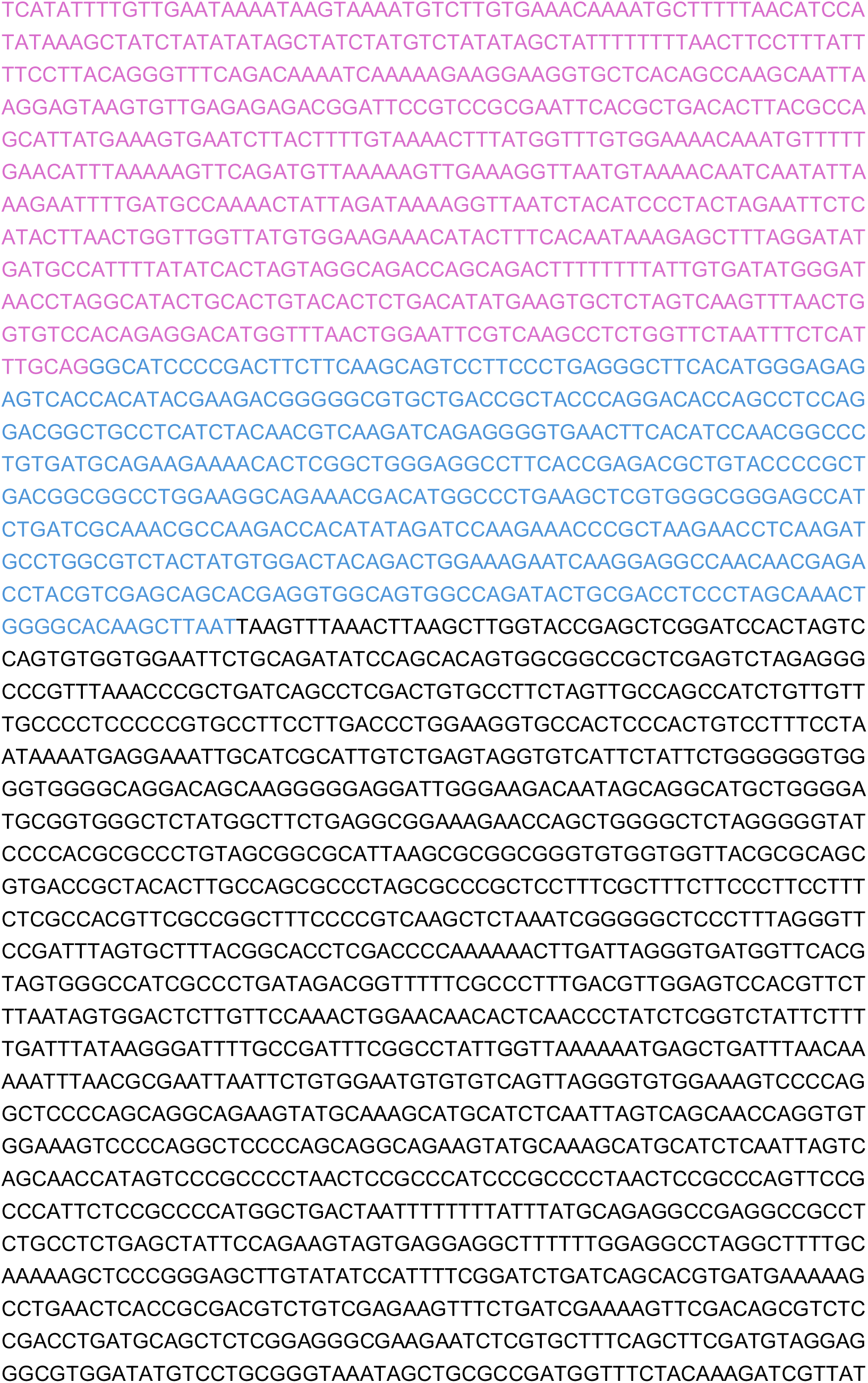

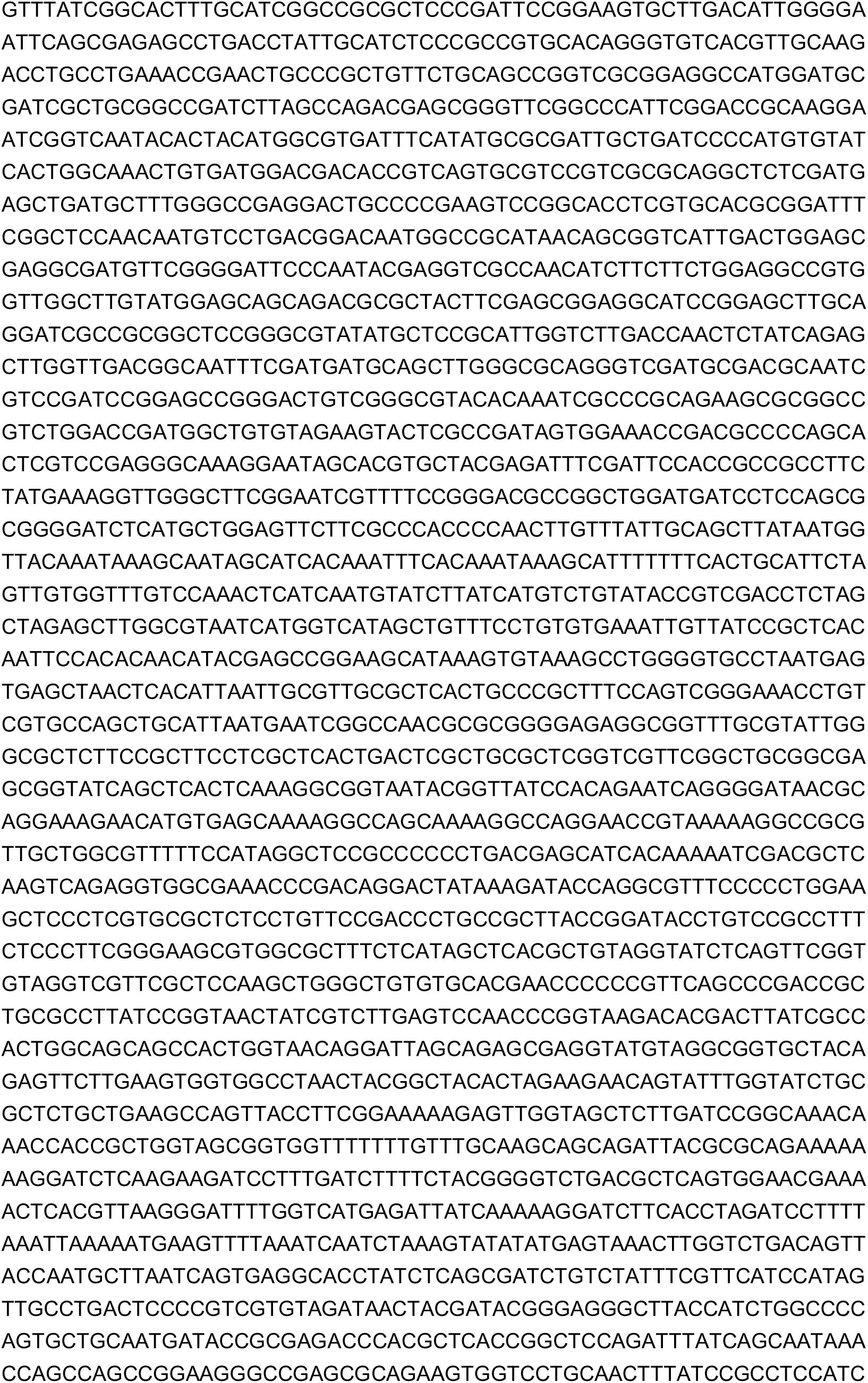

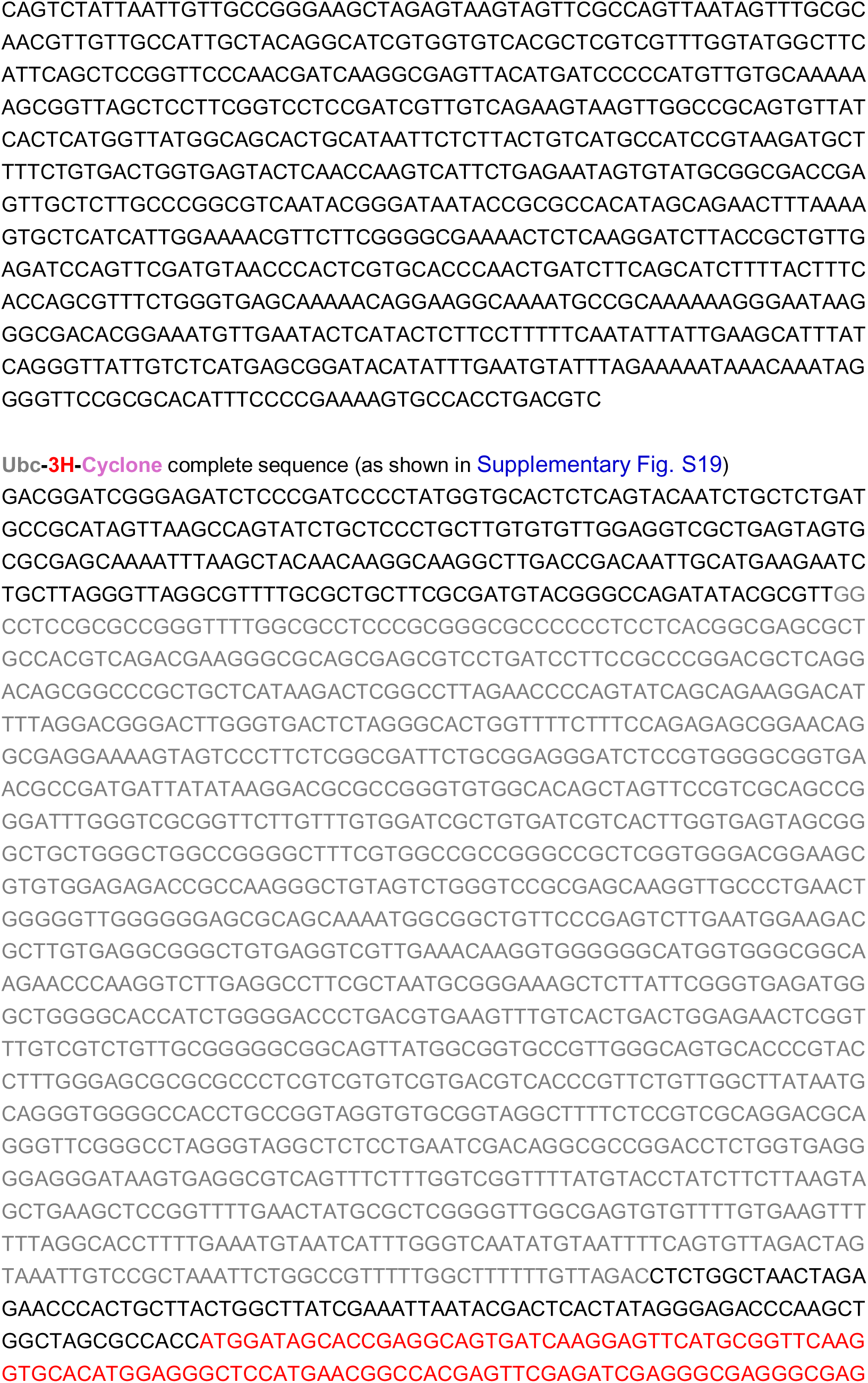

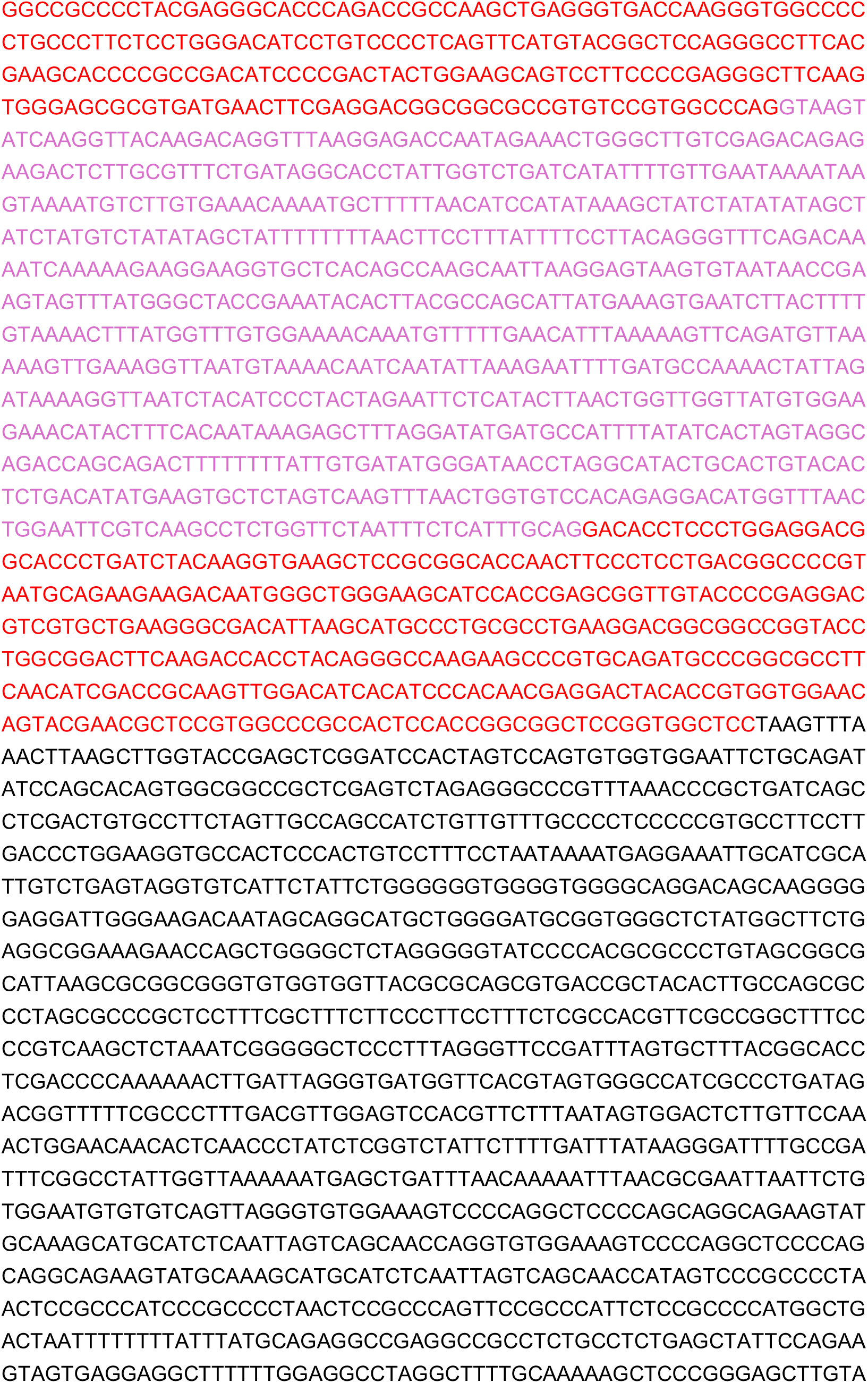

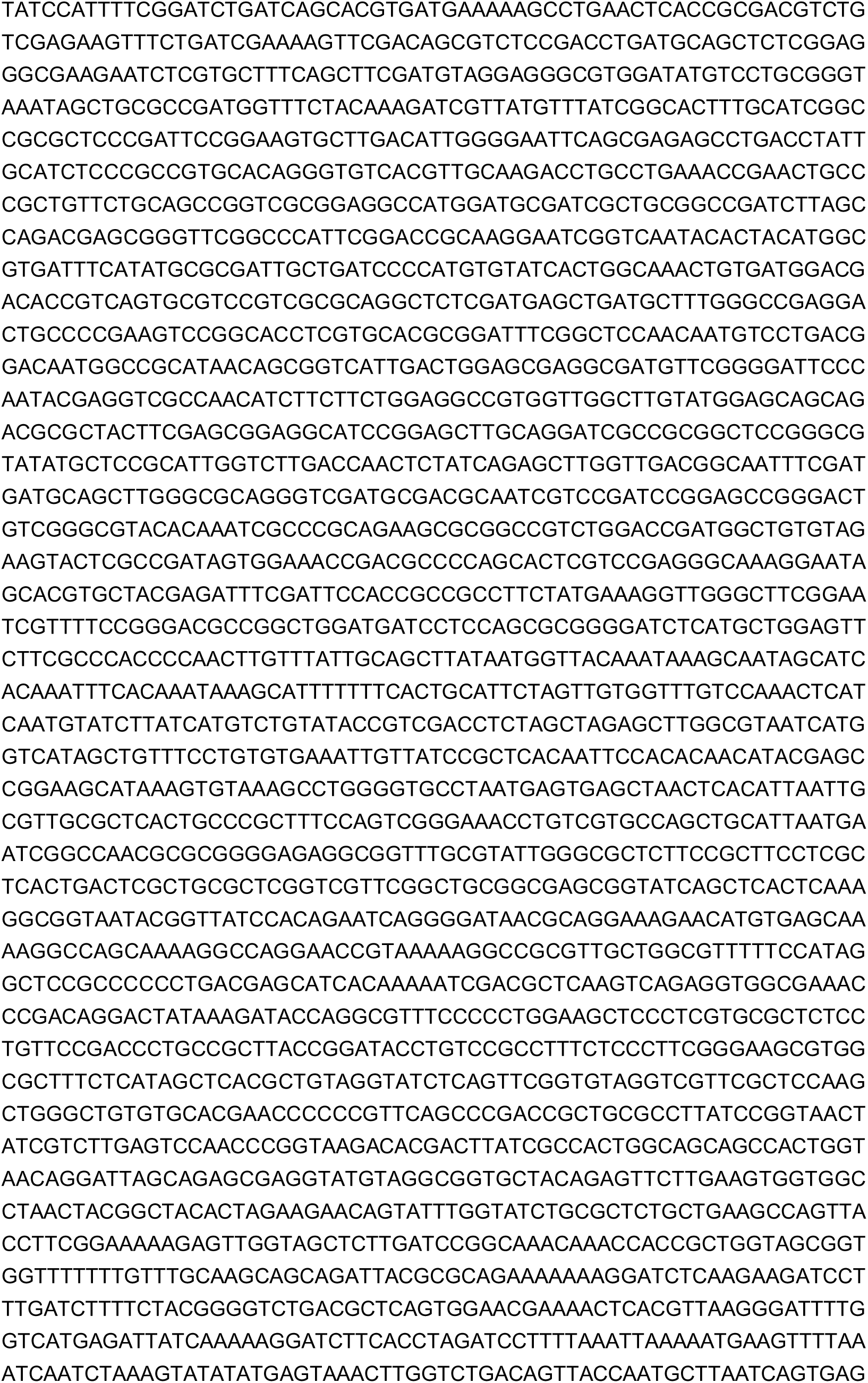

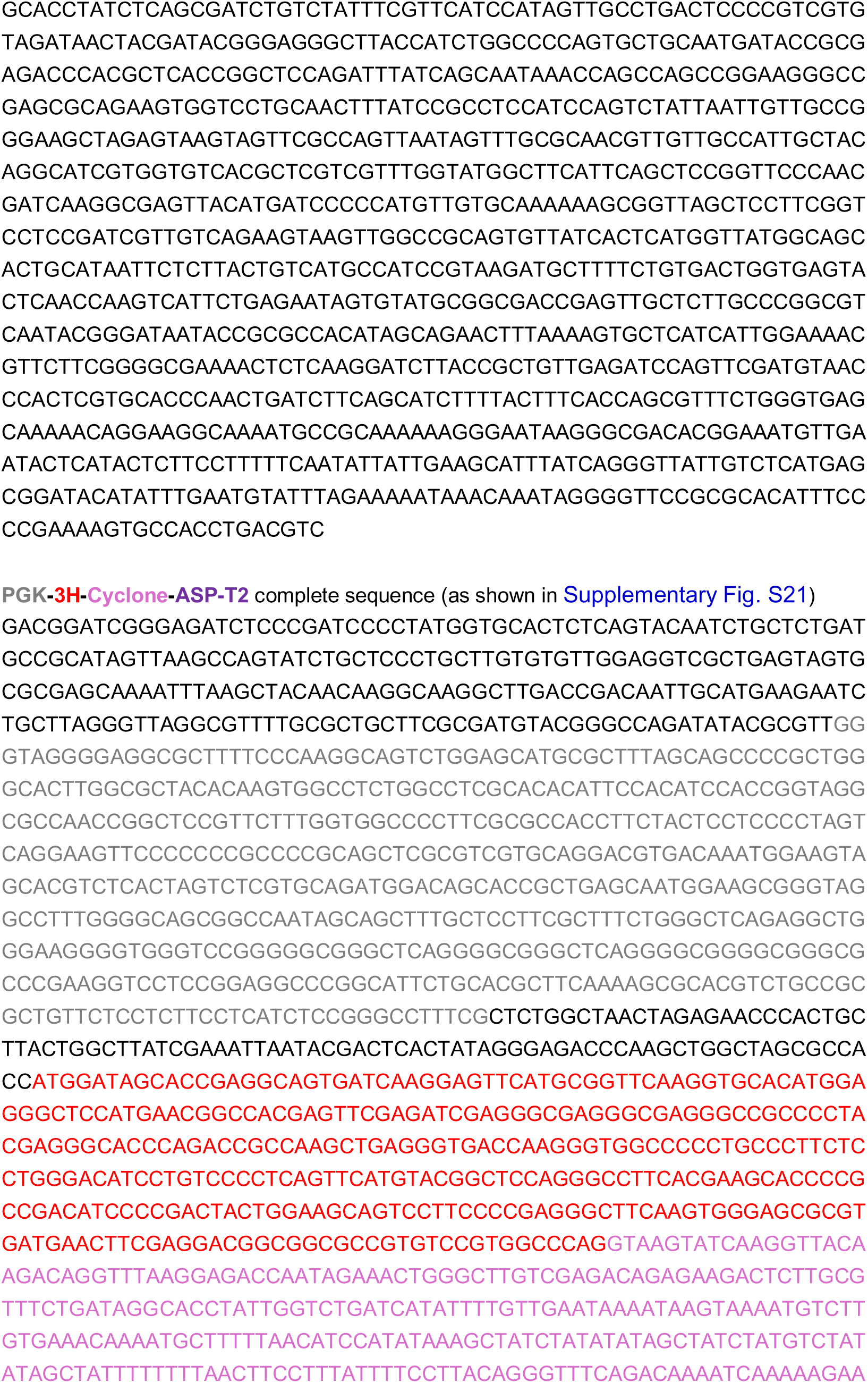

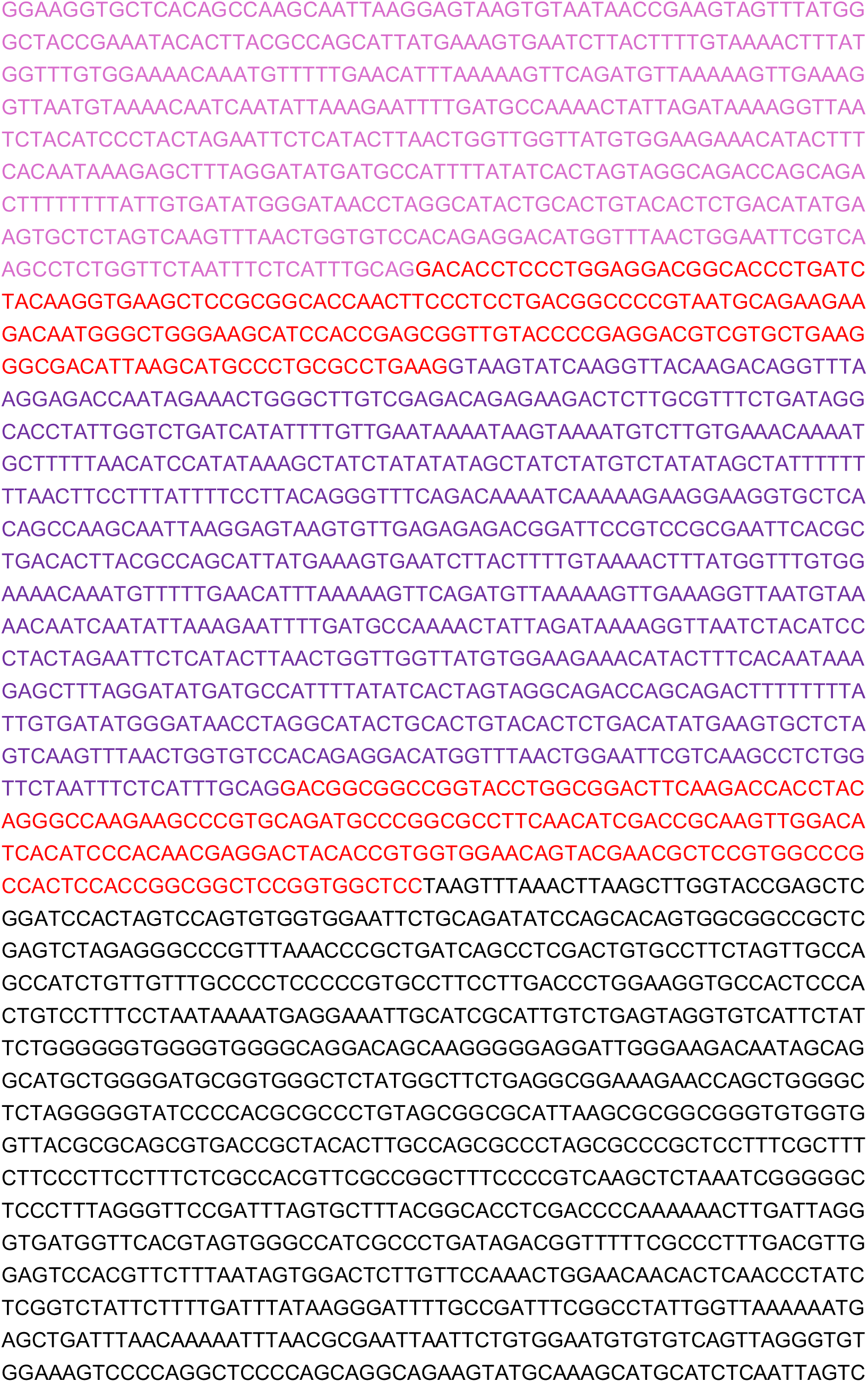

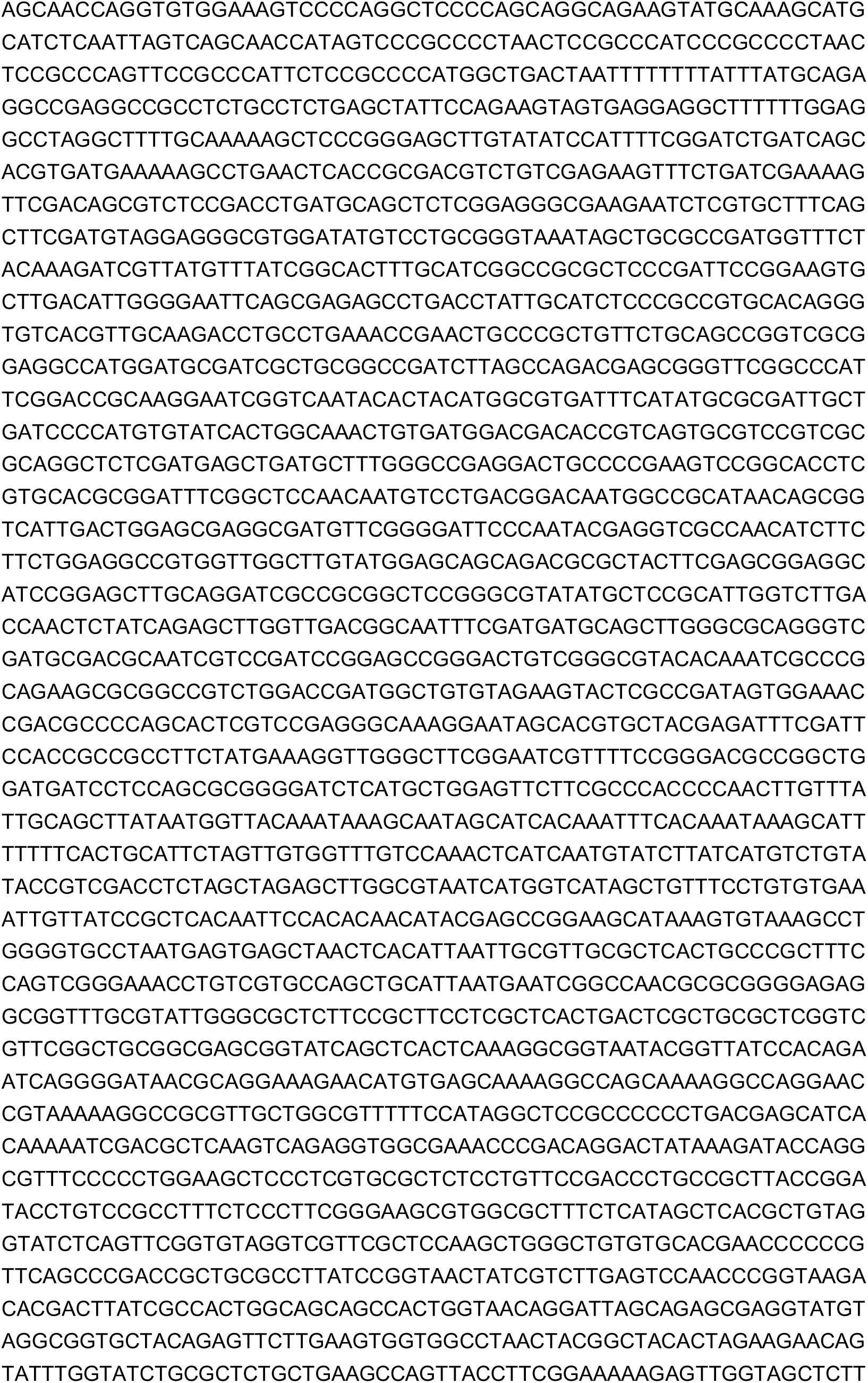

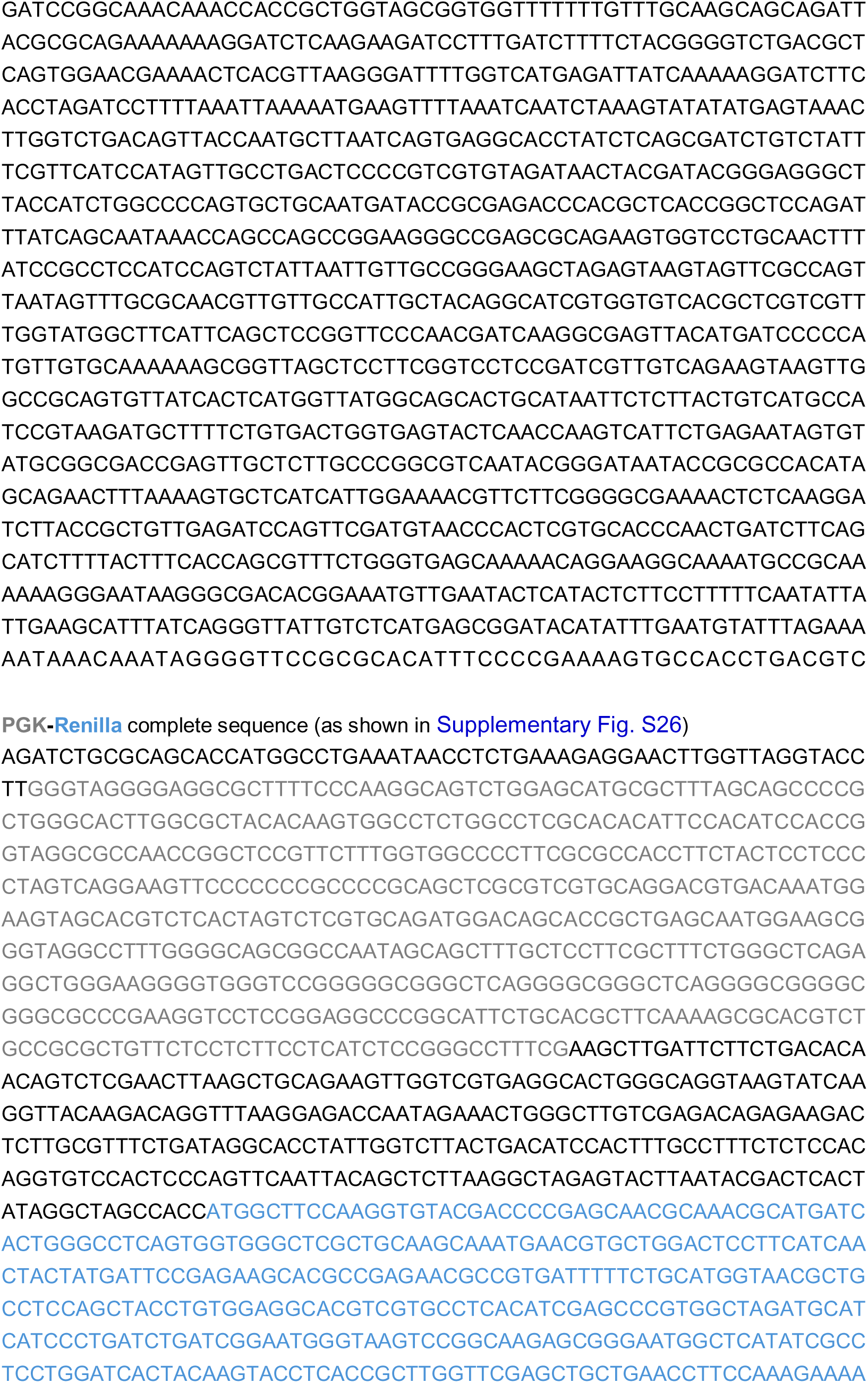

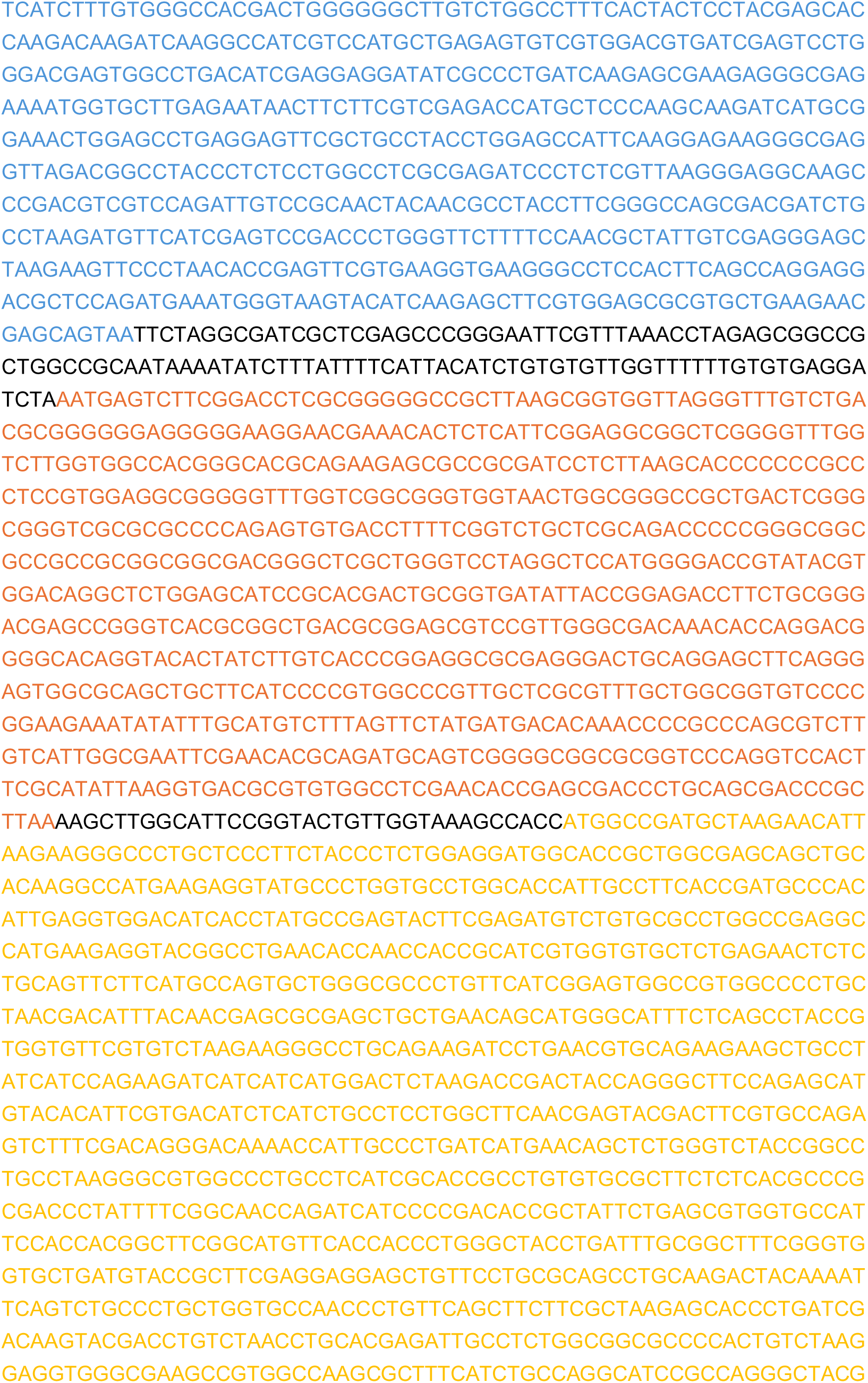

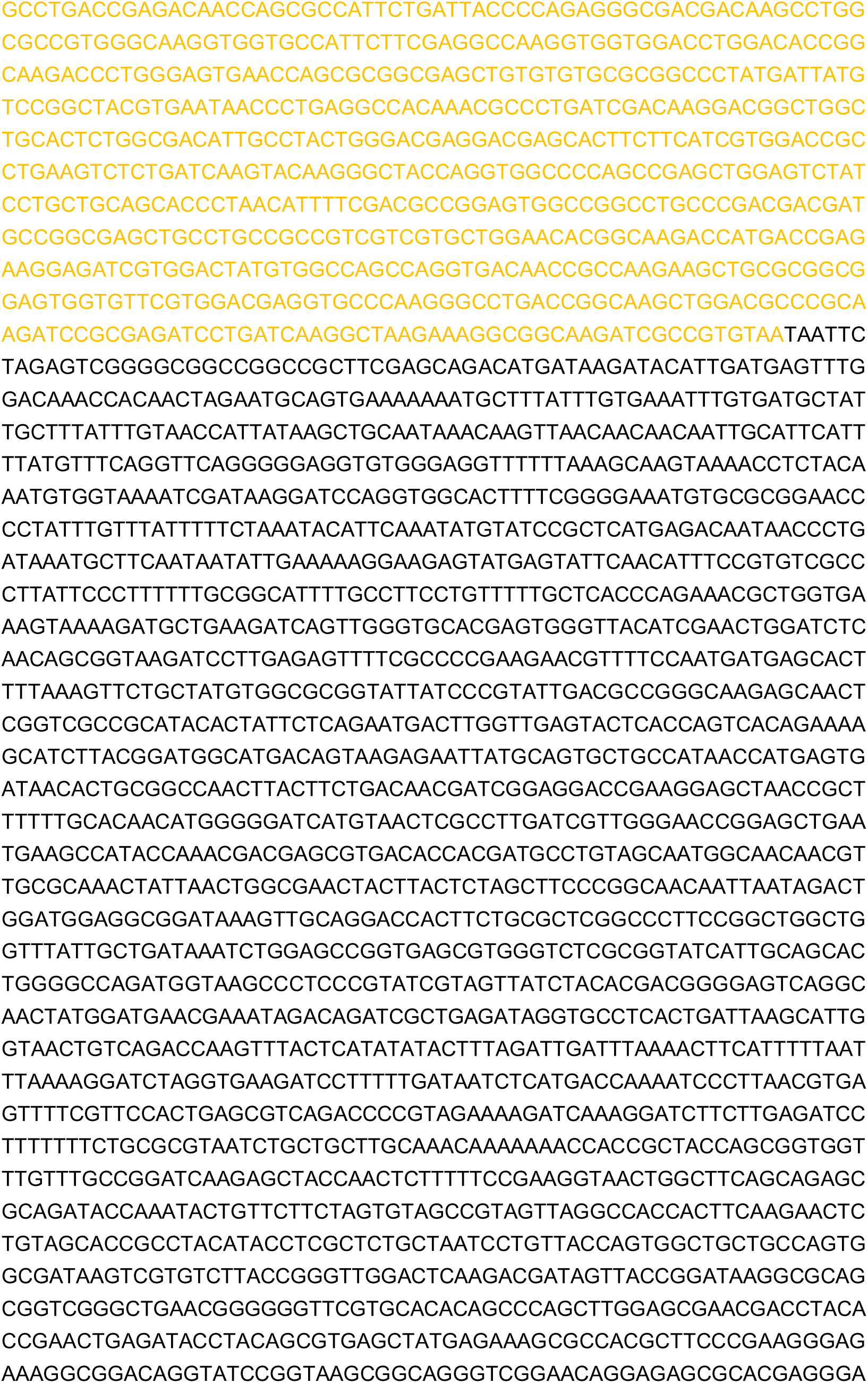

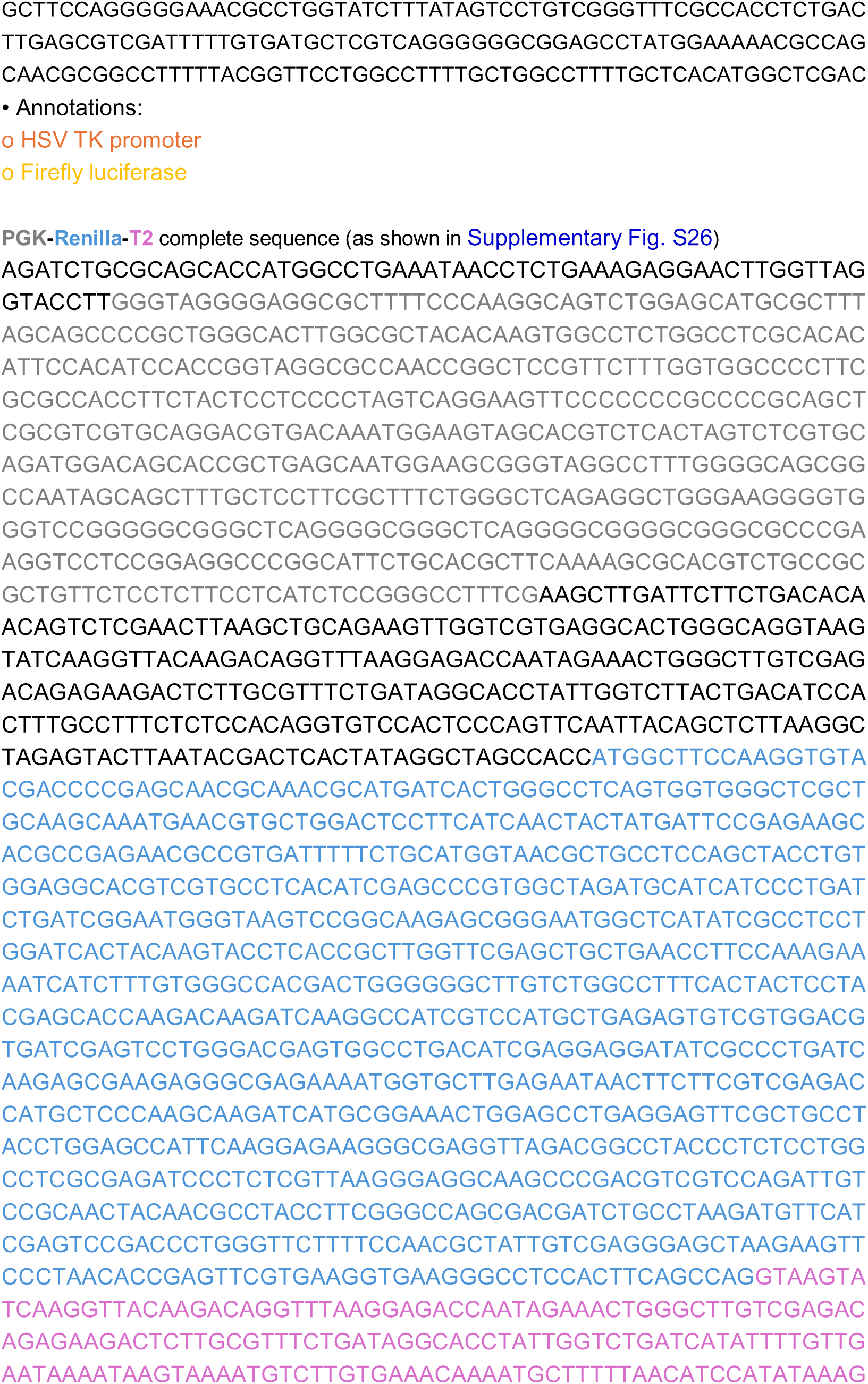

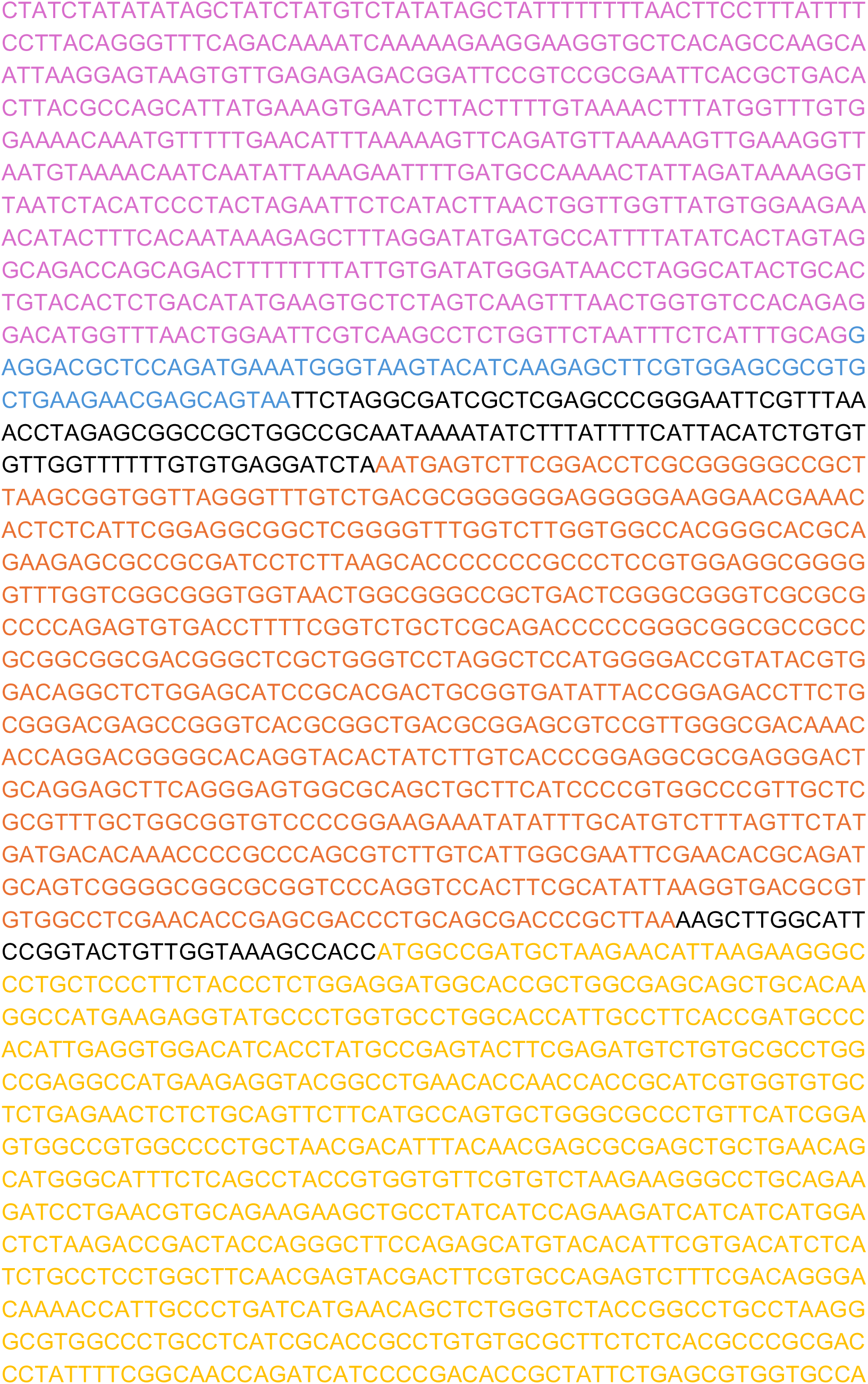

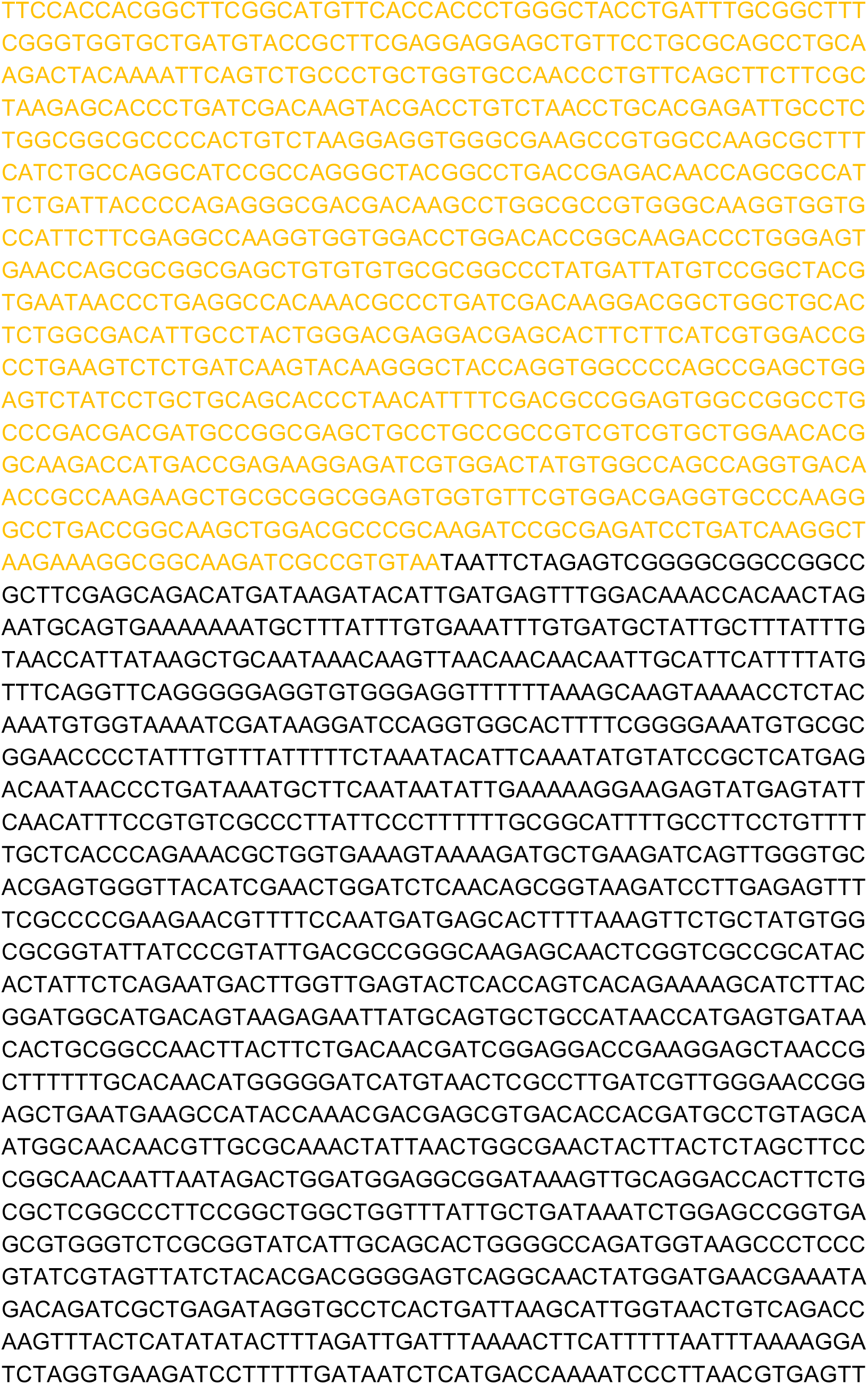

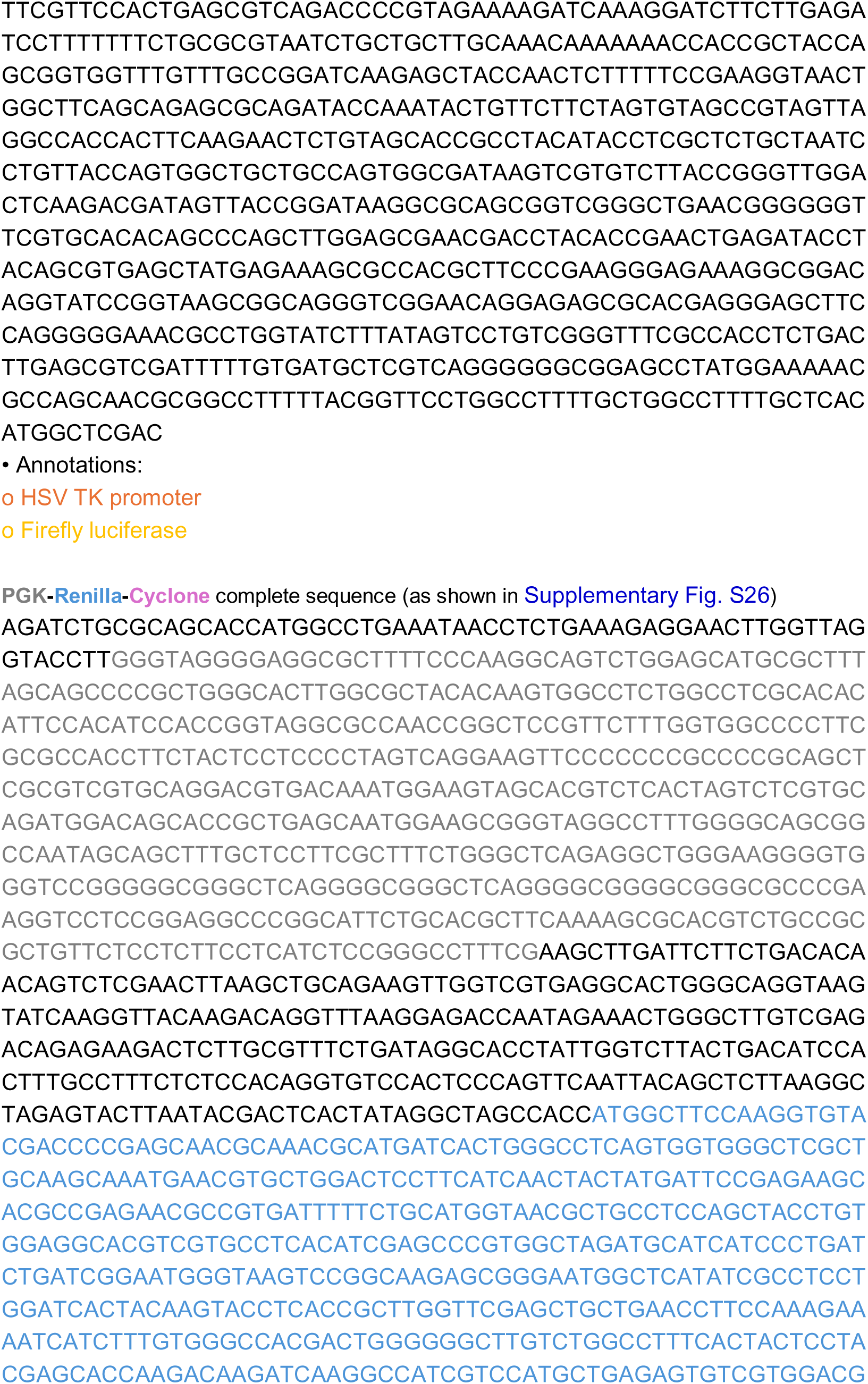

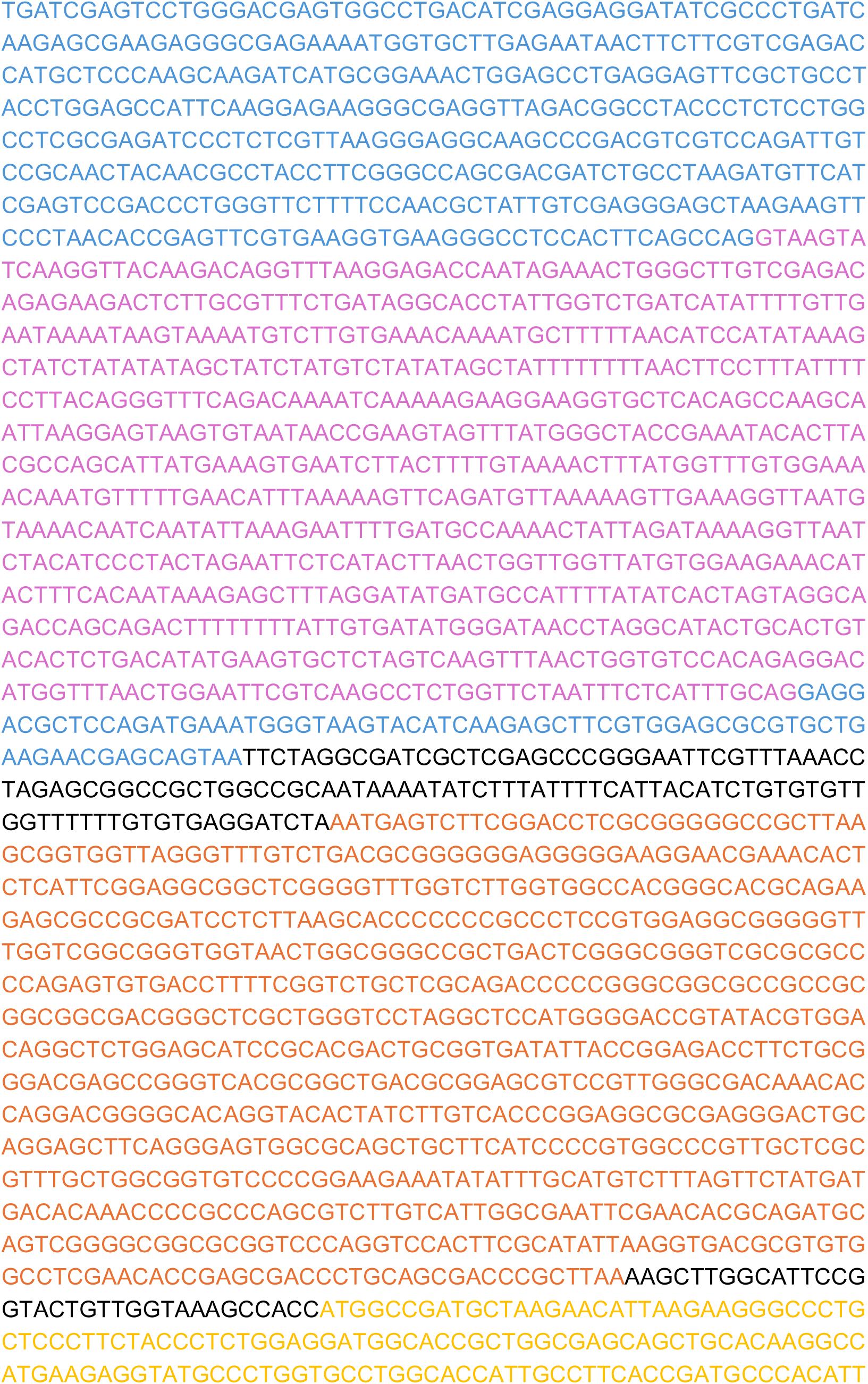

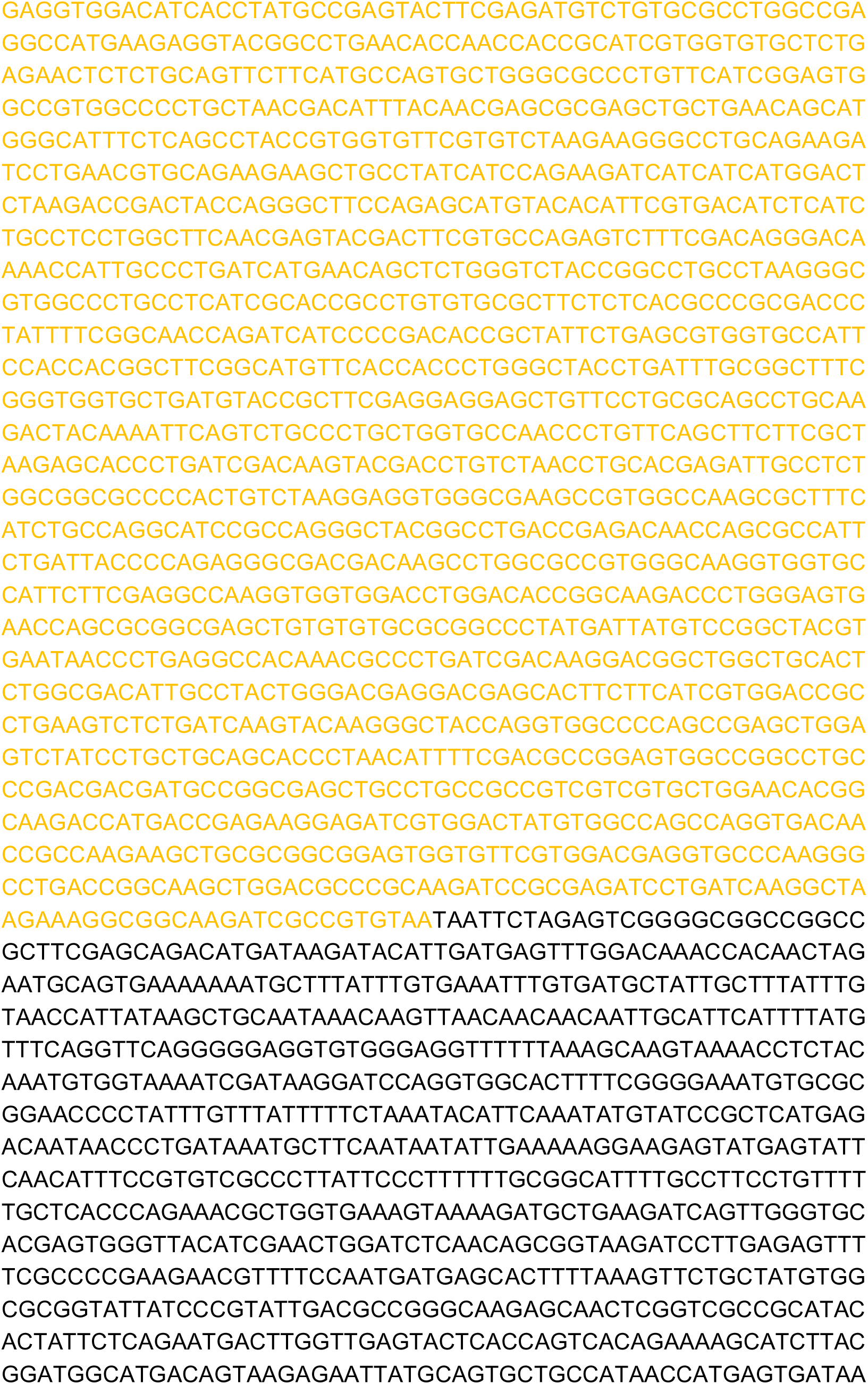

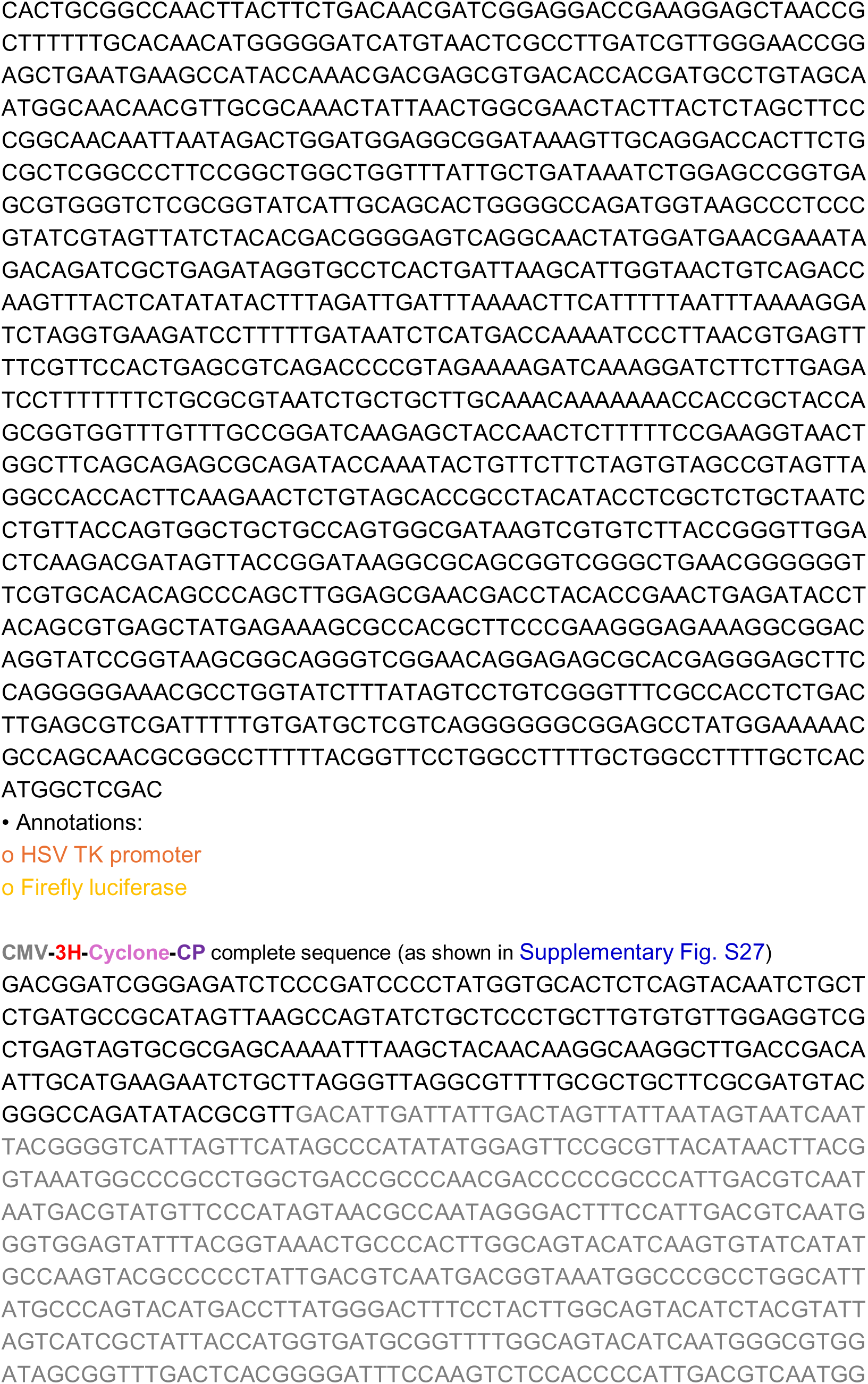

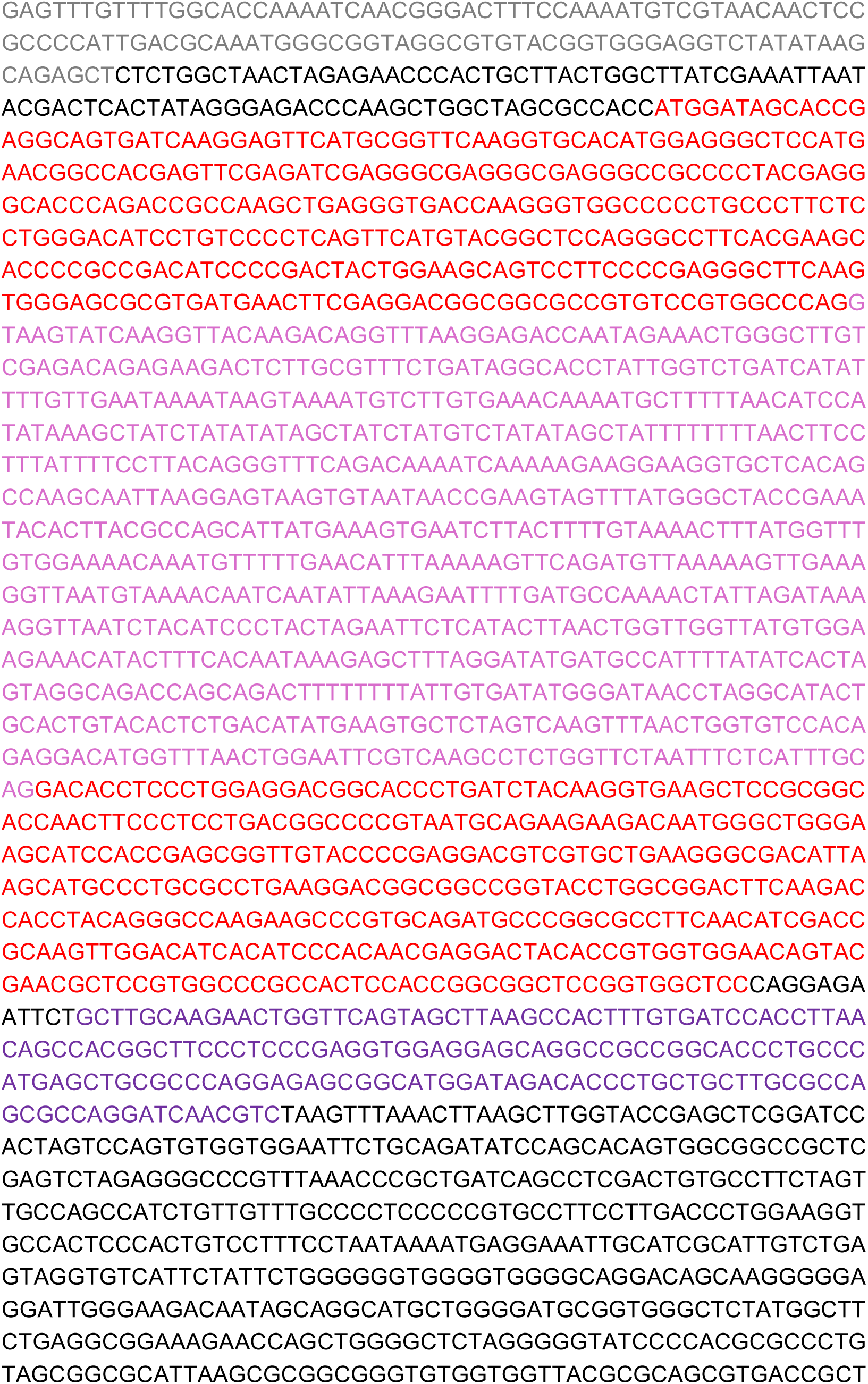

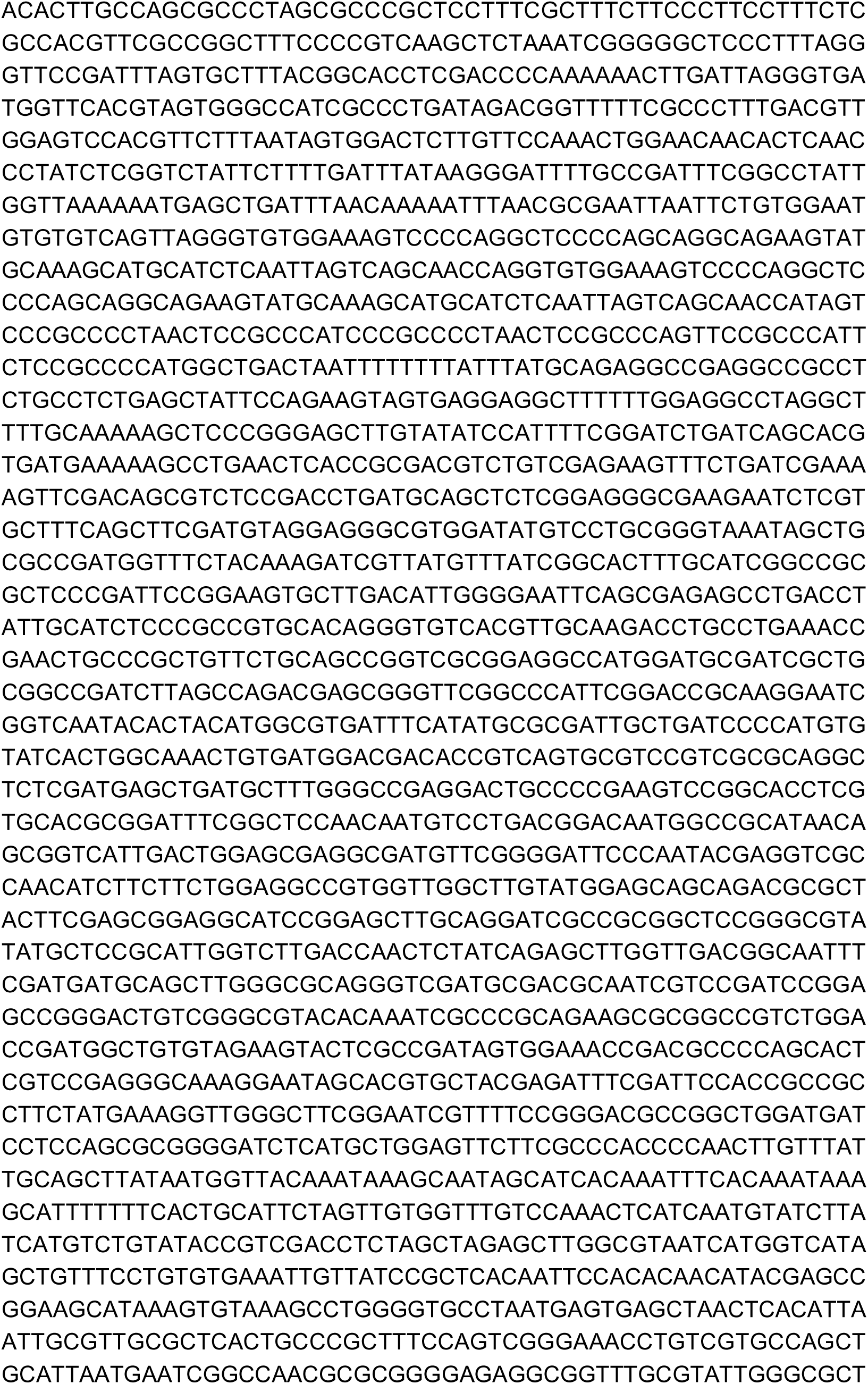

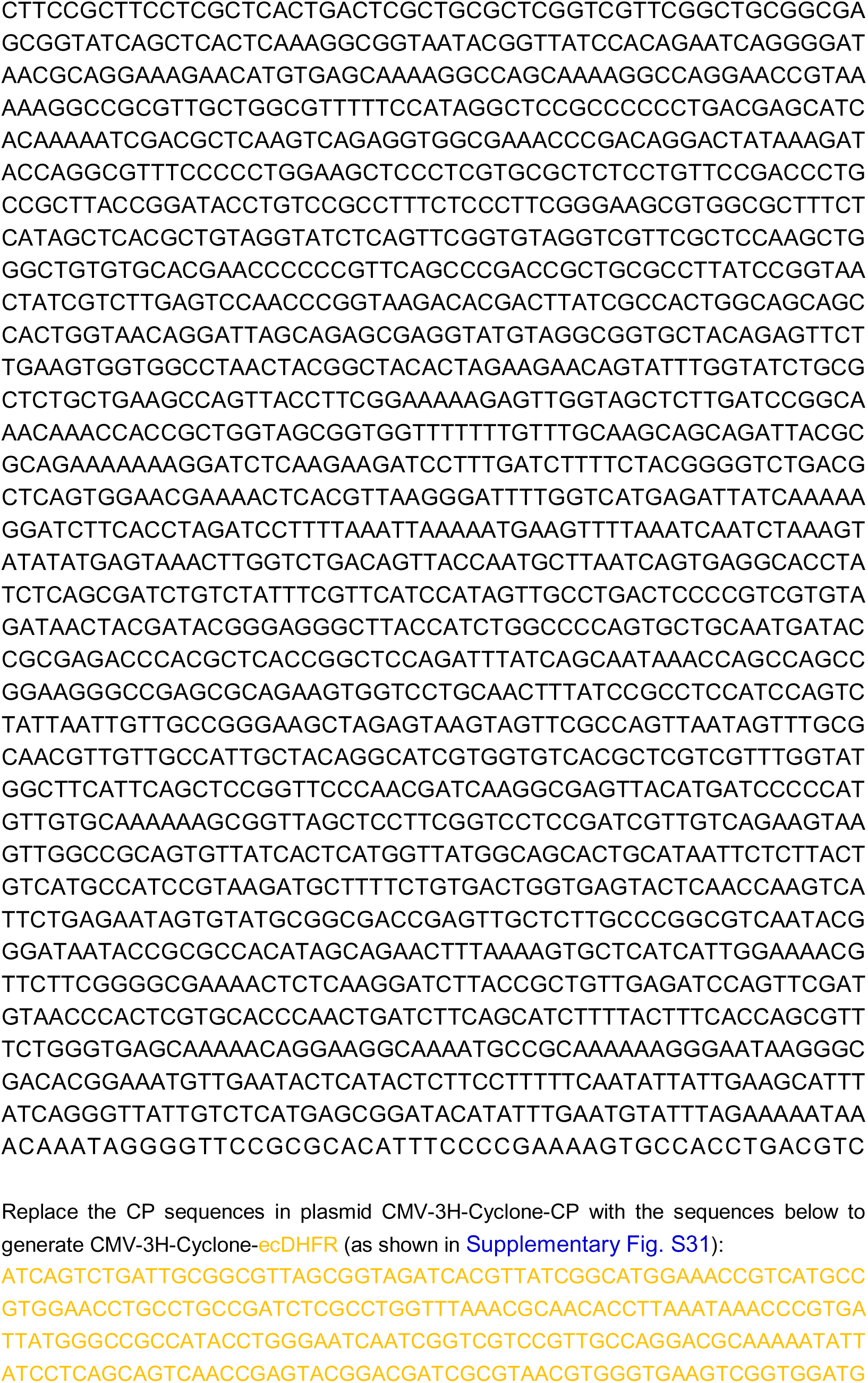

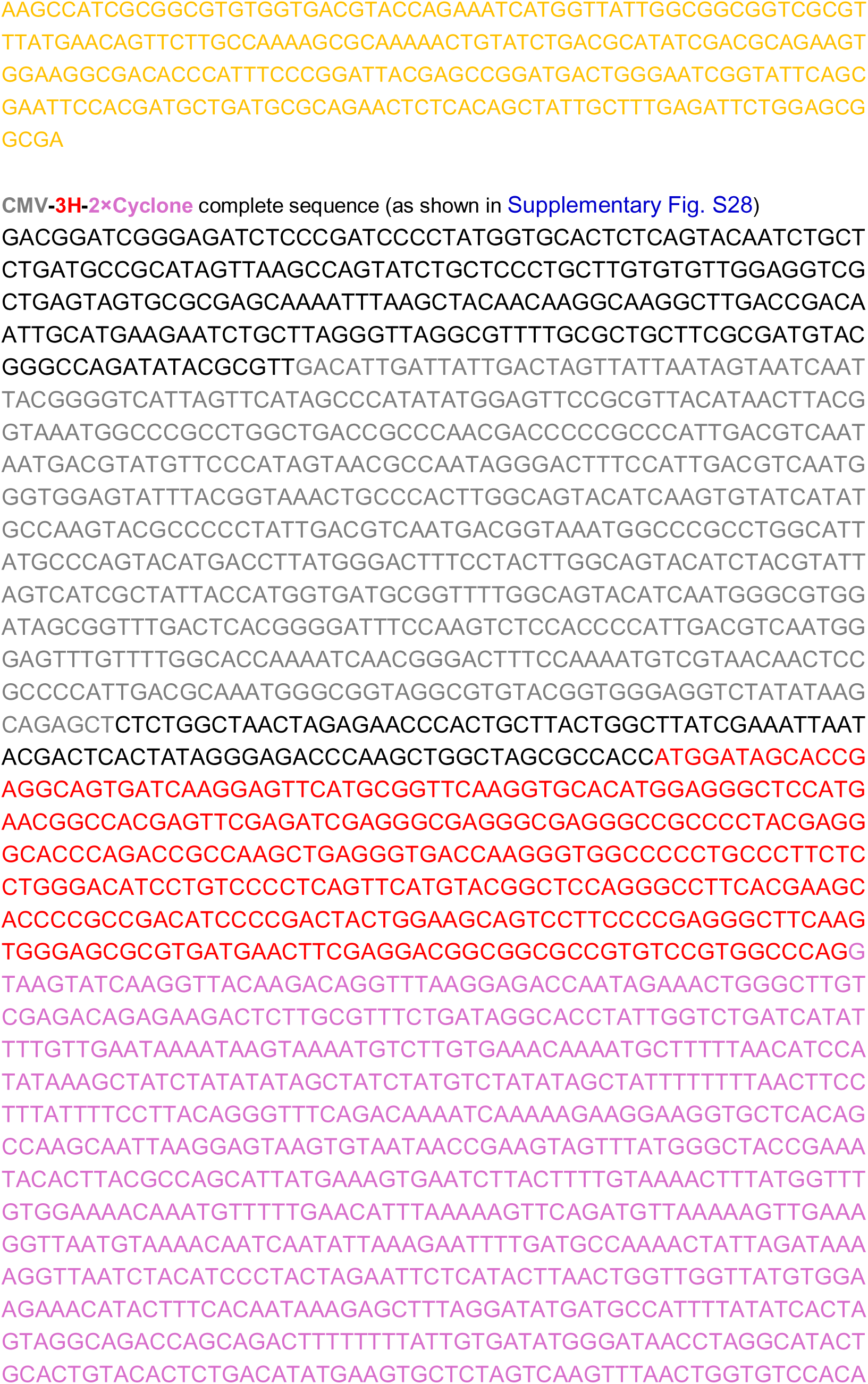

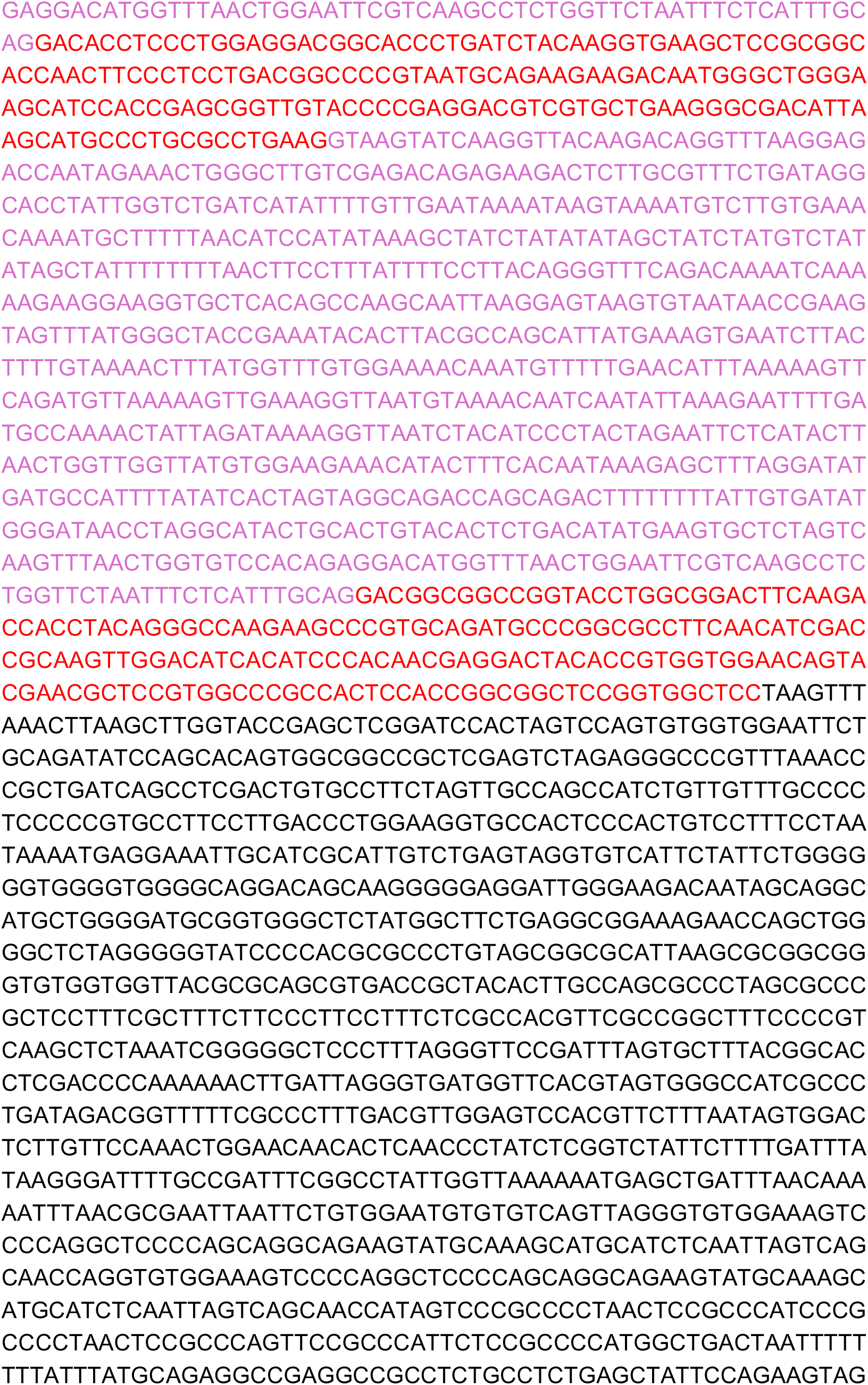

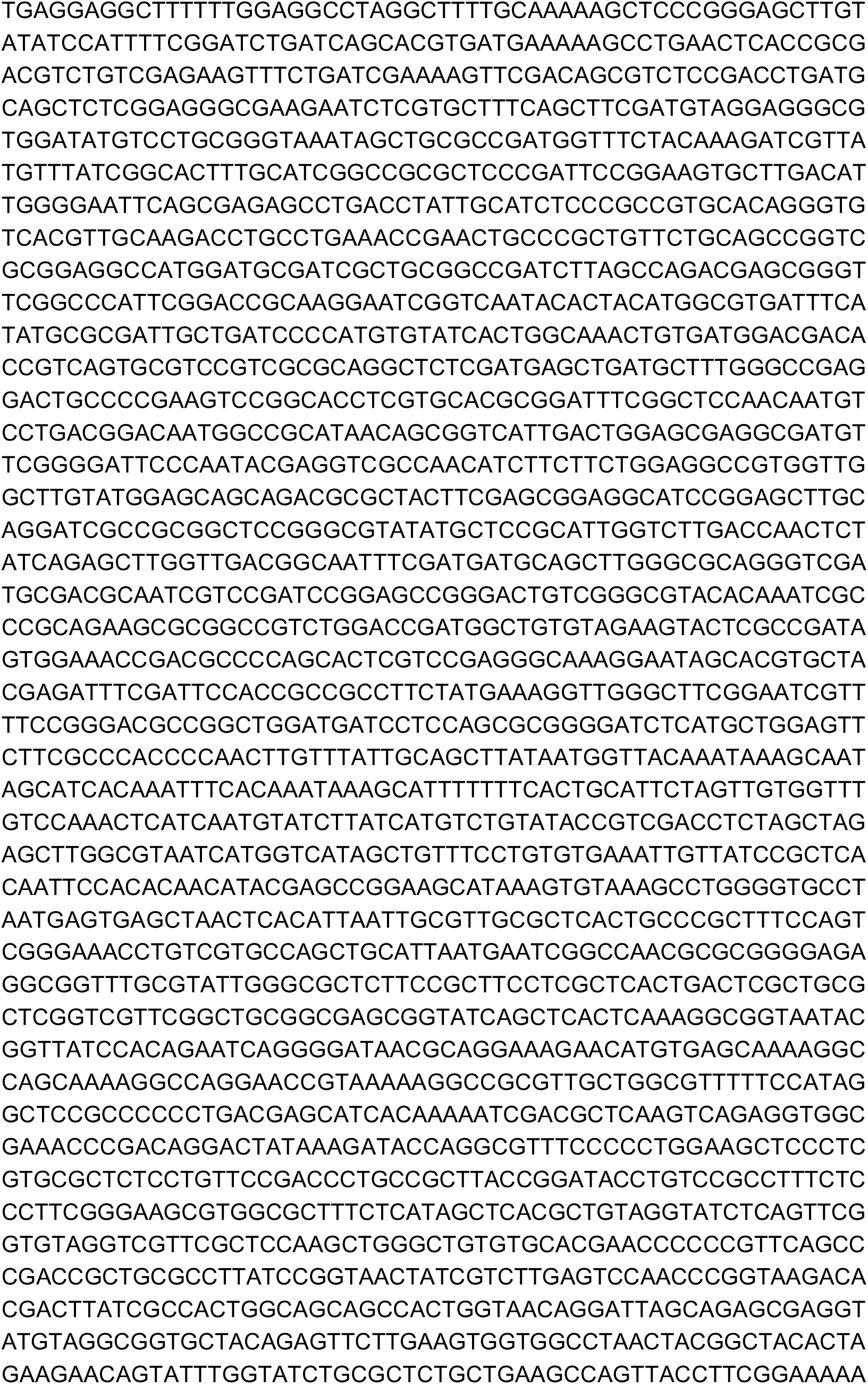

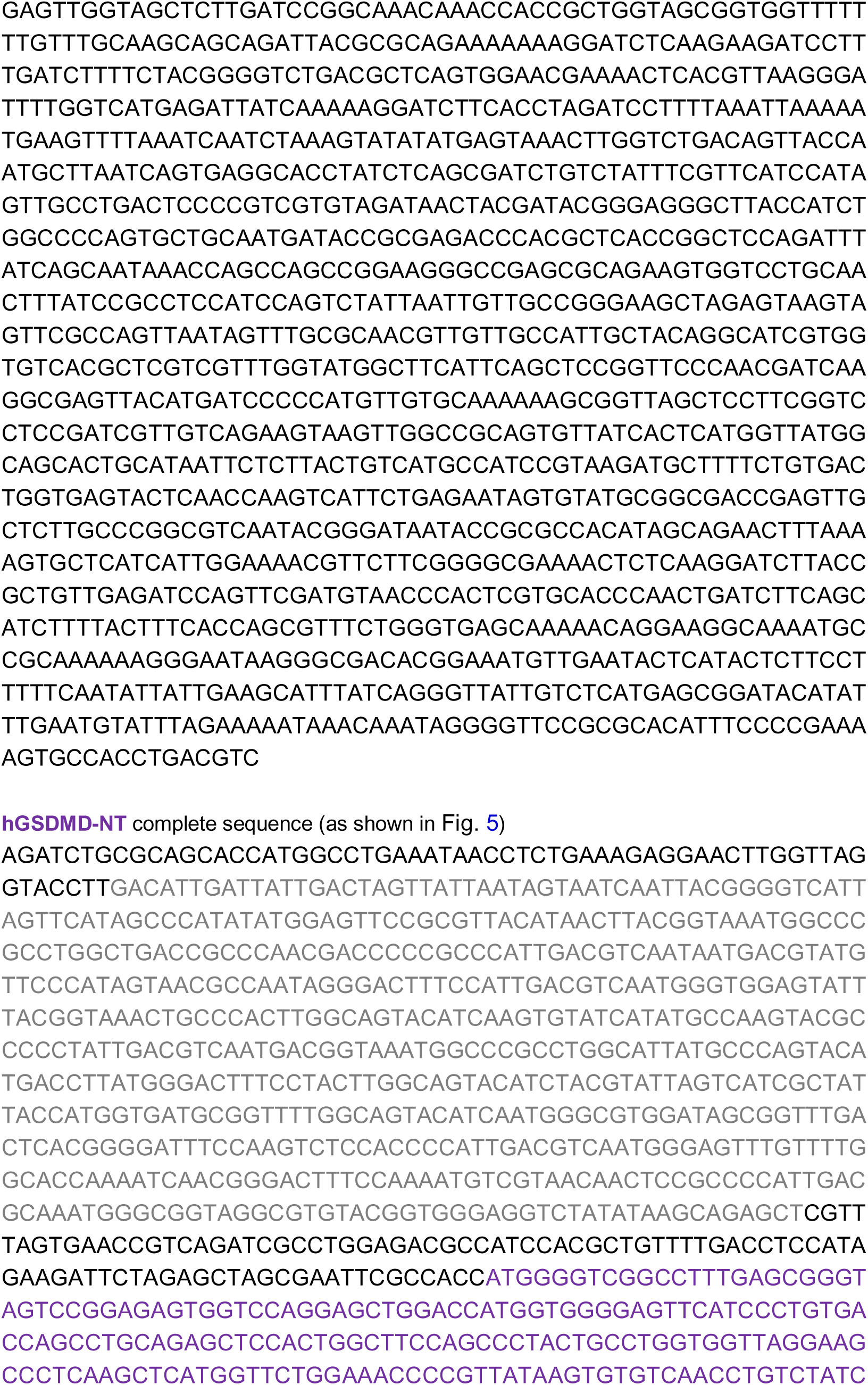

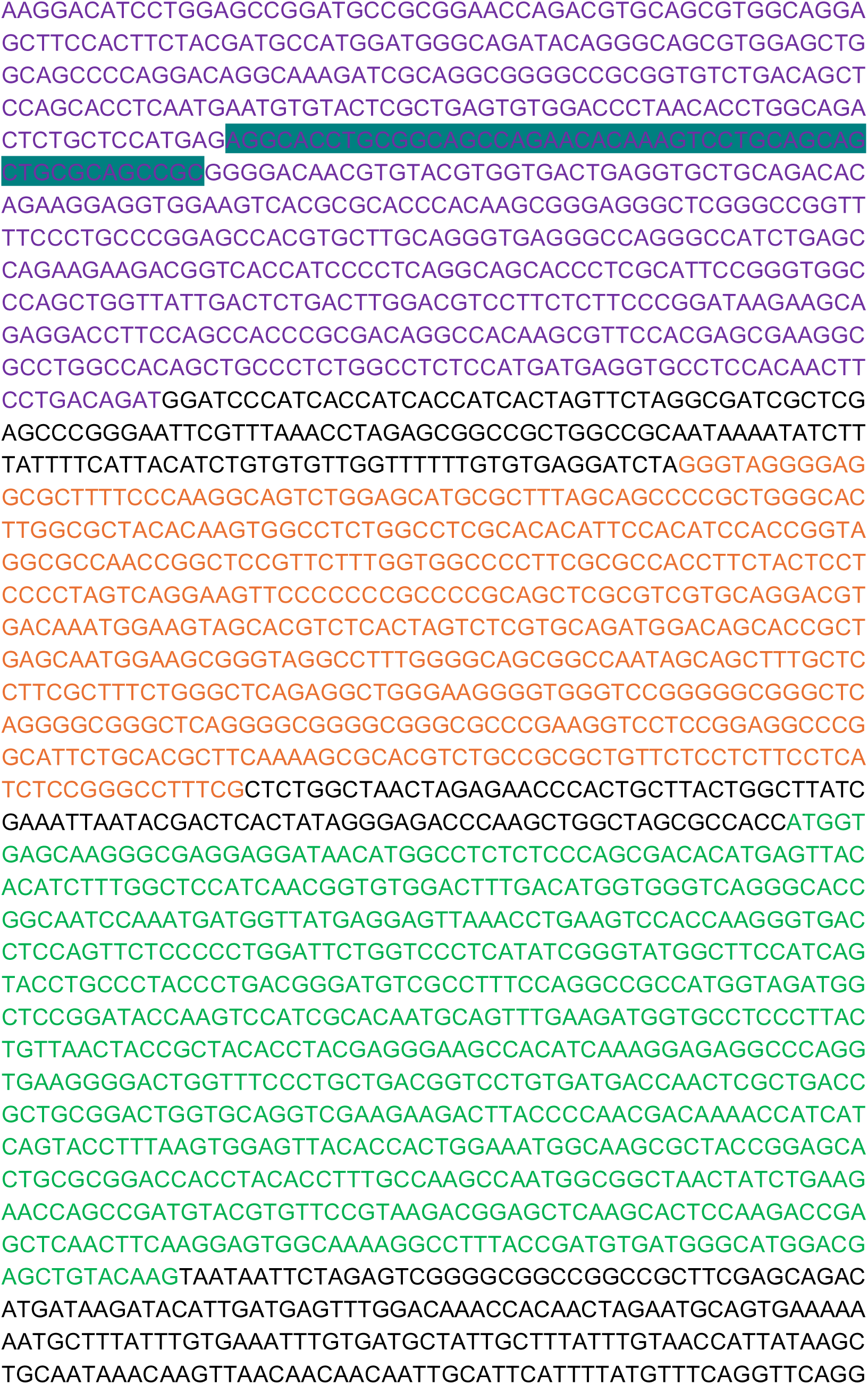

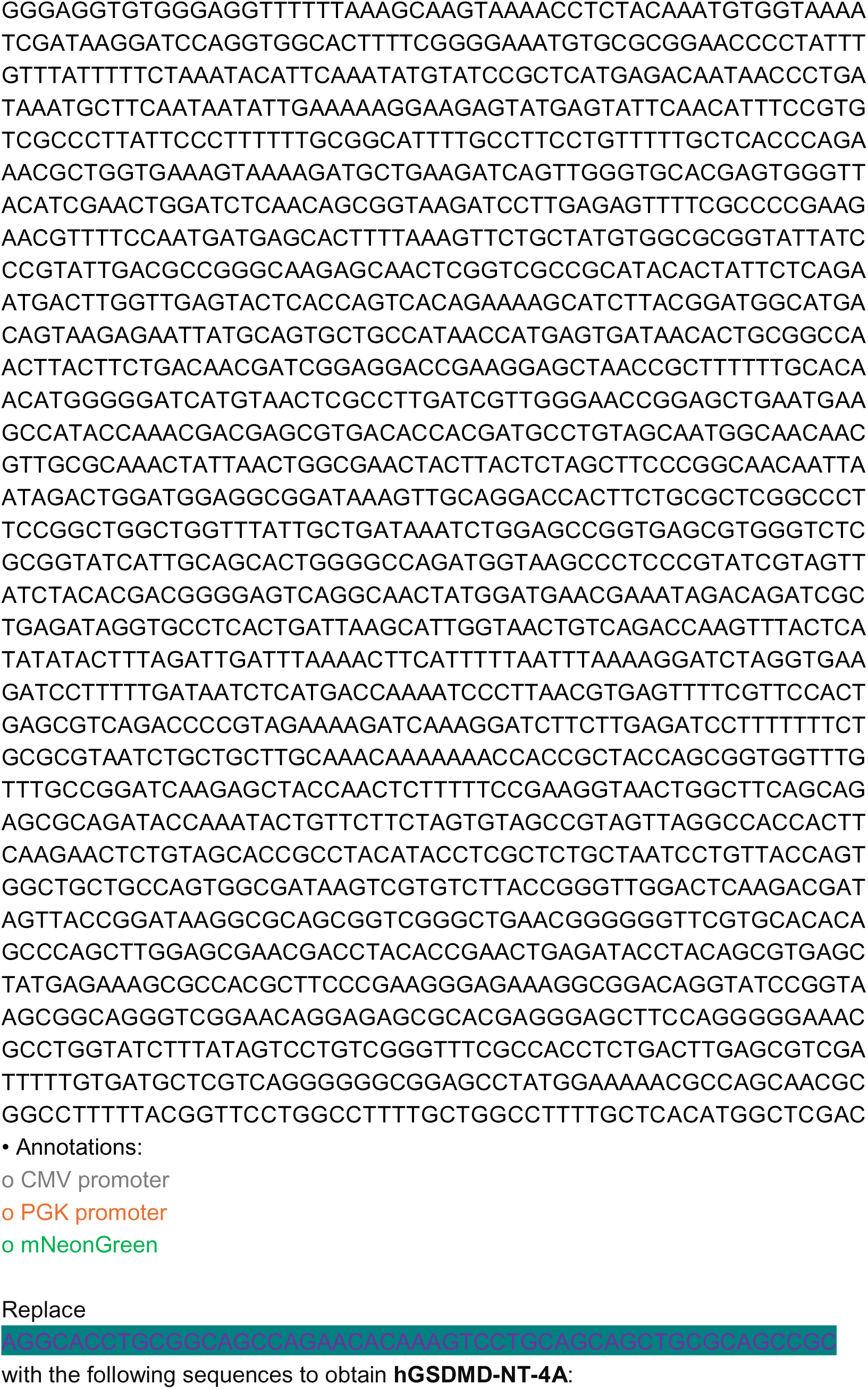

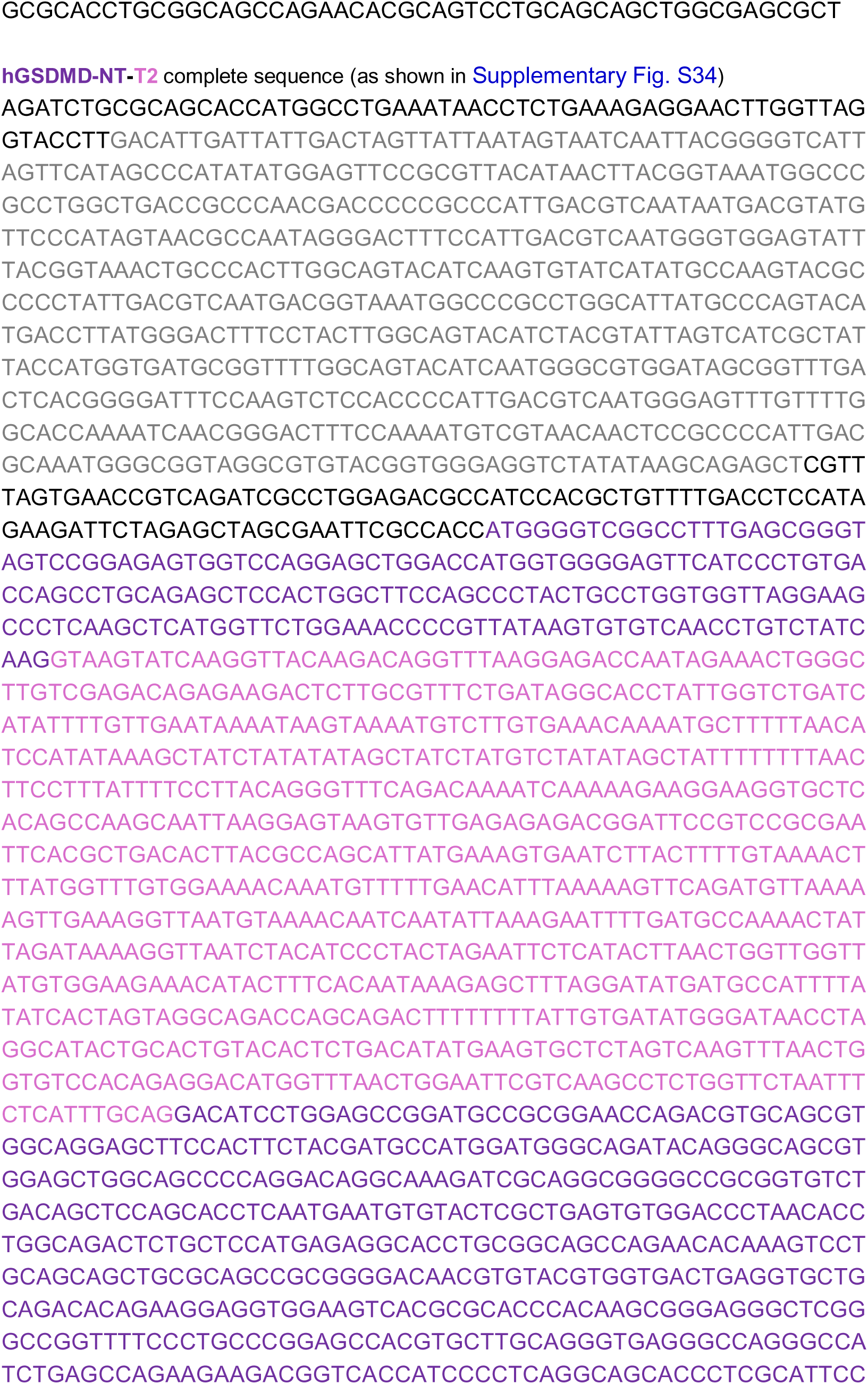

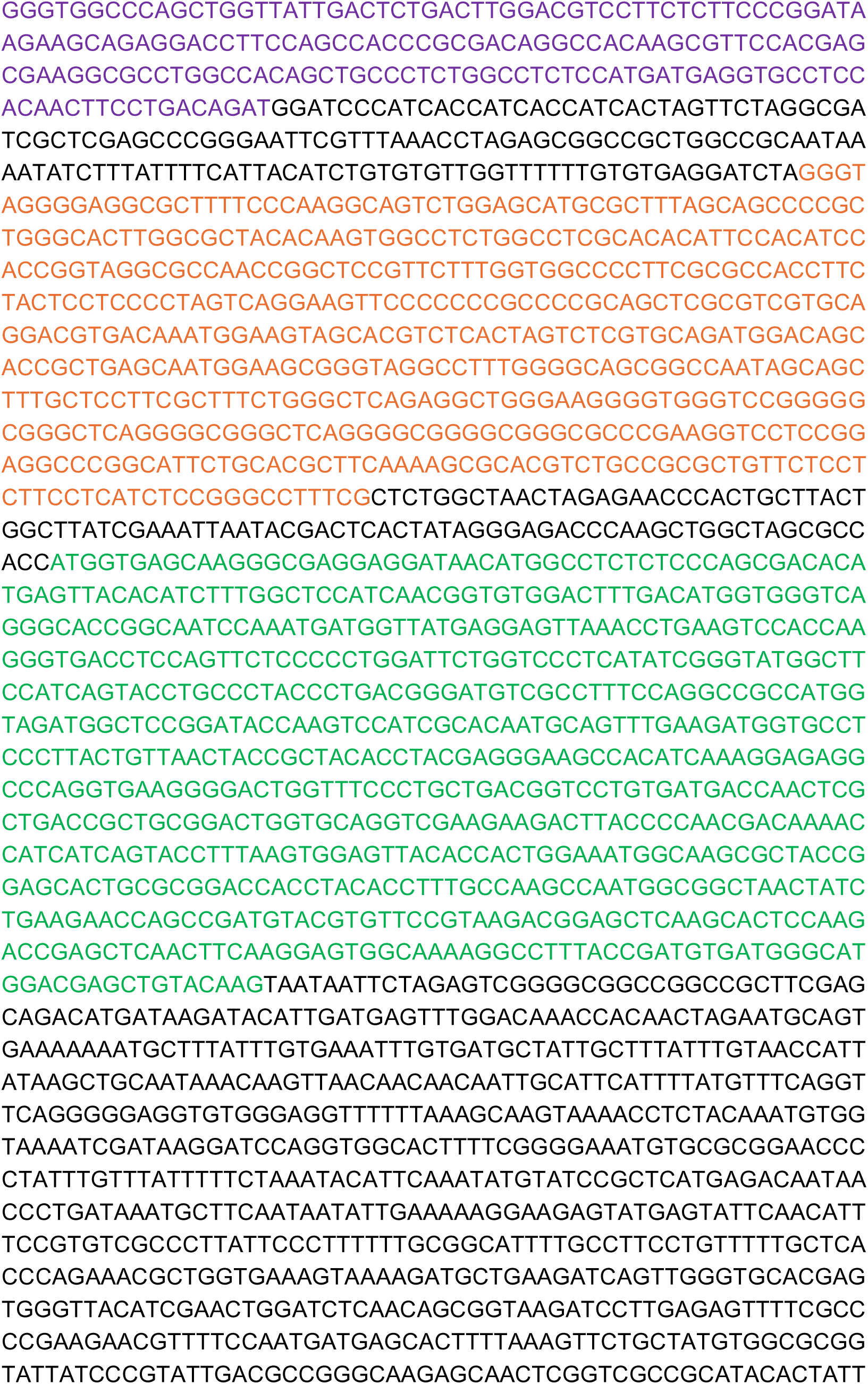

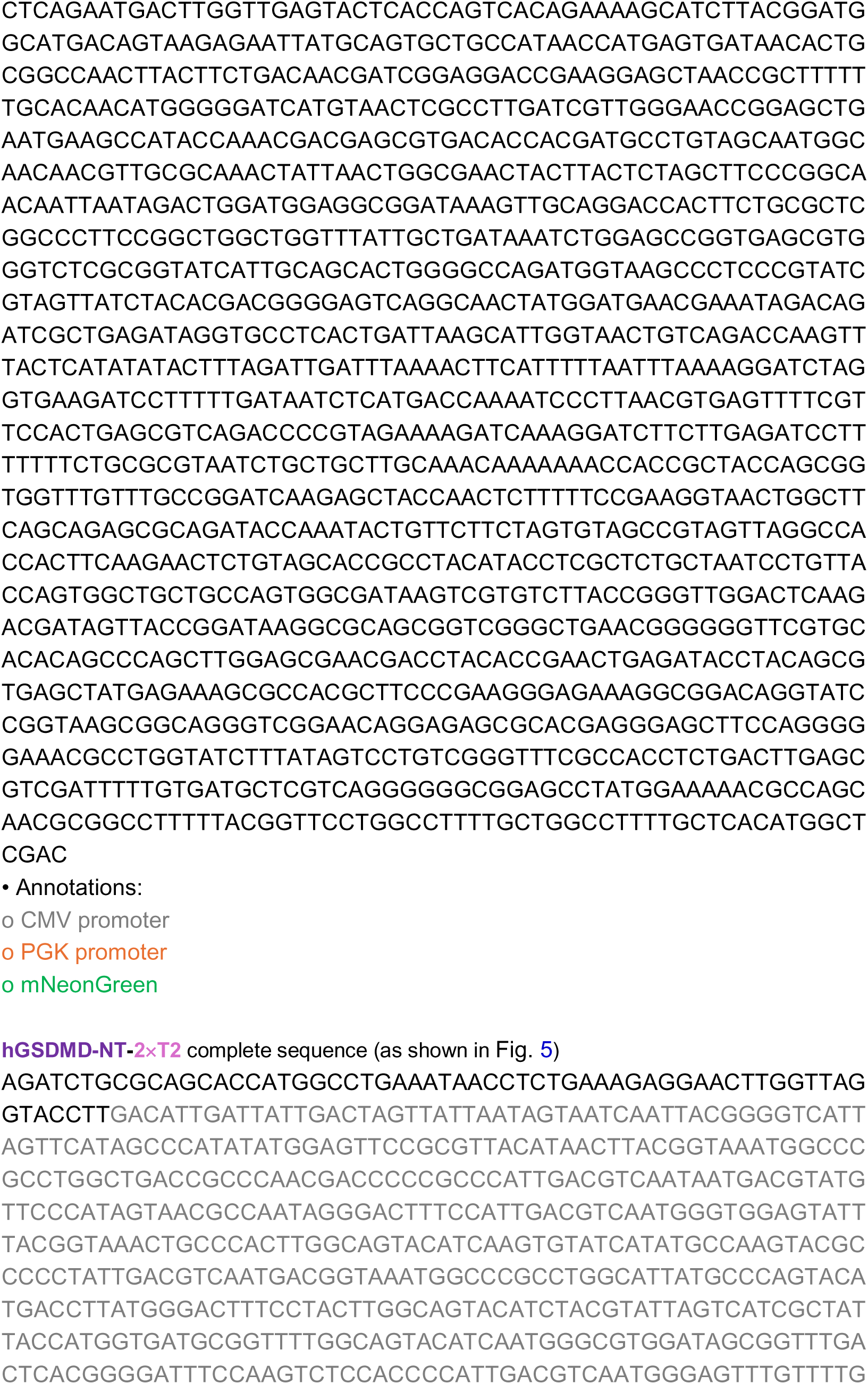

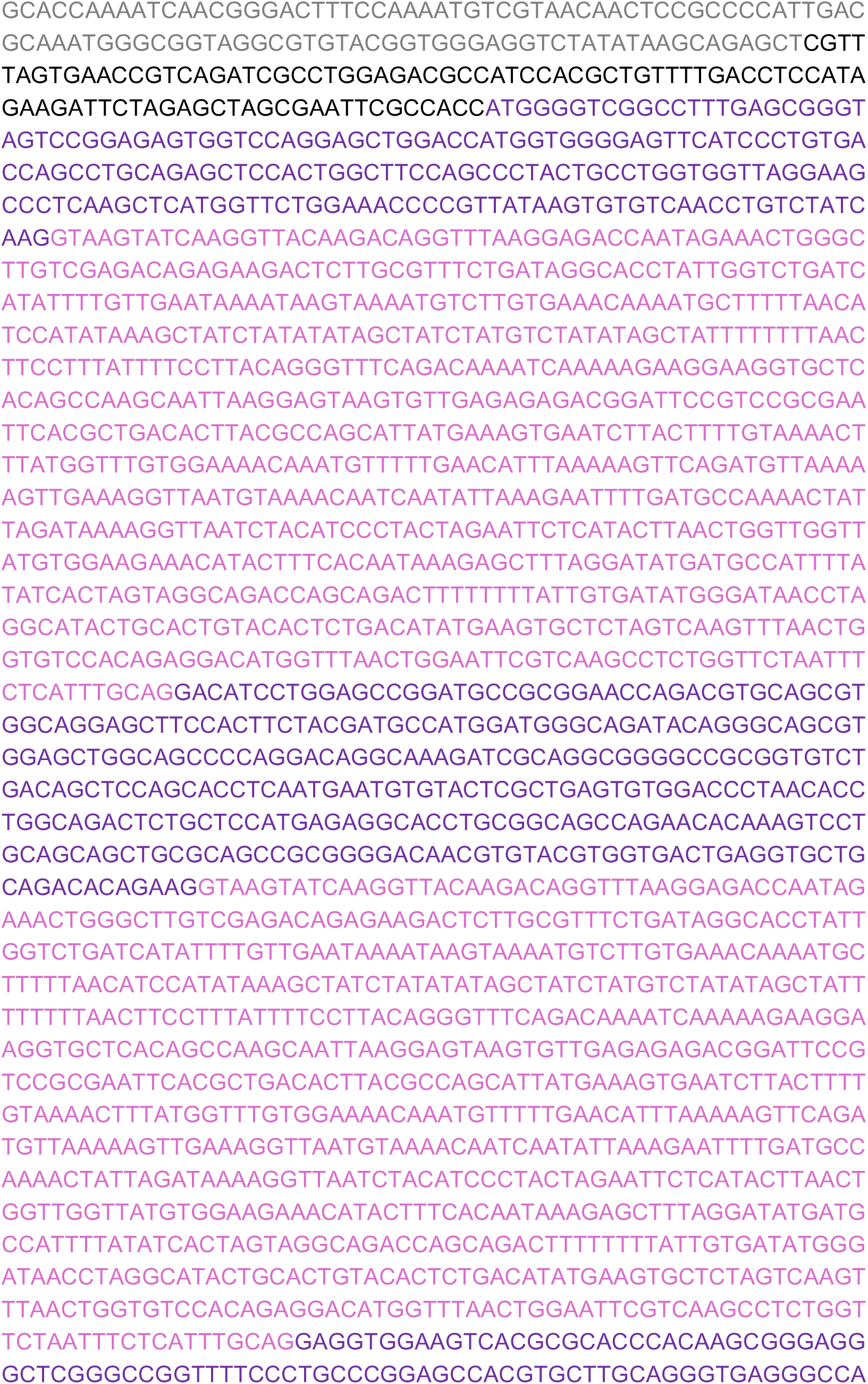

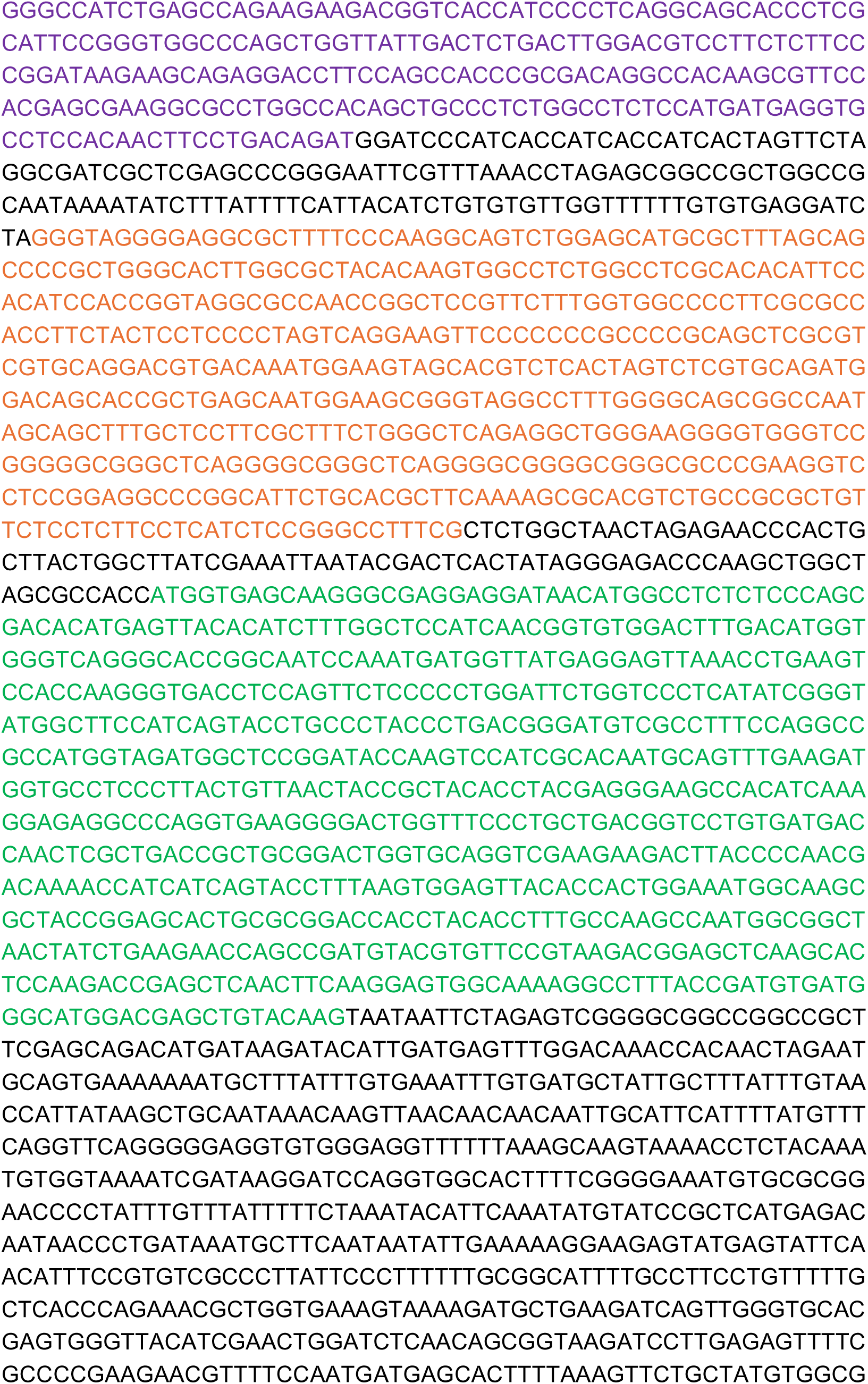

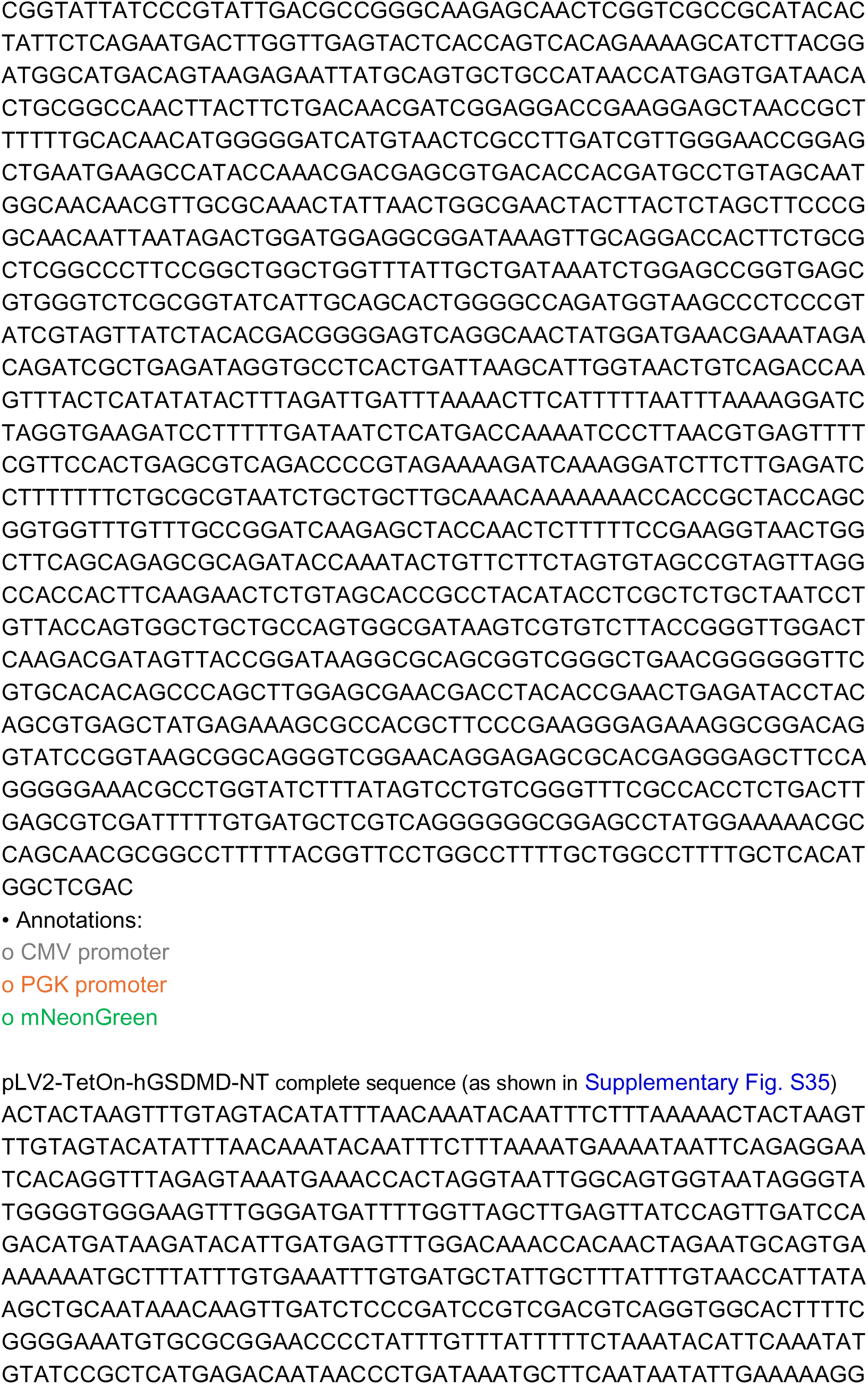

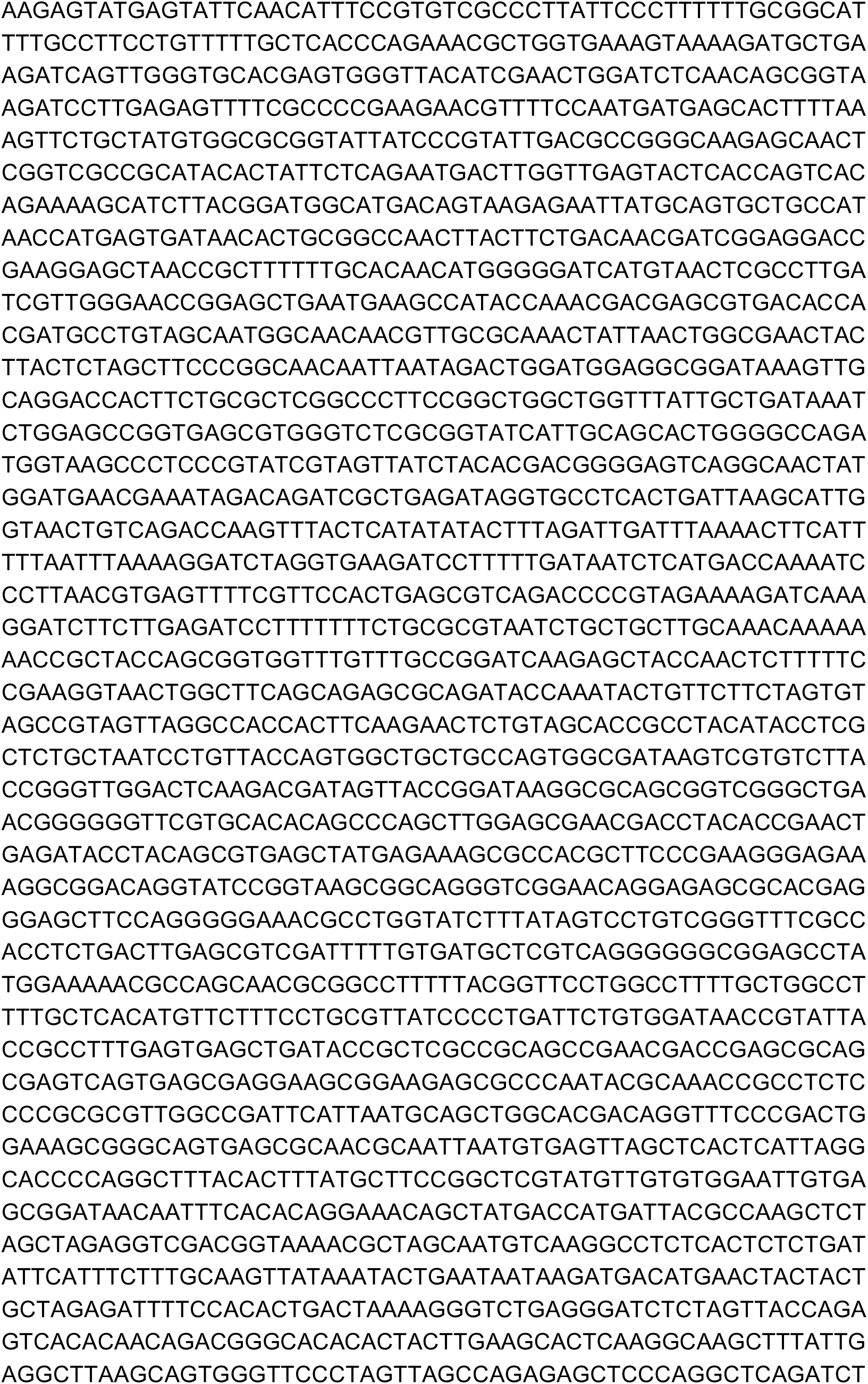

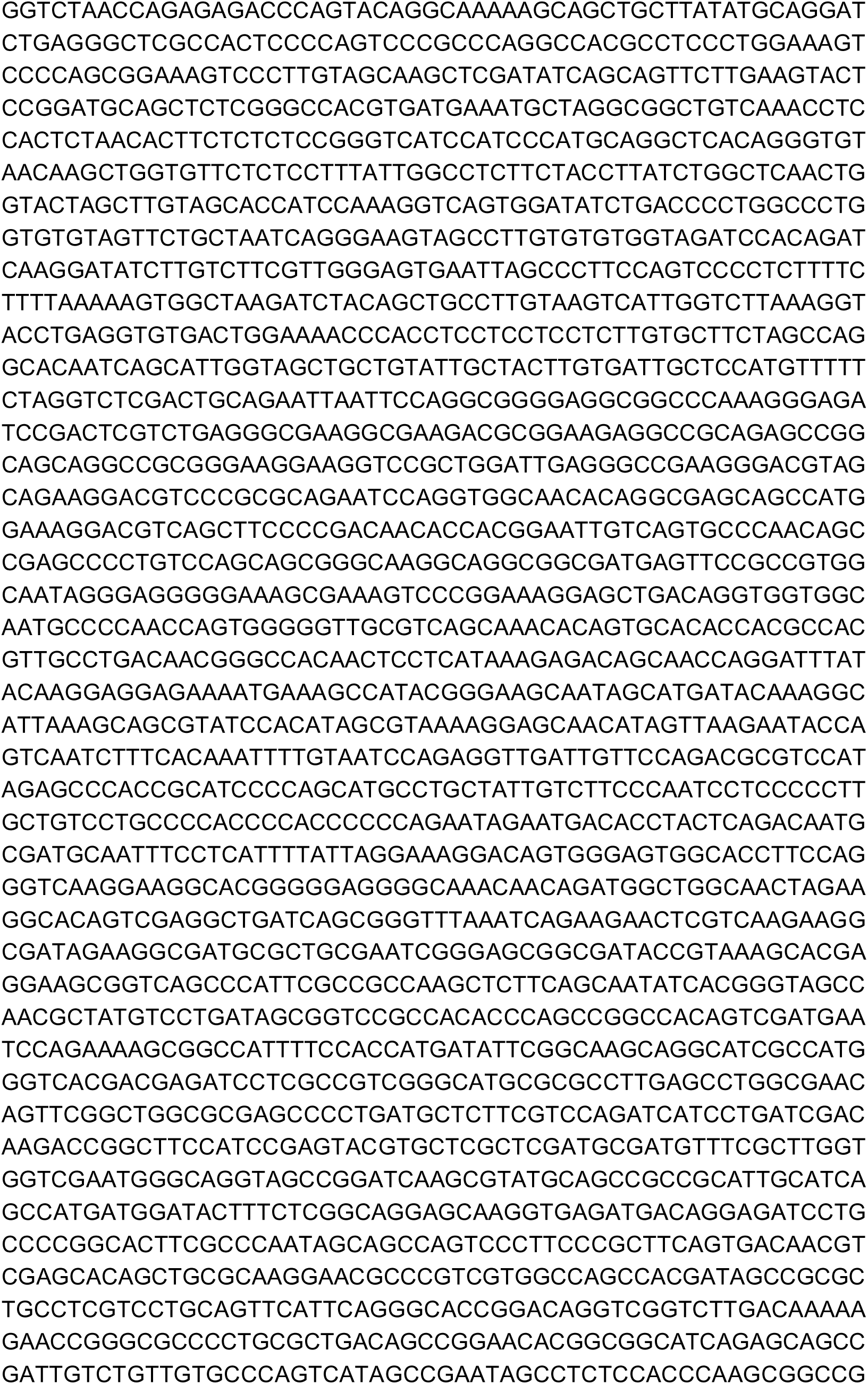

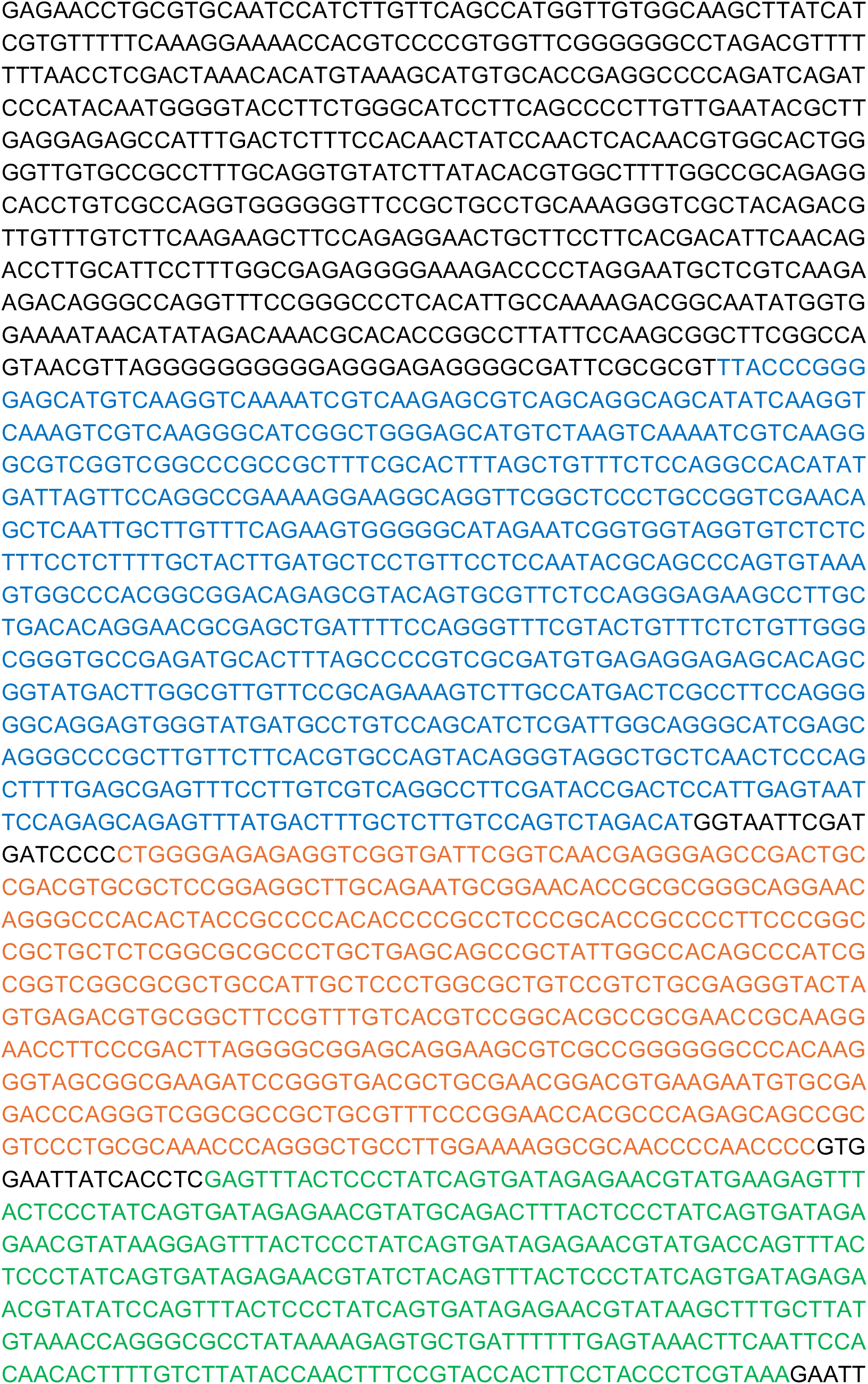

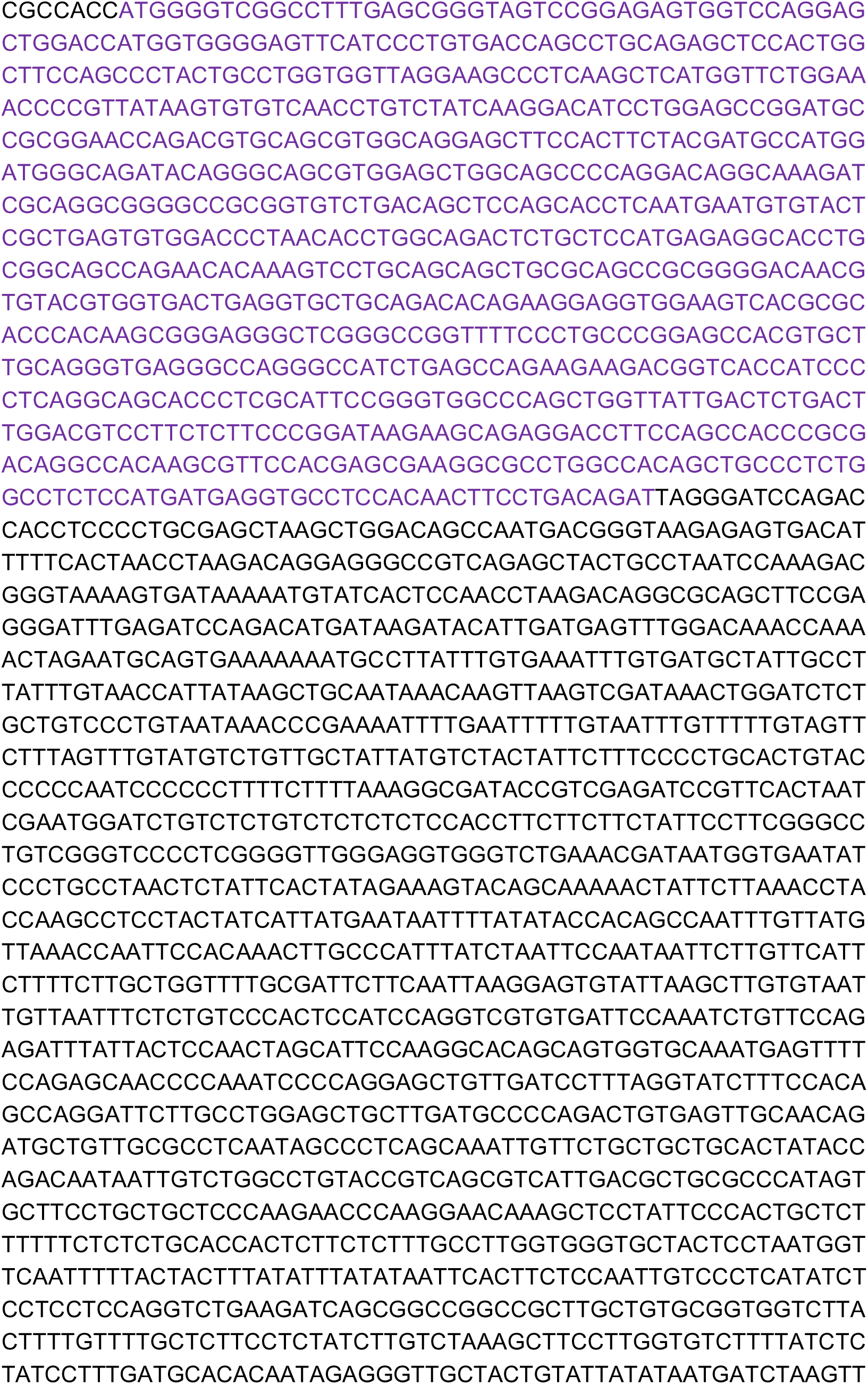

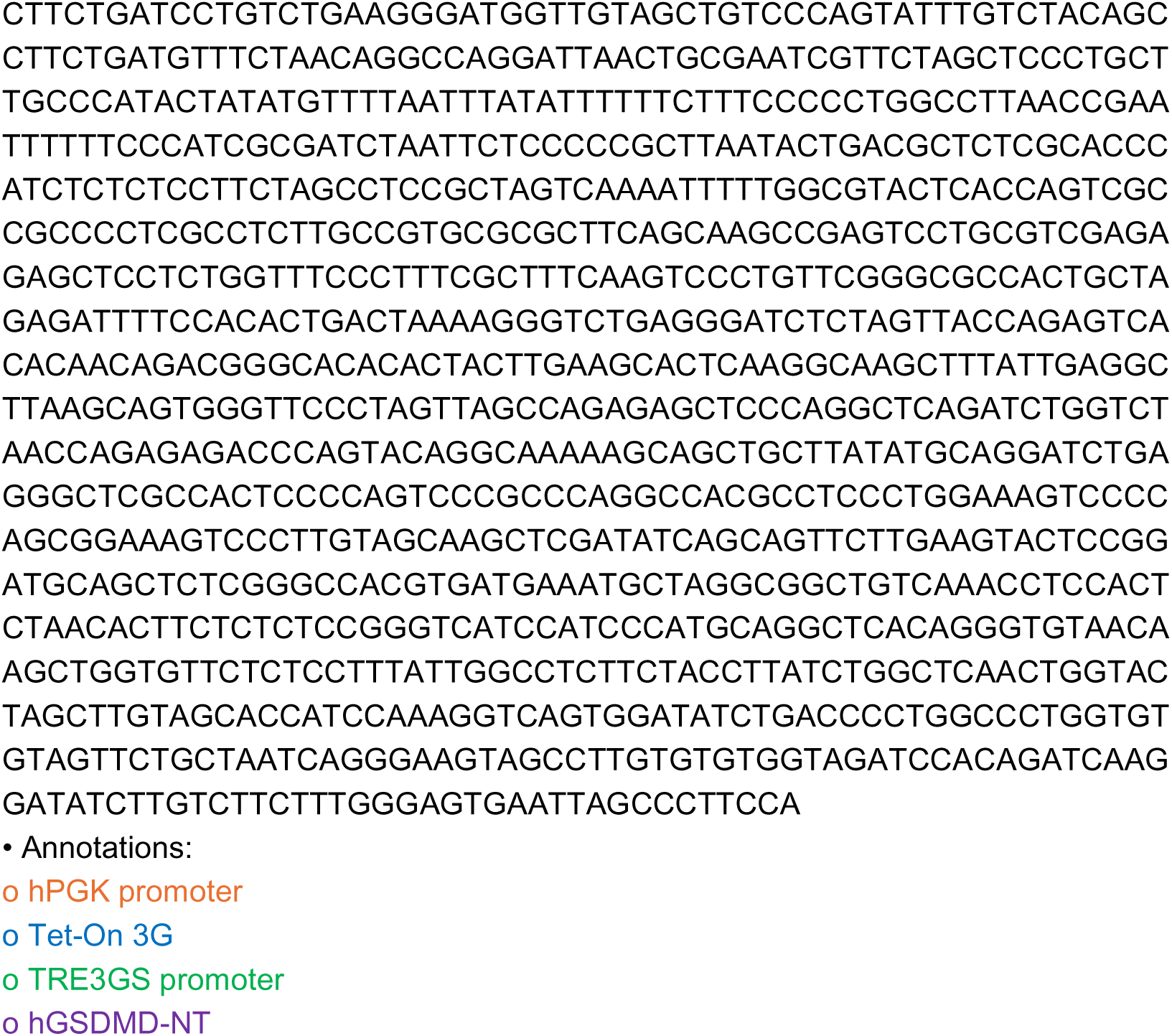

## References

1. Gao XJ, Chong LS, Kim MS et al. Programmable protein circuits in living cells. Science 2018;361: 1252–1258. doi:10.1126/science.aat5062

2. Chen Z, Kibler RD, Hunt A et al. De novo design of protein logic gates. Science 2020;368: 78–84. doi:10.1126/science.aay2790

3. Shao J, Qiu X, Zhang L et al. Multi-layered computational gene networks by engineered tristate logics. Cell 2024;187: 5064–5080.e5014. 10.1016/j.cell.2024.07.001

4. Chen Z, Linton JM, Xia S et al. A synthetic protein-level neural network in mammalian cells. Science 2024;386: 1243–1250. doi:10.1126/science.add8468

5. An B, Tang T-C, Zhang Q et al. Synthetic circuits for cell ratio control. Nature 2026;653: 587–598. 10.1038/s41586-026-10259-3

6. Gossen M and Bujard H. Tight control of gene expression in mammalian cells by tetracycline-responsive promoters. Proceedings of the National Academy of Sciences 1992;89: 5547–5551. doi:10.1073/pnas.89.12.5547

7. Gossen M, Freundlieb S, Bender G et al. Transcriptional Activation by Tetracyclines in Mammalian Cells. Science 1995;268: 1766–1769. doi:10.1126/science.7792603

8. Weinberg BH, Pham NTH, Caraballo LD et al. Large-scale design of robust genetic circuits with multiple inputs and outputs for mammalian cells. Nat Biotechnol 2017;35: 453–462. 10.1038/nbt.3805

9. Qi Lei S, Larson Matthew H, Gilbert Luke A et al. Repurposing CRISPR as an RNA-Guided Platform for Sequence-Specific Control of Gene Expression. Cell 2013;152: 1173–1183. 10.1016/j.cell.2013.02.022

10. Chavez A, Scheiman J, Vora S et al. Highly efficient Cas9-mediated transcriptional programming. Nature Methods 2015;12: 326–328. 10.1038/nmeth.3312

11. Ginhoux F, Turbant S, Gross DA et al. HLA-A*0201-restricted cytolytic responses to the rtTA transactivator dominant and cryptic epitopes compromise transgene expression induced by the tetracycline on system. Mol Ther 2004;10: 279–289. 10.1016/j.ymthe.2004.05.012

12. Chew WL, Tabebordbar M, Cheng JKW et al. A multifunctional AAV– CRISPR–Cas9 and its host response. Nature Methods 2016;13: 868–874. 10.1038/nmeth.3993

13. Charlesworth CT, Deshpande PS, Dever DP et al. Identification of preexisting adaptive immunity to Cas9 proteins in humans. Nature Medicine 2019;25: 249–254. 10.1038/s41591-018-0326-x

14. Wagner DL, Amini L, Wendering DJ et al. High prevalence of Streptococcus pyogenes Cas9-reactive T cells within the adult human population. Nature Medicine 2019;25: 242–248. 10.1038/s41591-018-0204-6

15. Dong J-Y, Fan P-D and Frizzell RA. Quantitative Analysis of the Packaging Capacity of Recombinant Adeno-Associated Virus. Human Gene Therapy 1996;7: 2101–2112. 10.1089/hum.1996.7.17-2101

16. Bentley WE, Mirjalili N, Andersen DC et al. Plasmid-encoded protein: The principal factor in the “metabolic burden” associated with recombinant bacteria. Biotechnology and Bioengineering 1990;35: 668–681. 10.1002/bit.260350704

17. Glick BR. Metabolic load and heterologous gene expression. Biotechnology Advances 1995;13: 247–261. 10.1016/0734-9750(95)00004-A

18. Hoetzel J, Wang T and Suess B. Beyond the niche - unlocking the full potential of synthetic riboswitches. Nature Communications 2025;16: 9897. 10.1038/s41467-025-65808-7

19. Vogel M, Weigand JE, Kluge B et al. A small, portable RNA device for the control of exon skipping in mammalian cells. Nucleic Acids Research 2018;46: e48–e48. 10.1093/nar/gky062

20. Finke M, Brecht D, Stifel J et al. Efficient splicing-based RNA regulators for tetracycline-inducible gene expression in human cell culture and C. elegans. Nucleic Acids Research 2021;49: e71–e71. 10.1093/nar/gkab233

21. Monteys AM, Hundley AA, Ranum PT et al. Regulated control of gene therapies by drug-induced splicing. Nature 2021;596: 291–295. 10.1038/s41586-021-03770-2

22. Fukunaga K, Dhamodharan V, Miyahira N et al. Small-Molecule Aptamer for Regulating RNA Functions in Mammalian Cells and Animals. Journal of the American Chemical Society 2023;145: 7820–7828. 10.1021/jacs.2c12332

23. Luo L, Jea JD-Y, Wang Y et al. Control of mammalian gene expression by modulation of polyA signal cleavage at 5′ UTR. Nature Biotechnology 2024;42: 1454–1466. 10.1038/s41587-023-01989-0

24. Hedwig V, Spöring M, Ottlinger J et al. Engineering oxypurinol-responsive riboswitches based on bacterial xanthine aptamers for gene expression control in mammalian cell culture. Nucleic Acids Research 2025;53. 10.1093/nar/gkae1189

25. Chen Z, Yang L, Zhang Y et al. The drug-elicitable alternative splicing module for tunable vector expression in the heart. Nature Cardiovascular Research 2025;4: 938–955. 10.1038/s44161-025-00665-7

26. Hou Q, Oleynikov M, Mei X et al. A portable poison exon for small-molecule control of mammalian gene expression. Nature Methods 2025;22: 2337– 2348. 10.1038/s41592-025-02860-7

27. Mendel M, Schwarz D, Sun T et al. Tunable gene control via RNA splicing with a clinically approved small molecule. Nature Communications 2026. 10.1038/s41467-026-73673-1

28. Chi Y, Lu X, Li S et al. A compact, versatile drug-induced splicing switch system with minimal background expression. Cell Reports Methods 2024;4. 10.1016/j.crmeth.2024.100842

29. Chow LT, Gelinas RE, Broker TR et al. An amazing sequence arrangement at the 5′ ends of adenovirus 2 messenger RNA. Cell 1977;12: 1–8. 10.1016/0092-8674(77)90180-5

30. Berget SM, Moore C and Sharp PA. Spliced segments at the 5’ terminus of adenovirus 2 late mRNA*. Proceedings of the National Academy of Sciences 1977;74: 3171–3175. doi:10.1073/pnas.74.8.3171

31. Gilbert W. Why genes in pieces? Nature 1978;271: 501–501. 10.1038/271501a0

32. Wang ET, Sandberg R, Luo S et al. Alternative isoform regulation in human tissue transcriptomes. Nature 2008;456: 470–476. 10.1038/nature07509

33. Pan Q, Shai O, Lee LJ et al. Deep surveying of alternative splicing complexity in the human transcriptome by high-throughput sequencing. Nature Genetics 2008;40: 1413–1415. 10.1038/ng.259

34. Maquat LE, Kinniburgh AJ, Rachmilewitz EA et al. Unstable β-globin mRNA in mRNA-deficient *β_0_* thalassemia. Cell 1981;27: 543–553. 10.1016/0092-8674(81)90396-2

35. Nomura Y, Kim N, Zhu B et al. Optimization of Exon-Skipping Riboswitches and Their Applications to Control Mammalian Cell Fate. ACS Synthetic Biology 2024;13: 3246–3255. 10.1021/acssynbio.4c00295

36. Xiong H, Chang Q, Ding J et al. A highly stable monomeric red fluorescent protein for advanced microscopy. Nature Methods 2025;22: 1288–1298. 10.1038/s41592-025-02676-5

37. Shaner NC, Lambert GG, Chammas A et al. A bright monomeric green fluorescent protein derived from Branchiostoma lanceolatum. Nature Methods 2013;10: 407–409. 10.1038/nmeth.2413

38. Feng H-Y, Wu X-J, Li X-H et al. Enhancing small-molecule-mediated translational control through multivalent RNA aptamers. Nucleic Acids Research 2026;54. 10.1093/nar/gkag592

39. Hedwig V, Müller E, Ketterer S et al. Engineering aptamer dimers (apdimers) for optimization of synthetic riboswitches. Nucleic Acids Research 2026;54. 10.1093/nar/gkag702

40. Tuerk C and Gold L. Systematic Evolution of Ligands by Exponential Enrichment: RNA Ligands to Bacteriophage T4 DNA Polymerase. Science 1990;249: 505–510. doi:10.1126/science.2200121

41. Ellington AD and Szostak JW. In vitro selection of RNA molecules that bind specific ligands. Nature 1990;346: 818–822. 10.1038/346818a0

42. Townshend B, Xiang JS, Manzanarez G et al. A multiplexed, automated evolution pipeline enables scalable discovery and characterization of biosensors. Nature Communications 2021;12: 1437. 10.1038/s41467-021-21716-0

43. Nandagopal N and Elowitz MB. Synthetic Biology: Integrated Gene Circuits. Science 2011;333: 1244–1248. doi:10.1126/science.1207084

44. Qin C, Xiang Y, Liu J et al. Precise programming of multigene expression stoichiometry in mammalian cells by a modular and programmable transcriptional system. Nature Communications 2023;14: 1500. 10.1038/s41467-023-37244-y

45. Sanford A, Kiriakov S and Khalil AS. A Toolkit for Precise, Multigene Control in Saccharomyces cerevisiae. ACS Synthetic Biology 2022;11: 3912–3920. 10.1021/acssynbio.2c00423

46. Subach OM, Cranfill PJ, Davidson MW et al. An Enhanced Monomeric Blue Fluorescent Protein with the High Chemical Stability of the Chromophore. PLOS ONE 2011;6: e28674. 10.1371/journal.pone.0028674

47. Takahashi S, Inamura K, Yarimizu J et al. Neurochemical and neuropharmacological characterization of ASP2905, a novel potent selective inhibitor of the potassium channel KCNH3. European Journal of Pharmacology 2017;810: 26–35. 10.1016/j.ejphar.2017.05.045

48. Mindel A, Carney O, Freris M et al. DOSAGE AND SAFETY OF LONG-TERM SUPPRESSIVE ACYCLOVIR THERAPY FOR RECURRENT GENITAL HERPES. The Lancet 1988;331: 926–928. 10.1016/S0140-6736(88)91725-4

49. Zhang J, Chen L, Zhu X et al. Coordinated regulation using small-molecule drugs enables controlled therapeutic genome editing and enhanced genomic precision in situ. Science Translational Medicine 2026;18: eadx7857. doi:10.1126/scitranslmed.adx7857

50. Kato Y. Extremely Low Leakage Expression Systems Using Dual Transcriptional-Translational Control for Toxic Protein Production. International journal of molecular sciences 2020;21: 705.

51. Bence NF, Sampat RM and Kopito RR. Impairment of the Ubiquitin-Proteasome System by Protein Aggregation. Science 2001;292: 1552–1555. doi:10.1126/science.292.5521.1552

52. Gilon T, Chomsky O and Kulka RG. Degradation signals for ubiquitin system proteolysis in Saccharomyces cerevisiae. Embo j 1998;17: 2759–2766. 10.1093/emboj/17.10.2759

53. Rogers S, Wells R and Rechsteiner M. Amino Acid Sequences Common to Rapidly Degraded Proteins: The PEST Hypothesis. Science 1986;234: 364–368. doi:10.1126/science.2876518

54. Banaszynski LA, Chen L-c, Maynard-Smith LA et al. A Rapid, Reversible, and Tunable Method to Regulate Protein Function in Living Cells Using Synthetic Small Molecules. Cell 2006;126: 995–1004. 10.1016/j.cell.2006.07.025

55. Iwamoto M, Björklund T, Lundberg C et al. A General Chemical Method to Regulate Protein Stability in the Mammalian Central Nervous System. Chemistry & Biology 2010;17: 981–988. 10.1016/j.chembiol.2010.07.009

56. Shi J, Zhao Y, Wang K et al. Cleavage of GSDMD by inflammatory caspases determines pyroptotic cell death. Nature 2015;526: 660–665. 10.1038/nature15514

57. Liu X, Zhang Z, Ruan J et al. Inflammasome-activated gasdermin D causes pyroptosis by forming membrane pores. Nature 2016;535: 153–158. 10.1038/nature18629

58. Ding J, Wang K, Liu W et al. Pore-forming activity and structural autoinhibition of the gasdermin family. Nature 2016;535: 111–116. 10.1038/nature18590

59. Xia S, Zhang Z, Magupalli VG et al. Gasdermin D pore structure reveals preferential release of mature interleukin-1. Nature 2021;593: 607–611. 10.1038/s41586-021-03478-3

60. Liu X, Xia S, Zhang Z et al. Channelling inflammation: gasdermins in physiology and disease. Nature Reviews Drug Discovery 2021;20: 384–405. 10.1038/s41573-021-00154-z

61. Lu Y, He W, Huang X et al. Strategies to package recombinant Adeno-Associated Virus expressing the N-terminal gasdermin domain for tumor treatment. Nature Communications 2021;12: 7155. 10.1038/s41467-021-27407-0

62. He J, Zheng P, Chen Y et al. A new personalized vaccine strategy based on inducing the pyroptosis of tumor cells in vivo by transgenic expression of a truncated GSDMD N-terminus. Frontiers in immunology 2022;Volume 13 - 2022. 10.3389/fimmu.2022.991857

63. Evavold CL, Hafner-Bratkovič I, Devant P et al. Control of gasdermin D oligomerization and pyroptosis by the Ragulator-Rag-mTORC1 pathway. Cell 2021;184: 4495–4511.e4419. 10.1016/j.cell.2021.06.028

64. Venezia O, Kane H, Du G et al. Functional chimeric mRNAs encode proteins in mammalian immunity. Nature 2026. 10.1038/s41586-026-10982-x

65. Zhang Y, Yang Y, Liu Z et al. The DreAM-plus integrative RNA switch enhances transient AAV expression and reduces side effects of gene editing. Molecular Therapy 2026. 10.1016/j.ymthe.2026.08.006

66. Orehek S, Ramuta TŽ, Lainšček D et al. Cytokine-armed pyroptosis induces antitumor immunity against diverse types of tumors. Nature Communications 2024;15: 10801. 10.1038/s41467-024-55083-3

67. Takahashi K and Yamanaka S. Induction of pluripotent stem cells from mouse embryonic and adult fibroblast cultures by defined factors. Cell 2006;126: 663–676. 10.1016/j.cell.2006.07.024

68. Takahashi K, Tanabe K, Ohnuki M et al. Induction of Pluripotent Stem Cells from Adult Human Fibroblasts by Defined Factors. Cell 2007;131: 861–872. 10.1016/j.cell.2007.11.019

69. Hochedlinger K, Yamada Y, Beard C et al. Ectopic Expression of Oct-4 Blocks Progenitor-Cell Differentiation and Causes Dysplasia in Epithelial Tissues. Cell 2005;121: 465–477. 10.1016/j.cell.2005.02.018

70. Nakagawa M, Koyanagi M, Tanabe K et al. Generation of induced pluripotent stem cells without Myc from mouse and human fibroblasts. Nature Biotechnology 2008;26: 101–106. 10.1038/nbt1374

